# Linear and partially linear models of behavioral trait variation using admixture regression

**DOI:** 10.1101/2021.05.14.444173

**Authors:** Gregory Connor, Gerard R. Fuerst

## Abstract

Admixture regression methodology exploits the natural experiment of random mating between individuals with different ancestral backgrounds to infer the environmental and genetic components to trait variation across racial and ethnic groups. This paper provides a statistical framework for admixture regression based on the linear polygenic index model and applies it to neuropsychological performance data from the Adolescent Brain Cognitive Development (ABCD) database. We develop and apply a new test of the differential impact of multi-racial identities on trait variation, an orthogonalization procedure for added explanatory variables, and a partially linear semiparametric functional form. We find a statistically significant genetic component to neuropsychological performance differences across racial identities, and find some possible evidence of nonlinearity in the link between admixture and neuropsychological performance scores in the ABCD data.

## 1. Introduction

Racial/ethnic group identities such as Black, White, Hispanic, Native American, East Asian and South Asian show empirically strong linkages to medical and behavioral traits such as obesity (Wang et al. 2007), type 2 diabetes (Cheng et al. 2013), hypertension (Lackland 2014), asthma (Choudry et al. 2006), neuropsychological performance (Llibre-Guerra et al. 2018), smoking behaviors (Choquet et al. 2021), and sleep disorders (Halder et al 2015). An important research question is to what degree any such observed trait variation arises from differences in the typical diets, cultural practices and other environmental particularities of the racial/ethnic groups, or from similarity in genetic pools within each group traceable to shared geographic ancestry. Many diverse national populations descend demographically from isolated continental groups within a few hundred years. Modern genetic technology can measure with high accuracy the proportion of an individual’s ancestry associated with these continental groups. Also, in many culturally diverse nations, most individuals can reliably self-identify as members of one or more racial or ethnic groups. Admixture regression leverages these two data sources, self-identified race or ethnicity (SIRE) and genetically-measured admixture proportions, to decompose trait variation correspondingly. Admixture regression has been widely applied to medical and behavioral traits including asthma (Salari et al. 2005), body mass index (Klimentidis et al. 2009), type 2 diabetes (Cheng et al. 2013), blood pressure (Klimentidis et al. 2012), neuropsychological performance (Lasker et al. 2019), and sleep depth (Halder et al. 2015). It has particular value in the case of complex behavioral traits where reliably identifying genetic loci associated with trait variation is beyond the current reach of science. Admixture mapping is a more technically challenging methodology, often used in conjuction with admixture regression, which uses ancestral population trait differences to attempt to identify genetic loci associated with a trait. This paper focusses exclusively on admixture regression.

This paper first develops a simple statistical framework for admixture regression of behavioral traits by linking it to the linear polygenic index model from behavioral genetics; this framework clarifies the key assumptions that are implicit in this simple and powerful statistical technique. The paper then extends the admixture regression methodology in several ways. We provide a new test statistic for identifying whether a given multi-racial identity differs in its trait impact from the average impact of its component single-SIRE categories. We examine the role of additional explanatory variables in the admixture regression and their interpretation with and without orthogonalization with respect to the core explanatory variables. We generalize the linear admixture regression specification to a partially linear semiparametric form.

We apply our methodology to neuropsychological performance data from the Adolescent Brain Cognitive Development database. Neuropsychological performance is one of the most complex traits to which admixture regression analysis has been applied. Our findings corroborate existing evidence that genetic variation plays a statistically significant role in explaining neuropsychological performance differences across racial identities (Lasker et al. 2019). Using our new test statistic, we find that some multi-racial categories have identifiably distinct impact relative to their component categories. We find that orthogonalization of additional variables can substantially change the interpretation of the core coefficients in the admixture regression. Our analysis also indicates (although not conclusively) that a partially linear semiparametric specification potentially adds empirical value.

## 2. A statistical framework for admixture regression tests of trait variation

### 2.1. Variable definitions

We assume that the database consists of n individuals indexed by *i* = 1,…, *n* who have each self-identified their racial or ethnic group membership(s), recorded a score on a behavioral trait, *s_i_*, and provided a personal DNA sample. The *k* racial or ethnic group self-identification choices are captured by a matrix of zero-one dummy variables SIRE*_ij_, i* = 1,…, *n*; *j* = 1,…, *k*. We assume that every individual has self-identified as belonging to at least one and possibly more of the *k* groups.

We assume that a set of *m* geographic ancestries covered in the study have been chosen, such as African, European, Amerindian, South Asian, and East Asian, indexed by *h* = 1,…, *m*. The genotyped DNA samples are carefully decomposed into admixture proportions of geographic ancestry, as discussed in Section 4 below. For each individual the ancestry proportions across the chosen geographic ancestries sum to one. This gives a matrix of ancestry proportions *A_ih_, i* = 1,…, *n*; *h* = 1,…, *m* with 0 ≤ *A_ih_* ≤ 1 for all *i, h* and 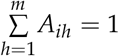 for each *i*.

In most applications of admixture regression, individuals’ racial or ethnic group identities will have statistical relationships with individuals’ genetically identified geographic ancestries and also with the observed trait *s_i_*. The objective of admixture regression is to decompose trait variation into linear components due to genetic ancestries and linear components due to racial/ethnic group related effects.

### 2.2. Ancestry proportions as a statistical proxy for ancestry-linked genetic trait variation

Admixture regression is an indirect method of analyzing group-related trait variation. In this subsection we provide a foundation for admixture regression by considering a more direct, but empirically much more challenging, alternative approach based on a linear polygenic index model. We show that the admixture regression model can be viewed as a statistically feasible simplification of this linear polygenic index model, in which proportional ancestries serve as statistical proxies for ancestry-related genetic differences.

The human genetic code contains a very large number of genetic variants (the allelles on the genome which vary between individuals) called single nucleotide polymorphisms or SNPs. Consider hypothetically a complete list of all genetic variants with any impact on variation in the observed trait. Assign a value of 0, 1 or 2 to each SNP for individual *i* depending upon the number of minor allelles for that SNP. Let SNP*_iz_ i* = 1,…, *n*; denote the number of minor allelles on the *z^th^* SNP of the *i^th^* individual in the sample.

The biochemical process linking human genetic variation to behavioral trait variation is unimaginably complex, and scientific understanding of the full biochemical process is very limited. Genome-wide association studies (GWAS) have made slow but steady progress in statistically modeling these linkages, although precise biochemical linkages are beyond the contemporary scientific frontier for most behavioral traits. A standard, admittedly highly simplified, model of the gene variation - trait variation nexus is the linear polygenic index model, in which the genetic component of a trait is a simple linear function of a relevant subset of the individual’s genetic variants. The linear polygenic index model has been applied to a wide range of medical and behavioral traits including body mass index (Yengo et al. 2018), neuroticism (Nagel et al. 2018), depression susceptibility (Wray et al. 2018), suicidal ideation (Mullins et al. 2014), schizophrenia (Mistry et al. 2018), educational attainment (Lee et al. 2018), neuropsychological test performance (Savage et al. 2018), and risk-taking (Clifton et al. 2018). The linear admixture regression model can be derived elegantly by invoking this standard linear polygenic index model, and hence we impose it, in order to provide a statistical underpinning for our admixture regression model.

Let *p_i_* denote the genetic potential of individual *i* regarding the observable trait *s_i_*. We assume that *p_i_* is a linear function of a large number of genetic variants SNP*_iz_* with associated linear coefficients *β_z_* and constant term *c*_1_:

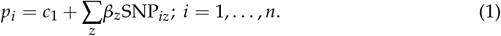

The key difference in the admixture regression methodology compared to GWAS is that there is no attempt to estimate the linear polygenic index (1). Rather, admixture regression uses the natural experiment of subpopulation mixing to infer differences in the conditional expected value of (1) arising from differences in the frequency distribution of genetic variants across ancestries. The assumed linearity of the polygenic index model (1), together with an assumption of random mating across ancestral populations, allows us to derive a linear regression model using admixture as a statistical proxy variable for the conditional expected value of *p_i_*.

The frequency distributions of many SNPs depend notably upon geographic ancestries. Consider a hypothetical individual with single-origin ancestry *h*, that is, an individual with *A_h_* = 1. Note that this also implies that *A_h′_* = 0 for all *h′* ≠ *h* since the ancestral proportions are non-negative and sum to one. Consider the expected value of *p* conditional on an individual having this single-origin ancestry. The expectation of 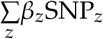 using a single-origin frequency distribution for each SNP*_z_* defines the average genetic trait potential of a single-origin ancestry:

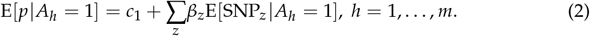

In admixture regression there is no attempt to measure (2) directly, but instead differences between (2) across *h* = 1,…, *m* will be inferred indirectly using regression methods.

A key assumption of the admixture regression model is that admixture arises from recent random mating between the previously geographically-isolated ancestral groups. Assuming recent random mating between ancestral lines, it follows from the fundamental processes of sexual reproduction that the expected value of any SNP for an admixed individual is the convex combination of the single-origin expected values, with linear coefficients equal to the individual’s admixture proportions. (The relationship between the multivariate distributions of the SNPs is more complicated, but the multivariate distributions do not impact the expected trait given the linear polygenic index assumption.) We use a subscript. to denote the vector created from the *i^th^* row of a matrix. We assume that mating across geographic ancestries is recent and random, and therefore in particular that the univariate frequency distribution of each SNP for any individual is the convex combination of the single-origin frequency distributions:

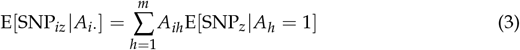

The linearity of genetic potential in the SNPs (1) and the random mixing assumption (3) imply that expected genetic potential of an admixed individual is a convex combination of the individual’s admixture proportions. Taking the expectation of (1) using (2) and (3) the conditional expected value of genetic potential for an individual with admixture proportions *A_i._* is the convex combination of the unobserved values E[*p*| *A_h_* = 1] with observed linear coefficients *A_ih_*:

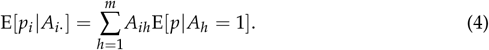

Equation (4) is a fundamental identification condition for the admixture regression methodology. As we discuss below, it allows differences between the single-origin expected values of genetic potential, E[*p*| *A_h_* = 1], *h* = 1,…, *m*, to be inferred by regression methods.

### 2.3. Adjusting for ancestry-related environmental influences on the trait

Define the environmental component of the trait, *e_i_*, as the observed trait minus genetic potential:

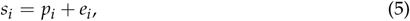

where *e_i_* is defined as all trait variation not captured by *p_i_*. Equation (5) is only definitional; later we will impose various conditions on *e_i_* to enable statistical identification of the model. Define 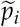 as the genetic component of the trait for each *i* which is not explained by ancestry proportions:

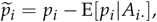

by simple substitution into (5) this gives:

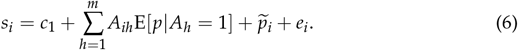

Recall that 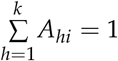, for all *i*, so that one term in (6) is redundant for the purposes of creating a regression model. Substitute 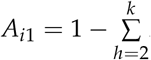 into (6) to get:

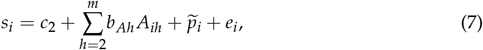

where *b_Ah_* = E[*p*|*A_h_* = 1] – E[*p*|*A*_1_ = 1]; *h* = 2,…, *m*, and *c*_2_ = *c*_1_ + E[*p*|*A*_1_ = 1].

Equation (7) is not well-specified as a regression model since the error term 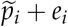 will not be mean zero conditional on *A_i._*, due to racial and ethnic group-related effects in *e_i_*. In order to transform (7) into a regression model it is necessary to add explanatory terms to the regression model to remove the expected value of *e_i_* conditional on *A_i._*. This is accomplished by assuming that the differences in *e_i_* conditional on *A_i._* are dependent on the group self-identification choices, but otherwise not dependent upon admixture proportions.

For expositional simplicity, in this subsection we assume that every individual included in the sample has self-identified as belonging to exactly one from the pre-specified set of *k* racial or ethnic groups, so that 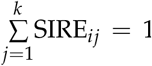 for all *i*. In this case, the *n* × *k* matrix of racial/ethnic group explanatory variables used in the admixture regression, denoted *G*, is simply set equal to the SIRE matrix: *G_ij_* = SIRE*_ij_* for *i* = 1,…, *n*; *j* = 1,…, *k*. Multi-racial individuals (those who have self-identified as belonging to two or more groups) will be introduced into the analysis in the next subsection.

We assume that after adjusting for the influence of the group identifiers *G_ij_*, the remaining error term in (7) is independent of the ancestry proportions:

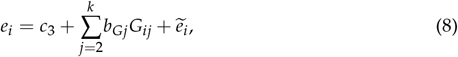

where *b_Gh_* captures the environmental component associated with membership in group *h* relative to the reference group *h* = 1, and 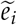 is assumed to be independent of *A_i._*, *G_i._* and 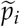, and *c*_3_ is a constant term. Combining (7) and (8) produces the key linear admixture regression specification:

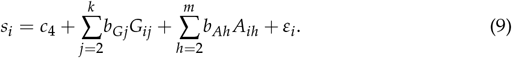

where 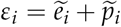 and *c*_4_ is a constant term. Note that *ε_i_* has zero mean and variance 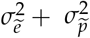 and is independent of *A_i._* and *G_i._*. Equation (9) is a well-specified linear regression model.

In many applications, the analyst also has information on the sampling substructure of the data, such as its division into site-specific subsamples. In this case, a linear mixed effects model can be used for estimating (9) rather than ordinary least squares. This involves partially decomposing the residual term *ε_i_* in (9) into linear random effects components linked to data collection site identifiers and/or other subsample identifiers, see Heeringa and Berglund (2020).

### 2.4. Adding multi-racial individuals to the regression

Recall that SIRE is the *n* × *k* matrix of race/ethnicity self-identifications. A key assumption of the admixture regression technique is that the environmental influences associated with racial/ethnic group membership are captured by these group membership self-identification choices. Many individuals self-identify as belonging to two or more racial or ethnic groups and the group variables used in the regression must be adapted to this reality. In the context of our statistical framework, there are essentially three approaches: evenly splitting the individual’s affiliation across their chosen groups, creating a new group for one or more particular multi-racial combinations, or deleting particular multi-racial observations where neither of the other two approaches seem appropriate.

We now allow that some individuals choose more than one category, so that 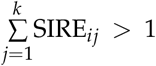 for some *i*. The simplest regression specification in this case is to assume that the group environment faced by a multi-racial individual is the average of the component group environments:

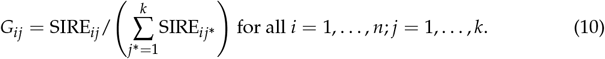

Although (10) is a reasonable specification, it is restrictive. It is possible to replace (10) with a more general specification at some loss of parsimony. Suppose that we are concerned about imposing the restrictive condition (10) for some common multi-racial choice (such as, for example, Black-White biracial in a US dataset). Let *V*_1_ denote a *k*–vector with ones for the included race/ethnicity groups in this particular multi-racial combination and zeros elsewhere. We can supplement (10) by adding a *k* + 1*^st^* group and using a different rule for this subset of multi-racials:

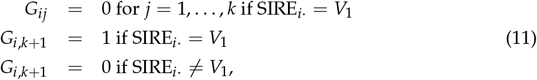

where SIRE*_i._* = *V*_1_ denotes vector equality between these two *k*–vectors. There are now *k* + 1 groups: the originally specified SIRE groups and a new group for the selected multiracial combination. *G* becomes a *n* × (*k* +1) matrix, and the regression (9) described in the previous subsection applies exactly as before but with one extra dimension to *G*. Any small number of defined multi-racial groups can be appended in this way. The only change to the regression methodology is that *G* becomes a *n* × *k*^*^ matrix (with an associated increase in the set of estimated parameters) where *k*^*^ – *k* is the number of multiracial combinations added as new categories.

It is not feasible to use rule (11) for all race/ethnicity choice combinations due to lack of parsimony; there are 2*^k^* – *k* potential multi-racial combinations and each one added requires an additional parameter in the regression. It can only be used for the common multi-racial choices where there is sufficient data of that combination in the sample. For all others, it is necessary to stick with the restrictive assumption (10) or drop the observations from the sample. This will be illustrated in the empirical application in Section 5.

Once a regression model is estimated using (11), it is possible to test the accuracy of restrictive assumption (10) for that multi-racial group. The restrictive assumption implicit in (10) requires that the average of the coefficients of the components equals the added-group coefficient in the unrestricted model:

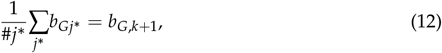

where #*j*^*^ denotes the number of components in the multiracial category (typically either two or three) and the sum runs over these element only. This is a linear restriction on the vector of coefficients, or multiple linear restrictions for *k*^*^ – *k* greater than one, which can be tested with a t-test (for each group coefficient singly) or a Wald test for all them, as detailed below.

Let 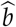 denote the (*m* + *k*^*^ – 1)–vector of all the coefficients in the admixture regression (9):

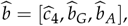

and let 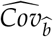 denote the estimated (*m* + *k*^*^ – 1) × (*m* + *k*^*^ – 1)–covariance matrix of these estimates.

First consider the case *k*^*^ – *k* = 1. Let *R* denote the (*m* + *k*^*^ – 1)–vector expressing restriction (12) imposed on b. For example, if the group combination consists of individuals who choose all three of the first, second, and third SIRE categories (recalling that the first SIRE category is not included in the regression) the restriction vector is:

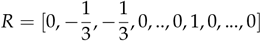

where the 1 is element *k*^*^ in the vector. Any other restriction of type (12) is easily stated in this way. In the case of one group, this gives rise to a standard t-test of the one coefficient restriction, and in particular:

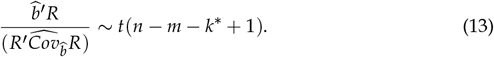

For the case *k*^*^ – *k* > 1 it is possible to test each multi-racial group equality individually as above using (13) or perform a joint Wald test on all of them. Let *R* denote the (*m* + *k*^*^ – 1) × (*k*^*^ – *k*)– matrix of all the linear restrictions, giving the standard Wald test:

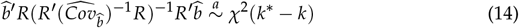

where 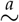 denotes the approximate distribution for large n. In the case of estimation by linear mixed effects modeling, both test statistics (13) and (14) are large–*n* asymptotic distributions rather than exact finite-sample distributions, but they remain valid tests.

## 3. Extensions of the linear admixture regression model

### 3.1. Additional explanatory variables with and without orthogonalization

It is straightforward to include additional explanatory variables in the admixture regression model. Let *x*_*i*1_, *x*_*i*2_,…, *x_il_* denote a set of explanatory variables that help to linearly explain the trait along with the ancestry proportions and group identities. We modify specification (9) to include these:

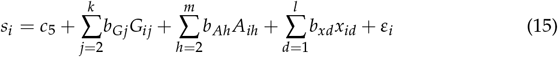

and keep all the other assumptions as before. The estimation theory for (15) is essentially identical to that of (9) as discussed above.

In some cases, the admixture regression model with additional explanatory variables (15) can be made more useful and informative by orthogonal rotation of one or more of the explanatory variables, in order to aggregate the full effects of proportional ancestries and group identities into their associated coefficients. To understand why such an orthogonal rotation might be useful, consider the hypothetical case of an admixture regression model of Body Mass Index (BMI) in which waist measurement is one of the explanatory variables. Waist measurement has such strong explanatory power for BMI that its presence in an admixture regression model like (15) will diminish the direct explanatory power of proportional ancestries and group identities; their total impact will be partly hidden within the waist measurement variable. This can be remedied by orthogonalizing the waist measurement variable with respect to the proportional ancestry and group identity variables before estimating the admixture regression, as explained next.

Suppose that variable *x*_1_ in (15) has strong explanatory power for *s* and substantial correlation with proportional ancestry and/or group identity variables, and therefore the analyst wishes to orthogonalize it with respect to *G_ij_* and *A_ih_*, *j* = 2,…, *k*; *h* = 2,…, *m*. In a first step, the analyst can perform a simple least square regression decomposition of *x*_1_ into the component linearly explained by these variables, and the residual, orthogonal component 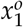:

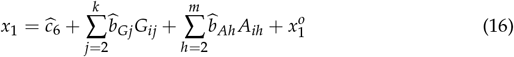

Since all the explanatory variables are deterministic (that is, conditionally fixed variables rather than random variables in the regression model), this orthogonalization step is interpreted as a matrix transformation of fixed vectors and does not alter any statistical assumptions of the main regression model. It merely serves to linearly rotate the deterministic explanatory variables used in the actual, second-stage, admixture regression. Replacing *x*_1_ with 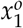 in (15) changes the interpretation of the coefficients 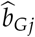 and 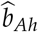, *j* = 2,…, *k*; *h* = 2,…, *m* since they now include the *G_ij_* and *A_ih_* related explanatory power from *x*_1_. An illustrative example will be provided in Section 5 below.

### 3.2. A semiparametric extension of the admixture regression model

The linear dependence of the trait on admixture proportions in our regression model is in part an artifact of the assumption of a linear polygenic index (1). It is possible to weaken this linearity assumption using nonparametric regression methods. We replace the restrictive assumption of a linear polygenic index (1) with a very general description of genetic potential as a function of the full vector of genetic variants:

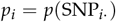

and instead of linearity as in (1) only require smoothness conditions on the conditional expectation of *p*(·) as a function of the ancestral proportions vector, as delineated below.

As in earlier subsections, we consider *p_i_* as a stochastic function of the ancestral proportions vector *A_i._*, but now without imposing the strict linearity (4) arising from the linear polygenic index assumption:

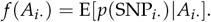

Define the unexplained component of *p_i_* as before:

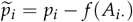

and we assume that 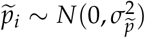 and independent of *A_i._* and *G_i._*. We impose the same assumptions on *e_i_* as in Section 2, giving:

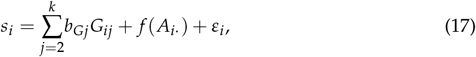

where 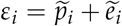 is assumed to be normally distributed with mean zero and variance 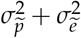 and independent of *A_i._* and *G_i._*. This equation (17) is a partially linear nonparametric regression model, see, e.g. Li and Racine (2007). This model can be consistently estimated using the three-step procedure of Robinson (1988). We will impose Condition 7.1 from Li and Racine (2007) in order to justify this procedure within our framework (see the Technical Appendix for details).

For the case *m* > 2 the general specification (17) suffers from the curse of dimensionality and is unlikely to be estimable on moderate-sized datasets. A more restrictive specification is needed to give the model sufficient parsimony for estimation. One reasonable specification choice is to restrict the nonlinearity in the impact of ancestries on the trait to a single ancestral category, which we assume is ancestry category 2, giving rise to the specification:

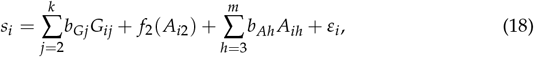

and we will now rely on this more restrictive specification throughout the remainder of this subsection.

We assume that the unconditional density Pr(*A*_2_) is continuous and strictly positive everywhere on the [0, 1] interval. Let 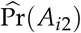 denote the nonparametrically estimated unconditional density of *A*_*i*2_:

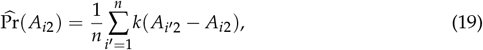

where *k*(•) is a kernel weighting function. In our empirical application in Section 5 we use the Gaussian kernel weighting function.

In the first step of the Robinson procedure, the conditional means of the dependent variable and linear-component explanatory variables are estimated nonparametrically as functions of the nonparametric-component explanatory variable, *A*_*i*2_:

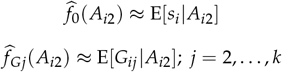

and

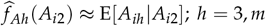

that is:

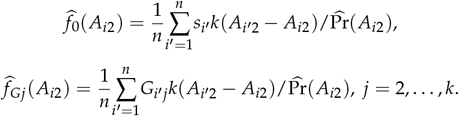

and

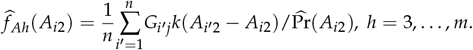

In the second step, the linear parameters of the model (17) are estimated by ordinary least squares, replacing the dependent variable and linear-component explanatory variables with the deviations from their conditional mean functions:

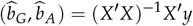

where

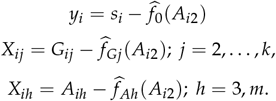

Note that 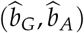 is a (*k* + *m* – 3)–vector and *X* is a *n* × (*k* + *m* – 3)–matrix where the index first runs from 2 to *k* over *j* and then from 3 to *m* over *h*.

In the third step, the nonparametric component of the model is estimated by subtracting the predicted linear component from both sides of (17) and then applying standard nonparametric regression:

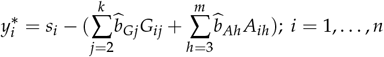

and then:

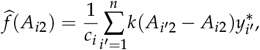

where 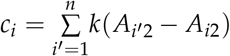.

The partially linear nonparametric approach to admixture regression is more empirically challenging than the linear specification. Proper implementation of the technique involves a tradeoff between parsimony, the generality of the specification used, and the distributional features of the available data. An example of (18) will be estimated in Section 5 below.

## 4. Materials and Methods

The Adolescent Brain and Cognitive Development (ABCD) study is the largest long-term study of brain development and child health in the United States, testing 11,000 children ages 9-10 at 21 testing sites; see Karcher and Barch (2021) for an overview. Our sample consists of age and gender-adjusted scores and genotyped DNA samples of the 9972 children in the ABCD study who met our sample selection criteria, along with questionaire responses of their parent(s)/guardian(s).

The dependent variable in our model is the composite neuropsychological performance score based on the NIH Toolbox (NIHTBX) neurocognitive battery provided in the ABCD database; this consists of tasks measuring attention, episodic memory, language abilities, executive function, processing speed, and working memory. Age-corrected composite scores, based on the seven tasks, were provided by ABCD. We regressed out sex from these age-corrected composite scores. The residuals were then standardized and serve as the dependent variable in our empirical analysis in Section 5 below.

Our core explanatory variables are seven SIRE variables, White, Black, Hispanic, Native American, East Asian, South Asian, and Other (and including multiple SIRE choices from among these) and five genetic ancestry proportions of European, African, Amerindian, East Asian and South Asian background obtained from the genotyped DNA samples. Children whose parent(s)/guardian(s) identified the child as belonging to Pacific Islander racial groups were excluded from our analyses owing to a lack of corresponding ancestry category in our chosen five categories. The ABCD Version 3 database provides 516,598 genotyped SNP variants for each individual’s DNA sample. After quality control, filtering, and pruning we were left with 99,642 SNP variants to determine the five ancestry proportions, employing the Admixture 1.3 software package (Alexander et al. 2015). We use the Pritchard et al. (2000) population structure algorithm, as implemented in R routine *Structure*, to estimate the ancestry proportions of each individual in the sample.

The ABCD database includes site identifiers for the data collection sites (in most cases, elementary schools) and family household identifiers (identifying multiple individuals in the sample from the same family household, usually twins). As recommended by Heeringa and Berglund (2020) for regression analysis using the ABCD database, we include random effects in our regression models to account for any site-specific and family-specific error correlation. We use the linear regression mixed effects estimate routine *lmer* from the R programming language library, see Bates et al. (2015). The one exception is regression Model 3 (see below) in which we estimate a semiparametric partially linear model. In that case, we use the the *npplr* routine in the R programming language subroutine library *NP* written and maintained by Hayfield and Racine (2020), and do not correct for site-specific and family-specific error correlation.

See the Supplemental Materials for more detailed description of the ABCD database, our sample selection procedure, and the construction of the variables that we use.

## 5. Results

In this section, we apply our admixture regression techniques to neuropsychological performance using the ABCD database. Table 1 shows means and standard deviations of the regression dependent variable on data subsets sorted by SIRE choice. On the full sample, by construction, the dependent variable has a mean of zero and standard deviation of one. There is considerable dispersion in the subsample means sorted by SIRE; for example, the means differ by 1.02 standard deviations (using the full-sample standard deviation for simplicity) between two of the largest SIRE categories shown, White-only SIRE and Black-only SIRE. The considerable variation in means for SIRE-based subsamples provides an initial justification for performing admixture regression analysis. This is a table of descriptive statistics; the standard errors shown are not appropriate for formal hypothesis testing since there is no adjustment for potential site-linked and family-linked correlations, particularly relevant in the case of the smaller subsample categories.

**Table 1.** Neuropsychological Performance Scores Sorted by Self-identified Race or Ethnicity (SIRE)

| Within-Category Means and Standard Deviations | Individuals with Single-SIRE Identities |  |  |  |  |  | All Hispanic |
| --- | --- | --- | --- | --- | --- | --- | --- |
|  | White Only | Black Only | Native American Only | East Asian Only | South Asian Only | Other Only |  |
| Mean | 0.25 | -0.77 | -0.42 | 0.57 | 0.45 | -.22 | -.23 |
| Standard Deviation | 0.92 | 0.87 | 0.79 | 1.02 | 1.02 | 1.12 | 0.96 |
| Number of Observations | 5593 | 1434 | 31 | 107 | 43 | 97 | 1869 |
|  | Individuals with Selected Multiple-SIRE Identities |  |  |  |  |  |  |
|  | Black-White | Hispanic-White | Native American - White | East Asian - White | South Asian - White | Hispanic - Black | Hispanic - Other |
| Mean | -0.13 | -0.19 | 0.01 | 0.58 | 0.83 | -0.34 | -0.45 |
| Standard Deviation | 0.95 | 0.96 | 0.86 | 0.98 | 0.84 | 0.93 | 0.92 |
| Number of Observations | 302 | 1171 | 131 | 249 | 40 | 84 | 411 |

Notes to Table: The table shows means and standard deviations of neuropsychological performance scores for individuals sorted into categories by self-identified race and ethnicity (SIRE). The categories used are: individuals who choose only one of the six race categories (White, Black, Native American, East Asian, South Asian, and Other), all individuals selecting Hispanic ethnicity along with any of the six race categories, and individuals choosing the seven most common two-SIRE choices. By construction the mean and standard deviation of the full sample are zero and one; there are 9972 individuals in the full sample.

Table 2 displays empirical results from three specifications of the admixture regression methodology. Recall that one SIRE variable and one ancestry proportion variable must be left out as an identification condition of the admixture regression: we leave out the White SIRE variable and the European ancestry proportion variable. Model 1 uses a linear regression specification and singleton SIRE categories for the group-identity variables *G*; individuals who choose multiple SIRE categories have *G* exposures equally divided between the chosen SIRE categories as in (10). Three of the four ancestral proportion variables and one of the six group-identity variables have statistically significant coefficients. Model 2 adds a selected set of multiple-SIRE composite categories to the *G* specification. We include the seven two-category choices with the largest number of observations in our sample. Individuals with one of these two-category choices has unit exposure to the associated explanatory variable, and no exposure to the weighted single-SIRE variables (see equation 11 above). The same three of four ancestral proportion variables as in Model 1 are significant in Model 2, with similar coefficients to Model 1. None of the single-SIRE group identity variables is significant. Three of the seven selected two-SIRE group identity variables have significantly different coefficients from that implied by equal weightings of the component single-category coefficients. One of these (Hispanic-Other) has a statistically significant coefficient; the other two are not significantly different from zero, but are significantly different from the value implied by the composite single-category coefficients. Random effects are included in all models except Model 3 to capture any common variation associated with the 22 individual data collection sites in the ABCD study or associated with those families having multiple individuals in the sample. We use the *lmer* maximum likelihood mixed effects model estimation routine from the *R* language library, see Bates et al. (2015), for all models except Model 3. See Nakagawa and Schielzeth (2013) for the definition and interpretation of conditional and marginal *R*^2^ in a linear mixed effects model. The marginal *R*^2^ (which does not include the explanatory power associated with site-specific and family-specific random effects) is approximately 0.16 in both model specifications. The African, Amerindian, and East Asian proportional ancestry variables have strong and significant explanatory power in Models 1 and 2. For single-SIRE individuals, the SIRE-based group identity variables are mostly indistinguishable from zero, but some of the multiple-SIRE group variables are significantly different from zero.

**Table 2.** Admixture Regression Results for Neuropsychological Performance Linear Specifications with and without Composite Groups and a Partially Linear Semiparametric Specification

|  | Core Explanatory Variables |  |  |  |  |  |  |  |  |  |  |
| --- | --- | --- | --- | --- | --- | --- | --- | --- | --- | --- | --- |
|  | Intercept | % African | % Amerindian | % East Asian | % South Asian | Black SIRE | Hispanic SIRE | Native American SIRE | East Asian SIRE | South Asian SIRE | Other SIRE |
| Model 1 | 0.3010 | -1.0242 | -1.3804 | 0.6439 | 0.4867 | -0.1420 | -0.0940 | -0.1428 | -0.2120 | -0.0735 | -0.1967 |
| t-statistic | 9.0640 | -8.3380 | -11.3700 | 3.1850 | 1.4910 | -1.4050 | -1.1150 | -1.3340 | -1.2060 | -0.2820 | -2.5210 |
| Model 2 | 0.2949 | -1.0298 | -1.3358 | 0.6602 | 0.5439 | -0.1503 | 0.3390 | -0.1856 | -0.3523 | -0.2679 | -0.1796 |
| t-statistic | 8.9330 | -8.2250 | -10.7830 | 3.2530 | 1.6590 | -1.4620 | 1.7250 | -1.3720 | -1.9280 | -0.9930 | -1.7640 |
| Model 3 | N/A | Figure 3 | -1.1914 | 0.6924 | 0.7377 | -0.1289 | -0.1202 | -0.2578 | -0.1369 | -0.1523 | -0.0761 |
| t-statistic |  |  | -9.9517 | 3.4772 | 2.3109 | -1.3291 | -1.4464 | -2.5228 | -0.7736 | -0.5955 | -0.9088 |
|  | Multiple-SIRE-Composite Explanatory Variables |  |  |  |  |  |  | Wald Test Statistic | Wald Test p-value |  |  |
|  | Black-White SIRE | Hispanic-White SIRE | Native America - White SIRE | East Asian - White SIRE | South Asian - White SIRE | Hispanic - Black SIRE | Hispanic - Other SIRE |  |  |  |  |
| Model 2 [cont.] | 0.0533 | -0.0813 | -0.0999 | 0.0043 | 0.3361 | 0.0361 | -0.1657 |  |  |  |  |
| t-statistic | 0.7110 | -1.7890 | -1.1850 | 0.0420 | 1.8000 | 0.2940 | -2.4000 |  |  |  |  |
| Test 2 | 2.2835 | -2.5520 | -0.0689 | 0.9357 | 1.5102 | -0.4236 | -2.0061 | 27.0810 | 0.0003 |  |  |
| Conditional R2 |  | Model 1: 0.550; Model 2: 0.550; Model 3:NA |  |  |  |  |  |  |  |  |  |
| Marginal R2 |  | Model 1: 0.157; Model 2: 0.160; Model 3:NA |  |  |  |  |  |  |  |  |  |

Notes to Table: Model 1 uses single-SIRE categories with multiple-SIRE choices allocated evenly across them; Model 2 adds seven multiple-SIRE categories. Model 4 uses semiparametric estimation and single-SIRE categories as in Model 1. Test 2 gives the z-statistic for testing if the multiple-SIRE group coefficient equals the average of the component coefficients; the Wald statistic provides a joint test of all the Test 2 restrictions.

Model 3 implements a partially linear nonparametric specification. This specification requires that the highlighted ancestry proportion (whose impact is estimated nonparametically) has observations throughout the [0, 1] range. For each of the five ancestry categories, Table 3 gives the number of sample observations of proportional ancestry in decile bins of percent ancestry, for each of the five genetic ancestry categories. We use African proportional ancestry as the highlighted variable since it fulfils the requirement for observations throughout the [0, 1] interval and therefore partially linear nonparametric estimation is feasible. Figure 1 shows the probability density of African ancestry for the full sample population; Figure 2 shows the density restricted to those individuals having measured African ancestry greater than 0.5%, this provides greater detail in the graph by excluding observations with near-zero ancestry. Interestingly, this density has three local peaks, at approximately 5%, 40% and 80% African ancestry.

**Table 3.** Number of Observations in Deciles of Proportional Ancestry for Each Ancestry Category

|  |  |  |  |  |  |  |
| --- | --- | --- | --- | --- | --- | --- |
| European | Interval | $A_{1i} \leq 10\%$ | $10\% < A_{1i} \leq 20\%$ | $20\% < A_{1i} \leq 30\%$ | $30\% < A_{1i} \leq 40\%$ | $40\% < A_{1i} \leq 50\%$ |
|  | Number of Obs. | 298 | 908 | 425 | 346 | 461 |
| | Interval | $50\% < A_{1i} \leq 60\%$ | $60\% < A_{1i} \leq 70\%$ | $70\% < A_{1i} \leq 80\%$ | $80\% < A_{1i} \leq 90\%$ | $90\% < A_{1i}$ |
|  | Number of Obs. | 700 | 406 | 462 | 514 | 5452 |
| African | Interval | $A_{2i} \leq 10\%$ | $10\% < A_{2i} \leq 20\%$ | $20\% < A_{2i} \leq 30\%$ | $30\% < A_{2i} \leq 40\%$ | $40\% < A_{2i} \leq 50\%$ |
|  | Number of Obs. | 7557 | 2935 | 286 | 125 | 165 |
| | Interval | $50\% < A_{2i} \leq 60\%$ | $60\% < A_{2i} \leq 70\%$ | $70\% < A_{2i} \leq 80\%$ | $80\% < A_{2i} \leq 90\%$ | $90\% < A_{2i}$ |
|  | Number of Obs. | 88 | 130 | 406 | 787 | 149 |
| Amerindian | Interval | $A_{3i} \leq 10\%$ | $10\% < A_{3i} \leq 20\%$ | $20\% < A_{3i} \leq 30\%$ | $30\% < A_{3i} \leq 40\%$ | $40\% < A_{3i} \leq 50\%$ |
|  | Number of Obs. | 8364 | 443 | 329 | 301 | 282 |
| | Interval | $50\% < A_{3i} \leq 60\%$ | $60\% < A_{3i} \leq 70\%$ | $70\% < A_{3i} \leq 80\%$ | $80\% < A_{3i} \leq 90\%$ | $90\% < A_{3i}$ |
|  | Number of Obs. | 156 | 75 | 18 | 0 | 4 |
| East Asian | Interval | $A_{4i} \leq 10\%$ | $10\% < A_{4i} \leq 20\%$ | $20\% < A_{4i} \leq 30\%$ | $30\% < A_{4i} \leq 40\%$ | $40\% < A_{4i} \leq 50\%$ |
|  | Number of Obs. | 9455 | 74 | 84 | 20 | 225 |
| | Interval | $50\% < A_{4i} \leq 60\%$ | $60\% < A_{4i} \leq 70\%$ | $70\% < A_{4i} \leq 80\%$ | $80\% < A_{4i} \leq 90\%$ | $90\% < A_{4i}$ |
|  | Number of Obs. | 18 | 4 | 8 | 10 | 74 |
| South Asian | Interval | $A_{5i} \leq 10\%$ | $10\% < A_{5i} \leq 20\%$ | $20\% < A_{5i} \leq 30\%$ | $30\% < A_{5i} \leq 40\%$ | $40\% < A_{5i} \leq 50\%$ |
|  | Number of Obs. | 9796 | 55 | 30 | 29 | 17 |
| | Interval | $50\% < A_{5i} \leq 60\%$ | $60\% < A_{5i} \leq 70\%$ | $70\% < A_{5i} \leq 80\%$ | $80\% < A_{5i} \leq 90\%$ | $90\% < A_{5i}$ |
|  | Number of Obs. | 4 | 11 | 9 | 21 | 0 |

Notes to Table: For each of the five geographic ancestries, the table shows the number of the total 9972 observations within each of the deciles of proportional ancestry.

**Figure 1.**
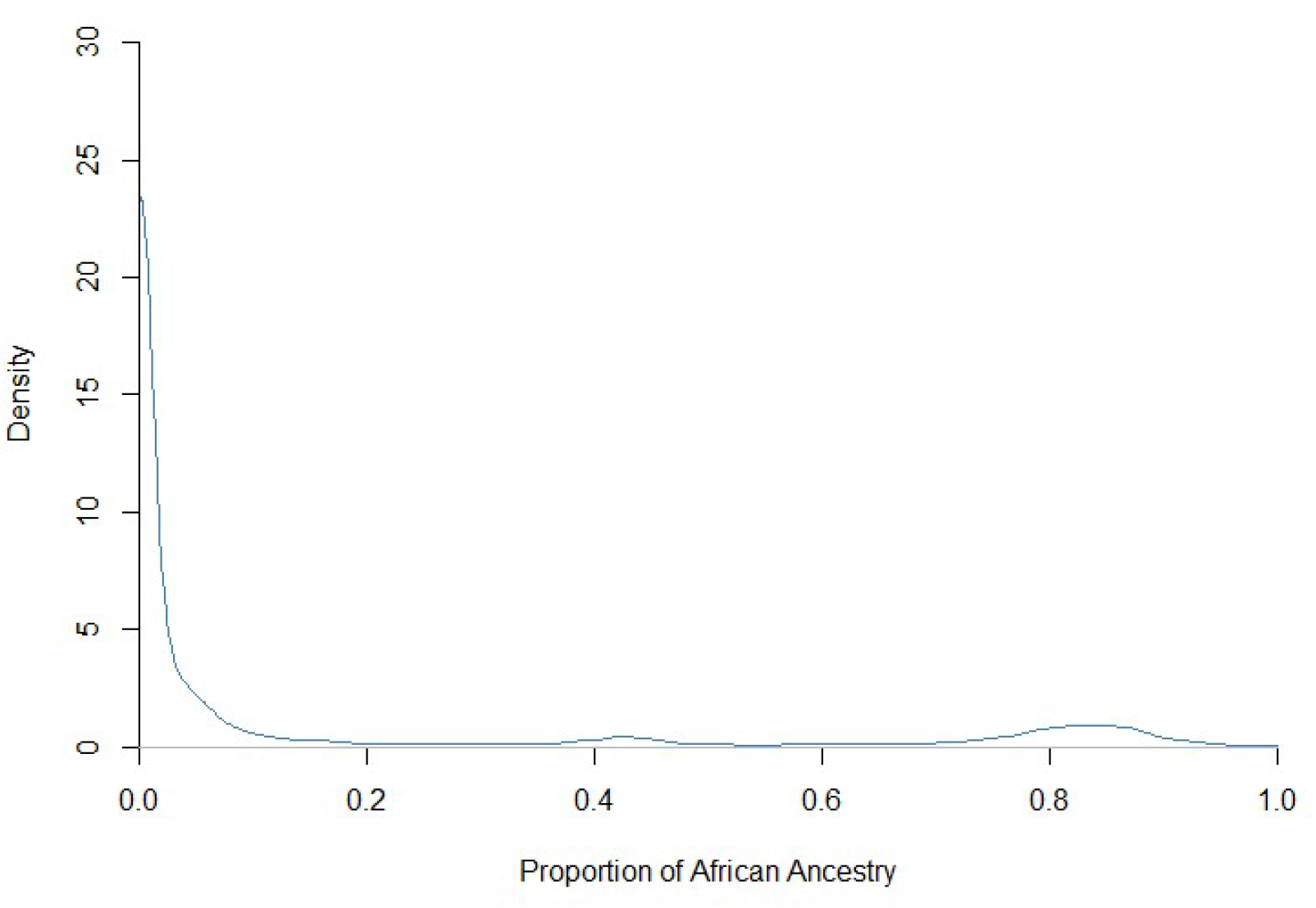
Estimated Density of African Ancestry for the Full Sample

**Figure 2.**
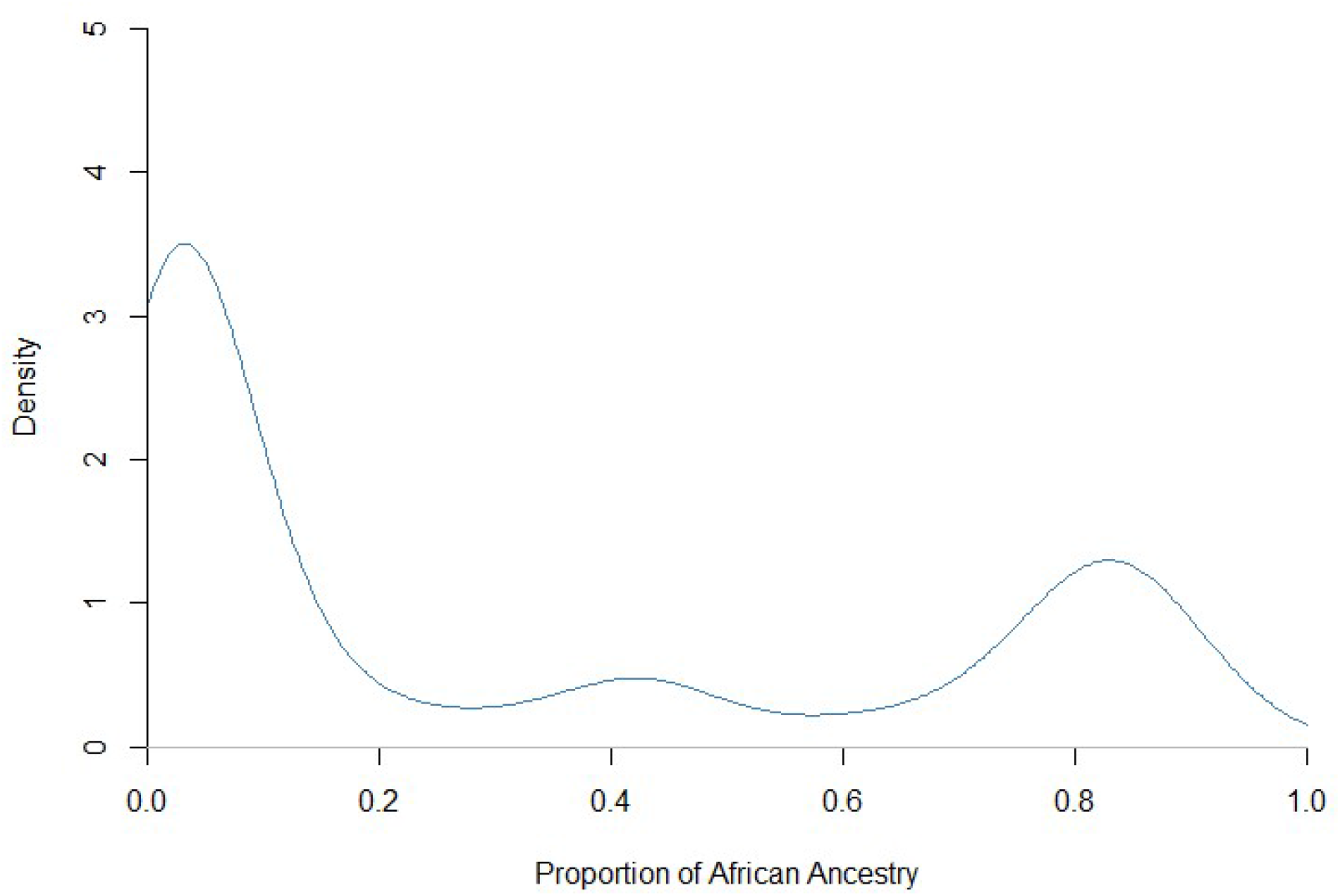
Estimated Density of African Ancestry for a Restricted Sample (Ancestry > 0.5%)

Partially linear semiparametric Model 3 (18) is estimated using the *npplr* routine in the R programming language subroutine library *NP* written and maintained by Hayfield and Racine (2020). We use the simple average SIRE specification of *G* as in Model 1. We use the Guassian kernel throughout, and all bandwidths are chosen by iterated least-squares cross-validation. The linear coefficient estimates in Model 3 do not differ notably from those in Model 1. Figure 3 displays the nonparametric estimate of the impact of African ancestry on the performance variable along with the corresponding linear impact estimate from Model 1, that is, 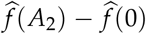 and 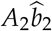 for *A*_2_ ∈ [0, 1]. There is some graphical evidence for an uptick in the nonlinear gradient for ancestry proportions above 90%. We now briefly examine this further.

**Figure 3.**
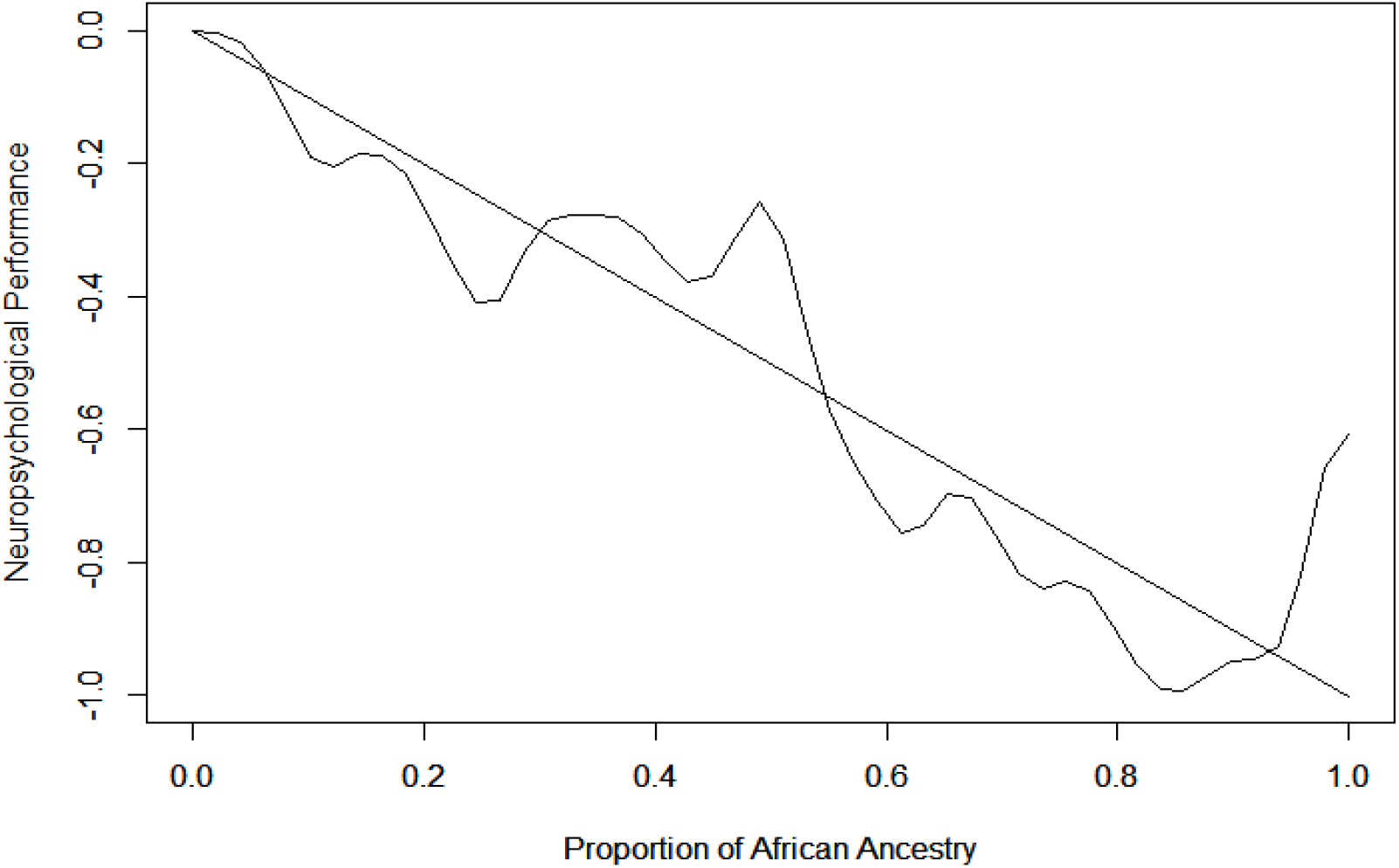
Linear and Nonlinear Gradients Measuring the Impact of African Ancestry

Model 3 does not capture the efficiency gain and test statistic bias reduction from the mixed effects modeling used in the estimation of the other models. Figure 3 of Model 3 is estimated in the second stage of a two-stage semiparametric estimation process and this weakens its empirical reliability. To examine more carefully the graphical pattern observed in Figure 3, but with single-stage estimation and the advantage of mixed effects modeling, we estimate a piecewise linear specification for *A*_*i*2_ ≥ 0.9. This was chosen in order to mimick the observed nonlinear uptick seen in Figure 3 within a linear regression functional form. Recall that African ancestry proportion is ancestry variable 2, giving the formulation:

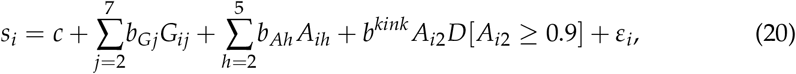

where *D*[•] is a zero-one dummy variable and *b^kink^* is the added coefficient. The results are shown as Models 4 and 5 in Table 4. In Model 4 we use the simple average SIRE specification of *G* as in Model 1; Model 5 adds the same seven two-SIRE combination groups as in Model 2. The coefficient *b^kink^* is significantly positive in one of the two models; the significance of this finding must be treated with caution since the particular kink specification (20) is based on examination of Figure 3 using the same data.

**Table 4.**
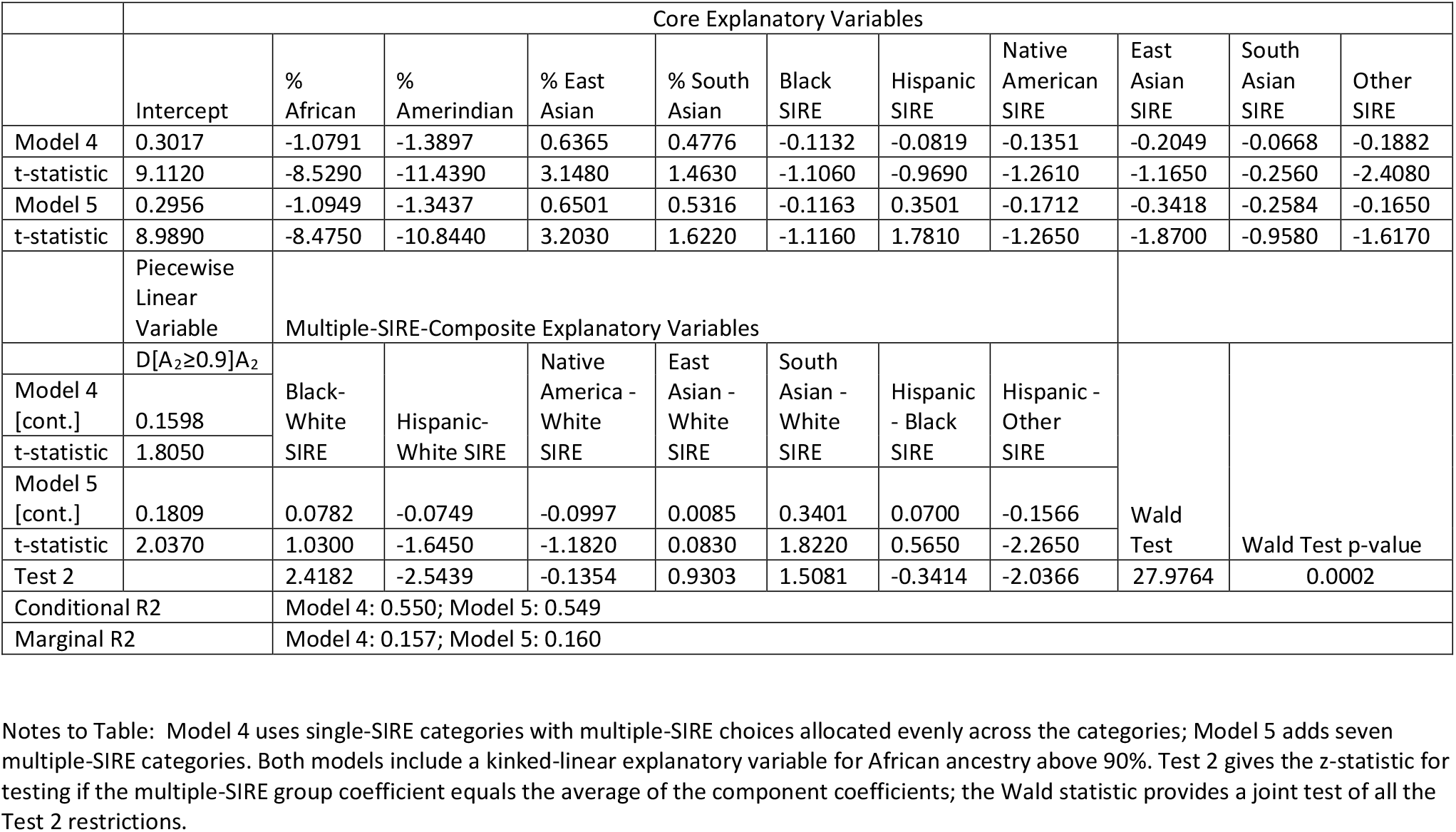
Piecewise Linear Admixture Regression Results with and without Composite Groups

Table 5 adds two new variables, US born child and Social-Economic Status (SES), to the admixture regression model. US born child equals one if the child was born in the USA and zero if born elsewhere. SES is a factor-analytic composite of underlying variables from the ABCD database including neighborhood SES, subjective SES as determined from a set of questionaire answers by the parent(s)/guardian(s) of the child on parental/guardian marital status, completed level of parental/guardian education, reported neighborhood safety, and parental/guardian employment. See the Supplemental Materials for more detailed discussion. Models 6 and 7 are identical to Models 4 and 5 (respectively) from Table 3, except for the addition of these two variables. As discussed in Section 3 above, including additional explanatory variables complicates the interpretation of an admixture regression model in terms of the implied decomposition of trait variation into linear components linked to group identities and components linked to genetic ancestries. The SES variable covaries strongly with both genetic and environmental components of neuropsychological performance scores. To retain the standard interpretability of the admixture regression it is important to orthogonalize SES with respect to the group identity and ancestry variables before running the regression. For completeness, Models 6 and 7 are shown with and without the orthogonalization of SES (versions a and b of each model). If the purpose of the estimation is to identify the total impact of SES on the trait, the regression with raw SES is more appropriate (version a). For admixture analysis intended to capture the total effects of group identity and genetic ancestry on the trait, orthogonalized SES is more appropriate (version b).

**Table 5.**
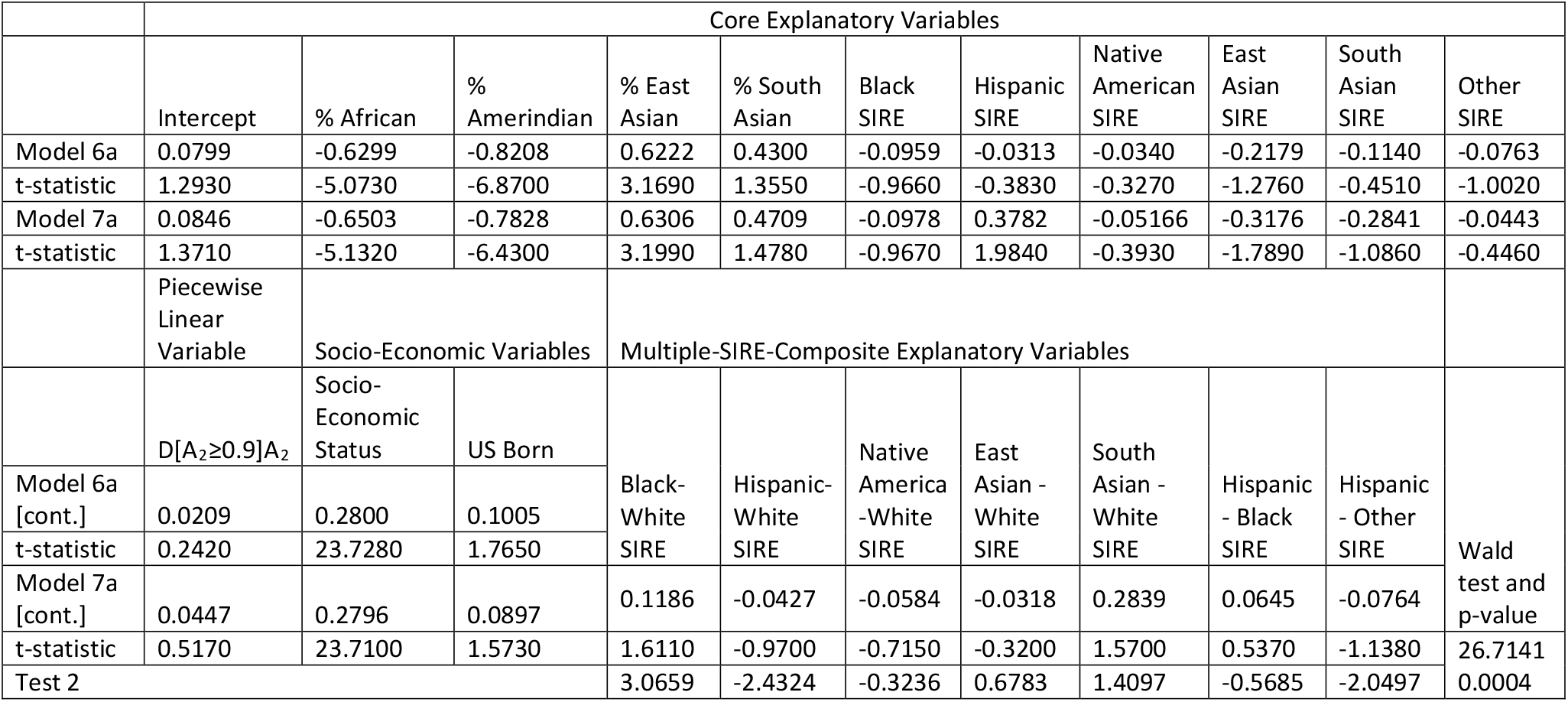
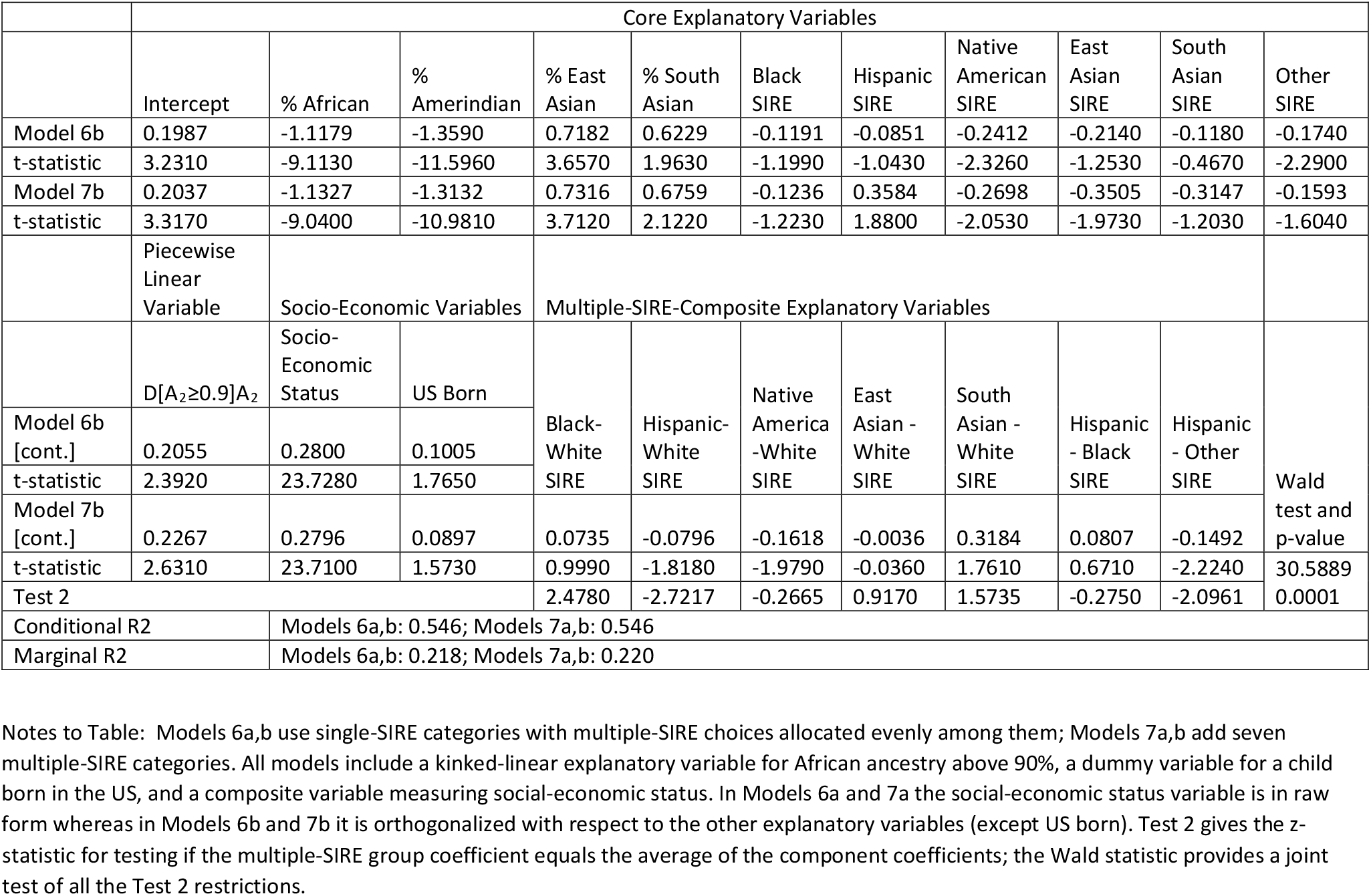
Linear Admixture Regression Results Including Social-Economic Status (SES) and US Born Variables 5a: Using Raw SES 5b: Using Orthogonalized SES

Adding SES to the admixture regression model increases the marginal *R*^2^ from approximately 0.16 to 0.22. If SES is used in its raw form, the coefficients associated with proportional ancestries tend to decrease in magnitude, but the coefficients on African, Amerindian, and East Asian proportional ancestry remain strong and significant. There is no clear and reliable impact on the SIRE-based group-identity coefficients from using SES, in either its raw or orthogonalized form.

## 6. Discussion and Limitations

Many behavioral traits covary strongly with racial/ethnic self-identities, but it is often ambigous whether this covariance reflects environmental causes associated with racial/ethnic identity groups or reflects underlying genetic similarity among group members arising from shared geographic ancestry. Admixture regression relies on the natural experiment of recent genetic admixture of previously geographically-isolated ancestral groups to measure the explanatory power arising from racial/ethnic group identities and that arising from ancestry-based similarities of genetic background. The admixture regression methodology, in various formulations, has been applied to a wide range of medical and behavioral traits including asthma, obesity, type 2 diabetes, hypertension, neuropsychological performance, and sleep depth.

This paper provides a statistical framework for admixture regression based on the linear polygenic index model of behavioral genetics, and develops refinements and extensions of the methodology within this framework. We provide a simple new test procedure for determining whether multiple-SIRE categories have independent explanatory power not captured by the individual component categories. We consider additional explanatory variable in the admixture regression and their interpretation with and without orthogonalization with respect to core variables. We weaken the linearity assumption and develop a partially linear semiparametric regression specification.

We apply our methodology to neuropsychological performance test data from the Adolescent Brain Cognitive Development database. We confirm existing findings that genetic variation plays a role in neuropsychological performance differences across self-identified races (Lasker et al. 2019). We find mixed evidence regarding the independent explanatory power of multi-racial identities relative to their component single-race categories. We find that when social econonomic status (SES) is included as an explanatory variable in the admixture regression, pre-regression orthogonalization of SES has a substantial impact on the measured magnitude of the ancestry proportion coefficients. We find that the proportional ancestry variable associated with African ancestry shows some evidence of nonlinearity in its impact on neuropsychological performance.

The techniques that we propose for admixture regression studies have broad applicability, but they do have some limitations. We describe three approaches to accommodating multiple-identity individuals in admixture regression studies (equal weighting, adding new groups, deletion of some observations) but none of the three methods is fool-proof in terms of correctly capturing identity-related environmental influences in a parsimonious way. We describe how to orthogonalize additional explanatory variables in order to accommodate them in an admixture regression while still capturing the full effect of ancestry-related genetic variation in the ancestry proportions coefficients. A limitation of this orthogonalization procedure is that it does not fundamentally alter the underlying regression being estimated, it merely rotates the estimated coefficients to aid in their interpretation. The partially linear admixture regression method that we describe has the usual limitations of nonparametric and semi-parametric estimation methods. It cannot be applied with complete generality due to the curse of dimensionality, and is data-intensive due to the nonparametric estimation component of the procedure.

## Data Availability Statement

All data used in this study comes from the Adolescent Behavior Cognitive Development (ABCD) database. Qualified researchers can request access to the ABCD database by applying through the National Institute of Mental Health Data Archive.

## Appendix A

In this technical appendix we re-state condition 7.1 from (Racine and Li 2007, p. 224) in the context of our partially linear admixture regression model (18).

We assume that the (*k* + *m* – 1)–vector of observations (*s_i_, G_ij_, A_ih_*) *j* = 2,…, *k*; *h* = 2,…, *m* has an i.i.d. distribution over observations *i* = 1,…, *n* and that the conditional mean functions E [*G_ij_*|*A*_*i*2_] and E[*A_ih_*|*A*_*i*2_] are twice differentiable throughout the interior of the domain of *A*_2_, the closed unit interval. Let *m*(•) denote any of these conditional mean functions or their first or second derivative functions. As in Racine and Li, we impose the following Lipschitz-type smoothness condition on these conditional mean functions and their first and second derivatives: 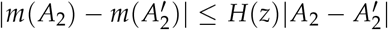 where *H*(•) is some continuous function such that E[*H*(*A*_2_)^2^] is finite. The expectation of *H*(*A*_2_)^2^ is over the probability distribution of *A*_2_.

We continue to assume that *ε_i_* is mean-zero normally distributed with constant variance. Since *G_ij_* only takes the values of zero and one and *A_ih_* is confined to the unit interval, it necessarily follows that both have bounded fourth moments. We assume that *k*(•) is a bounded second-order kernel.

To formally derive the limiting distribution of the Robinson estimator, it is necessary to define a trimming parameter which ensures that the estimates 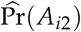 are bounded away from zero. Let *t* denote a trimming parameter and consider the estimator described in the text but where observations such that 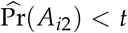 in (19) are dropped from the subsequent estimation steps. Let *ϕ* denote the kernel bandwidth for sample size *n*. Assume that the trimming parameter obeys the following two limiting conditions as *n* → ∞: *nϕ*^2^*t*^4^ → ∞ and *nt*^-4^*ϕ*^8^ → 0.

Under these conditions we have from (Robinson 1988) that

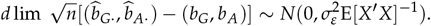

where the matrix *X* is defined in the main text of the paper above, in step two of the Robinson procedure.

