## Supplemental File 1 for "Linear and partially linear models of behavioral trait variation using admixture regression"

Supplemental File 1: Supplementary material for “Linear and partially linear models of behavioral trait variation using admixture regression” in which the dataset, variables, and methods for the empirical analysis are detailed.

### 1. Materials and Methods

#### 1.1 Dataset

The Adolescent Brain Cognitive Development Study (ABCD) is a collaborative longitudinal project between 21 sites across the US. Its goal is to further research into the psychological and neurobiological basis of development. At baseline, around 11,000 9-10 year old children were sampled, using a probabilistic sampling strategy, from public and private elementary schools and through non-school-based community outreach between 2016 and 2018, with the goal of creating a broadly representative sample of US children of this age. Children who were not fluent in English (or whose parents were not fluent in either English or Spanish) were excluded, along with those with severe medical, neurological, or psychiatric conditions. Informed consent was provided by parents.

For phenotypic data, ABCD 2.1 data release was used. For this analysis, we excluded individuals who did not have NIH Toolbox® results, who did not have admixture data, and who were identified as being a Pacific Islander. This left 9972 individuals.

#### 1.2 Variables

##### *1.2.1. Admixture*

Subjects were genotyped using Illumina XX, with 516,598 variants directly genotyped and surviving the quality control done by the data provider. We used (updated with) the 3.0 release of the genotypic dataset, which also includes an edition with imputed variants using TOPMED and Eagle 2.4. Because we had very few samples from Pacific Islanders, we excluded these from further analysis to simplify the reference populations needed (n = 69). All our work was done on build 38. Files in hg17/37 were lifted to hg38 using liftOver (https://github.com/sritchie73/liftOverPlink) and the GRC chain file at ftp://ftp.ensembl.org/pub/assembly_mapping/homo_sapiens/ (GRCh37_to_GRCh38.chain.gz).

Before global admixture estimation, we applied quality control using plink 1.9. We used only directly genotyped, bi-allelic, autosomal SNP variants (494,433, 493,196, before and after lifting). We pruned variants for linkage disequilibrium at the 0.1 $R^2$ level using plink 1.9 (--indep-pairwise 10000 100 0.1), as recommended in the admixture documentation (https://vcru.wisc.edu/simonlab/bioinformatics/programs/admixture/admixture-manual.pdf). This variant filtering was done in the reference population dataset to reduce bias from sample representativeness. After pruning, we were left with 99,642 variants. To ensure a reasonable balance in the estimation dataset, we merged the target samples from ABCD, with reference population data for the populations of interest. We desired a k=5 solution (European, Amerindian, African, East Asian, and South Asian), so we merged with relevant samples from 1000 Genomes and from the HGDP. The following populations were excluded: Adygei, Balochi, Bedouin, Bougainville, Brahui, Burusho, Druze, Hazara, Makrani, Mozabite, Palestinian, Papuan, San, Sindhi, Uygur, Yakut. These reference populations were excluded because they were overly admixed or because, in the case of Melanesians and San, the individuals in the ABCD sample lacked significant portions of these ancestries.

Because the estimation sample would still be very skewed towards European ancestry using this joint sample, we used repeated subsetting to achieve balance. Specifically, we split the ABCD target samples into 50 random subsets, each with about 222 persons, and merged them one at a time with the reference data, followed by running admixture k=5 on each merged subset. We verified that these subsets produced stable results by examining the stability of the estimates for the reference samples. There was very little variation across runs, e.g. for the reference sample with the most variance (European, NA12342), the mean estimate was 98.3% with SD=0.17% across the 50 runs. Since Admixture does not label the resulting clusters, we used 5 reference samples to index the populations so the data would be merged correctly. In no case did this produce any inconsistencies.

*1.2.2. Neuropsychological Performance*

The NIH Toolbox® (NIHTBX) neuropsychological battery was designed to measure a broad range of cognitive abilities. It consists of seven tasks which index attention (Flanker Inhibitory Control and Attention Task), episodic memory (Picture Sequence Memory Task), language abilities (Picture Vocabulary Task & Oral Reading Recognition Task), executive function (Dimensional Change Card Sort Task & Flanker Inhibitory Control and Attention Task), processing speed (Pattern Comparison Processing Speed Task), and working memory (List Sorting Working Memory Task) (Akshoomoff et al., 2014; Weintraub et al., 2013). NIHTBX was normed for samples between ages 3 and 85; tasks correlate highly with comparable ability assessmnents (Weintraub et al., 2013). Moreover, this battery has been shown to be measurement invariant across American ethnic groups (Lasker, Pesta, Fuerst, & Kirkegaard, 2019).

Age-corrected composite scores, based on the seven tasks, were provided by ABCD. We regressed out sex from these age-corrected composite scores. The residuals were then standardized.

*1.2.3. Self-identified Race and Ethnicity*

Self-identified race was based on parental responses to 18 questions asking about the child's race ("What race do you consider the child to be? Please check all that apply"). From these questions, six broad racial categories were created: European ("White"), African ("Black/African American"), Native American ("American/Native American" and "Alaska Native"), South Asian ("Asian Indian"), East Asian ("Chinese," "Filipino," "Japanese," "Korean," and "Vietnamese", "Other Asian,"), and Other ("Other race," "Refused to answer," "Don't know"). The Other Asian group ($N$ = 66) was classified as “East Asian” because the Asian ancestry component was predominantly East (44%;) not South (7%) Asian; the remaining ancestry was predominantly European (40%). The Pacific Islander groups (“Native Hawaiian,” “Guamanian,” “Samoan,” and “Other Pacific Islander”) were excluded as we did not have a corresponding admixture component. Self-identified ethnicity was based on parental responses to 1 question asking about Latin American ethnicity (“Do you consider yourself Hispanic/Latino/Latina?”). From this we created an additional ethnic category.

Descriptive statistics for the SIRE groups are shown in Table S1. Statistics are reported for single ethnic categories i.e., individuals reported as being only White, Black, East Asian, Native American, or Other, with no combinations (e.g., Hispanic & White), Hispanics, the seven top double combinations (i.e., Hispanic & White, Hispanic & Black, Hispanic & Other, non-Hispanic Black & White, non-Hispanic East Asian & White, non-Hispanic Native American & White, and non-Hispanic South Asian & White) and finally all other remaining groups combined.

Table 1. Descriptive Statistics for the SIRE Groups.

| | *N* | Age *M* | Eur. % | Afr. % | E.Asian % | S.Asian % | Amer. % | US Born *N* | Neuro psy. | SES |
|---|---|---|---|---|---|---|---|---|---|---|
| Total Sample | 9972 | 9.91 (0.63) | 0.74 (0.31) | 0.16 (0.29) | 0.03 (0.12) | 0.01 (0.06) | 0.06 (0.13) | 9703 | 0.00 (1.00) | 0.00 (1.00) |
| NH White Only | 5533 | 9.93 (0.63) | 0.97 (0.05) | 0.01 (0.02) | 0.00 (0.02) | 0.01 (0.02) | 0.01 (0.03) | 5459 | 0.25 (0.92) | 0.40 (0.75) |
| NH Black Only | 1434 | 9.91 (0.61) | 0.18 (0.11) | 0.80 (0.11) | 0.00 (0.02) | 0.00 (0.01) | 0.01 (0.02) | 1402 | -0.77 (0.87) | -1.00 (0.95) |
| NH East Asian Only | 107 | 10.02 (0.62) | 0.14 (0.18) | 0.01 (0.05) | 0.82 (0.23) | 0.02 (0.10) | 0.01 (0.02) | 88 | 0.57 (1.02) | 0.60 (0.75) |
| NH South Asian Only | 43 | 10.03 (0.68) | 0.24 (0.13) | 0.00 (0.00) | 0.03 (0.07) | 0.73 (0.14) | 0.01 (0.01) | 35 | 0.45 (1.02) | 0.88 (0.46) |

| | | | | | | | | | | |
|---|---|---|---|---|---|---|---|---|---|---|
| NH Native American Only | 31 | 9.70 (0.60) | 0.71 (0.30) | 0.11 (0.26) | 0.01 (0.02) | 0.01 (0.01) | 0.15 (0.19) | 31 | -0.42 (0.79) | -0.81 (0.72) |
| NH Other Only | 97 | 9.96 (0.61) | 0.55 (0.30) | 0.28 (0.31) | 0.06 (0.19) | 0.04 (0.11) | 0.07 (0.15) | 87 | -0.22 (1.12) | -0.53 (1.11) |
| Any Hispanic | 1869 | 9.88 (0.63) | 0.60 (0.20) | 0.10 (0.14) | 0.02 (0.06) | 0.01 (0.02) | 0.27 (0.18) | 1755 | -0.23 (0.96) | -0.41 (0.93) |
| White Hispanic | 1171 | 9.89 (0.64) | 0.67 (0.18) | 0.06 (0.06) | 0.01 (0.02) | 0.01 (0.01) | 0.26 (0.17) | 1097 | -0.19 (0.96) | -0.29 (0.91) |
| Black Hispanic | 84 | 9.80 (0.64) | 0.35 (0.13) | 0.53 (0.17) | 0.00 (0.01) | 0.00 (0.01) | 0.11 (0.09) | 79 | -0.34 (0.93) | -0.64 (0.98) |
| Other Hispanic | 411 | 9.89 (0.63) | 0.49 (0.15) | 0.09 (0.11) | 0.02 (0.03) | 0.00 (0.01) | 0.40 (0.17) | 383 | -0.45 (0.92) | -0.75 (0.88) |
| NH Black & White | 302 | 9.88 (0.62) | 0.58 (0.12) | 0.41 (0.12) | 0.00 (0.01) | 0.00 (0.01) | 0.01 (0.02) | 301 | -0.13 (0.95) | -0.45 (1.06) |
| NH East Asian & White | 249 | 9.99 (0.64) | 0.56 (0.12) | 0.01 (0.01) | 0.41 (0.14) | 0.02 (0.04) | 0.01 (0.01) | 244 | 0.58 (0.98) | 0.66 (0.67) |
| NH Native American & White | 131 | 9.78 (0.60) | 0.90 (0.10) | 0.01 (0.03) | 0.01 (0.02) | 0.01 (0.01) | 0.07 (0.09) | 131 | 0.01 (0.86) | -0.09 (0.91) |
| NH South Asian & White | 40 | 9.79 (0.54) | 0.63 (0.11) | 0.00 (0.00) | 0.02 (0.08) | 0.34 (0.12) | 0.01 (0.01) | 39 | 0.83 (0.84) | 0.78 (0.77) |
| Any_Other NH combination | 136 | 9.87 (0.62) | 0.37 (0.21) | 0.46 (0.25) | 0.13 (0.20) | 0.02 (0.07) | 0.02 (0.05) | 131 | -0.27 (1.06) | -0.63 (1.03) |

*Note*: Euro.% = European ancestry percentage, Afr.% = African ancestry percentage, E.Asian% = East Asian Ancestry percentage, S.Asian% = South Asian Ancestry percentage, Neuropsy. = Neuropsychiatric performance, SES = general socioeconomic component score. NH = non-Hispanic. Hispanic & White, Hispanic & Black, and Hispanic & Other are subsets of the Any Hispanic category.

The racial and ethnic variables were then recoded to create interval categories for which individuals are assigned a percentage of each SIRE category based on the number of responses chosen (Liebler & Halpern-Manners, 2008; Kirkegaard et al., 2019). By this coding, if someone was marked as White and Hispanic, they were assigned scores of .5 for white and .5 for Hispanic and 0 for the other 5 categories. The correlations between these interval scores and genetic ancestry components are shown in Table S2 below (*N* = 9972). These associations are similar to those found by others (for example: Guo et al. 2014); self-identified race generally corresponds with genetic ancestry.

Table S2. Correlations between Interval Coded SIRE and Genetic Ancestry.

| | European ancestry | African ancestry | East Asian ancestry | South Asian ancestry | Amerindian ancestry |
|---|---|---|---|---|---|
| White_SIRE | **0.91** | -0.74 | -0.21 | -0.05 | -0.31 |
| Black_SIRE | -0.76 | **0.96** | -0.08 | -0.09 | -0.18 |
| East_Asian_SIRE | -0.24 | -0.08 | **0.92** | 0.02 | -0.06 |
| South_Asian SIRE | -0.11 | -0.04 | 0.01 | **0.87** | -0.03 |
| Native_SIRE | -0.02 | 0.00 | -0.01 | -0.03 | 0.07 |
| Hispanic_SIRE | -0.21 | -0.10 | -0.04 | -0.06 | **0.77** |
| Other_SIRE | -0.16 | -0.01 | 0.00 | 0.00 | **0.38** |

##### *1.2.4. Region of Birth (US Born)*

Region of birth was based on the parental response to the question, “In which country was the child born?”. The response “United States” was recoded as 1 and all other responses were recoded as 0.

*1.2.5. Socioeconomic Status*

Socioeconomic status was based on seven indicators: financial adversity, area deprivation index, neighborhood safety protocol, parental education, parental income, parental marital status, and parental employment status. These are detailed below:

**1.2.5.1. Financial Adversity (Reverse Coded)**. Parents answered a seven item Financial Adversity Questionnaire (PRFQ). They were asked: “In the past 12 months, has there been a time when you and your immediate family experienced any of the following:

(1) “Needed food but could not afford to buy it or could not afford to go out to get it?”,
(2) “Were without telephone service because you could not afford it?”
(3) “Did not pay the full amount of the rent or mortgage because you could not afford it?”,
(4) “Were evicted from your home for not paying the rent or mortgage?”,
(5) “Had services turned off by the gas or electric company, or the oil company would not deliver oil because payments were not made?”,
(6) “Had someone who needed to see a doctor or go to the hospital but did not go because you could not afford it?”, and
(7) “Had someone who needed a dentist but could not go because you could not afford it?”
For each of the seven items they answered “yes” (1) or “no” (0). We summed responses. Thus the maximum was 7 and the minimum was 0.

This variable was reverse coded, so that higher scores indicated less financial adversity, and then standardized.

**1.2.5.2. Area Deprivation Index (ADI) (Reverse Coded)**. Parents completed a residential history questionnaire. They provided the residential addresses and the number of full years they lived at each residence. For each address an Area Deprivation Index (ADI) was computed by ABCD and the national percentile of the area's socioeconomic status was given. ADI was based on the following variables:

1. "Percentage of occupied housing units without complete plumbing (log)"
2. "Percentage of occupied housing units without a telephone"
3. "Percentage of occupied housing units without a motor vehicle"
4. "Percentage of single"
5. "Percentage of population below 138% of the poverty threshold"
6. "Percentage of families below the poverty level"
7. "Percentage of civilian labor force population aged >=16 y unemployed (unemployment rate)"
8. "Percentage of occupied housing units with >1 person per room (crowding)"
9. "Percentage of owner"
10. "Median monthly mortgage"
11. "Median gross rent"
12. "Median home value"
13. "Income disparity defined by Singh as the log of 100 x ratio of the number of households with <10000 annual income to the number of households with >50000 annual income"
14. "Median family income"
15. "Percentage of population aged >=25 y with at least a high school diploma"
16. "Percentage of population aged >=25 y with <9 y of education"

Scores were provided in terms of national percentiles. We used scores for the most recent residence (variable: reshist_addr1_adi_perc). The resultant values were reverse coded to make higher values indicate better neighborhoods, and then standardized.

**1.2.5.3. Neighborhood Safety Protocol**. Parents were asked three Likert scale (1 = strongly disagree; 5 = strongly agree) questions about neighborhood safety: "I feel safe walking in my neighborhood, day or night," "Violence is not a problem in my neighborhood," and "My neighborhood is safe from crime." We used the precomputed summary scores for which the three scores were summed and then divided them by three.

**1.2.5.3. Education.** Parents were asked, "What is the highest grade or level of school you have completed or the highest degree you have received." To create an interval variable, we recoded parental education as 0 to 18: Never attended/Kindergarten only = 0, 1st grade = 1, 2nd grade = 2, 3rd grade = 3, 4th grade = 4, 5th grade = 5, 6th grade =  6, 7th grade = 7, 8th grade = 8, 9th grade = 9, 10th grade = 10, 11th grade = 11, 12th grade = 12,  High school graduate =12, GED or equivalent Diploma General =12, Associate degree: Occupational Program =14, Associate degree: Academic Program = 14, Bachelor's degree = 16, Master's degree = 18, Professional school = 18, Doctoral degree = 18. We standardized the scores for each  educational scores for both parents and standardized the averaged scores.

**1.2.5.4. Income.** Family was an interval variable which reflected the parents' total combined family income in the past 12 months. The variable was recoded as follows: 1.00 = less than $5000 (recode: 4,500); 2.00 = $5000 to 11,999 (recode: 5,000); 3.00 = $12,000 to 15,999 (recode: 12,000); 4.00 = $16,000 to 24,999 (recode: 16,000); 5.00 = $25,000 to 34,999 (recode: 25,000); 6.00 = $35,000 to 49,999 (recode: 35,000); 7.00 = $50,000 to 74,999 (recode: 50,000); 8.00 = $75,000, to 99,999 (recode: 75,000); 9.00 = $100,000 to 199,999 (recode: 100,000); 10.00 = $200,000 and greater (recode: 200,000).

**1.2.5.5. Marital Status.** Parental marital status was coded as 1 if married and 0 for any other arrangement.

**1.2.5.6. Employment Status.** Parental employment was coded as 1 if at least one parent was working now either full or part time and 0 for all other cases.

**3.2.5.7. General SES.** Missing data for the seven economic indicators were imputed using the mice package (df, m=5, maxit = 50, method = 'pmm', seed = 500).  Descriptive statistics for the imputed SES indicators are provided in Table S3, while the correlation matrix for the imputed variables (N = 9972), along with neuropsychiatric performance, is shown in Table S4.

Table S3. Descriptive Statistics &Correlation Matrix for the SES indicators (*N* = 9972 in all cases)

| Variable | *M* | *SD* | 1 | 2 | 3 | 4 | 5 | 6 | 7 | 8 |
|---|---|---|---|---|---|---|---|---|---|---|
| 1. Financial Adversity | 0.00 | 1.00 | | | | | | | | |
| 2. ADI | 0.00 | 1.00 | .21** [.19, .23] | | | | | | | |
| 3. Neighborhood Safety Protocol | 0.00 | 1.00 | .24** [.22, .26] | .25** [.23, .27] | | | | | | |
| 4. Education | 0.00 | 1.00 | .28** [.27, .30] | .28** [.26, .30] | .28** [.27, .30] | | | | | |
| 5. Income | 0.00 | 1.00 | .34** [.33, .36] | .34** [.33, .36] | .31** [.29, .33] | .57** [.56, .58] | | | | |
| 6. Marital Status | 0.68 | 0.47 | .27** [.25, .29] | .23** [.21, .25] | .23** [.21, .25] | .33** [.31, .35] | .46** [.44, .47] | | | |
| 7. Employment Status | 0.91 | 0.28 | .16** [.14, .18] | .13** [.11, .15] | .15** [.13, .17] | .26** [.25, .28] | .27** [.25, .28] | .28** [.27, .30] | | |
| 8. General SES | 0.00 | 1.00 | .56** [.55, .58] | .54** [.53, .56] | .55** [.53, .56] | .73** [.72, .74] | .80** [.79, .81] | .66** [.64, .67] | .48** [.46, .49] | |
| 9. Neuropsy. | 0.00 | 1.00 | .20** [.18, .22] | .21** [.19, .23] | .18** [.16, .20] | .40** [.39, .42] | .35** [.33, .37] | .27** [.26, .29] | .18** [.16, .20] | .43** [.41, .44] |

*Note*. *M* and *SD* are used to represent mean and standard deviation, respectively. Values in square brackets indicate the 95% confidence interval for each correlation. The confidence interval is a plausible range of population correlations that could have caused the sample correlation (Cumming, 2014). * indicates $p < .05$. ** indicates $p < .01$.

We then submitted the seven SES indicators to Principal Component Analysis (PCA). For this, we used the R package PCAmixdata, which handles mixed categorical and continuous data (Chavent, Kuentz-Simonet, & Saracco, 2014). The first unrotated component explained 39% of the variance. The PCA_1 loadings for the seven SES indicators were as follows: financial adversity (.317), area deprivation index (.296), neighborhood safety protocol (.298), parental education (.529), parental income (.641), parental marital status (.431), and parental employment status (0.230). The vector of PCA_1 loadings correlated with the vector of SES indicator effects on Neuropsychiatric Performance at $r = .91$ ($N$=7). Similarly, the vector of PCA_1 loadings strongly correlated with the vectors of genetic ancestry, with absolute values of $r = .62$ to $r = .95$ ($N$=7). Table S4 shows the correlations between SES indicators and genetic ancestry for all five ancestry components. These results indicate that the better measures of general SES have stronger correlations with both Neuropsychiatric Performance and genetic ancestry.

Table S4. General SES-loadings and the Correlations between SES Indicators and Genetic Ancestry within the Full Sample.

| | General SES Loadings | *r*Europe | *r*African | *r*Amerindian | *r*East Asian | *r*South Asian |
|---|---|---|---|---|---|---|
| Financial Adversity | 0.317 | 0.24 | -0.26 | -0.07 | 0.05 | 0.05 |
| ADI | 0.296 | 0.26 | -0.32 | 0.00 | 0.06 | 0.04 |
| Neighborhood Safety Protocol | 0.298 | 0.30 | -0.28 | -0.12 | 0.02 | 0.04 |

| | | | | | | |
|---|---|---|---|---|---|---|
| Education | 0.529 | 0.38 | -0.29 | -0.39 | 0.10 | 0.12 |
| Income | 0.641 | 0.38 | -0.36 | -0.23 | 0.09 | 0.11 |
| Marital Status | 0.431 | 0.37 | -0.40 | -0.09 | 0.07 | 0.07 |
| Employment Status | 0.230 | 0.21 | -0.20 | -0.08 | 0.02 | 0.03 |
| ***r*****(SES loadings x ancestry correlations )** | | **0.89** | **-0.62** | **-0.73** | **0.87** | **0.95** |

*Note:* Of relevance is the magnitude, not direction, of the correlations between the vector of SES loadings and the vectors of ancestry correlations.

PCA_1 scores correlated with Neuropsychological Performance at $r = .43$ in the full sample. Among non-Hispanic Whites, non-Hispanic Blacks, and Hispanics, the correlations between PCA_1 and Neuropsychological Performance was $r = .25$ ($N = 5533$), $r = .31$ ($N = 1434$), and $r = .31$ ($N = 1869$), respectively. These magnitudes of child-parental SES correlations are consistent with those previously reported (Flores-Mendoza, Ardila, Gallegos, & Reategui-Colareta, 2021; Sirin, 2005). The congruence coefficients for the SES component loadings were greater or equal to $r = .97$ for the largest three SIRE groups (non-Hispanic Whites, non-Hispanic Blacks, and Hispanics), indicating identical structures across groups.

#### 2. Methods

A series of regression models were run with NIHTBX as the dependent variable. The NIHTBX and socioeconomic variables were standardized (based on the subsample of 9972 retained). The ancestry variables were left unstandardized, thus the coefficients from ancestries can be interpreted as a change in 100% ancestry over a change in one standardized unit of NIHTBX scores.

European ancestry and White SIRE were selected as reference values and thus not included as independents.

For the regression analyses, following the recommendations of Heeringa and Berglund (2021), we used a three-level (site, family, individual) multi-level mixed effects model. This model was applied to the pooled twin and regular ABCD baseline sample. This specification approximates the ABCD Data Exploration and Analysis Portal (DEAP) specification (Heeringa and Berglund, 2021).
