## Supplemental File 2 for "Linear and partially linear models of behavioral trait variation using admixture regression"

R estimation code library for Linear and partially linear models of behavioral trait variation

```
####

#merge csv files
library(pacman)
library(rlang)
library(data.table)

##
## Attaching package: 'data.table'

## The following object is masked from 'package:rlang':
##
##     :=

library (plyr)
library(dplyr)

##
## Attaching package: 'dplyr'

## The following objects are masked from 'package:plyr':
##
##     arrange, count, desc, failwith, id, mutate, rename, summarise,
##     summarize

## The following objects are masked from 'package:data.table':
##
##     between, first, last

## The following objects are masked from 'package:stats':
##
##     filter, lag

## The following objects are masked from 'package:base':
##
##     intersect, setdiff, setequal, union

library(dbplyr)

##
## Attaching package: 'dbplyr'

## The following objects are masked from 'package:dplyr':
##
##     ident, sql

library(mice)
```

```
##
## Attaching package: 'mice'

## The following object is masked from 'package:stats':
##
##     filter

## The following objects are masked from 'package:base':
##
##     cbind, rbind

library(survey)

## Loading required package: grid

## Loading required package: Matrix

## Loading required package: survival

##
## Attaching package: 'survey'

## The following object is masked from 'package:graphics':
##
##     dotchart

library(lme4)
library(nlme)

##
## Attaching package: 'nlme'

## The following object is masked from 'package:lme4':
##
##     lmList

## The following object is masked from 'package:dplyr':
##
##     collapse

library (plm)

##
## Attaching package: 'plm'

## The following objects are masked from 'package:dplyr':
##
##     between, lag, lead

## The following object is masked from 'package:data.table':
##
##     between
```

```
library(nlme)
library(sjstats)

##
## Attaching package: 'sjstats'

## The following object is masked from 'package:survey':
##
##     cv

library(fastDummies)
library(knitr)
library(Hmisc)

## Loading required package: lattice

## Loading required package: Formula

## Loading required package: ggplot2

##
## Attaching package: 'Hmisc'

## The following object is masked from 'package:survey':
##
##     deff

## The following objects are masked from 'package:dplyr':
##
##     src, summarize

## The following objects are masked from 'package:plyr':
##
##     is.discrete, summarize

## The following objects are masked from 'package:base':
##
##     format.pval, units

library(ggpubr)

##
## Attaching package: 'ggpubr'

## The following object is masked from 'package:plyr':
##
##     mutate

library(psych)

##
## Attaching package: 'psych'
```

```
## The following object is masked from 'package:Hmisc':
##
##     describe

## The following objects are masked from 'package:ggplot2':
##
##     %+%, alpha

## The following object is masked from 'package:sjstats':
##
##     phi

library(assertthat)

##
## Attaching package: 'assertthat'

## The following object is masked from 'package:rlang':
##
##     has_name

library(broom)

##
## Attaching package: 'broom'

## The following object is masked from 'package:sjstats':
##
##     bootstrap

library(modelr)

##
## Attaching package: 'modelr'

## The following object is masked from 'package:broom':
##
##     bootstrap

## The following objects are masked from 'package:sjstats':
##
##     bootstrap, mse, rmse

library(dplR)
library(merDeriv)

## Loading required package: nonnest2

## This is nonnest2 0.5-5.
## nonnest2 has not been tested with all combinations of model classes.

## Loading required package: sandwich

## Loading required package: lavaan
```

```
## This is lavaan 0.6-9
## lavaan is FREE software! Please report any bugs.

##
## Attaching package: 'lavaan'

## The following object is masked from 'package:psych':
##
##     cor2cov

library(np)

## Nonparametric Kernel Methods for Mixed Datatypes (version 0.60-11)
## [vignette("np_faq",package="np") provides answers to frequently asked ques
tions]
## [vignette("np",package="np") an overview]
## [vignette("entropy_np",package="np") an overview of entropy-based methods]

##
## Attaching package: 'np'

## The following object is masked from 'package:sjstats':
##
##     se

library(tidyverse)

## -- Attaching packages --------------------------------------- tidyverse 1.
3.1 --

## v tibble  3.1.2     v purrr   0.3.4
## v tidyr   1.1.3     v stringr 1.4.0
## v readr   1.4.0     v forcats 0.5.1

## -- Conflicts ------------------------------------------ tidyverse_conflict
s() --
## x purrr::%@%()         masks rlang::%@%()
## x psych::%+%()         masks ggplot2::%+%()
## x data.table:::=()     masks rlang:::=()
## x psych::alpha()       masks ggplot2::alpha()
## x dplyr::arrange()     masks plyr::arrange()
## x purrr::as_function() masks rlang::as_function()
## x plm::between()       masks dplyr::between(), data.table::between()
## x modelr::bootstrap()  masks broom::bootstrap(), sjstats::bootstrap()
## x nlme::collapse()     masks dplyr::collapse()
## x purrr::compact()     masks plyr::compact()
## x dplyr::count()       masks plyr::count()
## x tidyr::expand()      masks Matrix::expand()
## x dplyr::failwith()    masks plyr::failwith()
## x mice::filter()       masks dplyr::filter(), stats::filter()
## x dplyr::first()       masks data.table::first()
## x purrr::flatten()     masks rlang::flatten()
```

```
## x purrr::flatten_chr() masks rlang::flatten_chr()
## x purrr::flatten_dbl() masks rlang::flatten_dbl()
## x purrr::flatten_int() masks rlang::flatten_int()
## x purrr::flatten_lgl() masks rlang::flatten_lgl()
## x purrr::flatten_raw() masks rlang::flatten_raw()
## x tibble::has_name()   masks assertthat::has_name(), rlang::has_name()
## x dplyr::id()          masks plyr::id()
## x dbplyr::ident()      masks dplyr::ident()
## x purrr::invoke()      masks rlang::invoke()
## x plm::lag()           masks dplyr::lag(), stats::lag()
## x dplyr::last()        masks data.table::last()
## x plm::lead()          masks dplyr::lead()
## x purrr::list_along()  masks rlang::list_along()
## x purrr::modify()      masks rlang::modify()
## x modelr::mse()        masks sjstats::mse()
## x ggpubr::mutate()     masks dplyr::mutate(), plyr::mutate()
## x tidyr::pack()        masks Matrix::pack()
## x purrr::prepend()     masks rlang::prepend()
## x dplyr::rename()      masks plyr::rename()
## x modelr::rmse()       masks sjstats::rmse()
## x purrr::splice()      masks rlang::splice()
## x dbplyr::sql()        masks dplyr::sql()
## x Hmisc::src()         masks dplyr::src()
## x dplyr::summarise()   masks plyr::summarise()
## x Hmisc::summarize()   masks dplyr::summarize(), plyr::summarize()
## x purrr::transpose()   masks data.table::transpose()
## x tidyr::unpack()      masks Matrix::unpack()
```

```
library(rms)
```

```
## Loading required package: SparseM
```

```
##
## Attaching package: 'SparseM'
```

```
## The following object is masked from 'package:base':
##
##     backsolve
```

```
##
## Attaching package: 'rms'
```

```
## The following object is masked from 'package:dplR':
##
##     rcs
```

```
## The following object is masked from 'package:survey':
##
##     calibrate
```

```
library("survey") #
library(lavaan)
```

```
library(tidyverse)
library(corrr)
library(data.table)
library(PCAmixdata)
library(apaTables)

# Load data


merged_df_original=fread("E:/ABCD/Hippo_3.0/merged_df_withedu_withsite_new.cs
v")

#Cognitive ability

merged_df_original$CA_Z= scale(merged_df_original$nihtbx_totalcomp_agecorrect
ed, center = TRUE, scale = TRUE)


#Subset to cases with only cognitive ability

merged_df <- merged_df_original[complete.cases(merged_df_original$nihtbx_tota
lcomp_agecorrected), ]
str(merged_df)

## Classes 'data.table' and 'data.frame':   9972 obs. of  2417 variables:
##  $ subjectkey                         : chr  "NDAR_INV003RTV85" "NDAR_INV00BD7V
DC" "NDAR_INV00CY2MDM" "NDAR_INV00HEV6HB" ...
##  $ interview_age                      : int  131 112 130 124 110 109 121 118 11
4 130 ...
##  $ sex                                : chr  "F" "M" "M" "M" ...
##  $ eventname.x                        : chr  "baseline_year_1_arm_1" "baseline_
year_1_arm_1" "baseline_year_1_arm_1" "baseline_year_1_arm_1" ...
##  $ anthro_1_height_in                 : num  56.5 57.5 56.5 57.3 50.9 52 53.5 5
8.4 54 52.5 ...
##  $ anthro2heightin                    : num  56.5 57.5 56.5 57.3 50.9 52 53.5 5
8.2 54 52.5 ...
##  $ anthro3heightin                    : num  NA NA NA NA NA NA NA NA NA NA ...
##  $ anthroheightcalc                   : num  56.5 57.5 56.5 57.3 50.9 52 53.5 5
8.3 54 52.5 ...
##  $ anthroweightcast                   : int  0 0 0 0 0 0 0 0 0 0 ...
##  $ anthro_weight_a_location           : chr  "" "" "" "" ...
##  $ anthroweight1lb                    : num  93 76.8 91.5 70.8 70.3 80 81.4 85.
5 64 97.1 ...
##  $ anthroweight2lb                    : num  93 76.8 91.5 71 70.2 80 81.4 85.3
64 96.9 ...
##  $ anthroweight3lb                    : num  NA NA NA 70.8 NA NA NA 85.4 NA 96.
9 ...
##  $ anthroweightcalc                   : num  93 76.8 91.5 70.9 70.2 ...
##  $ anthro_waist_cm                    : num  31 23.5 30 28 26 30 25.2 26 25.5 2
```

```
9.5 ...
## $ anthro_timestamp : chr "10/1/2018 14:16" "6/12/2018 9:46"
"8/22/2017 9:52" "7/8/2017 10:32" ...
## $ demo_l_p_select_language___1 : int 0 0 0 0 0 1 0 0 0 0 ...
## $ demo_prim_l : int 1 1 1 1 1 1 1 1 1 1 ...
## $ demo_brthdat_v2_l : num 11 10 11 11 10 10 11 10 10 11 ...
## $ demo_ed_v2_l : int 6 5 6 6 5 4 6 5 4 5 ...
## $ demo_gender_id_v2_l : int 2 1 1 1 1 1 1 2 2 2 ...
## $ demo_nat_lang_l : int 58 58 58 58 58 47 58 58 58 58 ...
## $ demo_nat_lang_2_l : int 1 1 1 1 1 4 1 1 1 1 ...
## $ demo_dual_lang_v2_l : int 0 0 0 0 0 0 0 0 0 0 ...
## $ demo_dual_lang_years_p___1 : int 0 0 0 0 0 0 0 0 0 0 ...
## $ demo_dual_lang_years_p___2 : int 0 0 0 0 0 0 0 0 0 0 ...
## $ demo_dual_lang_years_p___3 : int 0 0 0 0 0 0 0 0 0 0 ...
## $ demo_dual_lang_years_p___4 : int 0 0 0 0 0 0 0 0 0 0 ...
## $ demo_dual_lang_years_p___5 : int 0 0 0 0 0 0 0 0 0 0 ...
## $ demo_dual_lang_years_p___6 : int 0 0 0 0 0 0 0 0 0 0 ...
## $ demo_dual_lang_years_p___7 : int 0 0 0 0 0 0 0 0 0 0 ...
## $ demo_dual_lang_years_p___8 : int 0 0 0 0 0 0 0 0 0 0 ...
## $ demo_dual_lang_years_p___9 : int 0 0 0 0 0 0 0 0 0 0 ...
## $ demo_dual_lang_years_p___10 : int 0 0 0 0 0 0 0 0 0 0 ...
## $ demo_relig_v2_l : int 2 9 13 13 17 2 2 13 13 13 ...
## $ demo_prnt_age_v2_l : int 44 40 40 37 46 43 39 51 39 36 ...
## $ demo_prnt_age_v2_refuse_l : int NA NA NA NA NA NA NA NA NA NA ...
## $ demo_prnt_gender_id_v2_l : int 2 2 2 2 2 2 2 2 2 2 ...
## $ demo_prnt_race_a_v2_l___10 : int 0 0 0 0 0 0 0 1 0 0 ...
## $ demo_prnt_race_a_v2_l___11 : int 0 0 0 0 0 0 0 0 0 0 ...
## $ demo_prnt_race_a_v2_l___12 : int 0 0 0 0 0 0 0 0 0 0 ...
## $ demo_prnt_race_a_v2_l___13 : int 0 0 0 0 0 0 0 0 0 0 ...
## $ demo_prnt_race_a_v2_l___14 : int 0 0 0 0 0 0 0 0 0 0 ...
## $ demo_prnt_race_a_v2_l___15 : int 0 0 0 0 0 0 0 0 0 0 ...
## $ demo_prnt_race_a_v2_l___16 : int 0 0 0 0 0 0 0 0 0 0 ...
## $ demo_prnt_race_a_v2_l___17 : int 0 0 0 0 0 0 0 0 0 0 ...
## $ demo_prnt_race_a_v2_l___18 : int 0 0 0 0 0 0 0 0 0 0 ...
## $ demo_prnt_race_a_v2_l___19 : int 0 0 0 0 0 0 0 0 0 0 ...
## $ demo_prnt_race_a_v2_l___20 : int 0 0 0 0 0 0 0 0 0 0 ...
## $ demo_prnt_race_a_v2_l___21 : int 0 0 0 0 0 0 0 0 0 0 ...
## $ demo_prnt_race_a_v2_l___22 : int 0 0 0 0 0 0 0 0 0 0 ...
## $ demo_prnt_race_a_v2_l___23 : int 0 0 0 0 0 0 0 0 0 0 ...
## $ demo_prnt_race_a_v2_l___24 : int 0 0 0 0 0 0 0 0 0 0 ...
## $ demo_prnt_race_a_v2_l___25 : int 0 0 0 0 0 0 0 0 0 0 ...
## $ demo_prnt_race_a_v2_l___77 : int 0 0 0 0 0 0 0 0 0 0 ...
## $ demo_prnt_race_a_v2_l___99 : int 0 0 0 0 0 0 0 0 0 0 ...
## $ demo_prnt_ethn_v2_l : int 2 2 2 2 2 1 2 2 2 2 ...
## $ demo_prnt_ethn2_v2_l : int NA NA NA NA NA 18 NA NA NA NA ...
## $ demo_prnt_nat_lang_l : int 58 58 58 58 15 47 58 58 58 58 ...
## $ demo_prnt_nat_lang_2_l : int 1 1 1 1 4 4 1 1 1 1 ...
## $ demo_prnt_marital_v2_l : int 1 1 4 1 777 1 5 4 1 1 ...
## $ demo_prnt_ed_v2_l : int 13 20 15 13 21 13 14 16 19 14 ...
## $ demo_prnt_empl_v2_l : int 1 1 1 1 1 6 1 1 1 1 ...
```

```
##  $ demo_prnt_empl_time_l          : int  1 2 1 2 1 NA 1 1 1 1 ...
##  $ demo_prnt_indust_refuse_l      : int  NA NA NA NA NA NA NA NA NA NA ...
##  $ demo_prnt_income_v2_l          : int  5 4 6 999 777 4 1 6 6 6 ...
##  $ demo_prnt_prtnr_v2_l           : int  1 1 2 1 2 1 2 1 1 1 ...
##  $ demo_prnt_prtnr_bio_l          : int  1 1 NA 1 NA 1 NA 1 1 1 ...
##  $ demo_prnt_prtnr_adopt_l        : int  NA NA NA NA NA NA NA NA NA NA ...
##  $ demo_prtnr_ed_v2_l             : int  13 20 NA 13 NA 13 NA 18 18 12 ...
##  $ demo_prtnr_empl_v2_l           : int  1 1 NA 1 NA 8 NA 1 1 1 ...
##  $ demo_prtnr_empl_time_l         : int  1 1 NA 1 NA NA NA 1 1 1 ...
##  $ demo_prtnr_indust_refuse_l     : int  NA NA NA NA NA NA NA NA NA NA ...
##  $ demo_prtnr_income_v2_l         : int  8 10 NA 999 NA 7 NA 7 9 5 ...
##  $ demo_child_time_v2_l           : int  0 1 0 0 1 0 0 0 1 0 ...
##  $ demo_child_time2_v2_l          : int  NA 5 NA NA NA NA NA NA 5 NA ...
##  $ demo_child_time2_v2_dk_l       : int  NA NA NA NA 777 NA NA NA NA NA ...
##  $ demo_child_time3_v2_l          : int  NA 4 NA NA 1 NA NA NA 8 NA ...
##  $ demo_comb_income_v2_l          : int  8 10 6 999 999 7 1 8 9 7 ...
##  $ demo_roster_v2_l               : int  6 4 5 5 2 6 5 4 4 8 ...
##  $ demo_roster_v2_refuse_l        : int  NA NA NA NA NA NA NA NA NA NA ...
##  $ fam_roster_2c_v2_l             : int  1 1 3 1 3 1 3 1 1 1 ...
##  $ fam_roster_3c_v2_l             : int  3 3 3 3 NA 3 3 3 3 3 ...
##  $ fam_roster_4c_v2_l             : int  3 2 3 3 NA 3 3 3 3 3 ...
##  $ fam_roster_5c_v2_l             : int  3 NA 3 3 NA 3 3 NA NA 3 ...
##  $ fam_roster_6c_v2_l             : int  3 NA NA NA NA 3 NA NA NA 14 ...
##  $ fam_roster_7c_v2_l             : int  NA NA NA NA NA NA NA NA NA 11 ...
##  $ fam_roster_8c_v2_l             : int  NA NA NA NA NA NA NA NA NA 4 ...
##  $ fam_roster_9c_v2_l             : int  NA NA NA NA NA NA NA NA NA NA ...
##  $ fam_roster_10c_v2_l            : int  NA NA NA NA NA NA NA NA NA NA ...
##  $ fam_roster_11c_v2_l            : int  NA NA NA NA NA NA NA NA NA NA ...
##  $ fam_roster_12c_v2_l            : int  NA NA NA NA NA NA NA NA NA NA ...
##  $ fam_roster_13c_v2_l            : int  NA NA NA NA NA NA NA NA NA NA ...
##  $ fam_roster_14c_v2_l            : int  NA NA NA NA NA NA NA NA NA NA ...
##  $ fam_roster_15c_v2_l            : int  NA NA NA NA NA NA NA NA NA NA ...
##  $ demo_fam_exp1_v2_l             : int  0 0 0 0 0 0 0 0 0 1 ...
##  $ demo_fam_exp2_v2_l             : int  0 0 0 0 0 0 0 0 0 1 ...
##  $ demo_fam_exp3_v2_l             : int  0 0 0 0 0 0 0 0 0 0 ...
##  $ demo_fam_exp4_v2_l             : int  0 0 0 0 0 0 0 0 0 0 ...
##   [list output truncated]
##  - attr(*, ".internal.selfref")=<externalptr>
```

```
summary(merged_df$CA_Z)
```

```
##     Min.  1st Qu.   Median     Mean  3rd Qu.     Max.
## -3.88552 -0.67504 -0.05547  0.00000  0.62042  5.57696
```

```
#age

merged_df$age = merged_df$interview_age %>% as.numeric()



# sex numeric
```

```
merged_df$sex_numeric[merged_df$sex=="M"] <- "0"
merged_df$sex_numeric[merged_df$sex=="F"] <- "1"
merged_df$sex_n = as.numeric(merged_df$sex_numeric)

#adjust CA for sex

d2 <- lm(CA_Z ~ sex_numeric, data = merged_df) # fit the model
merged_df$CA_Z_adj <- residuals(d2)

cor(merged_df$CA_Z_adj, merged_df$CA_Z,  use = "complete.obs")

## [1] 0.9999212

summary(merged_df$CA_Z_adj)

##    Min. 1st Qu.  Median    Mean 3rd Qu.    Max.
## -3.8989 -0.6884 -0.0437  0.0000  0.6322  5.5887

#Child_US_Born
pdem02_origin <- read.delim("E:/ABCD/Hippo_3.0/Files 3.0/ABCDStudyNDA/pdem02.
txt")

pdem02_origin$Child_US_Born <- NA
pdem02_origin= mutate(pdem02_origin, Child_US_Born  = case_when(
  demo_origin_v2 %in% 189 ~ 1,
  demo_origin_v2 <189 ~ 0,
  demo_origin_v2 >189 ~ 0,
  TRUE ~ as.numeric(Child_US_Born)) # This is for all other values
)

describe(pdem02_origin$Child_US_Born)

##    vars     n mean   sd median trimmed mad min max range  skew kurtosis se
## X1    1 11879 0.97 0.17      1       1   0   0   1     1 -5.43    27.48  0

table(pdem02_origin$Child_US_Born)

##
##     0     1
##   366 11513

pdem02_origin_USB<- c("Child_US_Born", "subjectkey")
pdem02_origin_USB_df <- pdem02_origin[pdem02_origin_USB]
merged_df=merge(merged_df, pdem02_origin_USB_df, by.x="subjectkey", by.y= "su
bjectkey")
merged_df$Child_US_Born= as.numeric(merged_df$Child_US_Born)


#recode SIRE Not used in this analysis
merged_df$race <- NA
```

```
merged_df= mutate(merged_df,
                  race = case_when(
                    demo_race_a_p___10 %in% "1" ~ "White_1",
                    demo_race_a_p___11 %in% "1" ~ "Black_1",
                    demo_race_a_p___12 %in%  "1" ~ "Native American_1",
                    demo_race_a_p___13 %in%  "1" ~ "Native American_1",
                    demo_race_a_p___14 %in%  "1" ~ "Pacific Islander_1",
                    demo_race_a_p___15 %in%  "1" ~ "Pacific Islander_1",
                    demo_race_a_p___16 %in%  "1" ~ "Pacific Islander_1",
                    demo_race_a_p___17 %in%  "1" ~ "Pacific Islander_1",
                    demo_race_a_p___18 %in%  "1" ~ "SouthAsian_1",
                    demo_race_a_p___19 %in%  "1" ~ "EastAsian_1",
                    demo_race_a_p___20 %in%  "1" ~ "EastAsian_1",
                    demo_race_a_p___21  %in%  "1" ~ "EastAsian_1",
                    demo_race_a_p___22 %in%  "1" ~ "EastAsian_1",
                    demo_race_a_p___23 %in%  "1" ~ "EastAsian_1",
                    demo_race_a_p___24 %in%  "1" ~ "EastAsian_1", #other Asia
n predominantly East Asian Ancestry so marked as EA
                    demo_race_a_p___25 %in%  "1" ~ "Other Race_1", #other Rac
e
                    demo_race_a_p___77 %in%  "1" ~ "Other Race_1",
                    demo_race_a_p___99 %in%  "1" ~ "Other Race_1",

                    TRUE ~ as.character(race)) # This is for all other values
)

table(merged_df$race, exclude=NULL)

##
##           Black_1       EastAsian_1 Native American_1      Other Race_1
##              1583               130                49               488
##      SouthAsian_1           White_1              <NA>
##                49              7653                20

merged_df$race=merged_df$race %>% replace_na("Other Race_1")
table(merged_df$race, exclude=NULL)

##
##           Black_1       EastAsian_1 Native American_1      Other Race_1
##              1583               130                49               508
##      SouthAsian_1           White_1
##                49              7653

merged_df$race= as.factor(merged_df$race)
merged_df$race <- relevel(merged_df$race, ref = "White_1")


#recode dummy SIRE

merged_df$White <-0
```

```
merged_df= mutate(merged_df,
                  White= case_when(
                    demo_race_a_p___10 == 1 ~ 1,
                    TRUE ~ as.numeric(White)) # This is for all other values
)

describe(merged_df$White)

##    vars    n mean   sd median trimmed mad min max range  skew kurtosis se
## X1    1 9972 0.77 0.42      1    0.83   0   0   1     1 -1.27     -0.4  0

summary(merged_df$White)

##    Min. 1st Qu.  Median    Mean 3rd Qu.    Max.
##  0.0000  1.0000  1.0000  0.7674  1.0000  1.0000

table(merged_df$White,exclude=NULL)

##
##    0    1
## 2319 7653

merged_df$Black <-0
merged_df= mutate(merged_df,
                  Black= case_when(
                    demo_race_a_p___11 == 1 ~ 1,
                    TRUE ~ as.numeric(Black)) # This is for all other values
)

describe(merged_df$Black)

##    vars    n mean  sd median trimmed mad min max range skew kurtosis se
## X1    1 9972  0.2 0.4      0    0.13   0   0   1     1 1.48     0.19  0

summary(merged_df$Black)

##    Min. 1st Qu.  Median    Mean 3rd Qu.    Max.
##  0.0000  0.0000  0.0000  0.2027  0.0000  1.0000

table(merged_df$Black, exclude=NULL)

##
##    0    1
## 7951 2021

merged_df$EastAsian <-0
merged_df= mutate(merged_df,
                  EastAsian= case_when(
                    demo_race_a_p___19 == 1 ~ 1,
                    demo_race_a_p___20 == 1 ~ 1,
                    demo_race_a_p___21 == 1 ~ 1,
                    demo_race_a_p___22 == 1 ~ 1,
```

```
                    demo_race_a_p___23 == 1 ~ 1,
                    demo_race_a_p___24 == 1 ~ 1,
                    TRUE ~ as.numeric(EastAsian)) # This is for all other val
ues
)

describe(merged_df$EastAsian)

##    vars    n mean   sd median trimmed mad min max range skew kurtosis se
## X1    1 9972 0.05 0.21      0       0   0   0   1     1 4.25    16.04  0

summary(merged_df$EastAsian)

##    Min. 1st Qu.  Median    Mean 3rd Qu.    Max.
## 0.00000 0.00000 0.00000 0.04763 0.00000 1.00000

table(merged_df$EastAsian, exclude=NULL)

##
##    0    1
## 9497  475

merged_df$Native_American <-0
merged_df= mutate(merged_df,
                  Native_American= case_when(
                    demo_race_a_p___12 == 1 ~ 1,
                    demo_race_a_p___13 == 1 ~ 1,
                    TRUE ~ as.numeric(Native_American)) # This is for all oth
er values
)

describe(merged_df$Native_American)

##    vars    n mean   sd median trimmed mad min max range skew kurtosis se
## X1    1 9972 0.03 0.18      0       0   0   0   1     1 5.09    23.93  0

summary(merged_df$Native_American)

##    Min. 1st Qu.  Median    Mean 3rd Qu.    Max.
##  0.0000  0.0000  0.0000  0.0346  0.0000  1.0000

table(merged_df$Native_American)

##
##    0    1
## 9627  345

merged_df$SouthAsian <-0
merged_df= mutate(merged_df,
                  SouthAsian= case_when(
                    demo_race_a_p___18 == 1 ~ 1,
                    TRUE ~ as.numeric(SouthAsian)) # This is for all other va
```

```
lues
)

describe(merged_df$SouthAsian)

##    vars    n mean  sd median trimmed mad min max range  skew kurtosis se
## X1    1 9972 0.01 0.1      0       0   0   0   1     1 10.04    98.86  0

summary(merged_df$SouthAsian)

##     Min.  1st Qu.   Median     Mean  3rd Qu.     Max. 
## 0.000000 0.000000 0.000000 0.009627 0.000000 1.000000

table(merged_df$SouthAsian)

## 
##    0    1 
## 9876   96

merged_df$Other_Race <-0
merged_df= mutate(merged_df,
                  Other_Race= case_when(
                    race == "Other Race_1" ~ 1,
                    TRUE ~ as.numeric(Other_Race)) # This is for all other va
lues
)



describe(merged_df$Other_Race)

##    vars    n mean   sd median trimmed mad min max range skew kurtosis se
## X1    1 9972 0.05 0.22      0       0   0   0   1     1 4.08    14.68  0

summary(merged_df$Other_Race)

##    Min. 1st Qu.  Median    Mean 3rd Qu.    Max. 
## 0.00000 0.00000 0.00000 0.05094 0.00000 1.00000

table(merged_df$Other_Race, exclude=NULL)

## 
##    0    1 
## 9464  508

#add Hispanic # define as people who are only positively identified as Hispan
ic


merged_df$Hispanic <-0
merged_df= mutate(merged_df,
                  Hispanic= case_when(
```

```
                         demo_ethn_v2 ==1 ~ 1,
                         TRUE ~ as.numeric(Hispanic)) # This is for all other valu
es
)

describe(merged_df$Hispanic)

##    vars    n mean   sd median trimmed mad min max range skew kurtosis se
## X1    1 9972 0.19 0.39      0    0.11   0   0   1     1  1.6     0.57  0

summary(merged_df$Hispanic)

##    Min. 1st Qu.  Median    Mean 3rd Qu.    Max. 
##  0.0000  0.0000  0.0000  0.1874  0.0000  1.0000

table(merged_df$Hispanic, exclude=NULL)

## 
##    0    1 
## 8103 1869

#non-Hispanic categories

merged_df$NH_White_only <-0
merged_df= mutate(merged_df,
                       NH_White_only= case_when(
                         White == 1 & !Black == 1 & !EastAsian ==1 & !Native_Ameri
can ==1 & !SouthAsian ==1 & !Other_Race ==1
                         & !Hispanic ==1  ~ 1,
                         TRUE ~ as.numeric(NH_White_only)) # This is for all other
values
)
describe(merged_df$NH_White_only)

##    vars    n mean  sd median trimmed mad min max range  skew kurtosis se
## X1    1 9972 0.55 0.5      1    0.57   0   0   1     1 -0.22    -1.95  0

summary(merged_df$NH_White_only)

##    Min. 1st Qu.  Median    Mean 3rd Qu.    Max. 
##  0.0000  0.0000  1.0000  0.5549  1.0000  1.0000

table(merged_df$NH_White_only)

## 
##    0    1 
## 4439 5533

merged_df$NH_Black_only <-0
merged_df= mutate(merged_df,
                       NH_Black_only= case_when(
                         Black == 1 & !White == 1 &!EastAsian ==1 & !Native_Americ
```

```
an ==1 & !SouthAsian ==1 & !Other_Race ==1
                    & !Hispanic ==1  ~ 1,
                    TRUE ~ as.numeric(NH_Black_only)) # This is for all other
values
)
describe(merged_df$NH_Black_only)

##    vars    n mean   sd median trimmed mad min max range skew kurtosis se
## X1    1 9972 0.14 0.35      0    0.05   0   0   1     1 2.03     2.12  0

summary(merged_df$NH_Black_only)

##    Min. 1st Qu.  Median    Mean 3rd Qu.    Max.
##  0.0000  0.0000  0.0000  0.1438  0.0000  1.0000

table(merged_df$NH_Black_only)

##
##    0    1
## 8538 1434

merged_df$NH_EastAsian_only <-0
merged_df= mutate(merged_df,
                  NH_EastAsian_only= case_when(
                    !Black == 1 & !White == 1 & EastAsian ==1 & !Native_Ameri
can ==1 & !SouthAsian ==1 & !Other_Race ==1
                    & !Hispanic ==1  ~ 1,
                    TRUE ~ as.numeric(NH_EastAsian_only)) # This is for all o
ther values
)
describe(merged_df$NH_EastAsian_only)

##    vars    n mean  sd median trimmed mad min max range skew kurtosis se
## X1    1 9972 0.01 0.1      0       0   0   0   1     1  9.5    88.19  0

summary(merged_df$NH_EastAsian_only)

##    Min. 1st Qu.  Median    Mean 3rd Qu.    Max.
## 0.00000 0.00000 0.00000 0.01073 0.00000 1.00000

table(merged_df$NH_EastAsian_only)

##
##    0    1
## 9865  107

merged_df$NH_SouthAsian_only <-0
merged_df= mutate(merged_df,
                  NH_SouthAsian_only= case_when(
                    !Black == 1 & !White == 1 & !EastAsian ==1 & !Native_Amer
ican ==1 & SouthAsian ==1 & !Other_Race ==1
                    & !Hispanic ==1  ~ 1,
```

```
                    TRUE ~ as.numeric(NH_SouthAsian_only)) # This is for all
other values
)
describe(merged_df$NH_SouthAsian_only)

##    vars    n mean   sd median trimmed mad min max range  skew kurtosis se
## X1    1 9972    0 0.07      0       0   0   0   1     1 15.13   226.87  0

summary(merged_df$NH_SouthAsian_only)

##     Min.  1st Qu.   Median     Mean  3rd Qu.     Max.
## 0.000000 0.000000 0.000000 0.004312 0.000000 1.000000

table(merged_df$NH_SouthAsian_only)

##
##    0    1
## 9929   43

merged_df$NH_Native_American_only <-0
merged_df= mutate(merged_df,
                  NH_Native_American_only= case_when(
                    !Black == 1 & !White == 1 & !EastAsian ==1 & Native_Ameri
can ==1 & !SouthAsian ==1 & !Other_Race ==1
                    & !Hispanic ==1  ~ 1,
                    TRUE ~ as.numeric(NH_Native_American_only)) # This is for
all other values
)
describe(merged_df$NH_Native_American_only)

##    vars    n mean   sd median trimmed mad min max range  skew kurtosis se
## X1    1 9972    0 0.06      0       0   0   0   1     1 17.85   316.62  0

summary(merged_df$NH_Native_American_only)

##     Min.  1st Qu.   Median     Mean  3rd Qu.     Max.
## 0.000000 0.000000 0.000000 0.003109 0.000000 1.000000

table(merged_df$NH_Native_American_only)

##
##    0    1
## 9941   31

merged_df$NH_Other_Race_only <-0
merged_df= mutate(merged_df,
                  NH_Other_Race_only= case_when(
                    !Black == 1 & !White == 1 & !EastAsian ==1 & !Native_Amer
ican ==1 & !SouthAsian ==1 & Other_Race ==1
                    & !Hispanic ==1  ~ 1,
                    TRUE ~ as.numeric(NH_Other_Race_only)) # This is for all
other values
```

```
)
describe(merged_df$NH_Other_Race_only)

##    vars    n mean  sd median trimmed mad min max range skew kurtosis se
## X1    1 9972 0.01 0.1      0       0   0   0   1     1 9.99    97.79  0

summary(merged_df$NH_Other_Race_only)

##     Min.  1st Qu.   Median     Mean  3rd Qu.     Max. 
## 0.000000 0.000000 0.000000 0.009727 0.000000 1.000000

table(merged_df$NH_Other_Race_only)

## 
##    0    1 
## 9875   97

#Mixed categories


merged_df$NH_Black_White_only <-0
merged_df= mutate(merged_df,
                  NH_Black_White_only= case_when(
                    Black == 1 & White == 1 & !EastAsian ==1 & !Native_Americ
an ==1 & !SouthAsian ==1 & !Other_Race ==1
                    & !Hispanic ==1  ~ 1,
                    TRUE ~ as.numeric(NH_Black_White_only)) # This is for all
other values
)

describe(merged_df$NH_Black_White_only)

##    vars    n mean   sd median trimmed mad min max range skew kurtosis se
## X1    1 9972 0.03 0.17      0       0   0   0   1     1 5.48    28.04  0

table(merged_df$NH_Black_White_only)

## 
##    0    1 
## 9670  302

merged_df$H_Black_White_only <-0
merged_df= mutate(merged_df,
                  H_Black_White_only= case_when(
                    Black == 1 & White == 1 & !EastAsian ==1 & !Native_Americ
an ==1 & !SouthAsian ==1 & !Other_Race ==1
                    & Hispanic ==1  ~ 1,
                    TRUE ~ as.numeric(H_Black_White_only)) # This is for all
other values
)

describe(merged_df$H_Black_White_only)
```

```
##    vars    n mean   sd median trimmed mad min max range  skew kurtosis se
## X1    1 9972 0.01 0.07      0       0   0   0   1     1 14.01   194.41  0

table(merged_df$H_Black_White_only)

##
##    0    1
## 9922   50

merged_df$H_Black_only <-0
merged_df= mutate(merged_df,
                  H_Black_only= case_when(
                    Black == 1 & !White ==1 & !EastAsian ==1 & !Native_Americ
an ==1 & !SouthAsian ==1 & !Other_Race ==1
                    & Hispanic ==1  ~ 1,
                    TRUE ~ as.numeric(H_Black_only)) # This is for all other
values
)

describe(merged_df$H_Black_only)

##    vars    n mean   sd median trimmed mad min max range  skew kurtosis se
## X1    1 9972 0.01 0.09      0       0   0   0   1     1 10.76    113.7  0

table(merged_df$H_Black_only)

##
##    0    1
## 9888   84

merged_df$H_White_only <-0
merged_df= mutate(merged_df,
                  H_White_only= case_when(
                    !Black == 1 & White ==1 & !EastAsian ==1 & !Native_Americ
an ==1 & !SouthAsian ==1 & !Other_Race ==1
                    & Hispanic ==1  ~ 1,
                    TRUE ~ as.numeric(H_White_only)) # This is for all other
values
)

describe(merged_df$H_White_only)

##    vars    n mean   sd median trimmed mad min max range skew kurtosis se
## X1    1 9972 0.12 0.32      0    0.02   0   0   1     1 2.38     3.65  0

table(merged_df$H_White_only)

##
##    0    1
## 8801 1171
```

```
merged_df$NH_EastAsian_White_only <-0
merged_df= mutate(merged_df,
                  NH_EastAsian_White_only = case_when(
                    EastAsian == 1 & White == 1 & !Black ==1 & !Native_Americ
an ==1 & !SouthAsian ==1 & !Other_Race ==1
                    & !Hispanic ==1  ~ 1,
                    TRUE ~ as.numeric(NH_EastAsian_White_only)) # This is for
all other values
)

describe(merged_df$NH_EastAsian_White_only)

##    vars    n mean   sd median trimmed mad min max range skew kurtosis se
## X1    1 9972 0.02 0.16      0       0   0   0   1     1 6.09    35.07  0

table(merged_df$NH_EastAsian_White_only)

##
##    0    1
## 9723  249

merged_df$NH_SouthAsian_White_only <-0
merged_df= mutate(merged_df,
                  NH_SouthAsian_White_only = case_when(
                    SouthAsian == 1 & White == 1 & !Black ==1 & !Native_Ameri
can ==1 & !EastAsian ==1 & !Other_Race ==1
                    & !Hispanic ==1  ~ 1,
                    TRUE ~ as.numeric(NH_SouthAsian_White_only)) # This is fo
r all other values
)

describe(merged_df$NH_SouthAsian_White_only)

##    vars    n mean   sd median trimmed mad min max range  skew kurtosis se
## X1    1 9972    0 0.06      0       0   0   0   1     1 15.69   244.25  0

table(merged_df$NH_SouthAsian_White_only)

##
##    0    1
## 9932   40

merged_df$NH_Native_American_White_only <-0
merged_df= mutate(merged_df,
                  NH_Native_American_White_only = case_when(
                    Native_American == 1 & White == 1 & !Black ==1 & !SouthAs
ian ==1 & !EastAsian ==1 & !Other_Race ==1
                    & !Hispanic ==1  ~ 1,
                    TRUE ~ as.numeric(NH_Native_American_White_only)) # This
is for all other values
)
```

```
describe(merged_df$NH_Native_American_White_only)

##    vars    n mean   sd median trimmed mad min max range skew kurtosis se
## X1    1 9972 0.01 0.11      0       0   0   0   1     1 8.55    71.12  0

table(merged_df$NH_Native_American_White_only)

##
##    0    1
## 9841  131

merged_df$H_Native_American_White_only <-0
merged_df= mutate(merged_df,
                  H_Native_American_White_only = case_when(
                    Native_American == 1 & White == 1 & !Black ==1 & !SouthAs
ian ==1 & !EastAsian ==1 & !Other_Race ==1
                    & Hispanic ==1  ~ 1,
                    TRUE ~ as.numeric(H_Native_American_White_only)) # This i
s for all other values
)

describe(merged_df$H_Native_American_White_only)

##    vars    n mean   sd median trimmed mad min max range  skew kurtosis se
## X1    1 9972    0 0.07      0       0   0   0   1     1 14.62   211.74  0

table(merged_df$H_Native_American_White_only)

##
##    0    1
## 9926   46

merged_df$H_Other_only <-0
merged_df= mutate(merged_df,
                  H_Other_only = case_when(
                    Other_Race == 1 & !White == 1& !Black ==1 & !SouthAsian =
=1 & !EastAsian ==1 & !Native_American ==1
                    & Hispanic ==1  ~ 1,
                    TRUE ~ as.numeric(H_Other_only)) # This is for all other
values
)

describe(merged_df$H_Other_only)

##    vars    n mean  sd median trimmed mad min max range skew kurtosis se
## X1    1 9972 0.04 0.2      0       0   0   0   1     1 4.62     19.3  0

table(merged_df$H_Other_only)
```

```
##
##    0    1
## 9561  411
```

```
merged_df$NH_Native_American_Black <-0
merged_df= mutate(merged_df,
                  NH_Native_American_Black = case_when(
                    Native_American == 1 & Black == 1 & !White ==1 & !SouthAs
ian ==1 & !EastAsian ==1 & !Other_Race ==1
                    & !Hispanic ==1  ~ 1,
                    TRUE ~ as.numeric(NH_Native_American_Black)) # This is fo
r all other values
)

describe(merged_df$NH_Native_American_Black)
```

```
##    vars    n mean   sd median trimmed mad min max range skew kurtosis se
## X1    1 9972    0 0.05      0       0   0   0   1     1 19.5   378.46  0
```

```
table(merged_df$NH_Native_American_Black)
```

```
##
##    0    1
## 9946   26
```

```
merged_df$H_Native_American <-0
merged_df= mutate(merged_df,
                  H_Native_American = case_when(
                    Native_American == 1 & !Black == 1 & !White ==1 & !SouthA
sian ==1 & !EastAsian ==1 & !Other_Race ==1
                    & !Hispanic ==1  ~ 1,
                    TRUE ~ as.numeric(H_Native_American)) # This is for all o
ther values
)

describe(merged_df$H_Native_American)
```

```
##    vars    n mean   sd median trimmed mad min max range  skew kurtosis se
## X1    1 9972    0 0.06      0       0   0   0   1     1 17.85   316.62  0
```

```
table(merged_df$H_Native_American)
```

```
##
##    0    1
## 9941   31
```

```
merged_df$White_only <-0
merged_df= mutate(merged_df,
                  White_only= case_when(
                    White == 1 & !Black ==1 & !EastAsian ==1 & !Native_Americ
an ==1 & !SouthAsian ==1 & !Other_Race ==1  ~ 1,
                    TRUE ~ as.numeric(White_only)) # This is for all other va
```

```
lues
)

describe(merged_df$White_only)

##    vars    n mean   sd median trimmed mad min max range  skew kurtosis se
## X1    1 9972 0.67 0.47      1    0.72   0   0   1     1 -0.73    -1.46  0

summary(merged_df$White_only)

##    Min. 1st Qu.  Median    Mean 3rd Qu.    Max.
##  0.0000  0.0000  1.0000  0.6723  1.0000  1.0000

table(merged_df$White_only)

##
##    0    1
## 3268 6704

merged_df$Black_only <-0
merged_df= mutate(merged_df,
                  Black_only= case_when(
                    Black == 1 & !White ==1 & !EastAsian ==1 & !Native_Americ
an ==1 & !SouthAsian ==1 & !Other_Race ==1  ~ 1,
                    TRUE ~ as.numeric(Black_only)) # This is for all other va
lues
)

describe(merged_df$Black_only)

##    vars    n mean   sd median trimmed mad min max range skew kurtosis se
## X1    1 9972 0.15 0.36      0    0.07   0   0   1     1 1.94     1.75  0

summary(merged_df$Black_only)

##    Min. 1st Qu.  Median    Mean 3rd Qu.    Max.
##  0.0000  0.0000  0.0000  0.1522  0.0000  1.0000

table(merged_df$Black_only)

##
##    0    1
## 8454 1518

merged_df$EastAsian_only <-0
merged_df= mutate(merged_df,
                  EastAsian_only= case_when(
                    EastAsian == 1 & !Black ==1 & !White ==1 & !Native_Americ
an ==1 & !SouthAsian ==1 & !Other_Race ==1  ~ 1,
                    TRUE ~ as.numeric(EastAsian_only)) # This is for all othe
r values
)
```

```
describe(merged_df$EastAsian_only)

##    vars    n mean   sd median trimmed mad min max range skew kurtosis se
## X1    1 9972 0.01 0.11      0       0   0   0   1     1 8.58    71.71  0

summary(merged_df$EastAsian_only)

##    Min. 1st Qu.  Median    Mean 3rd Qu.    Max.
## 0.00000 0.00000 0.00000 0.01304 0.00000 1.00000

table(merged_df$EastAsian_only)

##
##    0    1
## 9842  130

merged_df$Native_American_only <-0
merged_df= mutate(merged_df,
                  Native_American_only= case_when(
                    Native_American == 1 & !Black ==1 & !White ==1 & !EastAsi
an ==1 & !SouthAsian ==1 & !Other_Race ==1 ~ 1,
                    TRUE ~ as.numeric(Native_American_only)) # This is for al
l other values
)

describe(merged_df$Native_American_only)

##    vars    n mean   sd median trimmed mad min max range  skew kurtosis se
## X1    1 9972    0 0.07      0       0   0   0   1     1 14.46   207.13  0

summary(merged_df$Native_American_only)

##     Min.  1st Qu.   Median     Mean  3rd Qu.     Max.
## 0.000000 0.000000 0.000000 0.004713 0.000000 1.000000

table(merged_df$Native_American_only)

##
##    0    1
## 9925   47

merged_df$SouthAsian_only <-0
merged_df= mutate(merged_df,
                  SouthAsian_only= case_when(
                    SouthAsian == 1 & !Black ==1 & !White ==1 & !EastAsian ==
1 & !Native_American ==1 & !Other_Race ==1   ~ 1,
                    TRUE ~ as.numeric(SouthAsian_only)) # This is for all oth
er values
)

describe(merged_df$SouthAsian_only)
```

```
##    vars    n mean   sd median trimmed mad min max range  skew kurtosis se
## X1    1 9972    0 0.07      0       0   0   0   1     1 14.62   211.74  0

summary(merged_df$SouthAsian_only)

##     Min.  1st Qu.   Median     Mean  3rd Qu.     Max. 
## 0.000000 0.000000 0.000000 0.004613 0.000000 1.000000

table(merged_df$SouthAsian_only)

## 
##    0    1 
## 9926   46

merged_df$Hispanic_only <-0
merged_df= mutate(merged_df,
                  Hispanic_only= case_when(
                    Hispanic ==1 & !SouthAsian == 1 & !Black ==1 & !White ==1
& !EastAsian ==1 & !Native_American ==1 & !Other_Race ==1   ~ 1,
                    TRUE ~ as.numeric(Hispanic_only)) # This is for all other
values
)

describe(merged_df$Hispanic_only)

##    vars    n mean sd median trimmed mad min max range skew kurtosis se
## X1    1 9972    0  0      0       0   0   0   0     0  NaN      NaN  0

summary(merged_df$Hispanic_only)

##    Min. 1st Qu.  Median    Mean 3rd Qu.    Max. 
##       0       0       0       0       0       0

table(merged_df$Hispanic_only)

## 
##    0 
## 9972

merged_df$Any_Other <-1
merged_df= mutate(merged_df,
                  Any_Other= case_when(
                    Hispanic ==1 ~ 0,
                    NH_White_only ==1~ 0,
                    NH_Black_only ==1~ 0,
                    NH_EastAsian_only ==1~ 0,
                    NH_SouthAsian_only ==1~ 0,
                    NH_Native_American_only ==1~ 0,
                    NH_Other_Race_only ==1~ 0,
                    NH_Black_White_only ==1~ 0,
                    NH_EastAsian_White_only ==1~ 0,
                    NH_SouthAsian_White_only ==1~ 0,
```

```
                    NH_Native_American_White_only ==1~ 0,

                    TRUE ~ as.numeric(Any_Other)) # This is for all other val
ues
)

describe(merged_df$Any_Other)

##    vars    n mean   sd median trimmed mad min max range skew kurtosis se
## X1    1 9972 0.01 0.12      0       0   0   0   1     1 8.39    68.32  0

summary(merged_df$Any_Other)

##    Min. 1st Qu.  Median    Mean 3rd Qu.    Max.
## 0.00000 0.00000 0.00000 0.01364 0.00000 1.00000

table(merged_df$Any_Other)

##
##    0    1
## 9836  136

#Interval SIRE_regular


merged_df$sum_SIRE = merged_df$Black + merged_df$White + merged_df$EastAsian
+ merged_df$Native_American + merged_df$SouthAsian + merged_df$Other_Race + m
erged_df$Hispanic
summary(merged_df$sum_SIRE)

##    Min. 1st Qu.  Median    Mean 3rd Qu.    Max.
##     1.0     1.0     1.0     1.3     2.0     5.0

describe(merged_df$sum_SIRE)

##    vars    n mean   sd median trimmed mad min max range skew kurtosis   se
## X1    1 9972  1.3 0.52      1    1.22   0   1   5     4 1.63     2.63 0.01

table(merged_df$sum_SIRE)

##
##    1    2    3    4    5
## 7245 2487  214   24    2

merged_df$frac_Black_SIRE <-NA
merged_df$frac_Black_SIRE = merged_df$Black / merged_df$sum_SIRE
merged_df$frac_Black_SIRE[is.nan(merged_df$frac_Black_SIRE)]<-0
summary(merged_df$frac_Black_SIRE)

##    Min. 1st Qu.  Median    Mean 3rd Qu.    Max.
##  0.0000  0.0000  0.0000  0.1705  0.0000  1.0000

describe(merged_df$frac_Black_SIRE)
```

```
##    vars    n mean   sd median trimmed mad min max range skew kurtosis se
## X1    1 9972 0.17 0.36      0    0.09   0   0   1     1 1.76     1.28  0

table(merged_df$frac_Black_SIRE)

##
##                 0               0.2              0.25 0.333333333333333
##              7951                 2                21               126
##               0.5                 1
##               438              1434

merged_df$frac_White_SIRE <-NA
merged_df$frac_White_SIRE = merged_df$White / merged_df$sum_SIRE
merged_df$frac_White_SIRE[is.nan(merged_df$frac_White_SIRE)]<-0
summary(merged_df$frac_White_SIRE)

##    Min. 1st Qu.  Median    Mean 3rd Qu.    Max.
##  0.0000  0.3333  1.0000  0.6571  1.0000  1.0000

describe(merged_df$frac_White_SIRE)

##    vars    n mean   sd median trimmed mad min max range  skew kurtosis se
## X1    1 9972 0.66 0.42      1     0.7   0   0   1     1 -0.63    -1.28  0

table(merged_df$frac_White_SIRE)

##
##                 0               0.2              0.25 0.333333333333333
##              2319                 2                24               201
##               0.5                 1
##              1893              5533

merged_df$frac_EastAsian_SIRE <-NA
merged_df$frac_EastAsian_SIRE = merged_df$EastAsian / merged_df$sum_SIRE
merged_df$frac_EastAsian_SIRE[is.nan(merged_df$frac_EastAsian_SIRE)]<-0
summary(merged_df$frac_EastAsian_SIRE)

##    Min. 1st Qu.  Median    Mean 3rd Qu.    Max.
## 0.00000 0.00000 0.00000 0.02793 0.00000 1.00000

describe(merged_df$frac_EastAsian_SIRE)

##    vars    n mean   sd median trimmed mad min max range skew kurtosis se
## X1    1 9972 0.03 0.13      0       0   0   0   1     1 5.38    30.71  0

table(merged_df$frac_EastAsian_SIRE)

##
##                 0               0.2              0.25 0.333333333333333
##              9497                 2                10                56
##               0.5                 1
##               300               107
```

```
merged_df$frac_Native_American_SIRE <-NA
merged_df$frac_Native_American_SIRE = merged_df$Native_American / merged_df$s
um_SIRE
merged_df$frac_Native_American_SIRE[is.nan(merged_df$frac_Native_American_SIR
E)]<-0
summary(merged_df$frac_Native_American_SIRE)

##    Min. 1st Qu.  Median    Mean 3rd Qu.    Max.
## 0.00000 0.00000 0.00000 0.01632 0.00000 1.00000

describe(merged_df$frac_Native_American_SIRE)

##    vars    n mean   sd median trimmed mad min max range skew kurtosis se
## X1    1 9972 0.02 0.09      0       0   0   0   1     1 6.63    50.29  0

table(merged_df$frac_Native_American_SIRE)

##
##                0               0.2              0.25 0.333333333333333
##             9627                 2                22               115
##              0.5                 1
##              175                31

merged_df$frac_SouthAsian_SIRE <-NA
merged_df$frac_SouthAsian_SIRE = merged_df$SouthAsian / merged_df$sum_SIRE
merged_df$frac_SouthAsian_SIRE[is.nan(merged_df$frac_SouthAsian_SIRE)]<-0
summary(merged_df$frac_SouthAsian_SIRE)

##     Min.  1st Qu.   Median     Mean  3rd Qu.     Max.
## 0.000000 0.000000 0.000000 0.006903 0.000000 1.000000

describe(merged_df$frac_SouthAsian_SIRE)

##    vars    n mean   sd median trimmed mad min max range  skew kurtosis se
## X1    1 9972 0.01 0.07      0       0   0   0   1     1 11.71   143.38  0

table(merged_df$frac_SouthAsian_SIRE)

##
##                0 0.333333333333333               0.5                 1
##             9876                 4                49                43

merged_df$frac_Other_SIRE <-NA
merged_df$frac_Other_SIRE = merged_df$Other_Race / merged_df$sum_SIRE
merged_df$frac_Other_SIRE[is.nan(merged_df$frac_Other_SIRE)]<-0
summary(merged_df$frac_Other_SIRE)

##    Min. 1st Qu.  Median    Mean 3rd Qu.    Max.
## 0.00000 0.00000 0.00000 0.03033 0.00000 1.00000

describe(merged_df$frac_Other_SIRE)
```

```
##    vars    n mean   sd median trimmed mad min max range skew kurtosis se
## X1    1 9972 0.03 0.14      0       0   0   0   1     1 4.96    26.03  0

table(merged_df$frac_Other_SIRE)

## 
##    0  0.5    1 
## 9464  411   97

merged_df$frac_Hispanic_SIRE <-NA
merged_df$frac_Hispanic_SIRE = merged_df$Hispanic / merged_df$sum_SIRE
merged_df$frac_Hispanic_SIRE[is.nan(merged_df$frac_Hispanic_SIRE)]<-0
summary(merged_df$frac_Hispanic_SIRE)

##    Min. 1st Qu.  Median    Mean 3rd Qu.    Max. 
## 0.00000 0.00000 0.00000 0.09084 0.00000 0.50000

describe(merged_df$frac_Hispanic_SIRE)

##    vars    n mean   sd median trimmed mad min max range skew kurtosis se
## X1    1 9972 0.09 0.19      0    0.05   0   0 0.5   0.5 1.64     0.73  0

table(merged_df$frac_Hispanic_SIRE)

## 
##                 0               0.2              0.25 0.333333333333333 
##              8103                 2                19               140 
##               0.5 
##              1708

#See if add up to 1.00

merged_df$frac_SIRE_SIRE = merged_df$frac_Black_SIRE + merged_df$frac_White_S
IRE + merged_df$frac_EastAsian_SIRE + merged_df$frac_Native_American_SIRE + m
erged_df$frac_SouthAsian_SIRE + merged_df$frac_Other_SIRE + merged_df$frac_Hi
spanic_SIRE

table(merged_df$frac_SIRE_SIRE)

## 
##    1 
## 9972

#Interval SIRE with combinations, code 0 when combinattion  is 1.


merged_df$common_combination <- 0
merged_df= mutate (merged_df,
                   common_combination = case_when(
                     NH_Black_White_only == "1" ~ 1,
                     NH_SouthAsian_White_only == "1" ~ 1,
                     NH_EastAsian_White_only == "1" ~ 1,
```

```
                     NH_Native_American_White_only == "1" ~ 1,
                     H_White_only == "1" ~ 1,
                     H_Black_only == "1" ~ 1,
                     H_Other_only == "1" ~ 1,
                     TRUE ~ as.numeric(common_combination)) # This is for all
other values
)

table(merged_df$common_combination)

##
##    0    1
## 7584 2388

describe(merged_df$common_combination)

##    vars    n mean   sd median trimmed mad min max range skew kurtosis  se
## X1    1 9972 0.24 0.43      0    0.17   0   0   1     1 1.22    -0.51   0

merged_df$Black_woc <- 0
merged_df= mutate (merged_df,
                   Black_woc = case_when(
                     Black == 1 & !common_combination == 1 ~ 1,
                     TRUE ~ as.numeric(Black_woc)) # This is for all other va
lues
)

table(merged_df$Black)

##
##    0    1
## 7951 2021

table(merged_df$Black_woc)

##
##    0    1
## 8337 1635

merged_df$White_woc <- 0
merged_df= mutate (merged_df,
                   White_woc = case_when(
                     White == 1 & !common_combination == 1 ~ 1,
                     TRUE ~ as.numeric(White_woc)) # This is for all other va
lues
)

table(merged_df$White)

##
##    0    1
## 2319 7653
```

```
table(merged_df$White_woc)

##
##    0    1
## 4212 5760

merged_df$EastAsian_woc <- 0
merged_df= mutate (merged_df,
                   EastAsian_woc = case_when(
                     EastAsian == 1 & !common_combination == 1 ~ 1,
                     TRUE ~ as.numeric(EastAsian_woc)) # This is for all othe
r values
)

table(merged_df$EastAsian)

##
##    0    1
## 9497  475

table(merged_df$EastAsian_woc)

##
##    0    1
## 9746  226

merged_df$SouthAsian_woc <- 0
merged_df= mutate (merged_df,
                   SouthAsian_woc = case_when(
                     SouthAsian == 1 & !common_combination == 1 ~ 1,
                     TRUE ~ as.numeric(SouthAsian_woc)) # This is for all oth
er values
)

table(merged_df$SouthAsian)

##
##    0    1
## 9876   96

table(merged_df$SouthAsian_woc)

##
##    0    1
## 9916   56

merged_df$Native_American_woc <- 0
merged_df= mutate (merged_df,
                   Native_American_woc = case_when(
                     Native_American == 1 & !common_combination == 1 ~ 1,
                     TRUE ~ as.numeric(Native_American_woc)) # This is for al
l other values
```

```
)

table(merged_df$Native_American)

##
##    0    1
## 9627  345

table(merged_df$Native_American_woc)

##
##    0    1
## 9758  214

merged_df$Native_American_woc <- 0
merged_df= mutate (merged_df,
                   Native_American_woc = case_when(
                     Native_American == 1 & !common_combination == 1 ~ 1,
                     TRUE ~ as.numeric(Native_American_woc)) # This is for al
l other values
)

table(merged_df$Native_American)

##
##    0    1
## 9627  345

table(merged_df$Native_American_woc)

##
##    0    1
## 9758  214

merged_df$Other_Race_woc <- 0
merged_df= mutate (merged_df,
                   Other_Race_woc = case_when(
                     Other_Race == 1 & !common_combination == 1 ~ 1,
                     TRUE ~ as.numeric(Other_Race_woc)) # This is for all oth
er values
)

table(merged_df$Other_Race)

##
##    0    1
## 9464  508

table(merged_df$Other_Race_woc)
```

```
##
##    0    1
## 9875   97
```

```
merged_df$Hispanic_woc <- 0
merged_df= mutate (merged_df,
                   Hispanic_woc = case_when(
                     Hispanic == 1 & !common_combination == 1 ~ 1,
                     TRUE ~ as.numeric(Hispanic_woc)) # This is for all other
values
)

table(merged_df$Hispanic)
```

```
##
##    0    1
## 8103 1869
```

```
table(merged_df$Hispanic_woc)
```

```
##
##    0    1
## 9769  203
```

```
merged_df$sum_SIRE_woc <- NA
merged_df$sum_SIRE_woc = merged_df$Black_woc + merged_df$White_woc + merged_d
f$EastAsian_woc + merged_df$Native_American_woc + merged_df$SouthAsian_woc +
merged_df$Other_Race_woc + merged_df$Hispanic_woc
merged_df$sum_SIRE_woc  = as.numeric(merged_df$sum_SIRE_woc)

summary(merged_df$sum_SIRE_woc)
```

```
##    Min. 1st Qu.  Median    Mean 3rd Qu.    Max.
##  0.0000  1.0000  1.0000  0.8214  1.0000  5.0000
```

```
describe(merged_df$sum_SIRE_woc)
```

```
##    vars    n mean   sd median trimmed mad min max range skew kurtosis   se
## X1    1 9972 0.82 0.57      1    0.83   0   0   5     5 1.07     5.53 0.01
```

```
table(merged_df$sum_SIRE_woc)
```

```
##
##    0    1    2    3    4    5
## 2388 7245   99  214   24    2
```

```
merged_df$frac_White_SIRE_woc <-NA
merged_df$frac_White_SIRE_woc = merged_df$White_woc / merged_df$sum_SIRE_woc
merged_df$frac_White_SIRE_woc[is.nan(merged_df$frac_White_SIRE_woc)]<-0
table(merged_df$frac_White_SIRE_woc)
```

```
##
##                0              0.2             0.25 0.333333333333333
##             4212                2               24               201
##                1
##             5533
```

```
merged_df$frac_Black_SIRE_woc <-NA
merged_df$frac_Black_SIRE_woc = merged_df$Black_woc / merged_df$sum_SIRE_woc
merged_df$frac_Black_SIRE_woc[is.nan(merged_df$frac_Black_SIRE_woc)]<-0
table(merged_df$frac_Black_SIRE_woc)
```

```
##
##                0              0.2             0.25 0.333333333333333
##             8337                2               21               126
##              0.5                1
##               52             1434
```

```
merged_df$frac_EastAsian_SIRE_woc <-NA
merged_df$frac_EastAsian_SIRE_woc = merged_df$EastAsian_woc / merged_df$sum_S
IRE_woc
merged_df$frac_EastAsian_SIRE_woc[is.nan(merged_df$frac_EastAsian_SIRE_woc)]<
-0
table(merged_df$frac_EastAsian_SIRE_woc)
```

```
##
##                0              0.2             0.25 0.333333333333333
##             9746                2               10                56
##              0.5                1
##               51              107
```

```
merged_df$frac_SouthAsian_SIRE_woc <-NA
merged_df$frac_SouthAsian_SIRE_woc = merged_df$SouthAsian_woc / merged_df$sum
_SIRE_woc
merged_df$frac_SouthAsian_SIRE_woc[is.nan(merged_df$frac_SouthAsian_SIRE_woc)
]<-0
table(merged_df$frac_SouthAsian_SIRE_woc)
```

```
##
##                0 0.333333333333333              0.5                 1
##             9916                4                 9                43
```

```
merged_df$frac_Native_American_SIRE_woc <-NA
merged_df$frac_Native_American_SIRE_woc = merged_df$Native_American_woc / mer
ged_df$sum_SIRE_woc
merged_df$frac_Native_American_SIRE_woc[is.nan(merged_df$frac_Native_American
_SIRE_woc)]<-0
table(merged_df$frac_Native_American_SIRE_woc)
```

```
##
##                0              0.2             0.25 0.333333333333333
##             9758                2               22               115
```

```
##                0.5                1
##                 44               31
```

```
merged_df$frac_Other_Race_SIRE_woc <-NA
merged_df$frac_Other_Race_SIRE_woc = merged_df$Other_Race_woc / merged_df$sum
_SIRE_woc
merged_df$frac_Other_Race_SIRE_woc[is.nan(merged_df$frac_Other_Race_SIRE_woc)
]<-0
table(merged_df$frac_Other_Race_SIRE_woc)
```

```
##
##    0    1
## 9875   97
```

```
merged_df$frac_Hispanic_SIRE_woc <-NA
merged_df$frac_Hispanic_SIRE_woc = merged_df$Hispanic_woc / merged_df$sum_SIR
E_woc
merged_df$frac_Hispanic_SIRE_woc[is.nan(merged_df$frac_Hispanic_SIRE_woc)]<-0
table(merged_df$frac_Hispanic_SIRE_woc)
```

```
##
##                0               0.2              0.25 0.333333333333333
##             9769                 2                19               140
##              0.5
##               42
```

```
#See if add up to 1.00, except for xxxx combinations


table(merged_df$common_combination) #2388 combo
```

```
##
##    0    1
## 7584 2388
```

```
merged_df$frac_SIRE_SIRE_woc <- NA
merged_df$frac_SIRE_SIRE_woc = merged_df$frac_Hispanic_SIRE_woc + merged_df$f
rac_Other_Race_SIRE_woc + merged_df$frac_Native_American_SIRE_woc + merged_df
$frac_SouthAsian_SIRE_woc + merged_df$frac_EastAsian_SIRE_woc + merged_df$fra
c_Black_SIRE_woc + merged_df$frac_White_SIRE_woc

table(merged_df$frac_SIRE_SIRE_woc) #2388 combo
```

```
##
##    0    1
## 2388 7584
```

```
#Create SES variables (personal subjective ses, personal objective, neighborh
ood ses)
#Subjective SES
#
```

```
#meim_p_ss_total Ethnic  identity


merged_df$ses_1 <- NA
merged_df= mutate (merged_df,
                   ses_1= case_when(
                     demo_fam_exp1_v2 == "0" ~ 0,
                     demo_fam_exp1_v2 == "1" ~ -1,
                     TRUE ~ as.numeric(ses_1)) # This is for all other values
)

merged_df$ses_2 <- NA
merged_df= mutate (merged_df,
                   ses_2= case_when(
                     demo_fam_exp2_v2 == "0" ~ 0,
                     demo_fam_exp2_v2 == "1" ~ -1,
                     TRUE ~ as.numeric(ses_2)) # This is for all other values
)

merged_df$ses_3 <- NA
merged_df= mutate (merged_df,
                   ses_3= case_when(
                     demo_fam_exp3_v2 == "0" ~ 0,
                     demo_fam_exp3_v2 == "1" ~ -1,
                     TRUE ~ as.numeric(ses_3)) # This is for all other values
)

merged_df$ses_4 <- NA
merged_df= mutate (merged_df,
                   ses_4= case_when(
                     demo_fam_exp4_v2 == "0" ~ 0,
                     demo_fam_exp4_v2 == "1" ~ -1,
                     TRUE ~ as.numeric(ses_4)) # This is for all other values
)


merged_df$ses_5 <- NA
merged_df= mutate (merged_df,
                   ses_5= case_when(
                     demo_fam_exp5_v2 == "0" ~ 0,
                     demo_fam_exp5_v2 == "1" ~ -1,
                     TRUE ~ as.numeric(ses_5)) # This is for all other values
)



merged_df$ses_6 <- NA
merged_df= mutate (merged_df,
```

```
                      ses_6= case_when(
                        demo_fam_exp6_v2 == "0" ~ 0,
                        demo_fam_exp6_v2 == "1" ~ -1,
                        TRUE ~ as.numeric(ses_6)) # This is for all other values
)


merged_df$ses_7 <- NA
merged_df= mutate (merged_df,
                      ses_7= case_when(
                        demo_fam_exp7_v2 == "0" ~ 0,
                        demo_fam_exp7_v2 == "1" ~ -1,
                        TRUE ~ as.numeric(ses_7)) # This is for all other values
)


n_ses=which(names(merged_df)%in%c("ses_1"))

#compute sum

merged_df$sub_SES_sum= merged_df$ses_1 + merged_df$ses_2 + merged_df$ses_3 +
merged_df$ses_4 + merged_df$ses_5 + merged_df$ses_6 + merged_df$ses_7
hist(merged_df$sub_SES_sum)
```

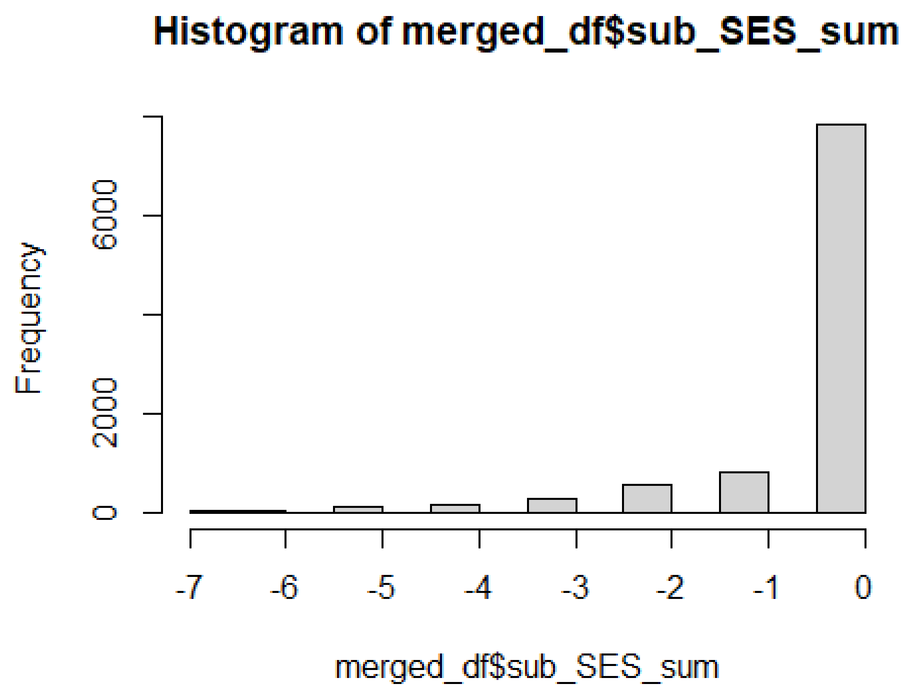


```
#standardize "sub_SES"
```

```
merged_df$sub_SES_Z= scale(merged_df$sub_SES_sum, center = TRUE, scale = TRUE
)
describe(merged_df$sub_SES_Z)

##    vars    n mean sd median trimmed mad   min  max range  skew kurtosis
se
## X1    1 9876    0  1   0.41    0.27   0 -6.04 0.41  6.45 -2.93     9.08 0.
01

hist(merged_df$sub_SES_Z)
```

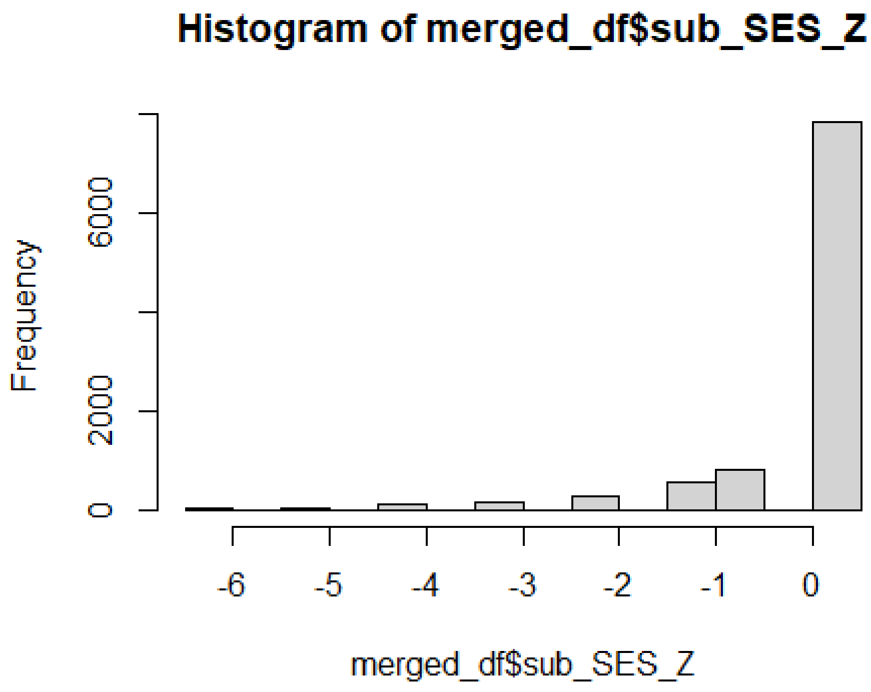


```
#Neighborhood SES simplified by just using address 1


table(merged_df$reshist_addr1_adi_perc)

##
##   1   2   3   4   5   6   7   8   9  10  11  12  13  14  15  16  17  18  1
9  20
## 544 189 172  73  98 121 134 125 178 134 130 128 115 126  89 175 124 177 17
4 171
##  21  22  23  24  25  26  27  28  29  30  31  32  33  34  35  36  37  38  3
9  40
## 183 128 217 142 179  91 191 188 141 137 207 107 127 116 125 142 145 167 12
6  88
##  41  42  43  44  45  46  47  48  49  50  51  52  53  54  55  56  57  58  5
9  60
```

```
## 113  82  94  80  96  71  63 108  76  82  56  73  83  76  50  64  84  54 10
2  52
##  61  62  63  64  65  66  67  68  69  70  71  72  73  74  75  76  77  78  7
9  80
##  67  89  68  40  44  77  56  50  28 108  33  48  51  52  31  42  49  38  4
6  28
##  81  82  83  84  85  86  87  88  89  90  91  92  93  94  95  96  97  98  9
9 100
## 117  48  29  51  60  34  50  49  54  31  47  85  61  57  26  51  50  42  6
5  67
## 101 102
##  66 104

describe(as.numeric(merged_df$reshist_addr1_adi_perc))

##    vars    n  mean    sd median trimmed   mad min max range skew kurtosis
se
## X1    1 9972 38.39 28.33     32   35.98 28.17   1 102   101 0.64    -0.61
0.28

merged_df$reshist_weighted_Z= scale(as.numeric(merged_df$reshist_addr1_adi_pe
rc, center = TRUE, scale = TRUE))*-1
describe(merged_df$reshist_weighted_Z)

##    vars    n mean sd median trimmed  mad   min  max range  skew kurtosis
se
## X1    1 9972    0  1   0.23    0.09 0.99 -2.25 1.32  3.56 -0.64    -0.61 0
.01

#Neighborhood Crime

#abcd_sscep01 <- read.delim("E:/ABCD/Hippo_3.0/Files 3.0/ABCDStudyNDA/abcd_ss
cep01.txt")
#abcd_sscep01_1_baseline=abcd_sscep01[abcd_sscep01$eventname=="baseline_year_
1_arm_1",]
#nsc_p_ss_mean_3_items <- c("nsc_p_ss_mean_3_items", "subjectkey")
#nsc_p_ss_mean_3_items_df <- abcd_sscep01_1_baseline[nsc_p_ss_mean_3_items]
#merged_df=merge(merged_df, nsc_p_ss_mean_3_items_df, by.x="subjectkey", by.y
= "subjectkey", all=T)

merged_df$nsc_p_ss_mean_3_items_z = scale(as.numeric(merged_df$nsc_p_ss_mean_
3_items), center = TRUE, scale = TRUE)
histogram(merged_df$nsc_p_ss_mean_3_items_z )
```

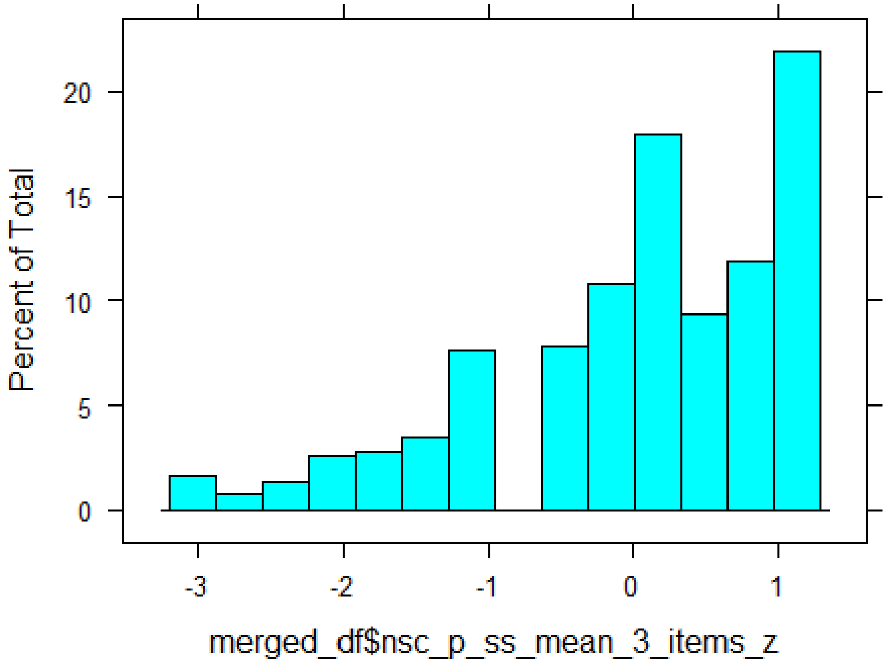


```
describe(merged_df$nsc_p_ss_mean_3_items_z)

##    vars    n mean sd median trimmed  mad   min  max range  skew kurtosis
se
## X1    1 9968    0  1   0.09    0.12 1.03 -3.04 1.13  4.16 -0.91     0.44 0
.01

#Personal SES

#Education

#Recode parental edu
#0=0, 1=1,â¦ 12=12, 13 = 12, 14 = 12, 15 = 14, 16 = 14, 17 = 14, 18 = 16, 19
=18, 20 = 18, 21=18



merged_df$edu_1 <- NA
merged_df= mutate (merged_df,
                   edu_1 = case_when(
                     demo_prnt_ed_v2_l %in% "0" ~ 0,
                     demo_prnt_ed_v2_l %in% "1" ~ 1,
                     demo_prnt_ed_v2_l %in% "2" ~ 2,
                     demo_prnt_ed_v2_l %in%  "3" ~ 3,
                     demo_prnt_ed_v2_l %in%  "4" ~ 4,
                     demo_prnt_ed_v2_l %in%  "5" ~ 5,
                     demo_prnt_ed_v2_l %in%  "6" ~ 6,
                     demo_prnt_ed_v2_l %in%  "7" ~ 7,
```

```
                      demo_prnt_ed_v2_l %in%  "8" ~ 8,
                      demo_prnt_ed_v2_l  %in%  "9" ~ 9,
                      demo_prnt_ed_v2_l %in%  "10" ~ 10,
                      demo_prnt_ed_v2_l %in% "11" ~ 11,
                      demo_prnt_ed_v2_l %in% "12" ~ 12,
                      demo_prnt_ed_v2_l %in%  "13" ~ 12,
                      demo_prnt_ed_v2_l %in%  "14" ~ 12,
                      demo_prnt_ed_v2_l %in%  "15" ~ 14,
                      demo_prnt_ed_v2_l %in%  "16" ~ 14,
                      demo_prnt_ed_v2_l %in%  "17" ~ 14,
                      demo_prnt_ed_v2_l %in%  "18" ~ 16,
                      demo_prnt_ed_v2_l  %in%  "19" ~ 18,
                      demo_prnt_ed_v2_l %in%  "20" ~ 18,
                      demo_prnt_ed_v2_l %in%  "21" ~ 18,
                      TRUE ~ as.numeric(edu_1)) # This is for all other values
)



merged_df$edu_2 <- NA
merged_df= mutate (merged_df,
                    edu_2 = case_when(
                      demo_prtnr_ed_v2_l %in% "0" ~ 0,
                      demo_prtnr_ed_v2_l %in% "1" ~ 1,
                      demo_prtnr_ed_v2_l %in% "2" ~ 2,
                      demo_prtnr_ed_v2_l %in%  "3" ~ 3,
                      demo_prtnr_ed_v2_l %in%  "4" ~ 4,
                      demo_prtnr_ed_v2_l %in%  "5" ~ 5,
                      demo_prtnr_ed_v2_l %in%  "6" ~ 6,
                      demo_prtnr_ed_v2_l %in%  "7" ~ 7,
                      demo_prtnr_ed_v2_l %in%  "8" ~ 8,
                      demo_prtnr_ed_v2_l  %in%  "9" ~ 9,
                      demo_prtnr_ed_v2_l %in%  "10" ~ 10,
                      demo_prtnr_ed_v2_l %in% "11" ~ 11,
                      demo_prtnr_ed_v2_l %in% "12" ~ 12,
                      demo_prtnr_ed_v2_l %in%  "13" ~ 12,
                      demo_prtnr_ed_v2_l %in%  "14" ~ 12,
                      demo_prtnr_ed_v2_l %in%  "15" ~ 14,
                      demo_prtnr_ed_v2_l %in%  "16" ~ 14,
                      demo_prtnr_ed_v2_l %in%  "17" ~ 14,
                      demo_prtnr_ed_v2_l %in%  "18" ~ 16,
                      demo_prtnr_ed_v2_l  %in%  "19" ~ 18,
                      demo_prtnr_ed_v2_l %in%  "20" ~ 18,
                      demo_prtnr_ed_v2_l %in%  "21" ~ 18,
                      TRUE ~ as.numeric(edu_2)) # This is for all other values
)


describe(merged_df$edu_1)
```

```
##    vars    n  mean   sd median trimmed  mad min max range  skew kurtosis
se
## X1    1 9914 15.24 2.29     16    15.43 2.97   1  18    17 -0.78     1.22 0
.02
```

```
describe(merged_df$edu_2)
```

```
##    vars    n  mean   sd median trimmed  mad min max range  skew kurtosis
se
## X1    1 7859 15.04 2.53     16    15.24 2.97   0  18    18 -0.94     1.82 0
.03
```

```
merged_df$edu_1_2 <-merged_df$edu_1
merged_df$edu_2_2 <-merged_df$edu_2

merged_df$education_mean <- rowMeans(merged_df[,2492:2493], na.rm=TRUE)
merged_df$edu_average_z= scale(merged_df$education_mean, center = TRUE, scale
= TRUE)
describe(merged_df$edu_average_z)
```

```
##    vars    n mean sd median trimmed  mad   min  max range  skew kurtosis
se
## X1    1 9922    0  1  -0.02    0.07 0.67 -5.45 1.33  6.78 -0.83     1.22 0
.01
```

```
#Marital status




merged_df$marital_status <- NA
merged_df= mutate(merged_df, marital_status
                      = case_when(
                       demo_prnt_marital_v2_l %in% 1 ~ 1,
                       demo_prnt_marital_v2_l %in% 2 ~ 0,
                       demo_prnt_marital_v2_l %in%  3 ~ 0,
                       demo_prnt_marital_v2_l %in%  4 ~ 0,
                       demo_prnt_marital_v2_l %in%  5 ~ 0,
                       demo_prnt_marital_v2_l %in%  6 ~ 0,
                       demo_prnt_marital_v2_l %in%  7 ~ 0,
                       demo_prnt_marital_v2_l %in%  777 ~ 0,
                       TRUE ~ as.numeric(marital_status)) # This is for all o
ther values
)


describe(merged_df$marital_status)
```

```
##    vars    n mean   sd median trimmed mad min max range  skew kurtosis se
## X1    1 9939 0.68 0.47      1    0.73   0   0   1     1 -0.79    -1.38  0
```

```r
table(merged_df$marital_status)

##
##    0    1
## 3147 6792

#Employed


merged_df$employed_1 <- NA
merged_df= mutate(merged_df, employed_1  = case_when(
  demo_prnt_empl_v2_l %in% 1 ~ 1,
  demo_prnt_empl_v2_l %in% 2 ~ 0,
  demo_prnt_empl_v2_l %in% 3 ~ 0,
  demo_prnt_empl_v2_l %in% 4 ~ 0,
  demo_prnt_empl_v2_l %in% 5 ~ 0,
  demo_prnt_empl_v2_l %in% 6 ~ 0,
  demo_prnt_empl_v2_l %in% 7 ~ 0,
  demo_prnt_empl_v2_l %in% 8 ~ 0,
  demo_prnt_empl_v2_l %in% 9 ~ 0,
  demo_prnt_empl_v2_l %in% 10 ~ 0,
  demo_prnt_empl_v2_l %in% 11 ~ 0,
  demo_prnt_empl_v2_l %in% 777 ~ 0,
  TRUE ~ as.numeric(employed_1)) # This is for all other values
)

merged_df$employed_2 <- NA
merged_df= mutate(merged_df, employed_2  = case_when(
  demo_prtnr_empl_v2_l %in% 1 ~ 1,
  demo_prtnr_empl_v2_l %in% 2 ~ 0,
  demo_prtnr_empl_v2_l %in% 3 ~ 0,
  demo_prtnr_empl_v2_l %in% 4 ~ 0,
  demo_prtnr_empl_v2_l %in% 5 ~ 0,
  demo_prtnr_empl_v2_l %in% 6 ~ 0,
  demo_prtnr_empl_v2_l %in% 7 ~ 0,
  demo_prtnr_empl_v2_l %in% 8 ~ 0,
  demo_prtnr_empl_v2_l %in% 9 ~ 0,
  demo_prtnr_empl_v2_l %in% 10 ~ 0,
  demo_prtnr_empl_v2_l %in% 11 ~ 0,
  demo_prtnr_empl_v2_l %in% 777 ~ 0,
  TRUE ~ as.numeric(employed_2)) # This is for all other values
)


merged_df$employed_sum=rowSums(cbind(merged_df$employed_1, merged_df$employed
_2), na.rm = TRUE)
table(merged_df$employed_sum)
```

```
##
##    0    1    2
##  868 3941 5163
```

```
merged_df$employed <- NA
merged_df= mutate(merged_df, employed  = case_when(
  employed_sum %in% 0 ~ 0,
  employed_sum %in% 1 ~ 1,
  employed_sum %in% 2 ~ 1,
  TRUE ~ as.numeric(employed)) # This is for all other values
)

table(merged_df$employed)
```

```
##
##    0    1
##  868 9104
```

```
#parental income

merged_df$income <- NA
merged_df= mutate (merged_df,
                income = case_when(
                  demo_comb_income_v2 %in% "1" ~ 4500,
                  demo_comb_income_v2 %in% "2" ~ 5000,
                  demo_comb_income_v2 %in%  "3" ~ 12000,
                  demo_comb_income_v2 %in%  "4" ~ 16000,
                  demo_comb_income_v2 %in%  "5" ~ 25000,
                  demo_comb_income_v2 %in%  "6" ~ 35000,
                  demo_comb_income_v2 %in%  "7" ~ 50000,
                  demo_comb_income_v2 %in%  "8" ~ 75000,
                  demo_comb_income_v2  %in%  "9" ~ 100000,
                  demo_comb_income_v2 %in%  "10" ~ 200000,
                  TRUE ~ as.numeric(income)) # This is for all other values
)


merged_df$income_z= scale(merged_df$income, center = TRUE, scale = TRUE)
describe(merged_df$income_z)
```

```
##    vars    n mean sd median trimmed  mad   min  max range skew kurtosis
se
## X1    1 9204    0  1  -0.08    -0.11 0.68 -1.37 2.21   3.58  0.9     0.33 0.
01
```

```
#create "parent objective SES" factor: edu_average + income + marital_status
+ employed_tot

#descriptives
```

```
describe(merged_df$sub_SES_Z)

##    vars    n mean sd median trimmed mad   min  max range  skew kurtosis
se
## X1    1 9876    0  1   0.41    0.27   0 -6.04 0.41  6.45 -2.93     9.08 0.
01

describe(merged_df$reshist_weighted_Z)

##    vars    n mean sd median trimmed  mad   min  max range  skew kurtosis
se
## X1    1 9972    0  1   0.23    0.09 0.99 -2.25 1.32  3.56 -0.64    -0.61 0
.01

describe(merged_df$nsc_p_ss_mean_3_items_z)

##    vars    n mean sd median trimmed  mad   min  max range  skew kurtosis
se
## X1    1 9968    0  1   0.09    0.12 1.03 -3.04 1.13  4.16 -0.91     0.44 0
.01

describe(merged_df$edu_average_z)

##    vars    n mean sd median trimmed  mad   min  max range  skew kurtosis
se
## X1    1 9922    0  1  -0.02    0.07 0.67 -5.45 1.33  6.78 -0.83     1.22 0
.01

describe(merged_df$income_z)

##    vars    n mean sd median trimmed  mad   min  max range skew kurtosis
se
## X1    1 9204    0  1  -0.08   -0.11 0.68 -1.37 2.21  3.58  0.9     0.33 0.
01

describe(merged_df$marital_status)

##    vars    n mean   sd median trimmed mad min max range  skew kurtosis se
## X1    1 9939 0.68 0.47      1    0.73   0   0   1     1 -0.79    -1.38  0

describe(merged_df$employed)

##    vars    n mean   sd median trimmed mad min max range  skew kurtosis se
## X1    1 9972 0.91 0.28      1       1   0   0   1     1 -2.93     6.58  0

describe(merged_df$CA_Z_adj)

##    vars    n mean sd median trimmed  mad  min  max range skew kurtosis   s
e
## X1    1 9972    0  1  -0.04   -0.02 0.96 -3.9 5.59  9.49 0.29     0.34 0.0
1
```

```
#subset to cases with CA

merged_dfs_CA_Z<-subset(merged_df,!merged_df$CA_Z_adj=="NA")
merged_df <- merged_dfs_CA_Z

#Create general SES based on neighborhood, personal, and subjective alternati
ve

df_general_SES_alt=data.frame(merged_df$subjectkey, merged_df$sub_SES_Z, merg
ed_df$reshist_weighted_Z, merged_df$nsc_p_ss_mean_3_items_z, merged_df$edu_av
erage_z, merged_df$income_z, merged_df$marital_status, merged_df$employed)
df.imputed_general_SES_alt=mice(df_general_SES_alt, m=5, maxit = 50, method =
'pmm', seed = 500)

##
##  iter imp variable
##   1   1  merged_df.sub_SES_Z  merged_df.nsc_p_ss_mean_3_items_z  merged_df
.edu_average_z  merged_df.income_z  merged_df.marital_status
##   1   2  merged_df.sub_SES_Z  merged_df.nsc_p_ss_mean_3_items_z  merged_df
.edu_average_z  merged_df.income_z  merged_df.marital_status
##   1   3  merged_df.sub_SES_Z  merged_df.nsc_p_ss_mean_3_items_z  merged_df
.edu_average_z  merged_df.income_z  merged_df.marital_status
##   1   4  merged_df.sub_SES_Z  merged_df.nsc_p_ss_mean_3_items_z  merged_df
.edu_average_z  merged_df.income_z  merged_df.marital_status
##   1   5  merged_df.sub_SES_Z  merged_df.nsc_p_ss_mean_3_items_z  merged_df
.edu_average_z  merged_df.income_z  merged_df.marital_status
##   2   1  merged_df.sub_SES_Z  merged_df.nsc_p_ss_mean_3_items_z  merged_df
.edu_average_z  merged_df.income_z  merged_df.marital_status
##   2   2  merged_df.sub_SES_Z  merged_df.nsc_p_ss_mean_3_items_z  merged_df
.edu_average_z  merged_df.income_z  merged_df.marital_status
##   2   3  merged_df.sub_SES_Z  merged_df.nsc_p_ss_mean_3_items_z  merged_df
.edu_average_z  merged_df.income_z  merged_df.marital_status
##   2   4  merged_df.sub_SES_Z  merged_df.nsc_p_ss_mean_3_items_z  merged_df
.edu_average_z  merged_df.income_z  merged_df.marital_status
##   2   5  merged_df.sub_SES_Z  merged_df.nsc_p_ss_mean_3_items_z  merged_df
.edu_average_z  merged_df.income_z  merged_df.marital_status
##   3   1  merged_df.sub_SES_Z  merged_df.nsc_p_ss_mean_3_items_z  merged_df
.edu_average_z  merged_df.income_z  merged_df.marital_status
##   3   2  merged_df.sub_SES_Z  merged_df.nsc_p_ss_mean_3_items_z  merged_df
.edu_average_z  merged_df.income_z  merged_df.marital_status
##   3   3  merged_df.sub_SES_Z  merged_df.nsc_p_ss_mean_3_items_z  merged_df
.edu_average_z  merged_df.income_z  merged_df.marital_status
##   3   4  merged_df.sub_SES_Z  merged_df.nsc_p_ss_mean_3_items_z  merged_df
.edu_average_z  merged_df.income_z  merged_df.marital_status
##   3   5  merged_df.sub_SES_Z  merged_df.nsc_p_ss_mean_3_items_z  merged_df
.edu_average_z  merged_df.income_z  merged_df.marital_status
##   4   1  merged_df.sub_SES_Z  merged_df.nsc_p_ss_mean_3_items_z  merged_df
.edu_average_z  merged_df.income_z  merged_df.marital_status
##   4   2  merged_df.sub_SES_Z  merged_df.nsc_p_ss_mean_3_items_z  merged_df
.edu_average_z  merged_df.income_z  merged_df.marital_status
```

```
##   4   3  merged_df.sub_SES_Z  merged_df.nsc_p_ss_mean_3_items_z  merged_df
.edu_average_z  merged_df.income_z  merged_df.marital_status
##   4   4  merged_df.sub_SES_Z  merged_df.nsc_p_ss_mean_3_items_z  merged_df
.edu_average_z  merged_df.income_z  merged_df.marital_status
##   4   5  merged_df.sub_SES_Z  merged_df.nsc_p_ss_mean_3_items_z  merged_df
.edu_average_z  merged_df.income_z  merged_df.marital_status
##   5   1  merged_df.sub_SES_Z  merged_df.nsc_p_ss_mean_3_items_z  merged_df
.edu_average_z  merged_df.income_z  merged_df.marital_status
##   5   2  merged_df.sub_SES_Z  merged_df.nsc_p_ss_mean_3_items_z  merged_df
.edu_average_z  merged_df.income_z  merged_df.marital_status
##   5   3  merged_df.sub_SES_Z  merged_df.nsc_p_ss_mean_3_items_z  merged_df
.edu_average_z  merged_df.income_z  merged_df.marital_status
##   5   4  merged_df.sub_SES_Z  merged_df.nsc_p_ss_mean_3_items_z  merged_df
.edu_average_z  merged_df.income_z  merged_df.marital_status
##   5   5  merged_df.sub_SES_Z  merged_df.nsc_p_ss_mean_3_items_z  merged_df
.edu_average_z  merged_df.income_z  merged_df.marital_status
##   6   1  merged_df.sub_SES_Z  merged_df.nsc_p_ss_mean_3_items_z  merged_df
.edu_average_z  merged_df.income_z  merged_df.marital_status
##   6   2  merged_df.sub_SES_Z  merged_df.nsc_p_ss_mean_3_items_z  merged_df
.edu_average_z  merged_df.income_z  merged_df.marital_status
##   6   3  merged_df.sub_SES_Z  merged_df.nsc_p_ss_mean_3_items_z  merged_df
.edu_average_z  merged_df.income_z  merged_df.marital_status
##   6   4  merged_df.sub_SES_Z  merged_df.nsc_p_ss_mean_3_items_z  merged_df
.edu_average_z  merged_df.income_z  merged_df.marital_status
##   6   5  merged_df.sub_SES_Z  merged_df.nsc_p_ss_mean_3_items_z  merged_df
.edu_average_z  merged_df.income_z  merged_df.marital_status
##   7   1  merged_df.sub_SES_Z  merged_df.nsc_p_ss_mean_3_items_z  merged_df
.edu_average_z  merged_df.income_z  merged_df.marital_status
##   7   2  merged_df.sub_SES_Z  merged_df.nsc_p_ss_mean_3_items_z  merged_df
.edu_average_z  merged_df.income_z  merged_df.marital_status
##   7   3  merged_df.sub_SES_Z  merged_df.nsc_p_ss_mean_3_items_z  merged_df
.edu_average_z  merged_df.income_z  merged_df.marital_status
##   7   4  merged_df.sub_SES_Z  merged_df.nsc_p_ss_mean_3_items_z  merged_df
.edu_average_z  merged_df.income_z  merged_df.marital_status
##   7   5  merged_df.sub_SES_Z  merged_df.nsc_p_ss_mean_3_items_z  merged_df
.edu_average_z  merged_df.income_z  merged_df.marital_status
##   8   1  merged_df.sub_SES_Z  merged_df.nsc_p_ss_mean_3_items_z  merged_df
.edu_average_z  merged_df.income_z  merged_df.marital_status
##   8   2  merged_df.sub_SES_Z  merged_df.nsc_p_ss_mean_3_items_z  merged_df
.edu_average_z  merged_df.income_z  merged_df.marital_status
##   8   3  merged_df.sub_SES_Z  merged_df.nsc_p_ss_mean_3_items_z  merged_df
.edu_average_z  merged_df.income_z  merged_df.marital_status
##   8   4  merged_df.sub_SES_Z  merged_df.nsc_p_ss_mean_3_items_z  merged_df
.edu_average_z  merged_df.income_z  merged_df.marital_status
##   8   5  merged_df.sub_SES_Z  merged_df.nsc_p_ss_mean_3_items_z  merged_df
.edu_average_z  merged_df.income_z  merged_df.marital_status
##   9   1  merged_df.sub_SES_Z  merged_df.nsc_p_ss_mean_3_items_z  merged_df
.edu_average_z  merged_df.income_z  merged_df.marital_status
##   9   2  merged_df.sub_SES_Z  merged_df.nsc_p_ss_mean_3_items_z  merged_df
.edu_average_z  merged_df.income_z  merged_df.marital_status
```

```
##   9   3  merged_df.sub_SES_Z  merged_df.nsc_p_ss_mean_3_items_z  merged_df
.edu_average_z  merged_df.income_z  merged_df.marital_status
##   9   4  merged_df.sub_SES_Z  merged_df.nsc_p_ss_mean_3_items_z  merged_df
.edu_average_z  merged_df.income_z  merged_df.marital_status
##   9   5  merged_df.sub_SES_Z  merged_df.nsc_p_ss_mean_3_items_z  merged_df
.edu_average_z  merged_df.income_z  merged_df.marital_status
##   10   1  merged_df.sub_SES_Z  merged_df.nsc_p_ss_mean_3_items_z  merged_d
f.edu_average_z  merged_df.income_z  merged_df.marital_status
##   10   2  merged_df.sub_SES_Z  merged_df.nsc_p_ss_mean_3_items_z  merged_d
f.edu_average_z  merged_df.income_z  merged_df.marital_status
##   10   3  merged_df.sub_SES_Z  merged_df.nsc_p_ss_mean_3_items_z  merged_d
f.edu_average_z  merged_df.income_z  merged_df.marital_status
##   10   4  merged_df.sub_SES_Z  merged_df.nsc_p_ss_mean_3_items_z  merged_d
f.edu_average_z  merged_df.income_z  merged_df.marital_status
##   10   5  merged_df.sub_SES_Z  merged_df.nsc_p_ss_mean_3_items_z  merged_d
f.edu_average_z  merged_df.income_z  merged_df.marital_status
##   11   1  merged_df.sub_SES_Z  merged_df.nsc_p_ss_mean_3_items_z  merged_d
f.edu_average_z  merged_df.income_z  merged_df.marital_status
##   11   2  merged_df.sub_SES_Z  merged_df.nsc_p_ss_mean_3_items_z  merged_d
f.edu_average_z  merged_df.income_z  merged_df.marital_status
##   11   3  merged_df.sub_SES_Z  merged_df.nsc_p_ss_mean_3_items_z  merged_d
f.edu_average_z  merged_df.income_z  merged_df.marital_status
##   11   4  merged_df.sub_SES_Z  merged_df.nsc_p_ss_mean_3_items_z  merged_d
f.edu_average_z  merged_df.income_z  merged_df.marital_status
##   11   5  merged_df.sub_SES_Z  merged_df.nsc_p_ss_mean_3_items_z  merged_d
f.edu_average_z  merged_df.income_z  merged_df.marital_status
##   12   1  merged_df.sub_SES_Z  merged_df.nsc_p_ss_mean_3_items_z  merged_d
f.edu_average_z  merged_df.income_z  merged_df.marital_status
##   12   2  merged_df.sub_SES_Z  merged_df.nsc_p_ss_mean_3_items_z  merged_d
f.edu_average_z  merged_df.income_z  merged_df.marital_status
##   12   3  merged_df.sub_SES_Z  merged_df.nsc_p_ss_mean_3_items_z  merged_d
f.edu_average_z  merged_df.income_z  merged_df.marital_status
##   12   4  merged_df.sub_SES_Z  merged_df.nsc_p_ss_mean_3_items_z  merged_d
f.edu_average_z  merged_df.income_z  merged_df.marital_status
##   12   5  merged_df.sub_SES_Z  merged_df.nsc_p_ss_mean_3_items_z  merged_d
f.edu_average_z  merged_df.income_z  merged_df.marital_status
##   13   1  merged_df.sub_SES_Z  merged_df.nsc_p_ss_mean_3_items_z  merged_d
f.edu_average_z  merged_df.income_z  merged_df.marital_status
##   13   2  merged_df.sub_SES_Z  merged_df.nsc_p_ss_mean_3_items_z  merged_d
f.edu_average_z  merged_df.income_z  merged_df.marital_status
##   13   3  merged_df.sub_SES_Z  merged_df.nsc_p_ss_mean_3_items_z  merged_d
f.edu_average_z  merged_df.income_z  merged_df.marital_status
##   13   4  merged_df.sub_SES_Z  merged_df.nsc_p_ss_mean_3_items_z  merged_d
f.edu_average_z  merged_df.income_z  merged_df.marital_status
##   13   5  merged_df.sub_SES_Z  merged_df.nsc_p_ss_mean_3_items_z  merged_d
f.edu_average_z  merged_df.income_z  merged_df.marital_status
##   14   1  merged_df.sub_SES_Z  merged_df.nsc_p_ss_mean_3_items_z  merged_d
f.edu_average_z  merged_df.income_z  merged_df.marital_status
##   14   2  merged_df.sub_SES_Z  merged_df.nsc_p_ss_mean_3_items_z  merged_d
f.edu_average_z  merged_df.income_z  merged_df.marital_status
```

```
##   14   3  merged_df.sub_SES_Z  merged_df.nsc_p_ss_mean_3_items_z  merged_d
f.edu_average_z  merged_df.income_z  merged_df.marital_status
##   14   4  merged_df.sub_SES_Z  merged_df.nsc_p_ss_mean_3_items_z  merged_d
f.edu_average_z  merged_df.income_z  merged_df.marital_status
##   14   5  merged_df.sub_SES_Z  merged_df.nsc_p_ss_mean_3_items_z  merged_d
f.edu_average_z  merged_df.income_z  merged_df.marital_status
##   15   1  merged_df.sub_SES_Z  merged_df.nsc_p_ss_mean_3_items_z  merged_d
f.edu_average_z  merged_df.income_z  merged_df.marital_status
##   15   2  merged_df.sub_SES_Z  merged_df.nsc_p_ss_mean_3_items_z  merged_d
f.edu_average_z  merged_df.income_z  merged_df.marital_status
##   15   3  merged_df.sub_SES_Z  merged_df.nsc_p_ss_mean_3_items_z  merged_d
f.edu_average_z  merged_df.income_z  merged_df.marital_status
##   15   4  merged_df.sub_SES_Z  merged_df.nsc_p_ss_mean_3_items_z  merged_d
f.edu_average_z  merged_df.income_z  merged_df.marital_status
##   15   5  merged_df.sub_SES_Z  merged_df.nsc_p_ss_mean_3_items_z  merged_d
f.edu_average_z  merged_df.income_z  merged_df.marital_status
##   16   1  merged_df.sub_SES_Z  merged_df.nsc_p_ss_mean_3_items_z  merged_d
f.edu_average_z  merged_df.income_z  merged_df.marital_status
##   16   2  merged_df.sub_SES_Z  merged_df.nsc_p_ss_mean_3_items_z  merged_d
f.edu_average_z  merged_df.income_z  merged_df.marital_status
##   16   3  merged_df.sub_SES_Z  merged_df.nsc_p_ss_mean_3_items_z  merged_d
f.edu_average_z  merged_df.income_z  merged_df.marital_status
##   16   4  merged_df.sub_SES_Z  merged_df.nsc_p_ss_mean_3_items_z  merged_d
f.edu_average_z  merged_df.income_z  merged_df.marital_status
##   16   5  merged_df.sub_SES_Z  merged_df.nsc_p_ss_mean_3_items_z  merged_d
f.edu_average_z  merged_df.income_z  merged_df.marital_status
##   17   1  merged_df.sub_SES_Z  merged_df.nsc_p_ss_mean_3_items_z  merged_d
f.edu_average_z  merged_df.income_z  merged_df.marital_status
##   17   2  merged_df.sub_SES_Z  merged_df.nsc_p_ss_mean_3_items_z  merged_d
f.edu_average_z  merged_df.income_z  merged_df.marital_status
##   17   3  merged_df.sub_SES_Z  merged_df.nsc_p_ss_mean_3_items_z  merged_d
f.edu_average_z  merged_df.income_z  merged_df.marital_status
##   17   4  merged_df.sub_SES_Z  merged_df.nsc_p_ss_mean_3_items_z  merged_d
f.edu_average_z  merged_df.income_z  merged_df.marital_status
##   17   5  merged_df.sub_SES_Z  merged_df.nsc_p_ss_mean_3_items_z  merged_d
f.edu_average_z  merged_df.income_z  merged_df.marital_status
##   18   1  merged_df.sub_SES_Z  merged_df.nsc_p_ss_mean_3_items_z  merged_d
f.edu_average_z  merged_df.income_z  merged_df.marital_status
##   18   2  merged_df.sub_SES_Z  merged_df.nsc_p_ss_mean_3_items_z  merged_d
f.edu_average_z  merged_df.income_z  merged_df.marital_status
##   18   3  merged_df.sub_SES_Z  merged_df.nsc_p_ss_mean_3_items_z  merged_d
f.edu_average_z  merged_df.income_z  merged_df.marital_status
##   18   4  merged_df.sub_SES_Z  merged_df.nsc_p_ss_mean_3_items_z  merged_d
f.edu_average_z  merged_df.income_z  merged_df.marital_status
##   18   5  merged_df.sub_SES_Z  merged_df.nsc_p_ss_mean_3_items_z  merged_d
f.edu_average_z  merged_df.income_z  merged_df.marital_status
##   19   1  merged_df.sub_SES_Z  merged_df.nsc_p_ss_mean_3_items_z  merged_d
f.edu_average_z  merged_df.income_z  merged_df.marital_status
##   19   2  merged_df.sub_SES_Z  merged_df.nsc_p_ss_mean_3_items_z  merged_d
f.edu_average_z  merged_df.income_z  merged_df.marital_status
```

```
##   19   3  merged_df.sub_SES_Z  merged_df.nsc_p_ss_mean_3_items_z  merged_d
f.edu_average_z  merged_df.income_z  merged_df.marital_status
##   19   4  merged_df.sub_SES_Z  merged_df.nsc_p_ss_mean_3_items_z  merged_d
f.edu_average_z  merged_df.income_z  merged_df.marital_status
##   19   5  merged_df.sub_SES_Z  merged_df.nsc_p_ss_mean_3_items_z  merged_d
f.edu_average_z  merged_df.income_z  merged_df.marital_status
##   20   1  merged_df.sub_SES_Z  merged_df.nsc_p_ss_mean_3_items_z  merged_d
f.edu_average_z  merged_df.income_z  merged_df.marital_status
##   20   2  merged_df.sub_SES_Z  merged_df.nsc_p_ss_mean_3_items_z  merged_d
f.edu_average_z  merged_df.income_z  merged_df.marital_status
##   20   3  merged_df.sub_SES_Z  merged_df.nsc_p_ss_mean_3_items_z  merged_d
f.edu_average_z  merged_df.income_z  merged_df.marital_status
##   20   4  merged_df.sub_SES_Z  merged_df.nsc_p_ss_mean_3_items_z  merged_d
f.edu_average_z  merged_df.income_z  merged_df.marital_status
##   20   5  merged_df.sub_SES_Z  merged_df.nsc_p_ss_mean_3_items_z  merged_d
f.edu_average_z  merged_df.income_z  merged_df.marital_status
##   21   1  merged_df.sub_SES_Z  merged_df.nsc_p_ss_mean_3_items_z  merged_d
f.edu_average_z  merged_df.income_z  merged_df.marital_status
##   21   2  merged_df.sub_SES_Z  merged_df.nsc_p_ss_mean_3_items_z  merged_d
f.edu_average_z  merged_df.income_z  merged_df.marital_status
##   21   3  merged_df.sub_SES_Z  merged_df.nsc_p_ss_mean_3_items_z  merged_d
f.edu_average_z  merged_df.income_z  merged_df.marital_status
##   21   4  merged_df.sub_SES_Z  merged_df.nsc_p_ss_mean_3_items_z  merged_d
f.edu_average_z  merged_df.income_z  merged_df.marital_status
##   21   5  merged_df.sub_SES_Z  merged_df.nsc_p_ss_mean_3_items_z  merged_d
f.edu_average_z  merged_df.income_z  merged_df.marital_status
##   22   1  merged_df.sub_SES_Z  merged_df.nsc_p_ss_mean_3_items_z  merged_d
f.edu_average_z  merged_df.income_z  merged_df.marital_status
##   22   2  merged_df.sub_SES_Z  merged_df.nsc_p_ss_mean_3_items_z  merged_d
f.edu_average_z  merged_df.income_z  merged_df.marital_status
##   22   3  merged_df.sub_SES_Z  merged_df.nsc_p_ss_mean_3_items_z  merged_d
f.edu_average_z  merged_df.income_z  merged_df.marital_status
##   22   4  merged_df.sub_SES_Z  merged_df.nsc_p_ss_mean_3_items_z  merged_d
f.edu_average_z  merged_df.income_z  merged_df.marital_status
##   22   5  merged_df.sub_SES_Z  merged_df.nsc_p_ss_mean_3_items_z  merged_d
f.edu_average_z  merged_df.income_z  merged_df.marital_status
##   23   1  merged_df.sub_SES_Z  merged_df.nsc_p_ss_mean_3_items_z  merged_d
f.edu_average_z  merged_df.income_z  merged_df.marital_status
##   23   2  merged_df.sub_SES_Z  merged_df.nsc_p_ss_mean_3_items_z  merged_d
f.edu_average_z  merged_df.income_z  merged_df.marital_status
##   23   3  merged_df.sub_SES_Z  merged_df.nsc_p_ss_mean_3_items_z  merged_d
f.edu_average_z  merged_df.income_z  merged_df.marital_status
##   23   4  merged_df.sub_SES_Z  merged_df.nsc_p_ss_mean_3_items_z  merged_d
f.edu_average_z  merged_df.income_z  merged_df.marital_status
##   23   5  merged_df.sub_SES_Z  merged_df.nsc_p_ss_mean_3_items_z  merged_d
f.edu_average_z  merged_df.income_z  merged_df.marital_status
##   24   1  merged_df.sub_SES_Z  merged_df.nsc_p_ss_mean_3_items_z  merged_d
f.edu_average_z  merged_df.income_z  merged_df.marital_status
##   24   2  merged_df.sub_SES_Z  merged_df.nsc_p_ss_mean_3_items_z  merged_d
f.edu_average_z  merged_df.income_z  merged_df.marital_status
```

```
##   24   3  merged_df.sub_SES_Z  merged_df.nsc_p_ss_mean_3_items_z  merged_d
f.edu_average_z  merged_df.income_z  merged_df.marital_status
##   24   4  merged_df.sub_SES_Z  merged_df.nsc_p_ss_mean_3_items_z  merged_d
f.edu_average_z  merged_df.income_z  merged_df.marital_status
##   24   5  merged_df.sub_SES_Z  merged_df.nsc_p_ss_mean_3_items_z  merged_d
f.edu_average_z  merged_df.income_z  merged_df.marital_status
##   25   1  merged_df.sub_SES_Z  merged_df.nsc_p_ss_mean_3_items_z  merged_d
f.edu_average_z  merged_df.income_z  merged_df.marital_status
##   25   2  merged_df.sub_SES_Z  merged_df.nsc_p_ss_mean_3_items_z  merged_d
f.edu_average_z  merged_df.income_z  merged_df.marital_status
##   25   3  merged_df.sub_SES_Z  merged_df.nsc_p_ss_mean_3_items_z  merged_d
f.edu_average_z  merged_df.income_z  merged_df.marital_status
##   25   4  merged_df.sub_SES_Z  merged_df.nsc_p_ss_mean_3_items_z  merged_d
f.edu_average_z  merged_df.income_z  merged_df.marital_status
##   25   5  merged_df.sub_SES_Z  merged_df.nsc_p_ss_mean_3_items_z  merged_d
f.edu_average_z  merged_df.income_z  merged_df.marital_status
##   26   1  merged_df.sub_SES_Z  merged_df.nsc_p_ss_mean_3_items_z  merged_d
f.edu_average_z  merged_df.income_z  merged_df.marital_status
##   26   2  merged_df.sub_SES_Z  merged_df.nsc_p_ss_mean_3_items_z  merged_d
f.edu_average_z  merged_df.income_z  merged_df.marital_status
##   26   3  merged_df.sub_SES_Z  merged_df.nsc_p_ss_mean_3_items_z  merged_d
f.edu_average_z  merged_df.income_z  merged_df.marital_status
##   26   4  merged_df.sub_SES_Z  merged_df.nsc_p_ss_mean_3_items_z  merged_d
f.edu_average_z  merged_df.income_z  merged_df.marital_status
##   26   5  merged_df.sub_SES_Z  merged_df.nsc_p_ss_mean_3_items_z  merged_d
f.edu_average_z  merged_df.income_z  merged_df.marital_status
##   27   1  merged_df.sub_SES_Z  merged_df.nsc_p_ss_mean_3_items_z  merged_d
f.edu_average_z  merged_df.income_z  merged_df.marital_status
##   27   2  merged_df.sub_SES_Z  merged_df.nsc_p_ss_mean_3_items_z  merged_d
f.edu_average_z  merged_df.income_z  merged_df.marital_status
##   27   3  merged_df.sub_SES_Z  merged_df.nsc_p_ss_mean_3_items_z  merged_d
f.edu_average_z  merged_df.income_z  merged_df.marital_status
##   27   4  merged_df.sub_SES_Z  merged_df.nsc_p_ss_mean_3_items_z  merged_d
f.edu_average_z  merged_df.income_z  merged_df.marital_status
##   27   5  merged_df.sub_SES_Z  merged_df.nsc_p_ss_mean_3_items_z  merged_d
f.edu_average_z  merged_df.income_z  merged_df.marital_status
##   28   1  merged_df.sub_SES_Z  merged_df.nsc_p_ss_mean_3_items_z  merged_d
f.edu_average_z  merged_df.income_z  merged_df.marital_status
##   28   2  merged_df.sub_SES_Z  merged_df.nsc_p_ss_mean_3_items_z  merged_d
f.edu_average_z  merged_df.income_z  merged_df.marital_status
##   28   3  merged_df.sub_SES_Z  merged_df.nsc_p_ss_mean_3_items_z  merged_d
f.edu_average_z  merged_df.income_z  merged_df.marital_status
##   28   4  merged_df.sub_SES_Z  merged_df.nsc_p_ss_mean_3_items_z  merged_d
f.edu_average_z  merged_df.income_z  merged_df.marital_status
##   28   5  merged_df.sub_SES_Z  merged_df.nsc_p_ss_mean_3_items_z  merged_d
f.edu_average_z  merged_df.income_z  merged_df.marital_status
##   29   1  merged_df.sub_SES_Z  merged_df.nsc_p_ss_mean_3_items_z  merged_d
f.edu_average_z  merged_df.income_z  merged_df.marital_status
##   29   2  merged_df.sub_SES_Z  merged_df.nsc_p_ss_mean_3_items_z  merged_d
f.edu_average_z  merged_df.income_z  merged_df.marital_status
```

```
##   29   3  merged_df.sub_SES_Z  merged_df.nsc_p_ss_mean_3_items_z  merged_d
f.edu_average_z  merged_df.income_z  merged_df.marital_status
##   29   4  merged_df.sub_SES_Z  merged_df.nsc_p_ss_mean_3_items_z  merged_d
f.edu_average_z  merged_df.income_z  merged_df.marital_status
##   29   5  merged_df.sub_SES_Z  merged_df.nsc_p_ss_mean_3_items_z  merged_d
f.edu_average_z  merged_df.income_z  merged_df.marital_status
##   30   1  merged_df.sub_SES_Z  merged_df.nsc_p_ss_mean_3_items_z  merged_d
f.edu_average_z  merged_df.income_z  merged_df.marital_status
##   30   2  merged_df.sub_SES_Z  merged_df.nsc_p_ss_mean_3_items_z  merged_d
f.edu_average_z  merged_df.income_z  merged_df.marital_status
##   30   3  merged_df.sub_SES_Z  merged_df.nsc_p_ss_mean_3_items_z  merged_d
f.edu_average_z  merged_df.income_z  merged_df.marital_status
##   30   4  merged_df.sub_SES_Z  merged_df.nsc_p_ss_mean_3_items_z  merged_d
f.edu_average_z  merged_df.income_z  merged_df.marital_status
##   30   5  merged_df.sub_SES_Z  merged_df.nsc_p_ss_mean_3_items_z  merged_d
f.edu_average_z  merged_df.income_z  merged_df.marital_status
##   31   1  merged_df.sub_SES_Z  merged_df.nsc_p_ss_mean_3_items_z  merged_d
f.edu_average_z  merged_df.income_z  merged_df.marital_status
##   31   2  merged_df.sub_SES_Z  merged_df.nsc_p_ss_mean_3_items_z  merged_d
f.edu_average_z  merged_df.income_z  merged_df.marital_status
##   31   3  merged_df.sub_SES_Z  merged_df.nsc_p_ss_mean_3_items_z  merged_d
f.edu_average_z  merged_df.income_z  merged_df.marital_status
##   31   4  merged_df.sub_SES_Z  merged_df.nsc_p_ss_mean_3_items_z  merged_d
f.edu_average_z  merged_df.income_z  merged_df.marital_status
##   31   5  merged_df.sub_SES_Z  merged_df.nsc_p_ss_mean_3_items_z  merged_d
f.edu_average_z  merged_df.income_z  merged_df.marital_status
##   32   1  merged_df.sub_SES_Z  merged_df.nsc_p_ss_mean_3_items_z  merged_d
f.edu_average_z  merged_df.income_z  merged_df.marital_status
##   32   2  merged_df.sub_SES_Z  merged_df.nsc_p_ss_mean_3_items_z  merged_d
f.edu_average_z  merged_df.income_z  merged_df.marital_status
##   32   3  merged_df.sub_SES_Z  merged_df.nsc_p_ss_mean_3_items_z  merged_d
f.edu_average_z  merged_df.income_z  merged_df.marital_status
##   32   4  merged_df.sub_SES_Z  merged_df.nsc_p_ss_mean_3_items_z  merged_d
f.edu_average_z  merged_df.income_z  merged_df.marital_status
##   32   5  merged_df.sub_SES_Z  merged_df.nsc_p_ss_mean_3_items_z  merged_d
f.edu_average_z  merged_df.income_z  merged_df.marital_status
##   33   1  merged_df.sub_SES_Z  merged_df.nsc_p_ss_mean_3_items_z  merged_d
f.edu_average_z  merged_df.income_z  merged_df.marital_status
##   33   2  merged_df.sub_SES_Z  merged_df.nsc_p_ss_mean_3_items_z  merged_d
f.edu_average_z  merged_df.income_z  merged_df.marital_status
##   33   3  merged_df.sub_SES_Z  merged_df.nsc_p_ss_mean_3_items_z  merged_d
f.edu_average_z  merged_df.income_z  merged_df.marital_status
##   33   4  merged_df.sub_SES_Z  merged_df.nsc_p_ss_mean_3_items_z  merged_d
f.edu_average_z  merged_df.income_z  merged_df.marital_status
##   33   5  merged_df.sub_SES_Z  merged_df.nsc_p_ss_mean_3_items_z  merged_d
f.edu_average_z  merged_df.income_z  merged_df.marital_status
##   34   1  merged_df.sub_SES_Z  merged_df.nsc_p_ss_mean_3_items_z  merged_d
f.edu_average_z  merged_df.income_z  merged_df.marital_status
##   34   2  merged_df.sub_SES_Z  merged_df.nsc_p_ss_mean_3_items_z  merged_d
f.edu_average_z  merged_df.income_z  merged_df.marital_status
```

```
##   34   3  merged_df.sub_SES_Z  merged_df.nsc_p_ss_mean_3_items_z  merged_d
f.edu_average_z  merged_df.income_z  merged_df.marital_status
##   34   4  merged_df.sub_SES_Z  merged_df.nsc_p_ss_mean_3_items_z  merged_d
f.edu_average_z  merged_df.income_z  merged_df.marital_status
##   34   5  merged_df.sub_SES_Z  merged_df.nsc_p_ss_mean_3_items_z  merged_d
f.edu_average_z  merged_df.income_z  merged_df.marital_status
##   35   1  merged_df.sub_SES_Z  merged_df.nsc_p_ss_mean_3_items_z  merged_d
f.edu_average_z  merged_df.income_z  merged_df.marital_status
##   35   2  merged_df.sub_SES_Z  merged_df.nsc_p_ss_mean_3_items_z  merged_d
f.edu_average_z  merged_df.income_z  merged_df.marital_status
##   35   3  merged_df.sub_SES_Z  merged_df.nsc_p_ss_mean_3_items_z  merged_d
f.edu_average_z  merged_df.income_z  merged_df.marital_status
##   35   4  merged_df.sub_SES_Z  merged_df.nsc_p_ss_mean_3_items_z  merged_d
f.edu_average_z  merged_df.income_z  merged_df.marital_status
##   35   5  merged_df.sub_SES_Z  merged_df.nsc_p_ss_mean_3_items_z  merged_d
f.edu_average_z  merged_df.income_z  merged_df.marital_status
##   36   1  merged_df.sub_SES_Z  merged_df.nsc_p_ss_mean_3_items_z  merged_d
f.edu_average_z  merged_df.income_z  merged_df.marital_status
##   36   2  merged_df.sub_SES_Z  merged_df.nsc_p_ss_mean_3_items_z  merged_d
f.edu_average_z  merged_df.income_z  merged_df.marital_status
##   36   3  merged_df.sub_SES_Z  merged_df.nsc_p_ss_mean_3_items_z  merged_d
f.edu_average_z  merged_df.income_z  merged_df.marital_status
##   36   4  merged_df.sub_SES_Z  merged_df.nsc_p_ss_mean_3_items_z  merged_d
f.edu_average_z  merged_df.income_z  merged_df.marital_status
##   36   5  merged_df.sub_SES_Z  merged_df.nsc_p_ss_mean_3_items_z  merged_d
f.edu_average_z  merged_df.income_z  merged_df.marital_status
##   37   1  merged_df.sub_SES_Z  merged_df.nsc_p_ss_mean_3_items_z  merged_d
f.edu_average_z  merged_df.income_z  merged_df.marital_status
##   37   2  merged_df.sub_SES_Z  merged_df.nsc_p_ss_mean_3_items_z  merged_d
f.edu_average_z  merged_df.income_z  merged_df.marital_status
##   37   3  merged_df.sub_SES_Z  merged_df.nsc_p_ss_mean_3_items_z  merged_d
f.edu_average_z  merged_df.income_z  merged_df.marital_status
##   37   4  merged_df.sub_SES_Z  merged_df.nsc_p_ss_mean_3_items_z  merged_d
f.edu_average_z  merged_df.income_z  merged_df.marital_status
##   37   5  merged_df.sub_SES_Z  merged_df.nsc_p_ss_mean_3_items_z  merged_d
f.edu_average_z  merged_df.income_z  merged_df.marital_status
##   38   1  merged_df.sub_SES_Z  merged_df.nsc_p_ss_mean_3_items_z  merged_d
f.edu_average_z  merged_df.income_z  merged_df.marital_status
##   38   2  merged_df.sub_SES_Z  merged_df.nsc_p_ss_mean_3_items_z  merged_d
f.edu_average_z  merged_df.income_z  merged_df.marital_status
##   38   3  merged_df.sub_SES_Z  merged_df.nsc_p_ss_mean_3_items_z  merged_d
f.edu_average_z  merged_df.income_z  merged_df.marital_status
##   38   4  merged_df.sub_SES_Z  merged_df.nsc_p_ss_mean_3_items_z  merged_d
f.edu_average_z  merged_df.income_z  merged_df.marital_status
##   38   5  merged_df.sub_SES_Z  merged_df.nsc_p_ss_mean_3_items_z  merged_d
f.edu_average_z  merged_df.income_z  merged_df.marital_status
##   39   1  merged_df.sub_SES_Z  merged_df.nsc_p_ss_mean_3_items_z  merged_d
f.edu_average_z  merged_df.income_z  merged_df.marital_status
##   39   2  merged_df.sub_SES_Z  merged_df.nsc_p_ss_mean_3_items_z  merged_d
f.edu_average_z  merged_df.income_z  merged_df.marital_status
```

```
##   39   3  merged_df.sub_SES_Z  merged_df.nsc_p_ss_mean_3_items_z  merged_d
f.edu_average_z  merged_df.income_z  merged_df.marital_status
##   39   4  merged_df.sub_SES_Z  merged_df.nsc_p_ss_mean_3_items_z  merged_d
f.edu_average_z  merged_df.income_z  merged_df.marital_status
##   39   5  merged_df.sub_SES_Z  merged_df.nsc_p_ss_mean_3_items_z  merged_d
f.edu_average_z  merged_df.income_z  merged_df.marital_status
##   40   1  merged_df.sub_SES_Z  merged_df.nsc_p_ss_mean_3_items_z  merged_d
f.edu_average_z  merged_df.income_z  merged_df.marital_status
##   40   2  merged_df.sub_SES_Z  merged_df.nsc_p_ss_mean_3_items_z  merged_d
f.edu_average_z  merged_df.income_z  merged_df.marital_status
##   40   3  merged_df.sub_SES_Z  merged_df.nsc_p_ss_mean_3_items_z  merged_d
f.edu_average_z  merged_df.income_z  merged_df.marital_status
##   40   4  merged_df.sub_SES_Z  merged_df.nsc_p_ss_mean_3_items_z  merged_d
f.edu_average_z  merged_df.income_z  merged_df.marital_status
##   40   5  merged_df.sub_SES_Z  merged_df.nsc_p_ss_mean_3_items_z  merged_d
f.edu_average_z  merged_df.income_z  merged_df.marital_status
##   41   1  merged_df.sub_SES_Z  merged_df.nsc_p_ss_mean_3_items_z  merged_d
f.edu_average_z  merged_df.income_z  merged_df.marital_status
##   41   2  merged_df.sub_SES_Z  merged_df.nsc_p_ss_mean_3_items_z  merged_d
f.edu_average_z  merged_df.income_z  merged_df.marital_status
##   41   3  merged_df.sub_SES_Z  merged_df.nsc_p_ss_mean_3_items_z  merged_d
f.edu_average_z  merged_df.income_z  merged_df.marital_status
##   41   4  merged_df.sub_SES_Z  merged_df.nsc_p_ss_mean_3_items_z  merged_d
f.edu_average_z  merged_df.income_z  merged_df.marital_status
##   41   5  merged_df.sub_SES_Z  merged_df.nsc_p_ss_mean_3_items_z  merged_d
f.edu_average_z  merged_df.income_z  merged_df.marital_status
##   42   1  merged_df.sub_SES_Z  merged_df.nsc_p_ss_mean_3_items_z  merged_d
f.edu_average_z  merged_df.income_z  merged_df.marital_status
##   42   2  merged_df.sub_SES_Z  merged_df.nsc_p_ss_mean_3_items_z  merged_d
f.edu_average_z  merged_df.income_z  merged_df.marital_status
##   42   3  merged_df.sub_SES_Z  merged_df.nsc_p_ss_mean_3_items_z  merged_d
f.edu_average_z  merged_df.income_z  merged_df.marital_status
##   42   4  merged_df.sub_SES_Z  merged_df.nsc_p_ss_mean_3_items_z  merged_d
f.edu_average_z  merged_df.income_z  merged_df.marital_status
##   42   5  merged_df.sub_SES_Z  merged_df.nsc_p_ss_mean_3_items_z  merged_d
f.edu_average_z  merged_df.income_z  merged_df.marital_status
##   43   1  merged_df.sub_SES_Z  merged_df.nsc_p_ss_mean_3_items_z  merged_d
f.edu_average_z  merged_df.income_z  merged_df.marital_status
##   43   2  merged_df.sub_SES_Z  merged_df.nsc_p_ss_mean_3_items_z  merged_d
f.edu_average_z  merged_df.income_z  merged_df.marital_status
##   43   3  merged_df.sub_SES_Z  merged_df.nsc_p_ss_mean_3_items_z  merged_d
f.edu_average_z  merged_df.income_z  merged_df.marital_status
##   43   4  merged_df.sub_SES_Z  merged_df.nsc_p_ss_mean_3_items_z  merged_d
f.edu_average_z  merged_df.income_z  merged_df.marital_status
##   43   5  merged_df.sub_SES_Z  merged_df.nsc_p_ss_mean_3_items_z  merged_d
f.edu_average_z  merged_df.income_z  merged_df.marital_status
##   44   1  merged_df.sub_SES_Z  merged_df.nsc_p_ss_mean_3_items_z  merged_d
f.edu_average_z  merged_df.income_z  merged_df.marital_status
##   44   2  merged_df.sub_SES_Z  merged_df.nsc_p_ss_mean_3_items_z  merged_d
f.edu_average_z  merged_df.income_z  merged_df.marital_status
```

```
##   44   3  merged_df.sub_SES_Z  merged_df.nsc_p_ss_mean_3_items_z  merged_d
f.edu_average_z  merged_df.income_z  merged_df.marital_status
##   44   4  merged_df.sub_SES_Z  merged_df.nsc_p_ss_mean_3_items_z  merged_d
f.edu_average_z  merged_df.income_z  merged_df.marital_status
##   44   5  merged_df.sub_SES_Z  merged_df.nsc_p_ss_mean_3_items_z  merged_d
f.edu_average_z  merged_df.income_z  merged_df.marital_status
##   45   1  merged_df.sub_SES_Z  merged_df.nsc_p_ss_mean_3_items_z  merged_d
f.edu_average_z  merged_df.income_z  merged_df.marital_status
##   45   2  merged_df.sub_SES_Z  merged_df.nsc_p_ss_mean_3_items_z  merged_d
f.edu_average_z  merged_df.income_z  merged_df.marital_status
##   45   3  merged_df.sub_SES_Z  merged_df.nsc_p_ss_mean_3_items_z  merged_d
f.edu_average_z  merged_df.income_z  merged_df.marital_status
##   45   4  merged_df.sub_SES_Z  merged_df.nsc_p_ss_mean_3_items_z  merged_d
f.edu_average_z  merged_df.income_z  merged_df.marital_status
##   45   5  merged_df.sub_SES_Z  merged_df.nsc_p_ss_mean_3_items_z  merged_d
f.edu_average_z  merged_df.income_z  merged_df.marital_status
##   46   1  merged_df.sub_SES_Z  merged_df.nsc_p_ss_mean_3_items_z  merged_d
f.edu_average_z  merged_df.income_z  merged_df.marital_status
##   46   2  merged_df.sub_SES_Z  merged_df.nsc_p_ss_mean_3_items_z  merged_d
f.edu_average_z  merged_df.income_z  merged_df.marital_status
##   46   3  merged_df.sub_SES_Z  merged_df.nsc_p_ss_mean_3_items_z  merged_d
f.edu_average_z  merged_df.income_z  merged_df.marital_status
##   46   4  merged_df.sub_SES_Z  merged_df.nsc_p_ss_mean_3_items_z  merged_d
f.edu_average_z  merged_df.income_z  merged_df.marital_status
##   46   5  merged_df.sub_SES_Z  merged_df.nsc_p_ss_mean_3_items_z  merged_d
f.edu_average_z  merged_df.income_z  merged_df.marital_status
##   47   1  merged_df.sub_SES_Z  merged_df.nsc_p_ss_mean_3_items_z  merged_d
f.edu_average_z  merged_df.income_z  merged_df.marital_status
##   47   2  merged_df.sub_SES_Z  merged_df.nsc_p_ss_mean_3_items_z  merged_d
f.edu_average_z  merged_df.income_z  merged_df.marital_status
##   47   3  merged_df.sub_SES_Z  merged_df.nsc_p_ss_mean_3_items_z  merged_d
f.edu_average_z  merged_df.income_z  merged_df.marital_status
##   47   4  merged_df.sub_SES_Z  merged_df.nsc_p_ss_mean_3_items_z  merged_d
f.edu_average_z  merged_df.income_z  merged_df.marital_status
##   47   5  merged_df.sub_SES_Z  merged_df.nsc_p_ss_mean_3_items_z  merged_d
f.edu_average_z  merged_df.income_z  merged_df.marital_status
##   48   1  merged_df.sub_SES_Z  merged_df.nsc_p_ss_mean_3_items_z  merged_d
f.edu_average_z  merged_df.income_z  merged_df.marital_status
##   48   2  merged_df.sub_SES_Z  merged_df.nsc_p_ss_mean_3_items_z  merged_d
f.edu_average_z  merged_df.income_z  merged_df.marital_status
##   48   3  merged_df.sub_SES_Z  merged_df.nsc_p_ss_mean_3_items_z  merged_d
f.edu_average_z  merged_df.income_z  merged_df.marital_status
##   48   4  merged_df.sub_SES_Z  merged_df.nsc_p_ss_mean_3_items_z  merged_d
f.edu_average_z  merged_df.income_z  merged_df.marital_status
##   48   5  merged_df.sub_SES_Z  merged_df.nsc_p_ss_mean_3_items_z  merged_d
f.edu_average_z  merged_df.income_z  merged_df.marital_status
##   49   1  merged_df.sub_SES_Z  merged_df.nsc_p_ss_mean_3_items_z  merged_d
f.edu_average_z  merged_df.income_z  merged_df.marital_status
##   49   2  merged_df.sub_SES_Z  merged_df.nsc_p_ss_mean_3_items_z  merged_d
f.edu_average_z  merged_df.income_z  merged_df.marital_status
```

```
##   49   3  merged_df.sub_SES_Z  merged_df.nsc_p_ss_mean_3_items_z  merged_d
f.edu_average_z  merged_df.income_z  merged_df.marital_status
##   49   4  merged_df.sub_SES_Z  merged_df.nsc_p_ss_mean_3_items_z  merged_d
f.edu_average_z  merged_df.income_z  merged_df.marital_status
##   49   5  merged_df.sub_SES_Z  merged_df.nsc_p_ss_mean_3_items_z  merged_d
f.edu_average_z  merged_df.income_z  merged_df.marital_status
##   50   1  merged_df.sub_SES_Z  merged_df.nsc_p_ss_mean_3_items_z  merged_d
f.edu_average_z  merged_df.income_z  merged_df.marital_status
##   50   2  merged_df.sub_SES_Z  merged_df.nsc_p_ss_mean_3_items_z  merged_d
f.edu_average_z  merged_df.income_z  merged_df.marital_status
##   50   3  merged_df.sub_SES_Z  merged_df.nsc_p_ss_mean_3_items_z  merged_d
f.edu_average_z  merged_df.income_z  merged_df.marital_status
##   50   4  merged_df.sub_SES_Z  merged_df.nsc_p_ss_mean_3_items_z  merged_d
f.edu_average_z  merged_df.income_z  merged_df.marital_status
##   50   5  merged_df.sub_SES_Z  merged_df.nsc_p_ss_mean_3_items_z  merged_d
f.edu_average_z  merged_df.income_z  merged_df.marital_status

## Warning: Number of logged events: 1

completeData_general_SES_alt <- complete(df.imputed_general_SES_alt,2)



#rescale

completeData_general_SES_alt$sub_SES_rescale= scale(completeData_general_SES_
alt$merged_df.sub_SES_Z, center = TRUE, scale = TRUE)
describe(completeData_general_SES_alt$sub_SES_rescale)

##    vars    n mean sd median trimmed mad min  max range  skew kurtosis   se
## X1    1 9972    0  1   0.42    0.27   0  -6 0.42  6.42 -2.91     8.96 0.01

hist(completeData_general_SES_alt$sub_SES_rescale)
```

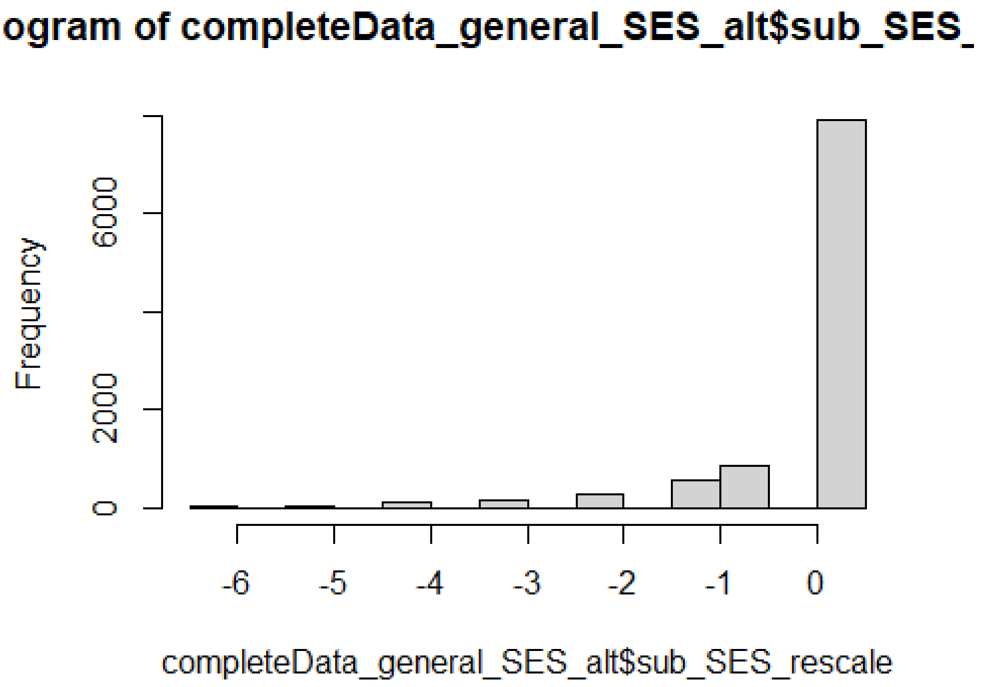


```
completeData_general_SES_alt$reshist_weighted_rescale= scale(completeData_gen
eral_SES_alt$merged_df.reshist_weighted_Z, center = TRUE, scale = TRUE)
describe(completeData_general_SES_alt$reshist_weighted_rescale)

##    vars    n mean sd median trimmed  mad   min  max range  skew kurtosis
se
## X1    1 9972    0  1   0.23    0.09 0.99 -2.25 1.32  3.56 -0.64    -0.61 0
.01

hist(completeData_general_SES_alt$reshist_weighted_rescale)
```

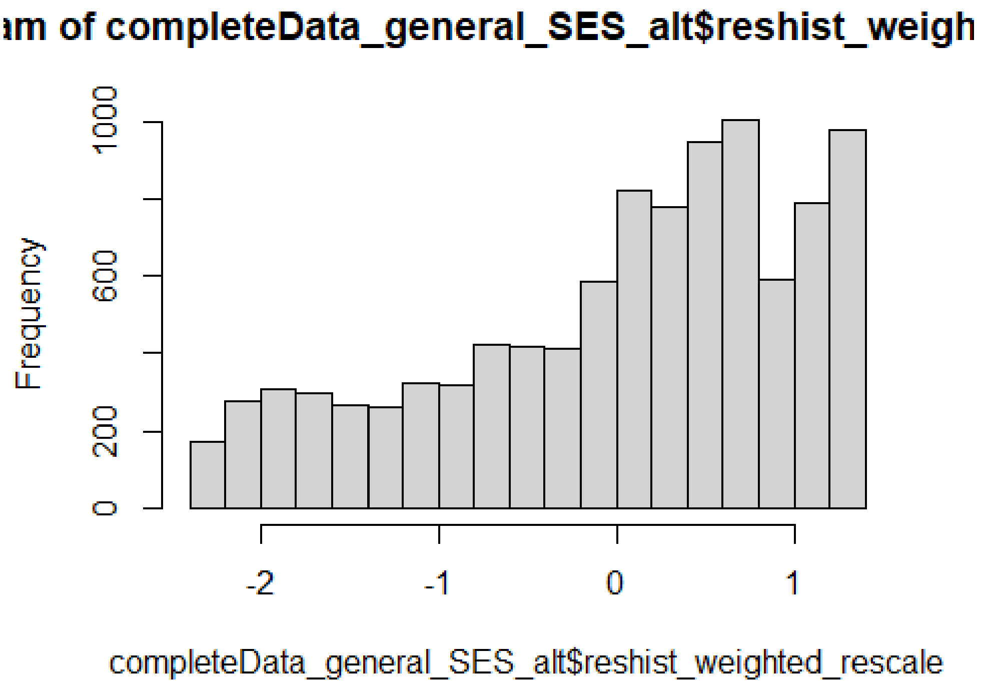


```
completeData_general_SES_alt$nsc_p_ss_mean_3_items_rescale= scale(completeDat
a_general_SES_alt$merged_df.nsc_p_ss_mean_3_items_z, center = TRUE, scale = T
RUE)
describe(completeData_general_SES_alt$nsc_p_ss_mean_3_items_rescale)

##    vars    n mean sd median trimmed  mad   min  max range  skew kurtosis
se
## X1    1 9972    0  1   0.09    0.12 1.03 -3.04 1.13  4.16 -0.91     0.44 0
.01

hist(completeData_general_SES_alt$nsc_p_ss_mean_3_items_rescale)
```

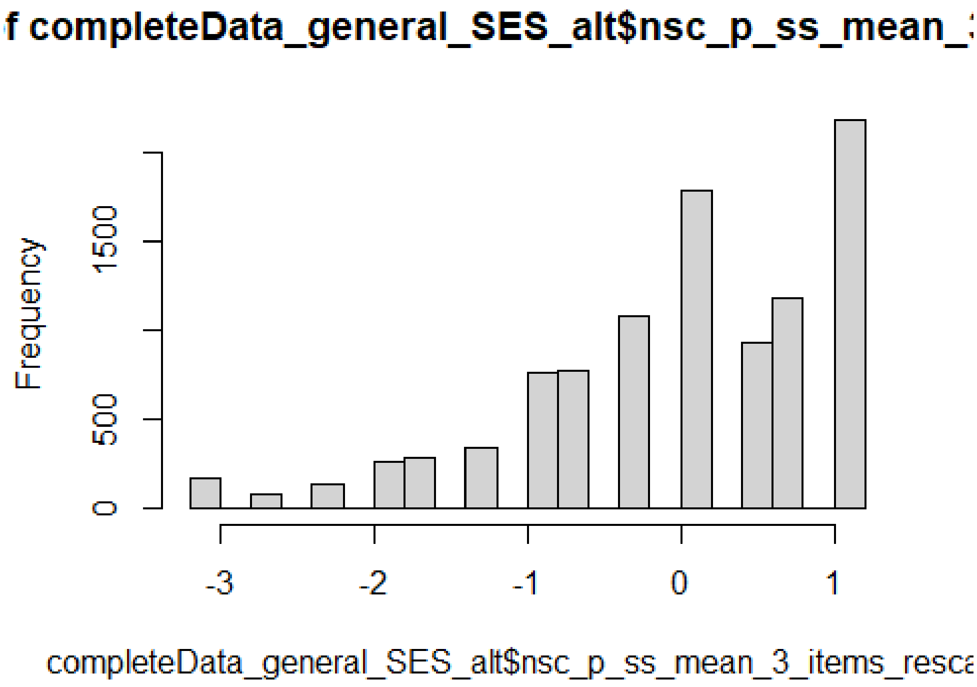


```
completeData_general_SES_alt$edu_average_rescale= scale(completeData_general_
SES_alt$merged_df.edu_average_z, center = TRUE, scale = TRUE)
describe(completeData_general_SES_alt$edu_average_rescale)

##    vars    n mean sd median trimmed  mad   min  max range  skew kurtosis
se
## X1    1 9972    0  1  -0.02    0.07 0.67 -5.45 1.33  6.78 -0.82     1.21 0
.01

hist(completeData_general_SES_alt$edu_average_rescale)
```

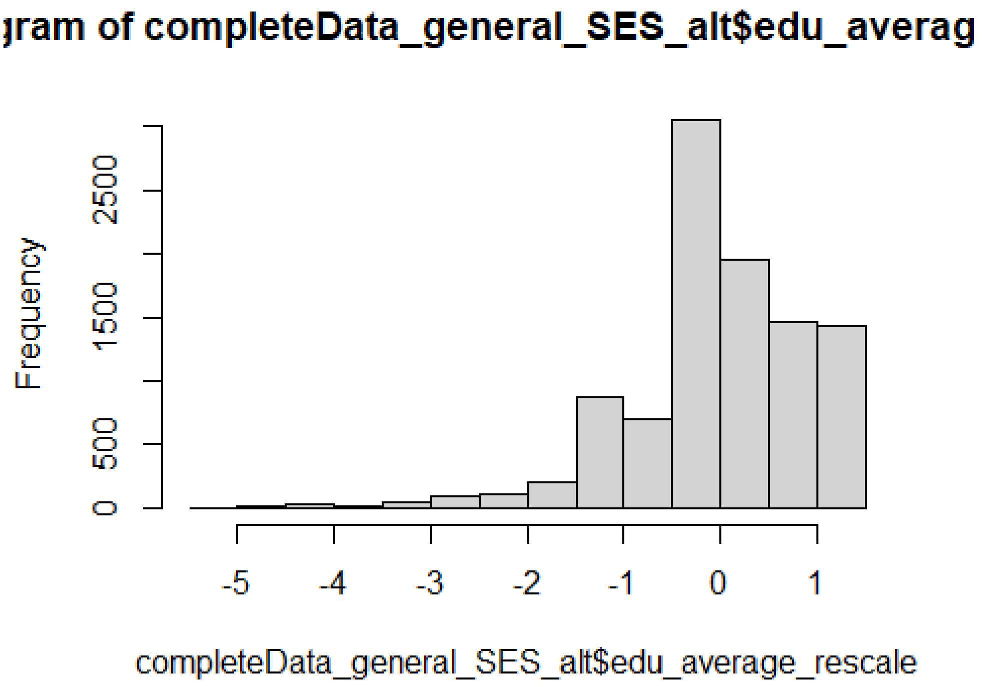


```
completeData_general_SES_alt$income_rescale= scale(completeData_general_SES_a
lt$merged_df.income_z, center = TRUE, scale = TRUE)
describe(completeData_general_SES_alt$income_rescale)

##    vars    n mean sd median trimmed  mad   min  max range skew kurtosis
se
## X1    1 9972    0  1  -0.04   -0.12 0.68 -1.34 2.25  3.58 0.92     0.37 0.
01

hist(completeData_general_SES_alt$income_rescale)
```

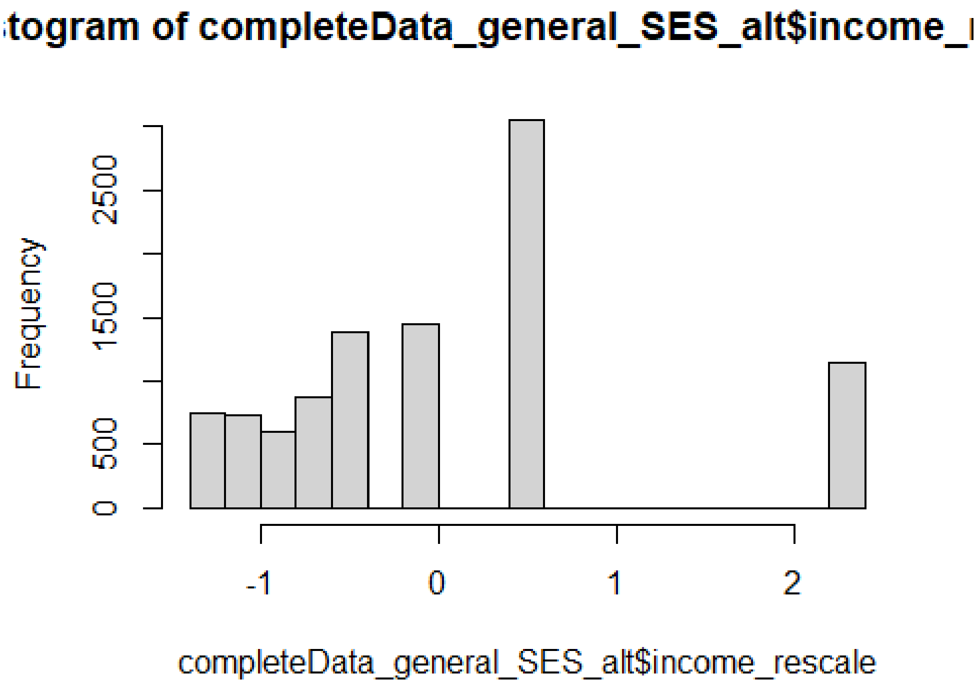


```
completeData_general_SES_alt_no_subject_key <- completeData_general_SES_alt[,
-c(1,2,3,4,5,6)]
apa.cor.table(completeData_general_SES_alt_no_subject_key, filename="APA_SES_
full_Connor.doc", table.number=1)

##
##
## Table 1
##
## Means, standard deviations, and correlations with confidence intervals
##
##
##   Variable                         M     SD   1          2          3
##   1. merged_df.marital_status      0.68  0.47
##
##   2. merged_df.employed            0.91  0.28 .28**
##                                               [.27, .30]
##
##   3. sub_SES_rescale               -0.00 1.00 .27**      .16**
##                                               [.25, .29] [.14, .18]
##
##   4. reshist_weighted_rescale      0.00  1.00 .23**      .13**      .21**
##                                               [.21, .25] [.11, .15] [.19,
.23]
##
##   5. nsc_p_ss_mean_3_items_rescale -0.00 1.00 .23**      .15**      .24**
##                                               [.21, .25] [.13, .17] [.22,
```

```
.26]
##
##   6. edu_average_rescale           -0.00 1.00 .33**       .26**       .28**
##                                               [.31, .35] [.25, .28] [.27,
.30]
##
##   7. income_rescale                -0.00 1.00 .46**       .27**       .34**
##                                               [.44, .47] [.25, .28] [.33,
.36]
##
##   4          5          6
##
##
##
##
##
##
##
##
##
##
##
##   .25**
##   [.23, .27]
##
##   .28**      .28**
##   [.26, .30] [.27, .30]
##
##   .34**      .31**      .57**
##   [.33, .36] [.29, .33] [.56, .58]
##
##
## Note. M and SD are used to represent mean and standard deviation, respecti
vely.
## Values in square brackets indicate the 95% confidence interval.
## The confidence interval is a plausible range of population correlations
## that could have caused the sample correlation (Cumming, 2014).
##  * indicates p < .05. ** indicates p < .01.
##

df_quanti=completeData_general_SES_alt[,c(9,10,11,12,13)]
df_quali=completeData_general_SES_alt[,c(7,8)]

df_quali[,c(1,2)]<-data.frame(apply(df_quali[,c(1,2)], 2, function(x){ as.fac
tor(x)}))#turn columns into factors

fit_PCAmix=PCAmix(X.quanti = df_quanti, X.quali = df_quali, ndim = 5, rename.
level = TRUE,
                  weight.col.quanti = NULL, weight.col.quali = NULL, graph =
TRUE)
```

**Individuals component map**

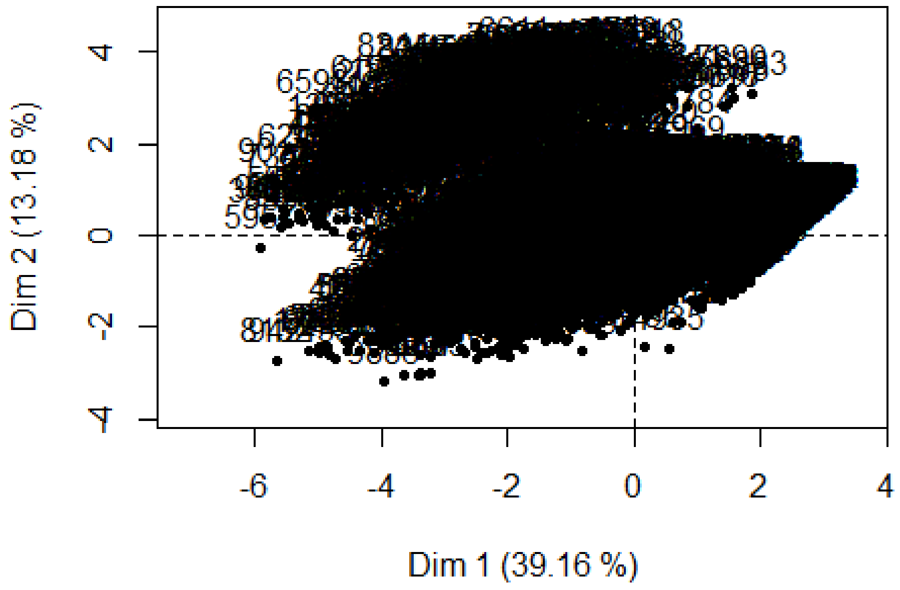


**Levels component map**

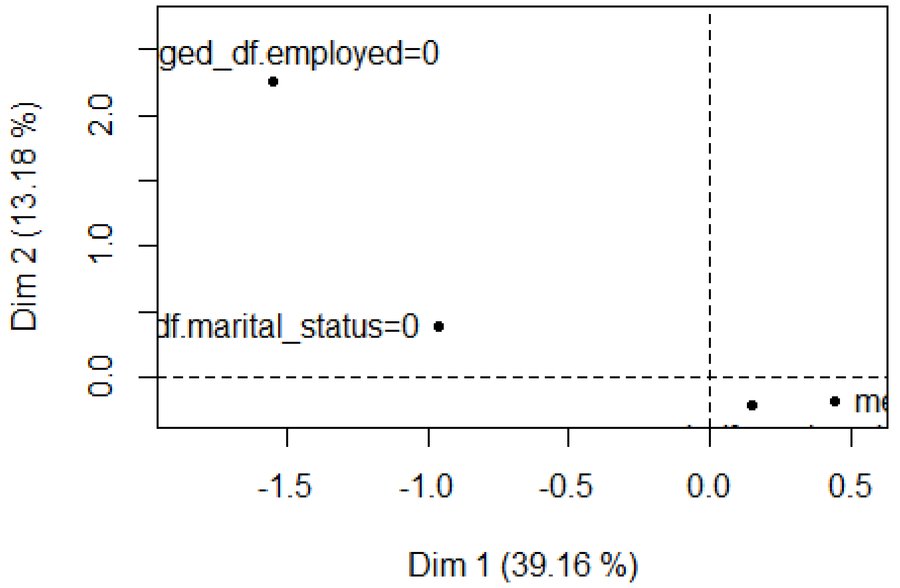

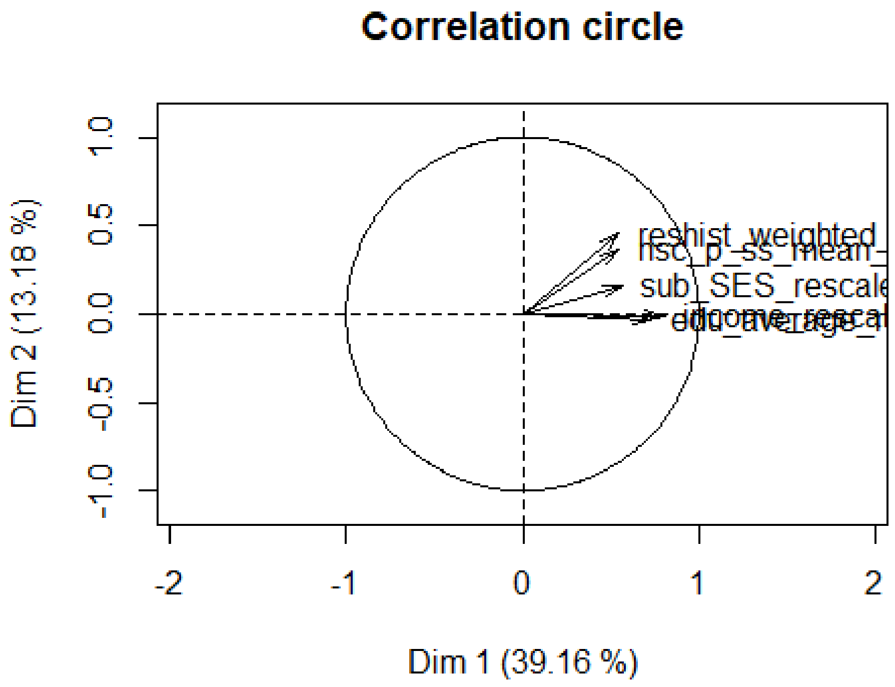


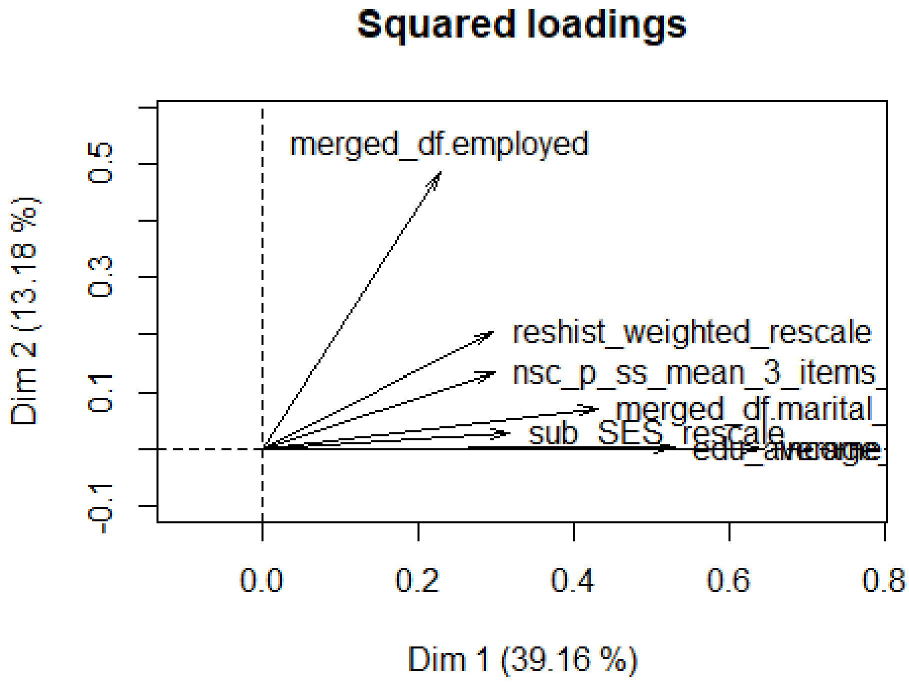


```
# SES_factor_scores ---------------------------------------------------------
#fit_PCAmix$ind$coord #factor (component) scores
fit_PCAmix$eig #eigenvalues and % variance explained by each component
```

```
##       Eigenvalue Proportion Cumulative
## dim 1  2.7413510  39.162157   39.16216
## dim 2  0.9223616  13.176594   52.33875
## dim 3  0.7848404  11.212005   63.55076
## dim 4  0.7673274  10.961820   74.51258
## dim 5  0.7162696  10.232423   84.74500
## dim 6  0.6693712   9.562446   94.30745
## dim 7  0.3984788   5.692554  100.00000

fit_PCAmix$sqload #factor(component) loadings

##                                  dim 1        dim 2        dim 3         d
im 4
## sub_SES_rescale              0.3174741 2.667689e-02 0.5208302181 1.484611
e-09
## reshist_weighted_rescale     0.2959331 2.043658e-01 0.2164826317 3.820776
e-02
## nsc_p_ss_mean_3_items_rescale 0.2982979 1.353574e-01 0.0036273894 5.065040
e-01
## edu_average_rescale          0.5288097 1.188647e-03 0.0058096501 4.025448
e-02
## income_rescale               0.6405717 5.607137e-05 0.0004570532 5.581240
e-02
## merged_df.marital_status     0.4305694 7.061298e-02 0.0037420828 2.373145
e-02
## merged_df.employed           0.2296951 4.841038e-01 0.0338913475 1.028173
e-01
##                                    dim 5
## sub_SES_rescale              0.113996241
## reshist_weighted_rescale     0.242501427
## nsc_p_ss_mean_3_items_rescale 0.043499265
## edu_average_rescale          0.151357719
## income_rescale               0.058355170
## merged_df.marital_status     0.001279327
## merged_df.employed           0.105280447

# SES_factor_scores -------------------------------------------------------



general_ses_fact_alt_PCA= fit_PCAmix$ind$coord[,1]
describe(general_ses_fact_alt_PCA)

##    vars    n mean   sd median trimmed  mad   min  max range  skew kurtosis
se
## X1    1 9972    0 1.66   0.34    0.12 1.59 -5.91 2.98  8.89 -0.66    -0.09
0.02

completeData_general_SES_alt$general_ses_PCA_z= scale(general_ses_fact_alt_PC
A, center = TRUE, scale = TRUE)
describe(completeData_general_SES_alt$general_ses_PCA_z)
```

```
##    vars    n mean sd median trimmed  mad   min max range  skew kurtosis
se
## X1    1 9972    0  1    0.2    0.07 0.96 -3.57 1.8  5.37 -0.66    -0.09 0.
01

general_ses_PCA_z <- c("merged_df.marital_status", "merged_df.employed", "sub
_SES_rescale", "reshist_weighted_rescale", "nsc_p_ss_mean_3_items_rescale", "
edu_average_rescale", "income_rescale","general_ses_PCA_z", "merged_df.subjec
tkey")
general_ses_PCA_z_df <- completeData_general_SES_alt[general_ses_PCA_z]
merged_df=merge(merged_df, general_ses_PCA_z_df, by.x="subjectkey", by.y= "me
rged_df.subjectkey", all=T)

# SES correlations

SES_correlations <- merged_df[,c(2505,2506,2507,2508,2509,2503,2504,2510)]
#apa.cor.table(SES_correlations, filename="APA_SES_full_Connor.doc", table.nu
mber=1)





#Kink variable


merged_df$African0_.9 <- ifelse(merged_df$African >= .9, "1", ifelse(merged_d
f$African < .9, "0", NA))
merged_df$African0_.9_num <- as.numeric(merged_df$African0_.9)
merged_df$upper_kink <- merged_df$African0_.9_num*merged_df$African
summary(merged_df$upper_kink)

##    Min. 1st Qu.  Median    Mean 3rd Qu.    Max.
## 0.00000 0.00000 0.00000 0.01397 0.00000 0.99996

describe(merged_df$upper_kink)

##    vars    n mean   sd median trimmed mad min max range skew kurtosis se
## X1    1 9972 0.01 0.11      0       0   0   0   1     1 8.01    62.22  0

#Desnisty Plots_AA
merged_df_African_ancestry005 = filter(merged_df, African >=.005)
dens_African_ancestry005 <- density(merged_df_African_ancestry005$African)

# plot density_AA

plot(dens_African_ancestry005, frame = FALSE, col = "steelblue",
     main = "",
     xlab="Proportion of African Ancestry",
     xlim=c(0,1),
```

```
     ylim=c(0,5),
     xaxs="i",
     yaxs="i")
```

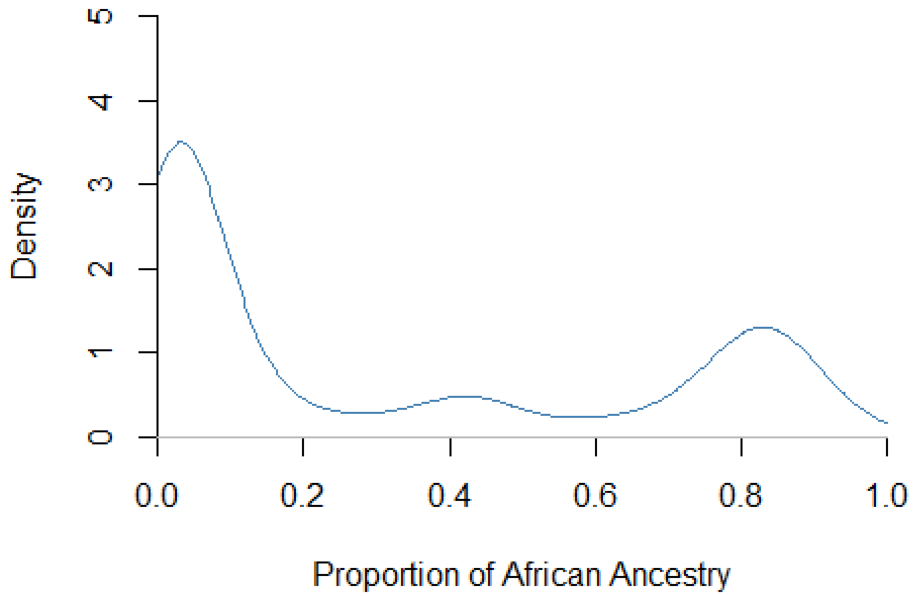


```
#Desnisty Plots_EA
merged_df_European_ancestry005 = filter(merged_df, European >=.005)
dens_European_ancestry005 <- density(merged_df_European_ancestry005$European)

# plot density_EA

plot(dens_European_ancestry005, frame = FALSE, col = "steelblue",
     main = "",
     xlab="% European Ancestry",
     xlim=c(0,1),
     ylim=c(0,5),
     xaxs="i",
     yaxs="i")
```

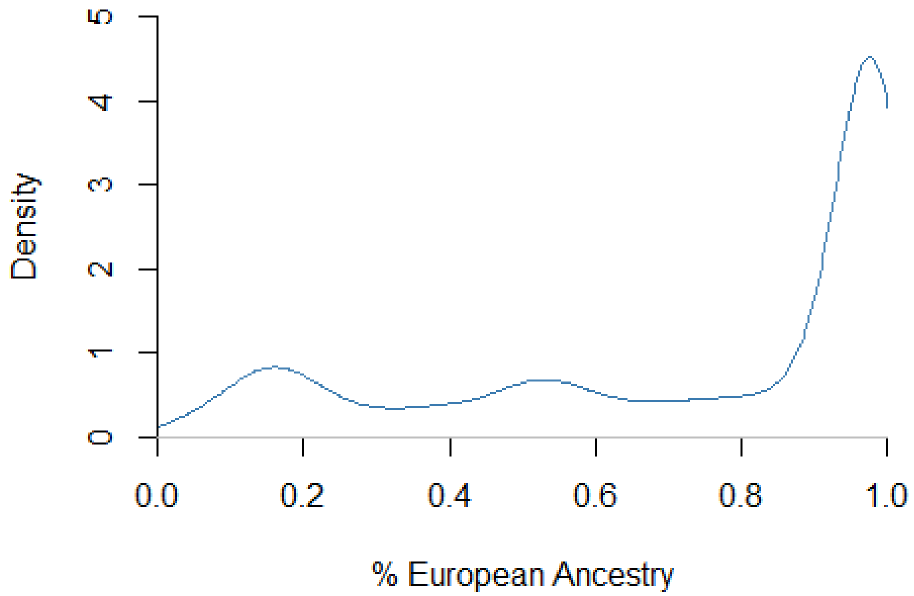


```
#npplreg analysis

#library(np)

#bw <- npplregbw(formula=CA_Z_adj ~ South_Asian + Amerindian + East_Asian + f
rac_Black_SIRE + frac_EastAsian_SIRE + frac_SouthAsian_SIRE +
 #                  frac_Native_American_SIRE + frac_Other_SIRE + frac_Hispani
c_SIRE | African, merged_df)


#summary(bw)
#pl <- npplreg(bws=bw, residuals=TRUE)
#summary(pl)
#coef(pl)
#coef(pl, errors = TRUE)
#summary(pl$resid)
#describe(pl$resid)

#par(mar = rep(3, 5))
#plot(pl$resid)


#merged_df$CA_Z_adj_hat <- 0.7376529*merged_df$South_Asian - 1.191356*merged_
df$Amerindian + 0.6923586*merged_df$East_Asian -0.1288999*merged_df$frac_Blac
k_SIRE -0.1368878*merged_df$frac_EastAsian_SIRE + 0.1523298*merged_df$frac_So
uthAsian_SIRE -0.2577826*merged_df$frac_Native_American_SIRE -0.07611056*merg
```

```
ed_df$frac_Other_SIRE -0.1202465*merged_df$frac_Hispanic_SIRE
#merged_df$CA_Z_adj_sub <- (merged_df$CA_Z_adj - merged_df$CA_Z_adj_hat) -.26
20
#bw2 <- npregbw(formula=merged_df$CA_Z_adj_sub ~ merged_df$African)


#summary(bw2)
#plot(npreg(bw2),xlim=c(0,1), ylim=c(-1,0), xaxs="i", xlab="Proportion of Afr
ican Ancestry", ylab="Test Scores")
#par(new=TRUE)
# create pairs of data points
#x = c(0,1)
#y =  c(0,-1.00136)
# Create a normal plot
#plot(x, y,type="l", xlim=c(0,1), ylim=c(-1,0),
#     xlab="Proportion of African Ancestry", ylab="Test Scores")


#  AA Deciles

merged_df_African_interval1 = filter(merged_df, African >=.0000 & African <.0
005)
describe(merged_df_African_interval1$African)

##    vars    n mean sd median trimmed mad min max range  skew kurtosis se
## X1    1 4622    0  0      0       0   0   0   0     0 16.91   307.58  0

merged_df_African_interval2 = filter(merged_df, African >= .0005 & African <
.1 )
describe(merged_df_African_interval2$African)

##    vars    n mean   sd median trimmed  mad min max range skew kurtosis se
## X1    1 2935 0.03 0.02   0.02    0.03 0.02   0 0.1   0.1 1.02     0.14  0

merged_df_African_interval3 = filter(merged_df, African >= .1 & African < .2
)
describe(merged_df_African_interval3$African)

##    vars   n mean   sd median trimmed  mad min max range skew kurtosis se
## X1    1 286 0.14 0.03   0.14    0.14 0.03 0.1 0.2   0.1 0.34    -1.02  0

merged_df_African_interval4 = filter(merged_df, African >= .2 & African < .3
)
describe(merged_df_African_interval4$African)

##    vars   n mean   sd median trimmed  mad min max range skew kurtosis se
## X1    1 125 0.24 0.03   0.24    0.24 0.04 0.2 0.3   0.1 0.27    -1.16  0

merged_df_African_interval5 = filter(merged_df, African >= .3 & African < .4
)
describe(merged_df_African_interval5$African)
```

```
##    vars   n mean   sd median trimmed  mad min max range  skew kurtosis se
## X1    1 165 0.36 0.03   0.36    0.36 0.04 0.3 0.4   0.1 -0.29    -1.35  0

merged_df_African_interval6 = filter(merged_df, African >= .4 & African < .5
)
describe(merged_df_African_interval6$African)

##    vars   n mean   sd median trimmed  mad min max range skew kurtosis se
## X1    1 279 0.44 0.03   0.44    0.44 0.03 0.4 0.5   0.1 0.51    -0.68  0

merged_df_African_interval7 = filter(merged_df, African >= .5 & African < .6
)
describe(merged_df_African_interval7$African)

##    vars  n mean   sd median trimmed  mad min max range  skew kurtosis se
## X1    1 88 0.55 0.03   0.56    0.55 0.04 0.5 0.6   0.1 -0.17    -1.42  0

merged_df_African_interval8 = filter(merged_df, African >= .6 & African < .7
)
describe(merged_df_African_interval8$African)

##    vars   n mean   sd median trimmed  mad min max range  skew kurtosis se
## X1    1 130 0.65 0.03   0.65    0.65 0.04 0.6 0.7   0.1 -0.13    -1.31  0

merged_df_African_interval9 = filter(merged_df, African >= .7 & African < .8
)
describe(merged_df_African_interval9$African)

##    vars   n mean   sd median trimmed  mad min max range  skew kurtosis se
## X1    1 406 0.76 0.03   0.77    0.77 0.03 0.7 0.8   0.1 -0.61    -0.74  0

merged_df_African_interval10 = filter(merged_df, African >= .8 & African < .9
)
describe(merged_df_African_interval10$African)

##    vars   n mean   sd median trimmed  mad min max range skew kurtosis se
## X1    1 787 0.85 0.03   0.84    0.85 0.03 0.8 0.9   0.1 0.11    -1.07  0

merged_df_African_interval11 = filter(merged_df, African >= .9 & African < .9
95 )
describe(merged_df_African_interval11$African)

##    vars   n mean   sd median trimmed  mad min  max range skew kurtosis se
## X1    1 137 0.93 0.02   0.92    0.93 0.02 0.9 0.99  0.09 1.03     0.28  0

merged_df_African_interval12 = filter(merged_df, African >= .995 & African <=
1 )
describe(merged_df_African_interval12$African)

##    vars  n mean sd median trimmed mad min max range skew kurtosis se
## X1    1 12    1  0      1       1   0   1   1     0  NaN      NaN  0
```

```
#  European Deciles

merged_df_Eur_interval1 = filter(merged_df, European >=.0000 & European <.000
5)
describe(merged_df_Eur_interval1$European)

##    vars  n mean sd median trimmed mad min max range skew kurtosis se
## X1    1 15    0  0      0       0   0   0   0     0  NaN      NaN  0

merged_df_Eur_interval2 = filter(merged_df, European >= .0005 & European < .1
)
describe(merged_df_Eur_interval2$European)

##    vars   n mean   sd median trimmed  mad min max range  skew kurtosis se
## X1    1 283 0.06 0.03   0.07    0.06 0.03   0 0.1   0.1 -0.33     -1.1  0

merged_df_Eur_interval3 = filter(merged_df, European >= .1 & European < .2 )
describe(merged_df_Eur_interval3$European)

##    vars   n mean   sd median trimmed  mad min max range  skew kurtosis se
## X1    1 908 0.15 0.03   0.15    0.15 0.03 0.1 0.2   0.1 -0.07    -1.11  0

merged_df_Eur_interval4 = filter(merged_df, European >= .2 & European < .3 )
describe(merged_df_Eur_interval4$European)

##    vars   n mean   sd median trimmed  mad min max range skew kurtosis se
## X1    1 425 0.24 0.03   0.24    0.24 0.03 0.2 0.3   0.1 0.43    -1.01  0

merged_df_Eur_interval5 = filter(merged_df, European >= .3 & European < .4 )
describe(merged_df_Eur_interval5$European)

##    vars   n mean   sd median trimmed  mad min max range  skew kurtosis se
## X1    1 346 0.35 0.03   0.36    0.35 0.04 0.3 0.4   0.1 -0.19    -1.21  0

merged_df_Eur_interval6 = filter(merged_df, European >= .4 & European < .5 )
describe(merged_df_Eur_interval6$European)

##    vars   n mean   sd median trimmed  mad min max range  skew kurtosis se
## X1    1 461 0.46 0.03   0.46    0.46 0.04 0.4 0.5   0.1 -0.29    -1.27  0

merged_df_Eur_interval7 = filter(merged_df, European >= .5 & European < .6 )
describe(merged_df_Eur_interval7$European)

##    vars   n mean   sd median trimmed  mad min max range skew kurtosis se
## X1    1 700 0.55 0.03   0.55    0.54 0.04 0.5 0.6   0.1 0.11    -1.22  0

merged_df_Eur_interval8 = filter(merged_df, European >= .6 & European < .7 )
describe(merged_df_Eur_interval8$European)

##    vars   n mean   sd median trimmed  mad min max range skew kurtosis se
## X1    1 406 0.65 0.03   0.65    0.65 0.04 0.6 0.7   0.1 0.01    -1.33  0
```

```
merged_df_Eur_interval9 = filter(merged_df, European >= .7 & European < .8 )
describe(merged_df_Eur_interval9$European)

##    vars   n mean   sd median trimmed  mad min max range skew kurtosis se
## X1    1 462 0.75 0.03   0.75    0.75 0.04 0.7 0.8   0.1 -0.1     -1.2  0

merged_df_Eur_interval10 = filter(merged_df, European >= .8 & European < .9 )
describe(merged_df_Eur_interval10$European)

##    vars   n mean   sd median trimmed  mad min max range  skew kurtosis se
## X1    1 514 0.86 0.03   0.86    0.86 0.04 0.8 0.9   0.1 -0.24    -1.18  0

merged_df_Eur_interval11 = filter(merged_df, European >= .9 & European < .995
)
describe(merged_df_Eur_interval11$European)

##    vars    n mean   sd median trimmed  mad min  max range  skew kurtosis s
e
## X1    1 5113 0.97 0.02   0.98    0.98 0.01 0.9 0.99  0.09 -1.51     2.03
0

merged_df_Eur_interval12 = filter(merged_df, European >= .995 & European <= 1
)
describe(merged_df_Eur_interval12$European)

##    vars   n mean sd median trimmed mad min max range  skew kurtosis se
## X1    1 339    1  0      1       1   0   1   1     0 -0.03    -1.55  0

#  East Asian Deciles

merged_df_East_Asian_interval1 = filter(merged_df, East_Asian >=.0000 & East_
Asian <.0005)
describe(merged_df_East_Asian_interval1$East_Asian)

##    vars    n mean sd median trimmed mad min max range skew kurtosis se
## X1    1 6230    0  0      0       0   0   0   0     0 9.54    94.64  0

merged_df_East_Asian_interval2 = filter(merged_df, East_Asian >= .0005 & East
_Asian < .1 )
describe(merged_df_East_Asian_interval2$East_Asian)

##    vars    n mean   sd median trimmed  mad min max range skew kurtosis se
## X1    1 3225 0.01 0.01   0.01    0.01 0.01   0 0.1   0.1 3.65     18.5  0

merged_df_East_Asian_interval3 = filter(merged_df, East_Asian >= .1 & East_As
ian < .2 )
describe(merged_df_East_Asian_interval3$East_Asian)

##    vars  n mean   sd median trimmed  mad min max range skew kurtosis se
## X1    1 74 0.15 0.03   0.15    0.15 0.04 0.1 0.2   0.1 0.15    -1.34  0
```

```
merged_df_East_Asian_interval4 = filter(merged_df, East_Asian >= .2 & East_As
ian < .3 )
describe(merged_df_East_Asian_interval4$East_Asian)

##    vars  n mean   sd median trimmed  mad  min max range skew kurtosis se
## X1    1 84 0.24 0.02   0.24    0.24 0.02 0.21 0.3  0.09 0.34     -0.4  0

merged_df_East_Asian_interval5 = filter(merged_df, East_Asian >= .3 & East_As
ian < .4 )
describe(merged_df_East_Asian_interval5$East_Asian)

##    vars  n mean   sd median trimmed  mad min  max range skew kurtosis   se
## X1    1 20 0.34 0.03   0.34    0.34 0.03 0.3 0.39  0.09 0.37    -1.37 0.01

merged_df_East_Asian_interval6 = filter(merged_df, East_Asian >= .4 & East_As
ian < .5 )
describe(merged_df_East_Asian_interval6$East_Asian)

##    vars   n mean   sd median trimmed  mad min max range  skew kurtosis se
## X1    1 225 0.47 0.02   0.48    0.48 0.02 0.4 0.5   0.1 -1.21     1.32  0

merged_df_East_Asian_interval7 = filter(merged_df, East_Asian >= .5 & East_As
ian < .6 )
describe(merged_df_East_Asian_interval7$East_Asian)

##    vars  n mean   sd median trimmed mad min  max range skew kurtosis   se
## X1    1 18 0.51 0.02    0.5    0.51   0 0.5 0.57  0.07 2.22     3.27 0.01

merged_df_East_Asian_interval8 = filter(merged_df, East_Asian >= .6 & East_As
ian < .7 )
describe(merged_df_East_Asian_interval8$East_Asian)

##    vars n mean   sd median trimmed  mad  min  max range  skew kurtosis   s
e
## X1    1 4 0.65 0.03   0.66    0.65 0.02 0.61 0.68  0.07 -0.57     -1.8 0.0
2

merged_df_East_Asian_interval9 = filter(merged_df, East_Asian >= .7 & East_As
ian < .8 )
describe(merged_df_East_Asian_interval9$East_Asian)

##    vars n mean   sd median trimmed  mad  min  max range skew kurtosis   se
## X1    1 8 0.75 0.02   0.75    0.75 0.03 0.72 0.78  0.05 0.21    -1.79 0.01

merged_df_East_Asian_interval10 = filter(merged_df, East_Asian >= .8 & East_A
sian < .9 )
describe(merged_df_East_Asian_interval10$East_Asian)

##    vars  n mean   sd median trimmed  mad min max range  skew kurtosis   se
## X1    1 10 0.87 0.03   0.88    0.87 0.02 0.8 0.9   0.1 -1.01    -0.04 0.01
```

```
merged_df_East_Asian_interval11 = filter(merged_df, East_Asian >= .9 & East_A
sian < .995 )
describe(merged_df_East_Asian_interval11$East_Asian)
```

```
##    vars  n mean   sd median trimmed  mad  min  max range  skew kurtosis se
## X1    1 74 0.95 0.02   0.96    0.95 0.02 0.91 0.98  0.08 -0.57    -1.12  0
```

```
merged_df_East_Asian_interval12 = filter(merged_df, East_Asian >= .995 & East
_Asian <= 1 )
describe(merged_df_East_Asian_interval12$East_Asian)
```

```
## Warning in min(x, na.rm = na.rm): no non-missing arguments to min; returni
ng Inf

## Warning in max(x, na.rm = na.rm): no non-missing arguments to max; returni
ng
## -Inf

##    vars n mean sd median trimmed mad min  max range skew kurtosis se
## X1    1 0  NaN NA     NA     NaN  NA Inf -Inf  -Inf   NA       NA NA
```

```
#  Amerindian Deciles

merged_df_Amerindian_interval1 = filter(merged_df, Amerindian >=.0000 & Ameri
ndian <.0005)
describe(merged_df_Amerindian_interval1$Amerindian)
```

```
##    vars    n mean sd median trimmed mad min max range skew kurtosis se
## X1    1 1229    0  0      0       0   0   0   0     0 4.15    16.62  0
```

```
merged_df_Amerindian_interval2 = filter(merged_df, Amerindian >= .0005 & Amer
indian < .1 )
describe(merged_df_Amerindian_interval2$Amerindian)
```

```
##    vars    n mean   sd median trimmed  mad min max range skew kurtosis se
## X1    1 7135 0.01 0.02   0.01    0.01 0.01   0 0.1   0.1 3.09    10.57  0
```

```
merged_df_Amerindian_interval3 = filter(merged_df, Amerindian >= .1 & Amerind
ian < .2 )
describe(merged_df_Amerindian_interval3$Amerindian)
```

```
##    vars   n mean   sd median trimmed  mad min max range skew kurtosis se
## X1    1 443 0.15 0.03   0.15    0.15 0.04 0.1 0.2   0.1 0.12    -1.12  0
```

```
merged_df_Amerindian_interval4 = filter(merged_df, Amerindian >= .2 & Amerind
ian < .3 )
describe(merged_df_Amerindian_interval4$Amerindian)
```

```
##    vars   n mean   sd median trimmed  mad min max range skew kurtosis se
## X1    1 329 0.25 0.03   0.25    0.25 0.03 0.2 0.3   0.1 0.12    -1.11  0
```

```
merged_df_Amerindian_interval5 = filter(merged_df, Amerindian >= .3 & Amerind
ian < .4 )
describe(merged_df_Amerindian_interval5$Amerindian)
```

```
##    vars   n mean   sd median trimmed  mad min max range skew kurtosis se
## X1    1 301 0.35 0.03   0.35    0.35 0.04 0.3 0.4   0.1 -0.1    -1.15  0
```

```
merged_df_Amerindian_interval6 = filter(merged_df, Amerindian >= .4 & Amerind
ian < .5 )
describe(merged_df_Amerindian_interval6$Amerindian)
```

```
##    vars   n mean   sd median trimmed  mad min max range skew kurtosis se
## X1    1 282 0.44 0.03   0.44    0.44 0.03 0.4 0.5   0.1 0.26    -1.14  0
```

```
merged_df_Amerindian_interval7 = filter(merged_df, Amerindian >= .5 & Amerind
ian < .6 )
describe(merged_df_Amerindian_interval7$Amerindian)
```

```
##    vars   n mean   sd median trimmed  mad min max range skew kurtosis se
## X1    1 156 0.54 0.03   0.54    0.54 0.03 0.5 0.6   0.1 0.32    -0.96  0
```

```
merged_df_Amerindian_interval8 = filter(merged_df, Amerindian >= .6 & Amerind
ian < .7 )
describe(merged_df_Amerindian_interval8$Amerindian)
```

```
##    vars  n mean   sd median trimmed  mad min max range skew kurtosis se
## X1    1 75 0.65 0.03   0.65    0.65 0.04 0.6 0.7   0.1 0.15    -1.21  0
```

```
merged_df_Amerindian_interval9 = filter(merged_df, Amerindian >= .7 & Amerind
ian < .8 )
describe(merged_df_Amerindian_interval9$Amerindian)
```

```
##    vars  n mean   sd median trimmed  mad min  max range skew kurtosis se
## X1    1 18 0.72 0.02   0.71    0.72 0.01 0.7 0.77  0.06 0.89    -0.14  0
```

```
merged_df_Amerindian_interval10 = filter(merged_df, Amerindian >= .8 & Amerin
dian < .9 )
describe(merged_df_Amerindian_interval10$Amerindian)
```

```
## Warning in min(x, na.rm = na.rm): no non-missing arguments to min; returni
ng Inf
```

```
## Warning in min(x, na.rm = na.rm): no non-missing arguments to max; returni
ng
## -Inf
```

```
##    vars n mean sd median trimmed mad min  max range skew kurtosis se
## X1    1 0  NaN NA     NA     NaN  NA Inf -Inf  -Inf   NA       NA NA
```

```
merged_df_Amerindian_interval11 = filter(merged_df, Amerindian >= .9 & Amerin
dian < .995 )
describe(merged_df_Amerindian_interval11$Amerindian)
```

```
##    vars n mean   sd median trimmed  mad  min  max range  skew kurtosis   s
e
## X1    1 4 0.96 0.03   0.96    0.96 0.03 0.92 0.98  0.06 -0.11    -2.24 0.0
1

merged_df_Amerindian_interval12 = filter(merged_df, Amerindian >= .995 & Amer
indian <= 1 )
describe(merged_df_Amerindian_interval12$Amerindian)

## Warning in min(x, na.rm = na.rm): no non-missing arguments to min; returni
ng Inf

## Warning in min(x, na.rm = na.rm): no non-missing arguments to max; returni
ng
## -Inf

##    vars n mean sd median trimmed mad min  max range skew kurtosis se
## X1    1 0  NaN NA     NA     NaN  NA Inf -Inf  -Inf   NA       NA NA

#  South_Asian Deciles

merged_df_South_Asian_interval1 = filter(merged_df, South_Asian >=.0000 & Sou
th_Asian <.0005)
describe(merged_df_South_Asian_interval1$South_Asian)

##    vars    n mean sd median trimmed mad min max range skew kurtosis se
## X1    1 5414    0  0      0       0   0   0   0     0 8.93    83.29  0

merged_df_South_Asian_interval2 = filter(merged_df, South_Asian >= .0005 & So
uth_Asian < .1 )
describe(merged_df_South_Asian_interval2$South_Asian)

##    vars    n mean   sd median trimmed  mad min max range skew kurtosis se
## X1    1 4382 0.02 0.02   0.01    0.01 0.01   0 0.1   0.1 1.97     4.38  0

merged_df_South_Asian_interval3 = filter(merged_df, South_Asian >= .1 & South
_Asian < .2 )
describe(merged_df_South_Asian_interval3$South_Asian)

##    vars  n mean   sd median trimmed  mad min max range skew kurtosis se
## X1    1 55 0.14 0.03   0.13    0.14 0.04 0.1 0.2   0.1 0.28    -1.52  0

merged_df_South_Asian_interval4 = filter(merged_df, South_Asian >= .2 & South
_Asian < .3 )
describe(merged_df_South_Asian_interval4$South_Asian)

##    vars  n mean   sd median trimmed  mad min max range skew kurtosis se
## X1    1 30 0.24 0.03   0.24    0.24 0.02 0.2 0.3  0.09 0.68    -0.52  0

merged_df_South_Asian_interval5 = filter(merged_df, South_Asian >= .3 & South
_Asian < .4 )
describe(merged_df_South_Asian_interval5$South_Asian)
```

```
##    vars  n mean   sd median trimmed  mad  min max range skew kurtosis se
## X1    1 29 0.35 0.03   0.35    0.35 0.02 0.31 0.4  0.09 0.03    -1.13  0
```

```
merged_df_South_Asian_interval6 = filter(merged_df, South_Asian >= .4 & South
_Asian < .5 )
describe(merged_df_South_Asian_interval6$South_Asian)
```

```
##    vars  n mean   sd median trimmed  mad  min  max range skew kurtosis   s
e
## X1    1 17 0.44 0.03   0.43    0.44 0.03 0.41 0.49  0.09 0.54    -1.21 0.0
1
```

```
merged_df_South_Asian_interval7 = filter(merged_df, South_Asian >= .5 & South
_Asian < .6 )
describe(merged_df_South_Asian_interval7$South_Asian)
```

```
##    vars n mean   sd median trimmed  mad  min  max range  skew kurtosis   s
e
## X1    1 4 0.53 0.02   0.53    0.53 0.02 0.51 0.56  0.05 -0.09       -2 0.0
1
```

```
merged_df_South_Asian_interval8 = filter(merged_df, South_Asian >= .6 & South
_Asian < .7 )
describe(merged_df_South_Asian_interval8$South_Asian)
```

```
##    vars  n mean   sd median trimmed  mad  min max range skew kurtosis   se
## X1    1 11 0.65 0.02   0.65    0.65 0.02 0.63 0.7  0.07 0.62    -0.46 0.01
```

```
merged_df_South_Asian_interval9 = filter(merged_df, South_Asian >= .7 & South
_Asian < .8 )
describe(merged_df_South_Asian_interval9$South_Asian)
```

```
##    vars n mean   sd median trimmed  mad  min max range skew kurtosis   se
## X1    1 9 0.75 0.03   0.75    0.75 0.03 0.71 0.8  0.09 0.23     -1.4 0.01
```

```
merged_df_South_Asian_interval10 = filter(merged_df, South_Asian >= .8 & Sout
h_Asian < .9 )
describe(merged_df_South_Asian_interval10$South_Asian)
```

```
##    vars  n mean   sd median trimmed  mad min  max range skew kurtosis   se
## X1    1 21 0.83 0.03   0.82    0.83 0.03 0.8 0.89  0.09 0.73    -0.95 0.01
```

```
merged_df_South_Asian_interval11 = filter(merged_df, South_Asian >= .9 & Sout
h_Asian < .995 )
describe(merged_df_South_Asian_interval11$South_Asian)
```

```
## Warning in min(x, na.rm = na.rm): no non-missing arguments to min; returni
ng Inf

## Warning in min(x, na.rm = na.rm): no non-missing arguments to max; returni
ng
## -Inf
```

```
##    vars n mean sd median trimmed mad min  max range skew kurtosis se
## X1    1 0  NaN NA     NA     NaN  NA Inf -Inf  -Inf   NA       NA NA
```

```
merged_df_South_Asian_interval12 = filter(merged_df, South_Asian >= .995 & So
uth_Asian <= 1 )
describe(merged_df_South_Asian_interval12$South_Asian)
```

```
## Warning in min(x, na.rm = na.rm): no non-missing arguments to min; returni
ng Inf

## Warning in min(x, na.rm = na.rm): no non-missing arguments to max; returni
ng
## -Inf

##    vars n mean sd median trimmed mad min  max range skew kurtosis se
## X1    1 0  NaN NA     NA     NaN  NA Inf -Inf  -Inf   NA       NA NA
```

```
#Alternative

density(merged_df$African)
```

```
##
## Call:
##  density.default(x = merged_df$African)
##
## Data: merged_df$African (9972 obs.); Bandwidth 'bw' = 0.009363
##
##        x                  y
##  Min.   :-0.02808   Min.   : 0.000635
##  1st Qu.: 0.23595   1st Qu.: 0.119599
##  Median : 0.49999   Median : 0.180692
##  Mean   : 0.49999   Mean   : 0.946076
##  3rd Qu.: 0.76402   3rd Qu.: 0.459462
##  Max.   : 1.02805   Max.   :23.383350
```

```
summary(merged_df$African)
```

```
##     Min.  1st Qu.   Median     Mean  3rd Qu.     Max.
## 0.000010 0.000010 0.006356 0.158984 0.087922 0.999960
```

```
describe(merged_df$African)
```

```
##    vars    n mean   sd median trimmed  mad min max range skew kurtosis se
## X1    1 9972 0.16 0.29   0.01    0.09 0.01   0   1     1 1.68     1.13  0
```

```
dens_all <- density(merged_df$African)

plot(dens_all, frame = FALSE, col = "steelblue",
     main = "Density Plot of African Ancestry",
     xlab="Proportion of African Ancestry",
     xlim=c(0,1),
     ylim=c(0,30),
```

```
     xaxs="i",
     yaxs="i")
```

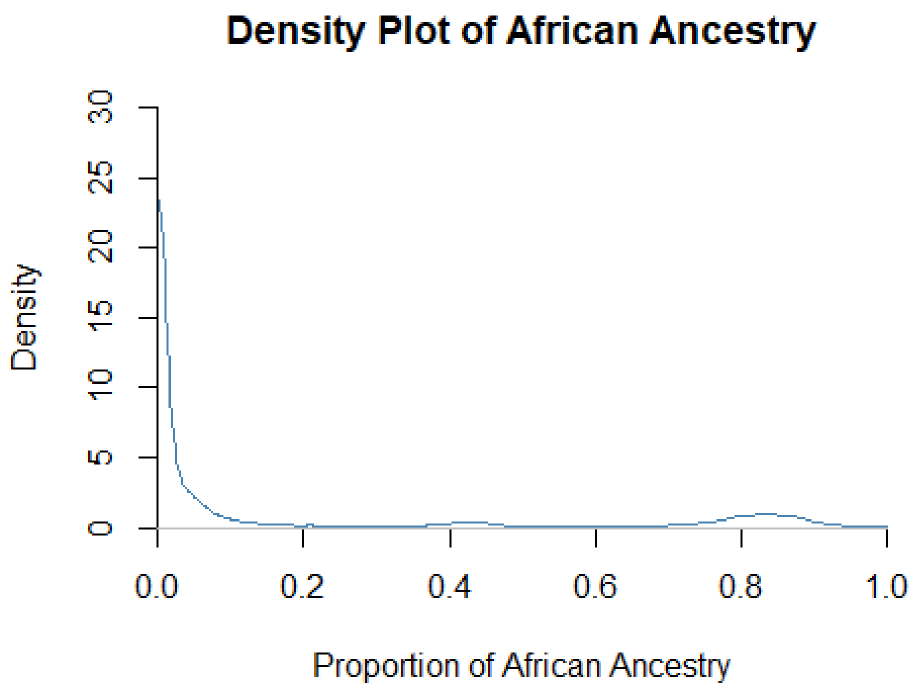


```
#merged_df_all_African_ancestry01 = filter(merged_df, African >=.01)
#summary(merged_df_all_African_ancestry01$African)
#describe(merged_df_all_African_ancestry01$African)
#density(merged_df_all_African_ancestry01$African)


# Compute the density data
#dens <- density(merged_df_all_African_ancestry$African)
# plot density
#plot(dens, frame = FALSE, col = "steelblue",
#    main = "Density Plot of African Ancestry")

# Compute the density data
#dens2 <- density(merged_df_all_African_ancestry01$African)
# plot density
#plot(dens2, frame = FALSE, col = "steelblue",
#     main = "Density Plot of African Ancestry")


#create subsets

##ethnicity
```

```
merged_df_all = merged_df
summary(merged_df_all$race)

##          White_1           Black_1       EastAsian_1 Native American_1
##             7653              1583               130                49
##     Other Race_1       SouthAsian_1
##              508                49

summary(merged_df_all$Hispanic)

##    Min. 1st Qu.  Median    Mean 3rd Qu.    Max.
##  0.0000  0.0000  0.0000  0.1874  0.0000  1.0000

merged_df_all_Model3_spec = filter(merged_df_all, frac_White_SIRE_woc > 0 | f
rac_Black_SIRE_woc > 0 | frac_EastAsian_SIRE_woc > 0 | frac_SouthAsian_SIRE_w
oc > 0 |
                                     frac_Native_American_SIRE_woc > 0 | frac
_Other_Race_SIRE_woc > 0 | frac_Hispanic_SIRE_woc > 0 | NH_Black_White_only >
0 |

                                     NH_SouthAsian_White_only > 0 | NH_EastAs
ian_White_only > 0 | NH_Native_American_White_only > 0 | H_White_only > 0 | H
_Black_only > 0 | H_Other_only > 0)

describe(merged_df_all_Model3_spec$CA_Z_adj)

##    vars    n mean sd median trimmed  mad  min  max range skew kurtosis   s
e
## X1    1 9972    0  1  -0.04   -0.02 0.96 -3.9 5.59  9.49 0.29     0.34 0.0
1

#Models

#Descriptives


aggregate(cbind(age/12, CA_Z_adj, general_ses_PCA_z, European, African, East_
Asian, South_Asian, Amerindian)~NH_White_only, data=merged_df, FUN=function(x
) c(mean = round(mean(x, na.rm=TRUE), 2), sd = round(sd(x, na.rm=TRUE), 2), c
ount=length(x)))

##   NH_White_only V1.mean   V1.sd V1.count CA_Z_adj.mean CA_Z_adj.sd
## 1             0    9.90    0.62  4439.00         -0.31        1.01
## 2             1    9.93    0.63  5533.00          0.25        0.92
##   CA_Z_adj.count general_ses_PCA_z.mean general_ses_PCA_z.sd
## 1        4439.00                  -0.50                 1.04
## 2        5533.00                   0.40                 0.75
##   general_ses_PCA_z.count European.mean European.sd European.count African
.mean
## 1                 4439.00          0.45        0.27        4439.00
0.35
```

```
## 2                     5533.00           0.97        0.05        5533.00
0.01
##   African.sd African.count East_Asian.mean East_Asian.sd East_Asian.count
## 1       0.35       4439.00            0.06          0.17          4439.00
## 2       0.02       5533.00            0.00          0.02          5533.00
##   South_Asian.mean South_Asian.sd South_Asian.count Amerindian.mean
## 1             0.02           0.09           4439.00            0.13
## 2             0.01           0.02           5533.00            0.01
##   Amerindian.sd Amerindian.count
## 1          0.18          4439.00
## 2          0.03          5533.00
```

```
table(merged_df$NH_White_only, merged_df$Child_US_Born)
```

```
##
##        0    1
##   0  195 4244
##   1   74 5459
```

```
aggregate(cbind(age/12, CA_Z_adj, general_ses_PCA_z, European, African, East_
Asian, South_Asian, Amerindian)~NH_Black_only, data=merged_df, FUN=function(x
) c(mean = round(mean(x, na.rm=TRUE), 2), sd = round(sd(x, na.rm=TRUE), 2), c
ount=length(x)))
```

```
##   NH_Black_only V1.mean   V1.sd V1.count CA_Z_adj.mean CA_Z_adj.sd
## 1             0    9.91    0.63  8538.00          0.13        0.96
## 2             1    9.91    0.61  1434.00         -0.77        0.87
##   CA_Z_adj.count general_ses_PCA_z.mean general_ses_PCA_z.sd
## 1        8538.00                   0.17                 0.91
## 2        1434.00                  -1.00                 0.95
##   general_ses_PCA_z.count European.mean European.sd European.count African
.mean
## 1                 8538.00          0.83        0.23        8538.00
0.05
## 2                 1434.00          0.18        0.11        1434.00
0.80
##   African.sd African.count East_Asian.mean East_Asian.sd East_Asian.count
## 1       0.13       8538.00            0.03          0.13          8538.00
## 2       0.11       1434.00            0.00          0.02          1434.00
##   South_Asian.mean South_Asian.sd South_Asian.count Amerindian.mean
## 1             0.02           0.06           8538.00            0.07
## 2             0.00           0.01           1434.00            0.01
##   Amerindian.sd Amerindian.count
## 1          0.14          8538.00
## 2          0.02          1434.00
```

```
table(merged_df$NH_Black_only, merged_df$Child_US_Born)
```

```
##
##        0    1
```

```
##    0  237 8301
##    1   32 1402
```

```
aggregate(cbind(age/12, CA_Z_adj, general_ses_PCA_z, European, African, East_
Asian, South_Asian, Amerindian)~NH_EastAsian_only, data=merged_df, FUN=functi
on(x) c(mean = round(mean(x, na.rm=TRUE), 2), sd = round(sd(x, na.rm=TRUE), 2
), count=length(x)))
```

```
##   NH_EastAsian_only V1.mean   V1.sd V1.count CA_Z_adj.mean CA_Z_adj.sd
## 1                 0    9.91    0.62  9865.00         -0.01        1.00
## 2                 1   10.02    0.62   107.00          0.57        1.02
##   CA_Z_adj.count general_ses_PCA_z.mean general_ses_PCA_z.sd
## 1        9865.00                  -0.01                 1.00
## 2         107.00                   0.60                 0.75
##   general_ses_PCA_z.count European.mean European.sd European.count African
.mean
## 1                 9865.00          0.74        0.31        9865.00
0.16
## 2                  107.00          0.14        0.18         107.00
0.01
##   African.sd African.count East_Asian.mean East_Asian.sd East_Asian.count
## 1       0.29       9865.00            0.02          0.08          9865.00
## 2       0.05        107.00            0.82          0.23           107.00
##   South_Asian.mean South_Asian.sd South_Asian.count Amerindian.mean
## 1             0.01           0.06           9865.00            0.06
## 2             0.02           0.10            107.00            0.01
##   Amerindian.sd Amerindian.count
## 1          0.13          9865.00
## 2          0.02           107.00
```

```
table(merged_df$NH_EastAsian_only, merged_df$Child_US_Born)
```

```
## 
##         0    1
##    0  250 9615
##    1   19   88
```

```
aggregate(cbind(age/12, CA_Z_adj, general_ses_PCA_z, European, African, East_
Asian, South_Asian, Amerindian)~NH_SouthAsian_only, data=merged_df, FUN=funct
ion(x) c(mean = round(mean(x, na.rm=TRUE), 2), sd = round(sd(x, na.rm=TRUE),
2), count=length(x)))
```

```
##   NH_SouthAsian_only V1.mean   V1.sd V1.count CA_Z_adj.mean CA_Z_adj.sd
## 1                  0    9.91    0.62  9929.00          0.00        1.00
## 2                  1   10.03    0.68    43.00          0.45        1.02
##   CA_Z_adj.count general_ses_PCA_z.mean general_ses_PCA_z.sd
## 1        9929.00                   0.00                 1.00
## 2          43.00                   0.88                 0.46
##   general_ses_PCA_z.count European.mean European.sd European.count African
.mean
## 1                 9929.00          0.74        0.31        9929.00
```

```
0.16
## 2                     43.00          0.24        0.13          43.00
0.00
##   African.sd African.count East_Asian.mean East_Asian.sd East_Asian.count
## 1       0.29       9929.00            0.03          0.12          9929.00
## 2       0.00         43.00            0.03          0.07            43.00
##   South_Asian.mean South_Asian.sd South_Asian.count Amerindian.mean
## 1             0.01           0.04           9929.00            0.06
## 2             0.73           0.14             43.00            0.01
##   Amerindian.sd Amerindian.count
## 1          0.13          9929.00
## 2          0.01            43.00
```

```
table(merged_df$NH_SouthAsian_only, merged_df$Child_US_Born)
```

```
##
##        0    1
##   0  261 9668
##   1    8   35
```

```
aggregate(cbind(age/12, CA_Z_adj, general_ses_PCA_z, European, African, East_
Asian, South_Asian, Amerindian)~NH_Native_American_only, data=merged_df, FUN=
function(x) c(mean = round(mean(x, na.rm=TRUE), 2), sd = round(sd(x, na.rm=TR
UE), 2), count=length(x)))
```

```
##   NH_Native_American_only V1.mean   V1.sd V1.count CA_Z_adj.mean CA_Z_adj.
sd
## 1                       0    9.91    0.63  9941.00          0.00        1.
00
## 2                       1    9.70    0.60    31.00         -0.42        0.
79
##   CA_Z_adj.count general_ses_PCA_z.mean general_ses_PCA_z.sd
## 1       9941.00                   0.00                 1.00
## 2         31.00                  -0.81                 0.72
##   general_ses_PCA_z.count European.mean European.sd European.count African
.mean
## 1                 9941.00          0.74        0.31        9941.00
0.16
## 2                   31.00          0.71        0.30          31.00
0.11
##   African.sd African.count East_Asian.mean East_Asian.sd East_Asian.count
## 1       0.29       9941.00            0.03          0.12          9941.00
## 2       0.26         31.00            0.01          0.02            31.00
##   South_Asian.mean South_Asian.sd South_Asian.count Amerindian.mean
## 1             0.01           0.06           9941.00            0.06
## 2             0.01           0.01             31.00            0.15
##   Amerindian.sd Amerindian.count
## 1          0.13          9941.00
## 2          0.19            31.00
```

```
table(merged_df$NH_Native_American_only, merged_df$Child_US_Born)
```

```
##
##         0    1
##   0   269 9672
##   1     0   31
```

```
aggregate(cbind(age/12, CA_Z_adj, general_ses_PCA_z, European, African, East_
Asian, South_Asian, Amerindian)~NH_Other_Race_only, data=merged_df, FUN=funct
ion(x) c(mean = round(mean(x, na.rm=TRUE), 2), sd = round(sd(x, na.rm=TRUE),
2), count=length(x)))
```

```
##   NH_Other_Race_only V1.mean   V1.sd V1.count CA_Z_adj.mean CA_Z_adj.sd
## 1                  0    9.91    0.63  9875.00          0.00        1.00
## 2                  1    9.96    0.61    97.00         -0.22        1.12
##   CA_Z_adj.count general_ses_PCA_z.mean general_ses_PCA_z.sd
## 1        9875.00                   0.01                 1.00
## 2          97.00                  -0.53                 1.11
##   general_ses_PCA_z.count European.mean European.sd European.count African
.mean
## 1                 9875.00          0.74        0.31        9875.00
0.16
## 2                   97.00          0.55        0.30          97.00
0.28
##   African.sd African.count East_Asian.mean East_Asian.sd East_Asian.count
## 1       0.29       9875.00            0.03          0.12          9875.00
## 2       0.31         97.00            0.06          0.19            97.00
##   South_Asian.mean South_Asian.sd South_Asian.count Amerindian.mean
## 1             0.01           0.06           9875.00            0.06
## 2             0.04           0.11             97.00            0.07
##   Amerindian.sd Amerindian.count
## 1          0.13          9875.00
## 2          0.15            97.00
```

```
table(merged_df$NH_Other_Race_only, merged_df$Child_US_Born)
```

```
##
##         0    1
##   0   259 9616
##   1    10   87
```

```
aggregate(cbind(age/12, CA_Z_adj, general_ses_PCA_z, European, African, East_
Asian, South_Asian, Amerindian)~Hispanic, data=merged_df, FUN=function(x) c(m
ean = round(mean(x, na.rm=TRUE), 2), sd = round(sd(x, na.rm=TRUE), 2), count=
length(x)))
```

```
##   Hispanic V1.mean   V1.sd V1.count CA_Z_adj.mean CA_Z_adj.sd CA_Z_adj.cou
nt
## 1        0    9.92    0.62  8103.00          0.05        1.00        8103.
00
## 2        1    9.88    0.63  1869.00         -0.23        0.96        1869.
00
##   general_ses_PCA_z.mean general_ses_PCA_z.sd general_ses_PCA_z.count
```

```
## 1                  0.10                  0.99                  8103.00
## 2                 -0.41                  0.93                  1869.00
##   European.mean European.sd European.count African.mean African.sd
## 1          0.77        0.33        8103.00         0.17       0.32
## 2          0.60        0.20        1869.00         0.10       0.14
##   African.count East_Asian.mean East_Asian.sd East_Asian.count South_Asian
.mean
## 1       8103.00            0.03          0.13          8103.00
0.02
## 2       1869.00            0.02          0.06          1869.00
0.01
##   South_Asian.sd South_Asian.count Amerindian.mean Amerindian.sd
## 1           0.07           8103.00            0.01          0.04
## 2           0.02           1869.00            0.27          0.18
##   Amerindian.count
## 1          8103.00
## 2          1869.00
```

```
table(merged_df$Hispanic, merged_df$Child_US_Born)
```

```
##
##        0    1
##   0  155 7948
##   1  114 1755
```

```
aggregate(cbind(age/12, CA_Z_adj, general_ses_PCA_z, European, African, East_
Asian, South_Asian, Amerindian)~H_White_only, data=merged_df, FUN=function(x)
c(mean = round(mean(x, na.rm=TRUE), 2), sd = round(sd(x, na.rm=TRUE), 2), cou
nt=length(x)))
```

```
##   H_White_only V1.mean   V1.sd V1.count CA_Z_adj.mean CA_Z_adj.sd
## 1            0    9.92    0.62  8801.00          0.03        1.00
## 2            1    9.89    0.64  1171.00         -0.19        0.96
##   CA_Z_adj.count general_ses_PCA_z.mean general_ses_PCA_z.sd
## 1        8801.00                   0.04                 1.00
## 2        1171.00                  -0.29                 0.91
##   general_ses_PCA_z.count European.mean European.sd European.count African
.mean
## 1                 8801.00          0.75        0.33        8801.00
0.17
## 2                 1171.00          0.67        0.18        1171.00
0.06
##   African.sd African.count East_Asian.mean East_Asian.sd East_Asian.count
## 1       0.31       8801.00            0.03          0.13          8801.00
## 2       0.06       1171.00            0.01          0.02          1171.00
##   South_Asian.mean South_Asian.sd South_Asian.count Amerindian.mean
## 1             0.02           0.06           8801.00            0.04
## 2             0.01           0.01           1171.00            0.26
##   Amerindian.sd Amerindian.count
## 1          0.10          8801.00
## 2          0.17          1171.00
```

```
table(merged_df$H_White_only, merged_df$Child_US_Born)
```

```
##
##         0    1
##   0   195 8606
##   1    74 1097
```

```
aggregate(cbind(age/12, CA_Z_adj, general_ses_PCA_z, European, African, East_
Asian, South_Asian, Amerindian)~H_Black_only, data=merged_df, FUN=function(x)
c(mean = round(mean(x, na.rm=TRUE), 2), sd = round(sd(x, na.rm=TRUE), 2), cou
nt=length(x)))
```

```
##   H_Black_only V1.mean   V1.sd V1.count CA_Z_adj.mean CA_Z_adj.sd
## 1            0    9.91    0.62  9888.00          0.00        1.00
## 2            1    9.80    0.64    84.00         -0.34        0.93
##   CA_Z_adj.count general_ses_PCA_z.mean general_ses_PCA_z.sd
## 1        9888.00                   0.01                 1.00
## 2          84.00                  -0.64                 0.98
##   general_ses_PCA_z.count European.mean European.sd European.count African
.mean
## 1                 9888.00          0.74        0.31        9888.00
0.16
## 2                   84.00          0.35        0.13          84.00
0.53
##   African.sd African.count East_Asian.mean East_Asian.sd East_Asian.count
## 1       0.29       9888.00            0.03          0.12          9888.00
## 2       0.17         84.00            0.00          0.01            84.00
##   South_Asian.mean South_Asian.sd South_Asian.count Amerindian.mean
## 1             0.01           0.06           9888.00            0.06
## 2             0.00           0.01             84.00            0.11
##   Amerindian.sd Amerindian.count
## 1          0.13          9888.00
## 2          0.09            84.00
```

```
table(merged_df$H_Black_only, merged_df$Child_US_Born)
```

```
##
##         0    1
##   0   264 9624
##   1     5   79
```

```
aggregate(cbind(age/12, CA_Z_adj, general_ses_PCA_z, European, African, East_
Asian, South_Asian, Amerindian)~H_Other_only, data=merged_df, FUN=function(x)
c(mean = round(mean(x, na.rm=TRUE), 2), sd = round(sd(x, na.rm=TRUE), 2), cou
nt=length(x)))
```

```
##   H_Other_only V1.mean   V1.sd V1.count CA_Z_adj.mean CA_Z_adj.sd
## 1            0    9.91    0.62  9561.00          0.02        1.00
## 2            1    9.89    0.63   411.00         -0.45        0.92
##   CA_Z_adj.count general_ses_PCA_z.mean general_ses_PCA_z.sd
## 1        9561.00                   0.03                 0.99
```

```
## 2         411.00                  -0.75                 0.88
##   general_ses_PCA_z.count European.mean European.sd European.count African
.mean
## 1                 9561.00          0.75        0.32        9561.00
0.16
## 2                  411.00          0.49        0.15         411.00
0.09
##   African.sd African.count East_Asian.mean East_Asian.sd East_Asian.count
## 1       0.30       9561.00            0.03          0.12          9561.00
## 2       0.11        411.00            0.02          0.03           411.00
##   South_Asian.mean South_Asian.sd South_Asian.count Amerindian.mean
## 1             0.01           0.06           9561.00            0.05
## 2             0.00           0.01            411.00            0.40
##   Amerindian.sd Amerindian.count
## 1          0.11          9561.00
## 2          0.17           411.00
```

```
table(merged_df$H_Other_only, merged_df$Child_US_Born)
```

```
## 
##        0    1
##   0  241 9320
##   1   28  383
```

```
#aggregate(cbind(age/12, CA_Z_adj, general_ses_PCA_z, European, African, East
_Asian, South_Asian, Amerindian)~Hispanic_only, data=merged_df, FUN=function(
x) c(mean = round(mean(x, na.rm=TRUE), 2), count=length(x)))
#table(merged_df$Hispanic_only, merged_df$Child_US_Born)

aggregate(cbind(age/12, CA_Z_adj, general_ses_PCA_z, European, African, East_
Asian, South_Asian, Amerindian)~NH_Black_White_only, data=merged_df, FUN=func
tion(x) c(mean = round(mean(x, na.rm=TRUE), 2), sd = round(sd(x, na.rm=TRUE),
2), count=length(x)))
```

```
##   NH_Black_White_only V1.mean   V1.sd V1.count CA_Z_adj.mean CA_Z_adj.sd
## 1                   0    9.91    0.63  9670.00          0.00        1.00
## 2                   1    9.88    0.62   302.00         -0.13        0.95
##   CA_Z_adj.count general_ses_PCA_z.mean general_ses_PCA_z.sd
## 1        9670.00                   0.01                 0.99
## 2         302.00                  -0.45                 1.06
##   general_ses_PCA_z.count European.mean European.sd European.count African
.mean
## 1                 9670.00          0.74        0.32        9670.00
0.15
## 2                  302.00          0.58        0.12         302.00
0.41
##   African.sd African.count East_Asian.mean East_Asian.sd East_Asian.count
## 1       0.29       9670.00            0.03          0.12          9670.00
## 2       0.12        302.00            0.00          0.01           302.00
##   South_Asian.mean South_Asian.sd South_Asian.count Amerindian.mean
## 1             0.01           0.06           9670.00            0.06
```

```
## 2              0.00            0.01             302.00            0.01
##   Amerindian.sd Amerindian.count
## 1          0.14          9670.00
## 2          0.02           302.00
```

```
table(merged_df$NH_Black_White_only, merged_df$Child_US_Born)
```

```
##
##        0    1
##    0  268 9402
##    1    1  301
```

```
aggregate(cbind(age/12, CA_Z_adj, general_ses_PCA_z, European, African, East_
Asian, South_Asian, Amerindian)~NH_EastAsian_White_only, data=merged_df, FUN=
function(x) c(mean = round(mean(x, na.rm=TRUE), 2), sd = round(sd(x, na.rm=TR
UE), 2), count=length(x)))
```

```
##   NH_EastAsian_White_only V1.mean   V1.sd V1.count CA_Z_adj.mean CA_Z_adj.
sd
## 1                       0    9.91    0.62  9723.00         -0.01        1.
00
## 2                       1    9.99    0.64   249.00          0.58        0.
98
##   CA_Z_adj.count general_ses_PCA_z.mean general_ses_PCA_z.sd
## 1        9723.00                  -0.02                 1.00
## 2         249.00                   0.66                 0.67
##   general_ses_PCA_z.count European.mean European.sd European.count African
.mean
## 1                 9723.00          0.74        0.32        9723.00
0.16
## 2                  249.00          0.56        0.12         249.00
0.01
##   African.sd African.count East_Asian.mean East_Asian.sd East_Asian.count
## 1       0.30       9723.00            0.02          0.10          9723.00
## 2       0.01        249.00            0.41          0.14           249.00
##   South_Asian.mean South_Asian.sd South_Asian.count Amerindian.mean
## 1             0.01           0.06           9723.00            0.06
## 2             0.02           0.04            249.00            0.01
##   Amerindian.sd Amerindian.count
## 1          0.13          9723.00
## 2          0.01           249.00
```

```
table(merged_df$NH_EastAsian_White_only, merged_df$Child_US_Born)
```

```
##
##        0    1
##    0  264 9459
##    1    5  244
```

```
aggregate(cbind(age/12, CA_Z_adj, general_ses_PCA_z, European, African, East_
Asian, South_Asian, Amerindian)~NH_Native_American_White_only, data=merged_df
```

```
, FUN=function(x) c(mean = round(mean(x, na.rm=TRUE), 2), sd = round(sd(x, na
.rm=TRUE), 2), count=length(x)))
```

```
##   NH_Native_American_White_only V1.mean   V1.sd V1.count CA_Z_adj.mean
## 1                             0    9.92    0.63  9841.00          0.00
## 2                             1    9.78    0.60   131.00          0.01
##   CA_Z_adj.sd CA_Z_adj.count general_ses_PCA_z.mean general_ses_PCA_z.sd
## 1        1.00        9841.00                   0.00                 1.00
## 2        0.86         131.00                  -0.09                 0.91
##   general_ses_PCA_z.count European.mean European.sd European.count African
.mean
## 1                 9841.00          0.73        0.32        9841.00
0.16
## 2                  131.00          0.90        0.10         131.00
0.01
##   African.sd African.count East_Asian.mean East_Asian.sd East_Asian.count
## 1       0.29       9841.00            0.03          0.12          9841.00
## 2       0.03        131.00            0.01          0.02           131.00
##   South_Asian.mean South_Asian.sd South_Asian.count Amerindian.mean
## 1             0.01           0.06           9841.00            0.06
## 2             0.01           0.01            131.00            0.07
##   Amerindian.sd Amerindian.count
## 1          0.13          9841.00
## 2          0.09           131.00
```

```
table(merged_df$NH_Native_American_White_only, merged_df$Child_US_Born)
```

```
##
##        0    1
##   0  269 9572
##   1    0  131
```

```
aggregate(cbind(age/12, CA_Z_adj, general_ses_PCA_z, European, African, East_
Asian, South_Asian, Amerindian)~NH_SouthAsian_White_only, data=merged_df, FUN
=function(x) c(mean = round(mean(x, na.rm=TRUE), 2), sd = round(sd(x, na.rm=T
RUE), 2), count=length(x)))
```

```
##   NH_SouthAsian_White_only V1.mean   V1.sd V1.count CA_Z_adj.mean CA_Z_adj
.sd
## 1                        0    9.91    0.63  9932.00          0.00        1
.00
## 2                        1    9.79    0.54    40.00          0.83        0
.84
##   CA_Z_adj.count general_ses_PCA_z.mean general_ses_PCA_z.sd
## 1        9932.00                   0.00                 1.00
## 2          40.00                   0.78                 0.77
##   general_ses_PCA_z.count European.mean European.sd European.count African
.mean
## 1                 9932.00          0.74        0.32        9932.00
0.16
## 2                   40.00          0.63        0.11          40.00
```

```
0.00
##   African.sd African.count East_Asian.mean East_Asian.sd East_Asian.count
## 1       0.29       9932.00            0.03          0.12          9932.00
## 2       0.00         40.00            0.02          0.08            40.00
##   South_Asian.mean South_Asian.sd South_Asian.count Amerindian.mean
## 1             0.01           0.06           9932.00            0.06
## 2             0.34           0.12             40.00            0.01
##   Amerindian.sd Amerindian.count
## 1          0.13          9932.00
## 2          0.01            40.00
```

```
table(merged_df$NH_SouthAsian_White_only, merged_df$Child_US_Born)
```

```
##
##        0    1
##   0  268 9664
##   1    1   39
```

```
aggregate(cbind(age/12, CA_Z_adj, general_ses_PCA_z, European, African, East_
Asian, South_Asian, Amerindian)~Any_Other, data=merged_df, FUN=function(x) c(
mean = round(mean(x, na.rm=TRUE), 2), sd = round(sd(x, na.rm=TRUE), 2), count
=length(x)))
```

```
##   Any_Other V1.mean   V1.sd V1.count CA_Z_adj.mean CA_Z_adj.sd CA_Z_adj.co
unt
## 1         0    9.91    0.63  9836.00          0.00        1.00        9836
.00
## 2         1    9.87    0.62   136.00         -0.27        1.06         136
.00
##   general_ses_PCA_z.mean general_ses_PCA_z.sd general_ses_PCA_z.count
## 1                   0.01                 1.00                 9836.00
## 2                  -0.63                 1.03                  136.00
##   European.mean European.sd European.count African.mean African.sd
## 1          0.74        0.31        9836.00         0.15       0.29
## 2          0.37        0.21         136.00         0.46       0.25
##   African.count East_Asian.mean East_Asian.sd East_Asian.count South_Asian
.mean
## 1       9836.00            0.03          0.12          9836.00
0.01
## 2        136.00            0.13          0.20           136.00
0.02
##   South_Asian.sd South_Asian.count Amerindian.mean Amerindian.sd
## 1           0.06           9836.00            0.06          0.13
## 2           0.07            136.00            0.02          0.05
##   Amerindian.count
## 1          9836.00
## 2           136.00
```

```
table(merged_df$Any_Other, merged_df$Child_US_Born)
```

```
##
##        0    1
##   0  264 9572
##   1    5  131
```

```r
#Model 1: ancestral proportions and single-SIRE categories - no restriction t
est

model_1=lmer(CA_Z_adj ~ South_Asian + Amerindian + East_Asian + African
             + frac_Black_SIRE
             + frac_EastAsian_SIRE
             + frac_SouthAsian_SIRE
             + frac_Native_American_SIRE
             + frac_Other_SIRE
             + frac_Hispanic_SIRE
             + (1|site_id_l) + (1|site_id_l:rel_family_id), data=merged_df, R
EML = FALSE)
summary(model_1)
```

```
## Warning in site_id_l:rel_family_id: numerical expression has 9972 elements
: only
## the first used

## Warning in site_id_l:rel_family_id: numerical expression has 9972 elements
: only
## the first used

## Warning in site_id_l:rel_family_id: numerical expression has 9972 elements
: only
## the first used

## Warning in site_id_l:rel_family_id: numerical expression has 9972 elements
: only
## the first used

## Warning in site_id_l:rel_family_id: numerical expression has 9972 elements
: only
## the first used

## Warning in site_id_l:rel_family_id: numerical expression has 9972 elements
: only
## the first used

## Warning in site_id_l:rel_family_id: numerical expression has 9972 elements
: only
## the first used

## Warning in site_id_l:rel_family_id: numerical expression has 9972 elements
: only
## the first used
```

```
## Warning in site_id_l:rel_family_id: numerical expression has 9972 elements
: only
## the first used

## Warning in site_id_l:rel_family_id: numerical expression has 9972 elements
: only
## the first used

## Warning in site_id_l:rel_family_id: numerical expression has 9972 elements
: only
## the first used

## Warning in site_id_l:rel_family_id: numerical expression has 9972 elements
: only
## the first used

## Warning in site_id_l:rel_family_id: numerical expression has 9972 elements
: only
## the first used

## Warning in site_id_l:rel_family_id: numerical expression has 9972 elements
: only
## the first used

## Warning in site_id_l:rel_family_id: numerical expression has 9972 elements
: only
## the first used

## Warning in site_id_l:rel_family_id: numerical expression has 9972 elements
: only
## the first used

## Linear mixed model fit by maximum likelihood  ['lmerMod']
## Formula: CA_Z_adj ~ South_Asian + Amerindian + East_Asian + African +
##     frac_Black_SIRE + frac_EastAsian_SIRE + frac_SouthAsian_SIRE +
##     frac_Native_American_SIRE + frac_Other_SIRE + frac_Hispanic_SIRE +
##     (1 | site_id_l) + (1 | site_id_l:rel_family_id)
##    Data: merged_df
##
##      AIC      BIC   logLik deviance df.resid
##  26073.5  26174.4 -13022.8  26045.5     9958
##
## Scaled residuals:
##     Min      1Q  Median      3Q     Max
## -3.7367 -0.5233 -0.0435  0.4714  4.6289
##
## Random effects:
##  Groups                  Name        Variance Std.Dev.
```

```
##  site_id_l:rel_family_id (Intercept) 0.37361  0.6112
##  site_id_l               (Intercept) 0.01885  0.1373
##  Residual                            0.44956  0.6705
## Number of obs: 9972, groups:  site_id_l:rel_family_id, 8419; site_id_l, 22
##
## Fixed effects:
##                            Estimate Std. Error t value
## (Intercept)                 0.30100    0.03321   9.064
## South_Asian                 0.48674    0.32639   1.491
## Amerindian                 -1.38037    0.12140 -11.370
## East_Asian                  0.64389    0.20217   3.185
## African                    -1.02422    0.12284  -8.338
## frac_Black_SIRE            -0.14201    0.10111  -1.405
## frac_EastAsian_SIRE        -0.21204    0.17580  -1.206
## frac_SouthAsian_SIRE       -0.07348    0.26062  -0.282
## frac_Native_American_SIRE -0.14276    0.10705  -1.334
## frac_Other_SIRE            -0.19670    0.07802  -2.521
## frac_Hispanic_SIRE         -0.09395    0.08427  -1.115
```

```
performance::icc(model_1)
```

```
## # Intraclass Correlation Coefficient
##
##      Adjusted ICC: 0.466
##   Conditional ICC: 0.393
```

```
performance::r2(model_1)
```

```
## # R2 for Mixed Models
##
##   Conditional R2: 0.550
##      Marginal R2: 0.157
```

```
#Model 2: ancestral proportions, single SIRE categories and 7 chosen multi-SI
RE categories - restriction test

model_2=lmer(CA_Z_adj ~ South_Asian + Amerindian + East_Asian + African
             + frac_Black_SIRE_woc
             + frac_EastAsian_SIRE_woc
             + frac_SouthAsian_SIRE_woc
             + frac_Native_American_SIRE_woc
             + frac_Other_Race_SIRE_woc
             + frac_Hispanic_SIRE_woc
             + NH_Black_White_only
             + NH_SouthAsian_White_only
             + NH_EastAsian_White_only
             + NH_Native_American_White_only
             + H_White_only
             + H_Black_only
             + H_Other_only
             + (1|site_id_l) + (1|site_id_l:rel_family_id), data=merged_df, R
```

```
EML = FALSE)
summary(model_2)

## Warning in site_id_l:rel_family_id: numerical expression has 9972 elements
: only
## the first used

## Warning in site_id_l:rel_family_id: numerical expression has 9972 elements
: only
## the first used

## Warning in site_id_l:rel_family_id: numerical expression has 9972 elements
: only
## the first used

## Warning in site_id_l:rel_family_id: numerical expression has 9972 elements
: only
## the first used

## Warning in site_id_l:rel_family_id: numerical expression has 9972 elements
: only
## the first used

## Warning in site_id_l:rel_family_id: numerical expression has 9972 elements
: only
## the first used

## Warning in site_id_l:rel_family_id: numerical expression has 9972 elements
: only
## the first used

## Warning in site_id_l:rel_family_id: numerical expression has 9972 elements
: only
## the first used

## Warning in site_id_l:rel_family_id: numerical expression has 9972 elements
: only
## the first used

## Warning in site_id_l:rel_family_id: numerical expression has 9972 elements
: only
## the first used

## Warning in site_id_l:rel_family_id: numerical expression has 9972 elements
: only
## the first used

## Warning in site_id_l:rel_family_id: numerical expression has 9972 elements
: only
```

```
## the first used

## Warning in site_id_l:rel_family_id: numerical expression has 9972 elements
: only
## the first used

## Warning in site_id_l:rel_family_id: numerical expression has 9972 elements
: only
## the first used

## Warning in site_id_l:rel_family_id: numerical expression has 9972 elements
: only
## the first used

## Warning in site_id_l:rel_family_id: numerical expression has 9972 elements
: only
## the first used

## Linear mixed model fit by maximum likelihood  ['lmerMod']
## Formula: CA_Z_adj ~ South_Asian + Amerindian + East_Asian + African +
##     frac_Black_SIRE_woc + frac_EastAsian_SIRE_woc + frac_SouthAsian_SIRE_w
oc +
##     frac_Native_American_SIRE_woc + frac_Other_Race_SIRE_woc +
##     frac_Hispanic_SIRE_woc + NH_Black_White_only + NH_SouthAsian_White_onl
y +
##     NH_EastAsian_White_only + NH_Native_American_White_only +
##     H_White_only + H_Black_only + H_Other_only + (1 | site_id_l) +
##     (1 | site_id_l:rel_family_id)
##    Data: merged_df
##
##      AIC      BIC   logLik deviance df.resid
##  26060.5  26211.9 -13009.2  26018.5     9951
##
## Scaled residuals:
##     Min      1Q  Median      3Q     Max
## -3.7237 -0.5245 -0.0432  0.4733  4.6672
##
## Random effects:
##  Groups                  Name        Variance Std.Dev.
##  site_id_l:rel_family_id (Intercept) 0.37088  0.6090
##  site_id_l               (Intercept) 0.01844  0.1358
##  Residual                            0.44980  0.6707
## Number of obs: 9972, groups:  site_id_l:rel_family_id, 8419; site_id_l, 22
##
## Fixed effects:
##                                Estimate Std. Error t value
## (Intercept)                    0.294890   0.033011   8.933
## South_Asian                    0.543943   0.327815   1.659
## Amerindian                    -1.335821   0.123882 -10.783
## East_Asian                     0.660174   0.202965   3.253
```

```
## African                          -1.029756   0.125197  -8.225
## frac_Black_SIRE_woc              -0.150326   0.102834  -1.462
## frac_EastAsian_SIRE_woc          -0.352299   0.182740  -1.928
## frac_SouthAsian_SIRE_woc         -0.267888   0.269723  -0.993
## frac_Native_American_SIRE_woc -0.185570   0.135215  -1.372
## frac_Other_Race_SIRE_woc         -0.179576   0.101808  -1.764
## frac_Hispanic_SIRE_woc            0.339005   0.196570   1.725
## NH_Black_White_only               0.053278   0.074907   0.711
## NH_SouthAsian_White_only          0.336091   0.186694   1.800
## NH_EastAsian_White_only           0.004296   0.102153   0.042
## NH_Native_American_White_only -0.099930   0.084357  -1.185
## H_White_only                     -0.081305   0.045454  -1.789
## H_Black_only                      0.036120   0.122835   0.294
## H_Other_only                     -0.165678   0.069026  -2.400
```

```
##
## Correlation matrix not shown by default, as p = 18 > 12.
## Use print(x, correlation=TRUE)  or
##     vcov(x)        if you need it
```

```
performance::icc(model_2)
```

```
## # Intraclass Correlation Coefficient
##
##      Adjusted ICC: 0.464
##   Conditional ICC: 0.390
```

```
performance::r2(model_2)
```

```
## # R2 for Mixed Models
##
##   Conditional R2: 0.550
##      Marginal R2: 0.160
```

```
#For Restriction Test
# the test applies to models 2, 3, 6, 8a and 8b

coeffvec <- coef(model_2)
varcov <- vcov(model_2, full=FALSE)
```

```
## Warning in site_id_l:rel_family_id: numerical expression has 9972 elements
: only
## the first used

## Warning in site_id_l:rel_family_id: numerical expression has 9972 elements
: only
## the first used

## Warning in site_id_l:rel_family_id: numerical expression has 9972 elements
: only
## the first used
```

```
## Warning in site_id_l:rel_family_id: numerical expression has 9972 elements
: only
## the first used

## Warning in site_id_l:rel_family_id: numerical expression has 9972 elements
: only
## the first used

## Warning in site_id_l:rel_family_id: numerical expression has 9972 elements
: only
## the first used

## Warning in site_id_l:rel_family_id: numerical expression has 9972 elements
: only
## the first used

## Warning in site_id_l:rel_family_id: numerical expression has 9972 elements
: only
## the first used
```

```
varcov
```

```
## 18 x 18 Matrix of class "dgeMatrix"
##                [,1]         [,2]         [,3]         [,4]         [,5
]
##  [1,]  1.089702e-03 -0.0016919742 -0.0003248982 -0.0004161887 -0.000273801
5
##  [2,] -1.691974e-03  0.1074627066  0.0029553784  0.0122650732  0.005085401
4
##  [3,] -3.248982e-04  0.0029553784  0.0153468402  0.0010638535  0.002086864
3
##  [4,] -4.161887e-04  0.0122650732  0.0010638535  0.0411949073  0.003421129
6
##  [5,] -2.738015e-04  0.0050854014  0.0020868643  0.0034211296  0.015674382
9
##  [6,] -4.863816e-06 -0.0026669657 -0.0015828064 -0.0025553051 -0.012303002
5
##  [7,]  1.198616e-04 -0.0104900199 -0.0004383571 -0.0327622849 -0.003103790
8
##  [8,]  9.761001e-04 -0.0760421212 -0.0019861684 -0.0095689419 -0.003743811
5
##  [9,] -9.513949e-05 -0.0008681135 -0.0024163513 -0.0017912201 -0.003542884
4
## [10,] -2.574126e-05 -0.0049257457 -0.0014905339 -0.0035434188 -0.004493307
8
## [11,] -1.966832e-04  0.0004304433 -0.0063185727  0.0011900806 -0.001517522
2
## [12,] -9.251167e-05 -0.0010833521 -0.0007913929 -0.0012994966 -0.006170715
7
```

```
## [13,]  3.580343e-04 -0.0351547697 -0.0009023477 -0.0046212964 -0.001649651
0
## [14,] -2.858465e-05 -0.0051248168 -0.0002297898 -0.0164516073 -0.001382117
2
## [15,] -1.581221e-04  0.0003398944 -0.0007907175 -0.0002333280 -0.000228752
2
## [16,] -1.400277e-04 -0.0001057062 -0.0036849175 -0.0004517136 -0.001210938
9
## [17,] -6.264296e-05 -0.0014575986 -0.0025535864 -0.0017061928 -0.008163383
8
## [18,] -1.040079e-04 -0.0003686854 -0.0055950994 -0.0008607627 -0.002110306
4
##                [,6]          [,7]          [,8]          [,9]         [,10
]
##  [1,] -4.863816e-06  0.0001198616  9.761001e-04 -9.513949e-05 -2.574126e-0
5
##  [2,] -2.666966e-03 -0.0104900199 -7.604212e-02 -8.681135e-04 -4.925746e-0
3
##  [3,] -1.582806e-03 -0.0004383571 -1.986168e-03 -2.416351e-03 -1.490534e-0
3
##  [4,] -2.555305e-03 -0.0327622849 -9.568942e-03 -1.791220e-03 -3.543419e-0
3
##  [5,] -1.230300e-02 -0.0031037908 -3.743812e-03 -3.542884e-03 -4.493308e-0
3
##  [6,]  1.057474e-02  0.0024790079  2.149139e-03  2.887397e-03  3.694942e-0
3
##  [7,]  2.479008e-03  0.0333937758  8.418767e-03  1.935152e-03  3.137613e-0
3
##  [8,]  2.149139e-03  0.0084187669  7.275023e-02  9.782450e-04  3.805566e-0
3
##  [9,]  2.887397e-03  0.0019351525  9.782450e-04  1.828305e-02  1.384932e-0
3
## [10,]  3.694942e-03  0.0031376131  3.805566e-03  1.384932e-03  1.036489e-0
2
## [11,]  1.292176e-03 -0.0033274506 -6.551623e-04 -5.399951e-03  9.293187e-0
4
## [12,]  5.090968e-03  0.0013585778  9.951562e-04  1.641914e-03  1.923894e-0
3
## [13,]  1.033680e-03  0.0042088532  2.519095e-02  4.766738e-04  1.815985e-0
3
## [14,]  1.203865e-03  0.0133874310  4.241236e-03  8.724652e-04  1.604841e-0
3
## [15,]  3.593683e-04  0.0003329228 -7.811707e-05  9.424408e-04  2.689939e-0
4
## [16,]  1.149161e-03  0.0005321098  2.647847e-04  9.354580e-04  7.356380e-0
4
## [17,]  6.667216e-03  0.0017305341  1.303781e-03  2.238406e-03  2.592881e-0
3
## [18,]  1.832296e-03  0.0009196182  4.815302e-04  1.373959e-03  1.128508e-0
3
```

```
##               [,11]         [,12]         [,13]         [,14]         [,15
]
##  [1,] -1.966832e-04 -9.251167e-05  3.580343e-04 -2.858465e-05 -1.581221e-0
4
##  [2,]  4.304433e-04 -1.083352e-03 -3.515477e-02 -5.124817e-03  3.398944e-0
4
##  [3,] -6.318573e-03 -7.913929e-04 -9.023477e-04 -2.297898e-04 -7.907175e-0
4
##  [4,]  1.190081e-03 -1.299497e-03 -4.621296e-03 -1.645161e-02 -2.333280e-0
4
##  [5,] -1.517522e-03 -6.170716e-03 -1.649651e-03 -1.382117e-03 -2.287522e-0
4
##  [6,]  1.292176e-03  5.090968e-03  1.033680e-03  1.203865e-03  3.593683e-0
4
##  [7,] -3.327451e-03  1.358578e-03  4.208853e-03  1.338743e-02  3.329228e-0
4
##  [8,] -6.551623e-04  9.951562e-04  2.519095e-02  4.241236e-03 -7.811707e-0
5
##  [9,] -5.399951e-03  1.641914e-03  4.766738e-04  8.724652e-04  9.424408e-0
4
## [10,]  9.293187e-04  1.923894e-03  1.815985e-03  1.604841e-03  2.689939e-0
4
## [11,]  3.863964e-02  8.578214e-04  9.202374e-05 -1.644750e-04  5.462919e-0
4
## [12,]  8.578214e-04  5.611085e-03  5.121305e-04  6.936420e-04  2.886982e-0
4
## [13,]  9.202374e-05  5.121305e-04  3.485468e-02  2.156628e-03  4.287971e-0
5
## [14,] -1.644750e-04  6.936420e-04  2.156628e-03  1.043516e-02  2.439338e-0
4
## [15,]  5.462919e-04  2.886982e-04  4.287971e-05  2.439338e-04  7.116111e-0
3
## [16,]  2.067421e-03  6.506733e-04  1.942084e-04  3.536636e-04  3.932445e-0
4
## [17,]  1.836495e-03  3.421402e-03  6.450000e-04  8.740130e-04  3.939479e-0
4
## [18,]  2.976524e-03  9.994005e-04  3.000158e-04  5.434390e-04  5.096368e-0
4
##               [,16]         [,17]         [,18]
##  [1,] -0.0001400277 -6.264296e-05 -0.0001040079
##  [2,] -0.0001057062 -1.457599e-03 -0.0003686854
##  [3,] -0.0036849175 -2.553586e-03 -0.0055950994
##  [4,] -0.0004517136 -1.706193e-03 -0.0008607627
##  [5,] -0.0012109389 -8.163384e-03 -0.0021103064
##  [6,]  0.0011491612  6.667216e-03  0.0018322956
##  [7,]  0.0005321098  1.730534e-03  0.0009196182
##  [8,]  0.0002647847  1.303781e-03  0.0004815302
##  [9,]  0.0009354580  2.238406e-03  0.0013739589
## [10,]  0.0007356380  2.592881e-03  0.0011285080
## [11,]  0.0020674209  1.836495e-03  0.0029765239
```

```
## [12,]  0.0006506733  3.421402e-03  0.0009994005
## [13,]  0.0001942084  6.450000e-04  0.0003000158
## [14,]  0.0003536636  8.740130e-04  0.0005434390
## [15,]  0.0003932445  3.939479e-04  0.0005096368
## [16,]  0.0020660902  1.286499e-03  0.0018062727
## [17,]  0.0012864989  1.508833e-02  0.0019533857
## [18,]  0.0018062727  1.953386e-03  0.0047645688
```

```
#Model 3: same as model 2 but with smaller data set (cutting out 50 multi-SIR
E observations which do not match the 7 categories) - restriction test

model_3=lmer(CA_Z_adj ~ South_Asian + Amerindian + East_Asian + African
             + frac_Black_SIRE_woc
             + frac_EastAsian_SIRE_woc
             + frac_SouthAsian_SIRE_woc
             + frac_Native_American_SIRE_woc
             + frac_Other_Race_SIRE_woc
             + frac_Hispanic_SIRE_woc
             + NH_Black_White_only
             + NH_SouthAsian_White_only
             + NH_EastAsian_White_only
             + NH_Native_American_White_only
             + H_White_only
             + H_Black_only
             + H_Other_only
             + (1|site_id_l) + (1|site_id_l:rel_family_id), data=merged_df_al
l_Model3_spec, REML = FALSE)
summary(model_3)
```

```
## Warning in site_id_l:rel_family_id: numerical expression has 9972 elements
: only
## the first used

## Warning in site_id_l:rel_family_id: numerical expression has 9972 elements
: only
## the first used

## Warning in site_id_l:rel_family_id: numerical expression has 9972 elements
: only
## the first used

## Warning in site_id_l:rel_family_id: numerical expression has 9972 elements
: only
## the first used

## Warning in site_id_l:rel_family_id: numerical expression has 9972 elements
: only
## the first used

## Warning in site_id_l:rel_family_id: numerical expression has 9972 elements
```

```
: only
## the first used

## Warning in site_id_l:rel_family_id: numerical expression has 9972 elements
: only
## the first used

## Warning in site_id_l:rel_family_id: numerical expression has 9972 elements
: only
## the first used

## Warning in site_id_l:rel_family_id: numerical expression has 9972 elements
: only
## the first used

## Warning in site_id_l:rel_family_id: numerical expression has 9972 elements
: only
## the first used

## Warning in site_id_l:rel_family_id: numerical expression has 9972 elements
: only
## the first used

## Warning in site_id_l:rel_family_id: numerical expression has 9972 elements
: only
## the first used

## Warning in site_id_l:rel_family_id: numerical expression has 9972 elements
: only
## the first used

## Warning in site_id_l:rel_family_id: numerical expression has 9972 elements
: only
## the first used

## Warning in site_id_l:rel_family_id: numerical expression has 9972 elements
: only
## the first used

## Warning in site_id_l:rel_family_id: numerical expression has 9972 elements
: only
## the first used

## Linear mixed model fit by maximum likelihood  ['lmerMod']
## Formula: CA_Z_adj ~ South_Asian + Amerindian + East_Asian + African +
##     frac_Black_SIRE_woc + frac_EastAsian_SIRE_woc + frac_SouthAsian_SIRE_w
oc +
##     frac_Native_American_SIRE_woc + frac_Other_Race_SIRE_woc +
##     frac_Hispanic_SIRE_woc + NH_Black_White_only + NH_SouthAsian_White_onl
```

```
y +
##     NH_EastAsian_White_only + NH_Native_American_White_only +
##     H_White_only + H_Black_only + H_Other_only + (1 | site_id_l) +
##     (1 | site_id_l:rel_family_id)
##    Data: merged_df_all_Model3_spec
##
##      AIC      BIC   logLik deviance df.resid
##  26060.5  26211.9 -13009.2  26018.5     9951
##
## Scaled residuals:
##     Min      1Q  Median      3Q     Max
## -3.7237 -0.5245 -0.0432  0.4733  4.6672
##
## Random effects:
##  Groups                 Name        Variance Std.Dev.
##  site_id_l:rel_family_id (Intercept) 0.37088  0.6090
##  site_id_l              (Intercept) 0.01844  0.1358
##  Residual                           0.44980  0.6707
## Number of obs: 9972, groups:  site_id_l:rel_family_id, 8419; site_id_l, 22
##
## Fixed effects:
##                                Estimate Std. Error t value
## (Intercept)                    0.294890   0.033011   8.933
## South_Asian                    0.543943   0.327815   1.659
## Amerindian                    -1.335821   0.123882 -10.783
## East_Asian                     0.660174   0.202965   3.253
## African                       -1.029756   0.125197  -8.225
## frac_Black_SIRE_woc           -0.150326   0.102834  -1.462
## frac_EastAsian_SIRE_woc       -0.352299   0.182740  -1.928
## frac_SouthAsian_SIRE_woc      -0.267888   0.269723  -0.993
## frac_Native_American_SIRE_woc -0.185570   0.135215  -1.372
## frac_Other_Race_SIRE_woc      -0.179576   0.101808  -1.764
## frac_Hispanic_SIRE_woc         0.339005   0.196570   1.725
## NH_Black_White_only            0.053278   0.074907   0.711
## NH_SouthAsian_White_only       0.336091   0.186694   1.800
## NH_EastAsian_White_only        0.004296   0.102153   0.042
## NH_Native_American_White_only -0.099930   0.084357  -1.185
## H_White_only                  -0.081305   0.045454  -1.789
## H_Black_only                   0.036120   0.122835   0.294
## H_Other_only                  -0.165678   0.069026  -2.400

##
## Correlation matrix not shown by default, as p = 18 > 12.
## Use print(x, correlation=TRUE)  or
##     vcov(x)        if you need it
```

```
performance::icc(model_3)
```

```
## # Intraclass Correlation Coefficient
##
```

```
##      Adjusted ICC: 0.464
##   Conditional ICC: 0.390
```

```
performance::r2(model_3)
```

```
## # R2 for Mixed Models
##
##   Conditional R2: 0.550
##      Marginal R2: 0.160
```

```
#For Restriction Test
# the test applies to models 2, 3, 6, 8a and 8b

coeffvec <- coef(model_3)
varcov <- vcov(model_3, full=FALSE)
```

```
## Warning in site_id_l:rel_family_id: numerical expression has 9972 elements
: only
## the first used

## Warning in site_id_l:rel_family_id: numerical expression has 9972 elements
: only
## the first used

## Warning in site_id_l:rel_family_id: numerical expression has 9972 elements
: only
## the first used

## Warning in site_id_l:rel_family_id: numerical expression has 9972 elements
: only
## the first used

## Warning in site_id_l:rel_family_id: numerical expression has 9972 elements
: only
## the first used

## Warning in site_id_l:rel_family_id: numerical expression has 9972 elements
: only
## the first used

## Warning in site_id_l:rel_family_id: numerical expression has 9972 elements
: only
## the first used

## Warning in site_id_l:rel_family_id: numerical expression has 9972 elements
: only
## the first used
```

```
varcov
```

```
## 18 x 18 Matrix of class "dgeMatrix"
##                [,1]          [,2]          [,3]          [,4]          [,5
]
##  [1,]  1.089702e-03 -0.0016919742 -0.0003248982 -0.0004161887 -0.000273801
5
##  [2,] -1.691974e-03  0.1074627066  0.0029553784  0.0122650732  0.005085401
4
##  [3,] -3.248982e-04  0.0029553784  0.0153468402  0.0010638535  0.002086864
3
##  [4,] -4.161887e-04  0.0122650732  0.0010638535  0.0411949073  0.003421129
6
##  [5,] -2.738015e-04  0.0050854014  0.0020868643  0.0034211296  0.015674382
9
##  [6,] -4.863816e-06 -0.0026669657 -0.0015828064 -0.0025553051 -0.012303002
5
##  [7,]  1.198616e-04 -0.0104900199 -0.0004383571 -0.0327622849 -0.003103790
8
##  [8,]  9.761001e-04 -0.0760421212 -0.0019861684 -0.0095689419 -0.003743811
5
##  [9,] -9.513949e-05 -0.0008681135 -0.0024163513 -0.0017912201 -0.003542884
4
## [10,] -2.574126e-05 -0.0049257457 -0.0014905339 -0.0035434188 -0.004493307
8
## [11,] -1.966832e-04  0.0004304433 -0.0063185727  0.0011900806 -0.001517522
2
## [12,] -9.251167e-05 -0.0010833521 -0.0007913929 -0.0012994966 -0.006170715
7
## [13,]  3.580343e-04 -0.0351547697 -0.0009023477 -0.0046212964 -0.001649651
0
## [14,] -2.858465e-05 -0.0051248168 -0.0002297898 -0.0164516073 -0.001382117
2
## [15,] -1.581221e-04  0.0003398944 -0.0007907175 -0.0002333280 -0.000228752
2
## [16,] -1.400277e-04 -0.0001057062 -0.0036849175 -0.0004517136 -0.001210938
9
## [17,] -6.264296e-05 -0.0014575986 -0.0025535864 -0.0017061928 -0.008163383
8
## [18,] -1.040079e-04 -0.0003686854 -0.0055950994 -0.0008607627 -0.002110306
4
##                [,6]          [,7]          [,8]          [,9]         [,10
]
##  [1,] -4.863816e-06  0.0001198616  9.761001e-04 -9.513949e-05 -2.574126e-0
5
##  [2,] -2.666966e-03 -0.0104900199 -7.604212e-02 -8.681135e-04 -4.925746e-0
3
##  [3,] -1.582806e-03 -0.0004383571 -1.986168e-03 -2.416351e-03 -1.490534e-0
3
##  [4,] -2.555305e-03 -0.0327622849 -9.568942e-03 -1.791220e-03 -3.543419e-0
3
##  [5,] -1.230300e-02 -0.0031037908 -3.743812e-03 -3.542884e-03 -4.493308e-0
```

```
3
##  [6,]  1.057474e-02  0.0024790079  2.149139e-03  2.887397e-03  3.694942e-0
3
##  [7,]  2.479008e-03  0.0333937758  8.418767e-03  1.935152e-03  3.137613e-0
3
##  [8,]  2.149139e-03  0.0084187669  7.275023e-02  9.782450e-04  3.805566e-0
3
##  [9,]  2.887397e-03  0.0019351525  9.782450e-04  1.828305e-02  1.384932e-0
3
## [10,]  3.694942e-03  0.0031376131  3.805566e-03  1.384932e-03  1.036489e-0
2
## [11,]  1.292176e-03 -0.0033274506 -6.551623e-04 -5.399951e-03  9.293187e-0
4
## [12,]  5.090968e-03  0.0013585778  9.951562e-04  1.641914e-03  1.923894e-0
3
## [13,]  1.033680e-03  0.0042088532  2.519095e-02  4.766738e-04  1.815985e-0
3
## [14,]  1.203865e-03  0.0133874310  4.241236e-03  8.724652e-04  1.604841e-0
3
## [15,]  3.593683e-04  0.0003329228 -7.811707e-05  9.424408e-04  2.689939e-0
4
## [16,]  1.149161e-03  0.0005321098  2.647847e-04  9.354580e-04  7.356380e-0
4
## [17,]  6.667216e-03  0.0017305341  1.303781e-03  2.238406e-03  2.592881e-0
3
## [18,]  1.832296e-03  0.0009196182  4.815302e-04  1.373959e-03  1.128508e-0
3
##                [,11]         [,12]         [,13]         [,14]         [,15
]
##  [1,] -1.966832e-04 -9.251167e-05  3.580343e-04 -2.858465e-05 -1.581221e-0
4
##  [2,]  4.304433e-04 -1.083352e-03 -3.515477e-02 -5.124817e-03  3.398944e-0
4
##  [3,] -6.318573e-03 -7.913929e-04 -9.023477e-04 -2.297898e-04 -7.907175e-0
4
##  [4,]  1.190081e-03 -1.299497e-03 -4.621296e-03 -1.645161e-02 -2.333280e-0
4
##  [5,] -1.517522e-03 -6.170716e-03 -1.649651e-03 -1.382117e-03 -2.287522e-0
4
##  [6,]  1.292176e-03  5.090968e-03  1.033680e-03  1.203865e-03  3.593683e-0
4
##  [7,] -3.327451e-03  1.358578e-03  4.208853e-03  1.338743e-02  3.329228e-0
4
##  [8,] -6.551623e-04  9.951562e-04  2.519095e-02  4.241236e-03 -7.811707e-0
5
##  [9,] -5.399951e-03  1.641914e-03  4.766738e-04  8.724652e-04  9.424408e-0
4
## [10,]  9.293187e-04  1.923894e-03  1.815985e-03  1.604841e-03  2.689939e-0
4
## [11,]  3.863964e-02  8.578214e-04  9.202374e-05 -1.644750e-04  5.462919e-0
```

```
4
## [12,]  8.578214e-04  5.611085e-03  5.121305e-04  6.936420e-04  2.886982e-0
4
## [13,]  9.202374e-05  5.121305e-04  3.485468e-02  2.156628e-03  4.287971e-0
5
## [14,] -1.644750e-04  6.936420e-04  2.156628e-03  1.043516e-02  2.439338e-0
4
## [15,]  5.462919e-04  2.886982e-04  4.287971e-05  2.439338e-04  7.116111e-0
3
## [16,]  2.067421e-03  6.506733e-04  1.942084e-04  3.536636e-04  3.932445e-0
4
## [17,]  1.836495e-03  3.421402e-03  6.450000e-04  8.740130e-04  3.939479e-0
4
## [18,]  2.976524e-03  9.994005e-04  3.000158e-04  5.434390e-04  5.096368e-0
4
##               [,16]         [,17]         [,18]
##  [1,] -0.0001400277 -6.264296e-05 -0.0001040079
##  [2,] -0.0001057062 -1.457599e-03 -0.0003686854
##  [3,] -0.0036849175 -2.553586e-03 -0.0055950994
##  [4,] -0.0004517136 -1.706193e-03 -0.0008607627
##  [5,] -0.0012109389 -8.163384e-03 -0.0021103064
##  [6,]  0.0011491612  6.667216e-03  0.0018322956
##  [7,]  0.0005321098  1.730534e-03  0.0009196182
##  [8,]  0.0002647847  1.303781e-03  0.0004815302
##  [9,]  0.0009354580  2.238406e-03  0.0013739589
## [10,]  0.0007356380  2.592881e-03  0.0011285080
## [11,]  0.0020674209  1.836495e-03  0.0029765239
## [12,]  0.0006506733  3.421402e-03  0.0009994005
## [13,]  0.0001942084  6.450000e-04  0.0003000158
## [14,]  0.0003536636  8.740130e-04  0.0005434390
## [15,]  0.0003932445  3.939479e-04  0.0005096368
## [16,]  0.0020660902  1.286499e-03  0.0018062727
## [17,]  0.0012864989  1.508833e-02  0.0019533857
## [18,]  0.0018062727  1.953386e-03  0.0047645688

#Model 4: nonparametric - no restriction test
#bw <- npplregbw(formula=CA_Z_adj ~ South_Asian + Amerindian + East_Asian + f
rac_Black_SIRE + frac_EastAsian_SIRE + frac_SouthAsian_SIRE +
#                   frac_Native_American_SIRE + frac_Other_SIRE + frac_Hispani
c_SIRE | African, merged_df)
#summary(bw)


#pl <- npplreg(bws=bw, residuals=TRUE)
#summary(pl)
#coef(pl)
#coef(pl, errors = TRUE)
#summary(pl$resid)
#describe(pl$resid)
```

```
#par(mar = rep(3, 5))
#plot(pl$resid)


#merged_df$CA_Z_adj_hat <- 0.7376529*merged_df$South_Asian - 1.191356*merged_
df$Amerindian + 0.6923586*merged_df$East_Asian -0.1288999*merged_df$frac_Blac
k_SIRE -0.1368878*merged_df$frac_EastAsian_SIRE + 0.1523298*merged_df$frac_So
uthAsian_SIRE -0.2577826*merged_df$frac_Native_American_SIRE -0.07611056*merg
ed_df$frac_Other_SIRE -0.1202465*merged_df$frac_Hispanic_SIRE
#merged_df$CA_Z_adj_sub <- (merged_df$CA_Z_adj - merged_df$CA_Z_adj_hat) -.26
20
#bw2 <- npregbw(formula=merged_df$CA_Z_adj_sub ~ merged_df$African)

# plot density_AA

#plot(dens_African_ancestry005, frame = FALSE, col = "steelblue",
#     main = "",
#     xlab="% African Ancestry",
#     xlim=c(0,1),
#     ylim=c(0,5),
#     xaxs="i",
#     yaxs="i")


#Desnisty Plots_EA
#merged_df_European_ancestry005 = filter(merged_df, European >=.005)
#dens_European_ancestry005 <- density(merged_df_European_ancestry005$European
)

# plot density_EA

#plot(dens_European_ancestry005, frame = FALSE, col = "steelblue",
#     main = "",
#     xlab="% European Ancestry",
#     xlim=c(0,1),
#     ylim=c(0,5),
#     xaxs="i",
#     yaxs="i")

#summary(bw2)
#plot(npreg(bw2),xlim=c(0,1), ylim=c(-1,0), xaxs="i", xlab="Proportion of Afr
ican Ancestry", ylab="Test Scores")
#par(new=TRUE)
# create pairs of data points
#x = c(0,1)
#y =  c(0,-1.00136)
# Create a normal plot
#plot(x, y,type="l", xlim=c(0,1), ylim=c(-1,0),
#     xlab="Proportion of African Ancestry", ylab="Test Scores")
```

```
#Model 5: upper kink, ancestry proportions and individual SIRE categories - n
o restriction test

model_5=lmer(CA_Z_adj ~ South_Asian + Amerindian + East_Asian + African
             + frac_Black_SIRE
             + frac_EastAsian_SIRE
             + frac_SouthAsian_SIRE
             + frac_Native_American_SIRE
             + frac_Other_SIRE
             + frac_Hispanic_SIRE
             + upper_kink
             + (1|site_id_l) + (1|site_id_l:rel_family_id), data=merged_df, R
EML = FALSE)
summary(model_5)
```

```
## Warning in site_id_l:rel_family_id: numerical expression has 9972 elements
: only
## the first used

## Warning in site_id_l:rel_family_id: numerical expression has 9972 elements
: only
## the first used

## Warning in site_id_l:rel_family_id: numerical expression has 9972 elements
: only
## the first used

## Warning in site_id_l:rel_family_id: numerical expression has 9972 elements
: only
## the first used

## Warning in site_id_l:rel_family_id: numerical expression has 9972 elements
: only
## the first used

## Warning in site_id_l:rel_family_id: numerical expression has 9972 elements
: only
## the first used

## Warning in site_id_l:rel_family_id: numerical expression has 9972 elements
: only
## the first used

## Warning in site_id_l:rel_family_id: numerical expression has 9972 elements
: only
## the first used
```

```
## Warning in site_id_l:rel_family_id: numerical expression has 9972 elements
: only
## the first used

## Warning in site_id_l:rel_family_id: numerical expression has 9972 elements
: only
## the first used

## Warning in site_id_l:rel_family_id: numerical expression has 9972 elements
: only
## the first used

## Warning in site_id_l:rel_family_id: numerical expression has 9972 elements
: only
## the first used

## Warning in site_id_l:rel_family_id: numerical expression has 9972 elements
: only
## the first used

## Warning in site_id_l:rel_family_id: numerical expression has 9972 elements
: only
## the first used

## Warning in site_id_l:rel_family_id: numerical expression has 9972 elements
: only
## the first used

## Warning in site_id_l:rel_family_id: numerical expression has 9972 elements
: only
## the first used

## Linear mixed model fit by maximum likelihood  ['lmerMod']
## Formula: CA_Z_adj ~ South_Asian + Amerindian + East_Asian + African +
##     frac_Black_SIRE + frac_EastAsian_SIRE + frac_SouthAsian_SIRE +
##     frac_Native_American_SIRE + frac_Other_SIRE + frac_Hispanic_SIRE +
##     upper_kink + (1 | site_id_l) + (1 | site_id_l:rel_family_id)
##    Data: merged_df
##
##      AIC      BIC   logLik deviance df.resid
##  26072.3  26180.4 -13021.1  26042.3     9957
##
## Scaled residuals:
##     Min      1Q  Median      3Q     Max
## -3.7384 -0.5247 -0.0448  0.4723  4.6302
##
## Random effects:
##  Groups                  Name        Variance Std.Dev.
```

```
##  site_id_l:rel_family_id (Intercept) 0.37322  0.6109
##  site_id_l               (Intercept) 0.01871  0.1368
##  Residual                            0.44965  0.6706
## Number of obs: 9972, groups:  site_id_l:rel_family_id, 8419; site_id_l, 22
##
## Fixed effects:
##                           Estimate Std. Error t value
## (Intercept)                0.30171    0.03311   9.112
## South_Asian                0.47757    0.32637   1.463
## Amerindian                -1.38966    0.12148 -11.439
## East_Asian                 0.63646    0.20217   3.148
## African                   -1.07908    0.12652  -8.529
## frac_Black_SIRE           -0.11323    0.10234  -1.106
## frac_EastAsian_SIRE       -0.20485    0.17581  -1.165
## frac_SouthAsian_SIRE      -0.06677    0.26060  -0.256
## frac_Native_American_SIRE -0.13510    0.10711  -1.261
## frac_Other_SIRE           -0.18816    0.07815  -2.408
## frac_Hispanic_SIRE        -0.08185    0.08451  -0.969
## upper_kink                 0.15979    0.08850   1.805
```

```
performance::icc(model_5)
```

```
## # Intraclass Correlation Coefficient
##
##      Adjusted ICC: 0.466
##   Conditional ICC: 0.393
```

```
performance::r2(model_5)
```

```
## # R2 for Mixed Models
##
##   Conditional R2: 0.550
##      Marginal R2: 0.157
```

```
#Model 6: upper kink, ancestry proportions, individual SIRE categories, 7 mul
ti-SIRE categories with smaller data set - restriction test

model_6=lmer(CA_Z_adj ~ South_Asian + Amerindian + East_Asian + African
             + frac_Black_SIRE_woc
             + frac_EastAsian_SIRE_woc
             + frac_SouthAsian_SIRE_woc
             + frac_Native_American_SIRE_woc
             + frac_Other_Race_SIRE_woc
             + frac_Hispanic_SIRE_woc
             + NH_Black_White_only
             + NH_SouthAsian_White_only
             + NH_EastAsian_White_only
             + NH_Native_American_White_only
             + H_White_only
             + H_Black_only
             + H_Other_only
```

```
             + upper_kink
             + (1|site_id_l) + (1|site_id_l:rel_family_id), data=merged_df_al
l_Model3_spec, REML = FALSE)
summary(model_6)
```

```
## Warning in site_id_l:rel_family_id: numerical expression has 9972 elements
: only
## the first used

## Warning in site_id_l:rel_family_id: numerical expression has 9972 elements
: only
## the first used

## Warning in site_id_l:rel_family_id: numerical expression has 9972 elements
: only
## the first used

## Warning in site_id_l:rel_family_id: numerical expression has 9972 elements
: only
## the first used

## Warning in site_id_l:rel_family_id: numerical expression has 9972 elements
: only
## the first used

## Warning in site_id_l:rel_family_id: numerical expression has 9972 elements
: only
## the first used

## Warning in site_id_l:rel_family_id: numerical expression has 9972 elements
: only
## the first used

## Warning in site_id_l:rel_family_id: numerical expression has 9972 elements
: only
## the first used

## Warning in site_id_l:rel_family_id: numerical expression has 9972 elements
: only
## the first used

## Warning in site_id_l:rel_family_id: numerical expression has 9972 elements
: only
## the first used

## Warning in site_id_l:rel_family_id: numerical expression has 9972 elements
: only
## the first used
```

```
## Warning in site_id_l:rel_family_id: numerical expression has 9972 elements
: only
## the first used

## Warning in site_id_l:rel_family_id: numerical expression has 9972 elements
: only
## the first used

## Warning in site_id_l:rel_family_id: numerical expression has 9972 elements
: only
## the first used

## Warning in site_id_l:rel_family_id: numerical expression has 9972 elements
: only
## the first used

## Warning in site_id_l:rel_family_id: numerical expression has 9972 elements
: only
## the first used

## Linear mixed model fit by maximum likelihood  ['lmerMod']
## Formula: CA_Z_adj ~ South_Asian + Amerindian + East_Asian + African +
##     frac_Black_SIRE_woc + frac_EastAsian_SIRE_woc + frac_SouthAsian_SIRE_w
oc +
##     frac_Native_American_SIRE_woc + frac_Other_Race_SIRE_woc +
##     frac_Hispanic_SIRE_woc + NH_Black_White_only + NH_SouthAsian_White_onl
y +
##     NH_EastAsian_White_only + NH_Native_American_White_only +
##     H_White_only + H_Black_only + H_Other_only + upper_kink +
##     (1 | site_id_l) + (1 | site_id_l:rel_family_id)
##    Data: merged_df_all_Model3_spec
##
##      AIC      BIC   logLik deviance df.resid
##  26058.3  26216.9 -13007.2  26014.3     9950
##
## Scaled residuals:
##     Min      1Q  Median      3Q     Max
## -3.7255 -0.5242 -0.0438  0.4725  4.6693
##
## Random effects:
##  Groups                  Name        Variance Std.Dev.
##  site_id_l:rel_family_id (Intercept) 0.37031  0.6085
##  site_id_l               (Intercept) 0.01826  0.1351
##  Residual                            0.44998  0.6708
## Number of obs: 9972, groups:  site_id_l:rel_family_id, 8419; site_id_l, 22
##
## Fixed effects:
##                                 Estimate Std. Error t value
## (Intercept)                     0.295577   0.032884   8.989
## South_Asian                     0.531618   0.327789   1.622
```

```
## Amerindian                    -1.343729   0.123909 -10.844
## East_Asian                     0.650113   0.202978   3.203
## African                       -1.094851   0.129180  -8.475
## frac_Black_SIRE_woc           -0.116278   0.104162  -1.116
## frac_EastAsian_SIRE_woc       -0.341833   0.182770  -1.870
## frac_SouthAsian_SIRE_woc      -0.258389   0.269696  -0.958
## frac_Native_American_SIRE_woc -0.171245   0.135368  -1.265
## frac_Other_Race_SIRE_woc      -0.165017   0.102038  -1.617
## frac_Hispanic_SIRE_woc         0.350075   0.196595   1.781
## NH_Black_White_only            0.078160   0.075878   1.030
## NH_SouthAsian_White_only       0.340133   0.186653   1.822
## NH_EastAsian_White_only        0.008492   0.102148   0.083
## NH_Native_American_White_only -0.099662   0.084335  -1.182
## H_White_only                  -0.074910   0.045549  -1.645
## H_Black_only                   0.070014   0.123923   0.565
## H_Other_only                  -0.156631   0.069150  -2.265
## upper_kink                     0.180855   0.088764   2.037
```

```
##
## Correlation matrix not shown by default, as p = 19 > 12.
## Use print(x, correlation=TRUE)  or
##     vcov(x)        if you need it
```

```
performance::icc(model_6)
```

```
## # Intraclass Correlation Coefficient
##
##      Adjusted ICC: 0.463
##   Conditional ICC: 0.389
```

```
performance::r2(model_6)
```

```
## # R2 for Mixed Models
##
##   Conditional R2: 0.549
##      Marginal R2: 0.160
```

```
#For Restriction Test
# the test applies to models 2, 3, 6, 8a and 8b

coeffvec <- coef(model_6)
varcov <- vcov(model_6, full=FALSE)
```

```
## Warning in site_id_l:rel_family_id: numerical expression has 9972 elements
: only
## the first used

## Warning in site_id_l:rel_family_id: numerical expression has 9972 elements
: only
## the first used
```

```
## Warning in site_id_l:rel_family_id: numerical expression has 9972 elements
: only
## the first used

## Warning in site_id_l:rel_family_id: numerical expression has 9972 elements
: only
## the first used

## Warning in site_id_l:rel_family_id: numerical expression has 9972 elements
: only
## the first used

## Warning in site_id_l:rel_family_id: numerical expression has 9972 elements
: only
## the first used

## Warning in site_id_l:rel_family_id: numerical expression has 9972 elements
: only
## the first used

## Warning in site_id_l:rel_family_id: numerical expression has 9972 elements
: only
## the first used
```

```
varcov
```

```
## 19 x 19 Matrix of class "dgeMatrix"
##                [,1]          [,2]          [,3]          [,4]          [,5
]
##  [1,]  1.081342e-03 -0.0016929455 -0.0003260876 -0.0004176675 -0.000285445
1
##  [2,] -1.692945e-03  0.1074454880  0.0029781999  0.0122896390  0.005282189
0
##  [3,] -3.260876e-04  0.0029781999  0.0153534935  0.0010827362  0.002213107
2
##  [4,] -4.176675e-04  0.0122896390  0.0010827362  0.0412001946  0.003580329
1
##  [5,] -2.854451e-04  0.0052821890  0.0022131072  0.0035803291  0.016687356
6
##  [6,]  1.332020e-06 -0.0027702367 -0.0016487579 -0.0026384459 -0.012831875
0
##  [7,]  1.216425e-04 -0.0105169277 -0.0004585429 -0.0327730674 -0.003266530
7
##  [8,]  9.772374e-04 -0.0760326435 -0.0020037457 -0.0095878872 -0.003893309
4
##  [9,] -9.249743e-05 -0.0009115948 -0.0024432081 -0.0018256879 -0.003767293
2
## [10,] -2.309961e-05 -0.0049680873 -0.0015183787 -0.0035779618 -0.004720129
8
## [11,] -1.945298e-04  0.0003957984 -0.0063368503  0.0011618958 -0.001689876
```

```
6
## [12,] -8.792783e-05 -0.0011591314 -0.0008396583 -0.0013603462 -0.006558048
8
## [13,]  3.585559e-04 -0.0351491082 -0.0009097298 -0.0046290394 -0.001712832
1
## [14,] -2.782752e-05 -0.0051351842 -0.0002379140 -0.0164543509 -0.001447389
8
## [15,] -1.579883e-04  0.0003387854 -0.0007909811 -0.0002339436 -0.000233666
2
## [16,] -1.387457e-04 -0.0001255133 -0.0036954849 -0.0004673686 -0.001310805
3
## [17,] -5.643086e-05 -0.0015610100 -0.0026184519 -0.0017891485 -0.008690488
8
## [18,] -1.022532e-04 -0.0003965641 -0.0056101283 -0.0008828274 -0.002251009
4
## [19,]  3.298522e-05 -0.0005543034 -0.0003527612 -0.0004462628 -0.002835713
4
##                [,6]          [,7]          [,8]          [,9]         [,10
]
##  [1,]  1.332020e-06  0.0001216425  9.772374e-04 -9.249743e-05 -2.309961e-0
5
##  [2,] -2.770237e-03 -0.0105169277 -7.603264e-02 -9.115948e-04 -4.968087e-0
3
##  [3,] -1.648758e-03 -0.0004585429 -2.003746e-03 -2.443208e-03 -1.518379e-0
3
##  [4,] -2.638446e-03 -0.0327730674 -9.587887e-03 -1.825688e-03 -3.577962e-0
3
##  [5,] -1.283188e-02 -0.0032665307 -3.893309e-03 -3.767293e-03 -4.720130e-0
3
##  [6,]  1.084981e-02  0.0025639424  2.227530e-03  3.004474e-03  3.813217e-0
3
##  [7,]  2.563942e-03  0.0334048971  8.439059e-03  1.970355e-03  3.173100e-0
3
##  [8,]  2.227530e-03  0.0084390589  7.273592e-02  1.011131e-03  3.837635e-0
3
##  [9,]  3.004474e-03  0.0019703553  1.011131e-03  1.832437e-02  1.434942e-0
3
## [10,]  3.813217e-03  0.0031731002  3.837635e-03  1.434942e-03  1.041170e-0
2
## [11,]  1.382273e-03 -0.0032977197 -6.288440e-04 -5.359291e-03  9.677262e-0
4
## [12,]  5.292981e-03  0.0014206837  1.052605e-03  1.727545e-03  2.010529e-0
3
## [13,]  1.066747e-03  0.0042170640  2.518740e-02  4.905243e-04  1.829467e-0
3
## [14,]  1.237861e-03  0.0133919515  4.248969e-03  8.865648e-04  1.618933e-0
3
## [15,]  3.618033e-04  0.0003335259 -7.735361e-05  9.425424e-04  2.699948e-0
4
## [16,]  1.201253e-03  0.0005480188  2.796852e-04  9.572301e-04  7.578196e-0
```

```
4
## [17,]  6.942231e-03  0.0018151881  1.382155e-03  2.354972e-03  2.710772e-0
3
## [18,]  1.905683e-03  0.0009419915  5.025055e-04  1.404710e-03  1.159776e-0
3
## [19,]  1.485558e-03  0.0004557376  4.206799e-04  6.279305e-04  6.352738e-0
4
##               [,11]         [,12]         [,13]         [,14]         [,15
]
##  [1,] -0.0001945298 -8.792783e-05  3.585559e-04 -2.782752e-05 -1.579883e-0
4
##  [2,]  0.0003957984 -1.159131e-03 -3.514911e-02 -5.135184e-03  3.387854e-0
4
##  [3,] -0.0063368503 -8.396583e-04 -9.097298e-04 -2.379140e-04 -7.909811e-0
4
##  [4,]  0.0011618958 -1.360346e-03 -4.629039e-03 -1.645435e-02 -2.339436e-0
4
##  [5,] -0.0016898766 -6.558049e-03 -1.712832e-03 -1.447390e-03 -2.336662e-0
4
##  [6,]  0.0013822729  5.292981e-03  1.066747e-03  1.237861e-03  3.618033e-0
4
##  [7,] -0.0032977197  1.420684e-03  4.217064e-03  1.339195e-02  3.335259e-0
4
##  [8,] -0.0006288440  1.052605e-03  2.518740e-02  4.248969e-03 -7.735361e-0
5
##  [9,] -0.0053592913  1.727545e-03  4.905243e-04  8.865648e-04  9.425424e-0
4
## [10,]  0.0009677262  2.010529e-03  1.829467e-03  1.618933e-03  2.699948e-0
4
## [11,]  0.0386494136  9.235635e-04  1.029876e-04 -1.532161e-04  5.469727e-0
4
## [12,]  0.0009235635  5.757454e-03  5.363769e-04  7.184990e-04  2.904552e-0
4
## [13,]  0.0001029876  5.363769e-04  3.483925e-02  2.159684e-03  4.316409e-0
5
## [14,] -0.0001532161  7.184990e-04  2.159684e-03  1.043428e-02  2.441201e-0
4
## [15,]  0.0005469727  2.904552e-04  4.316409e-05  2.441201e-04  7.112422e-0
3
## [16,]  0.0020831239  6.887562e-04  2.004803e-04  3.599695e-04  3.935318e-0
4
## [17,]  0.0019254287  3.622805e-03  6.781080e-04  9.078983e-04  3.963515e-0
4
## [18,]  0.0029988406  1.053102e-03  3.088375e-04  5.523224e-04  5.100844e-0
4
## [19,]  0.0004806715  1.084358e-03  1.779972e-04  1.829171e-04  1.405558e-0
5
##               [,16]         [,17]         [,18]         [,19]
##  [1,] -0.0001387457 -5.643086e-05 -0.0001022532  3.298522e-05
##  [2,] -0.0001255133 -1.561010e-03 -0.0003965641 -5.543034e-04
```

```
##  [3,] -0.0036954849 -2.618452e-03 -0.0056101283 -3.527612e-04
##  [4,] -0.0004673686 -1.789148e-03 -0.0008828274 -4.462628e-04
##  [5,] -0.0013108053 -8.690489e-03 -0.0022510094 -2.835713e-03
##  [6,]  0.0012012531  6.942231e-03  0.0019056827  1.485558e-03
##  [7,]  0.0005480188  1.815188e-03  0.0009419915  4.557376e-04
##  [8,]  0.0002796852  1.382155e-03  0.0005025055  4.206799e-04
##  [9,]  0.0009572301  2.354972e-03  0.0014047102  6.279305e-04
## [10,]  0.0007578196  2.710772e-03  0.0011597760  6.352738e-04
## [11,]  0.0020831239  1.925429e-03  0.0029988406  4.806715e-04
## [12,]  0.0006887562  3.622805e-03  0.0010531017  1.084358e-03
## [13,]  0.0002004803  6.781080e-04  0.0003088375  1.779972e-04
## [14,]  0.0003599695  9.078983e-04  0.0005523224  1.829171e-04
## [15,]  0.0003935318  3.963515e-04  0.0005100844  1.405558e-05
## [16,]  0.0020746800  1.337971e-03  0.0018191412  2.783945e-04
## [17,]  0.0013379707  1.535700e-02  0.0020260093  1.474992e-03
## [18,]  0.0018191412  2.026009e-03  0.0047817747  3.932758e-04
## [19,]  0.0002783945  1.474992e-03  0.0003932758  7.879004e-03

#Model 7a: upper kink, raw SES, US child, ancestry proportions and individual
SIRE categories - no restriction test


model_7a=lmer(CA_Z_adj ~ South_Asian + Amerindian + East_Asian + African
              + frac_Black_SIRE
              + frac_EastAsian_SIRE
              + frac_SouthAsian_SIRE
              + frac_Native_American_SIRE
              + frac_Other_SIRE
              + frac_Hispanic_SIRE
              + upper_kink
              + Child_US_Born
              + general_ses_PCA_z
              + (1|site_id_l) + (1|site_id_l:rel_family_id), data=merged_df,
REML = FALSE)
summary(model_7a)

## Warning in site_id_l:rel_family_id: numerical expression has 9972 elements
: only
## the first used

## Warning in site_id_l:rel_family_id: numerical expression has 9972 elements
: only
## the first used

## Warning in site_id_l:rel_family_id: numerical expression has 9972 elements
: only
## the first used

## Warning in site_id_l:rel_family_id: numerical expression has 9972 elements
: only
```

```
## the first used

## Warning in site_id_l:rel_family_id: numerical expression has 9972 elements
: only
## the first used

## Warning in site_id_l:rel_family_id: numerical expression has 9972 elements
: only
## the first used

## Warning in site_id_l:rel_family_id: numerical expression has 9972 elements
: only
## the first used

## Warning in site_id_l:rel_family_id: numerical expression has 9972 elements
: only
## the first used

## Warning in site_id_l:rel_family_id: numerical expression has 9972 elements
: only
## the first used

## Warning in site_id_l:rel_family_id: numerical expression has 9972 elements
: only
## the first used

## Warning in site_id_l:rel_family_id: numerical expression has 9972 elements
: only
## the first used

## Warning in site_id_l:rel_family_id: numerical expression has 9972 elements
: only
## the first used

## Warning in site_id_l:rel_family_id: numerical expression has 9972 elements
: only
## the first used

## Warning in site_id_l:rel_family_id: numerical expression has 9972 elements
: only
## the first used

## Warning in site_id_l:rel_family_id: numerical expression has 9972 elements
: only
## the first used

## Warning in site_id_l:rel_family_id: numerical expression has 9972 elements
: only
## the first used
```

```
## Linear mixed model fit by maximum likelihood  ['lmerMod']
## Formula: CA_Z_adj ~ South_Asian + Amerindian + East_Asian + African +
##     frac_Black_SIRE + frac_EastAsian_SIRE + frac_SouthAsian_SIRE +
##     frac_Native_American_SIRE + frac_Other_SIRE + frac_Hispanic_SIRE +
##     upper_kink + Child_US_Born + general_ses_PCA_z + (1 | site_id_l) +
##     (1 | site_id_l:rel_family_id)
##    Data: merged_df
##
##      AIC      BIC   logLik deviance df.resid
##  25536.1  25658.6 -12751.0  25502.1     9955
##
## Scaled residuals:
##     Min      1Q  Median      3Q     Max
## -3.9197 -0.5388 -0.0475  0.4950  4.9804
##
## Random effects:
##  Groups                  Name        Variance Std.Dev.
##  site_id_l:rel_family_id (Intercept) 0.321000 0.56657
##  site_id_l               (Intercept) 0.008193 0.09052
##  Residual                            0.454391 0.67409
## Number of obs: 9972, groups:  site_id_l:rel_family_id, 8419; site_id_l, 22
##
## Fixed effects:
##                            Estimate Std. Error t value
## (Intercept)                 0.07991    0.06179   1.293
## South_Asian                 0.42998    0.31742   1.355
## Amerindian                 -0.82075    0.11946  -6.870
## East_Asian                  0.62222    0.19635   3.169
## African                    -0.62986    0.12415  -5.073
## frac_Black_SIRE            -0.09594    0.09930  -0.966
## frac_EastAsian_SIRE        -0.21786    0.17079  -1.276
## frac_SouthAsian_SIRE       -0.11397    0.25288  -0.451
## frac_Native_American_SIRE -0.03397    0.10374  -0.327
## frac_Other_SIRE            -0.07625    0.07611  -1.002
## frac_Hispanic_SIRE         -0.03131    0.08167  -0.383
## upper_kink                  0.02086    0.08612   0.242
## Child_US_Born               0.10053    0.05697   1.765
## general_ses_PCA_z           0.27999    0.01180  23.728

##
## Correlation matrix not shown by default, as p = 14 > 12.
## Use print(x, correlation=TRUE)  or
##     vcov(x)        if you need it

performance::icc(model_7a)

## # Intraclass Correlation Coefficient
##
##      Adjusted ICC: 0.420
##   Conditional ICC: 0.329
```

```
performance::r2(model_7a)

## # R2 for Mixed Models
##
##   Conditional R2: 0.546
##      Marginal R2: 0.218

#Model 7b: upper kink, orthogonalized SES, US child, ancestry proportions and
individual SIRE categories - no restriction test
# Model 7b same as model 7a except using orthogonalized SES in place of raw S
ES
# using the LM command to get the projection of SES on the other variables (a
ll except Child_US_born)

orthostep = lm(general_ses_PCA_z ~ South_Asian + Amerindian + East_Asian + Af
rican
               + frac_Black_SIRE
               + frac_EastAsian_SIRE
               + frac_SouthAsian_SIRE
               + frac_Native_American_SIRE
               + frac_Other_SIRE
               + frac_Hispanic_SIRE
               + upper_kink, data=merged_df)
ortho_ses <- residuals(orthostep)


model_7b=lmer(CA_Z_adj ~ South_Asian + Amerindian + East_Asian + African
              + frac_Black_SIRE
              + frac_EastAsian_SIRE
              + frac_SouthAsian_SIRE
              + frac_Native_American_SIRE
              + frac_Other_SIRE
              + frac_Hispanic_SIRE
              + upper_kink
              + Child_US_Born
              + ortho_ses
              + (1|site_id_l) + (1|site_id_l:rel_family_id), data=merged_df,
REML = FALSE)
summary(model_7b)

## Warning in site_id_l:rel_family_id: numerical expression has 9972 elements
: only
## the first used

## Warning in site_id_l:rel_family_id: numerical expression has 9972 elements
: only
## the first used

## Warning in site_id_l:rel_family_id: numerical expression has 9972 elements
: only
```

```
## the first used

## Warning in site_id_l:rel_family_id: numerical expression has 9972 elements
: only
## the first used

## Warning in site_id_l:rel_family_id: numerical expression has 9972 elements
: only
## the first used

## Warning in site_id_l:rel_family_id: numerical expression has 9972 elements
: only
## the first used

## Warning in site_id_l:rel_family_id: numerical expression has 9972 elements
: only
## the first used

## Warning in site_id_l:rel_family_id: numerical expression has 9972 elements
: only
## the first used

## Warning in site_id_l:rel_family_id: numerical expression has 9972 elements
: only
## the first used

## Warning in site_id_l:rel_family_id: numerical expression has 9972 elements
: only
## the first used

## Warning in site_id_l:rel_family_id: numerical expression has 9972 elements
: only
## the first used

## Warning in site_id_l:rel_family_id: numerical expression has 9972 elements
: only
## the first used

## Warning in site_id_l:rel_family_id: numerical expression has 9972 elements
: only
## the first used

## Warning in site_id_l:rel_family_id: numerical expression has 9972 elements
: only
## the first used

## Warning in site_id_l:rel_family_id: numerical expression has 9972 elements
: only
## the first used
```

```
## Warning in site_id_l:rel_family_id: numerical expression has 9972 elements
: only
## the first used

## Linear mixed model fit by maximum likelihood  ['lmerMod']
## Formula: CA_Z_adj ~ South_Asian + Amerindian + East_Asian + African +
##     frac_Black_SIRE + frac_EastAsian_SIRE + frac_SouthAsian_SIRE +
##     frac_Native_American_SIRE + frac_Other_SIRE + frac_Hispanic_SIRE +
##     upper_kink + Child_US_Born + ortho_ses + (1 | site_id_l) +
##     (1 | site_id_l:rel_family_id)
##    Data: merged_df
##
##      AIC      BIC   logLik deviance df.resid
##  25536.1  25658.6 -12751.0  25502.1     9955
##
## Scaled residuals:
##     Min      1Q  Median      3Q     Max
## -3.9197 -0.5388 -0.0475  0.4950  4.9804
##
## Random effects:
##  Groups                  Name        Variance Std.Dev.
##  site_id_l:rel_family_id (Intercept) 0.321000 0.56657
##  site_id_l               (Intercept) 0.008193 0.09052
##  Residual                            0.454391 0.67409
## Number of obs: 9972, groups:  site_id_l:rel_family_id, 8419; site_id_l, 22
##
## Fixed effects:
##                            Estimate Std. Error t value
## (Intercept)                 0.19868    0.06149   3.231
## South_Asian                 0.62293    0.31740   1.963
## Amerindian                 -1.35902    0.11720 -11.596
## East_Asian                  0.71815    0.19635   3.657
## African                    -1.11791    0.12268  -9.113
## frac_Black_SIRE            -0.11908    0.09930  -1.199
## frac_EastAsian_SIRE        -0.21402    0.17079  -1.253
## frac_SouthAsian_SIRE       -0.11798    0.25289  -0.467
## frac_Native_American_SIRE -0.24121    0.10372  -2.326
## frac_Other_SIRE            -0.17395    0.07598  -2.290
## frac_Hispanic_SIRE         -0.08514    0.08165  -1.043
## upper_kink                  0.20551    0.08592   2.392
## Child_US_Born               0.10053    0.05697   1.765
## ortho_ses                   0.27999    0.01180  23.728

##
## Correlation matrix not shown by default, as p = 14 > 12.
## Use print(x, correlation=TRUE)  or
##     vcov(x)        if you need it

performance::icc(model_7b)
```

```
## # Intraclass Correlation Coefficient
##
##      Adjusted ICC: 0.420
##   Conditional ICC: 0.329
```

```
performance::r2(model_7b)
```

```
## # R2 for Mixed Models
##
##   Conditional R2: 0.546
##      Marginal R2: 0.218
```

```
#Model 8a: upper kink, raw SES, US child, ancestry proportions, individual SI
RE categories, and 7 multi-SIRE categories with smaller data set - restrictio
n test
# same steps for models 8 which splits into models 8a and 8b using raw vs. or
thogonalized SES
# model 8a (already run but included for completeness)

model_8a=lmer(CA_Z_adj ~ South_Asian + Amerindian + East_Asian + African
              + frac_Black_SIRE_woc
              + frac_EastAsian_SIRE_woc
              + frac_SouthAsian_SIRE_woc
              + frac_Native_American_SIRE_woc
              + frac_Other_Race_SIRE_woc
              + frac_Hispanic_SIRE_woc
              + NH_Black_White_only
              + NH_SouthAsian_White_only
              + NH_EastAsian_White_only
              + NH_Native_American_White_only
              + H_White_only
              + H_Black_only
              + H_Other_only
              + upper_kink
              + Child_US_Born
              + general_ses_PCA_z
              + (1|site_id_l) + (1|site_id_l:rel_family_id), data=merged_df_a
ll_Model3_spec, REML = FALSE)
summary(model_8a)
```

```
## Warning in site_id_l:rel_family_id: numerical expression has 9972 elements
: only
## the first used

## Warning in site_id_l:rel_family_id: numerical expression has 9972 elements
: only
## the first used

## Warning in site_id_l:rel_family_id: numerical expression has 9972 elements
: only
```

```
## the first used

## Warning in site_id_l:rel_family_id: numerical expression has 9972 elements
: only
## the first used

## Warning in site_id_l:rel_family_id: numerical expression has 9972 elements
: only
## the first used

## Warning in site_id_l:rel_family_id: numerical expression has 9972 elements
: only
## the first used

## Warning in site_id_l:rel_family_id: numerical expression has 9972 elements
: only
## the first used

## Warning in site_id_l:rel_family_id: numerical expression has 9972 elements
: only
## the first used

## Warning in site_id_l:rel_family_id: numerical expression has 9972 elements
: only
## the first used

## Warning in site_id_l:rel_family_id: numerical expression has 9972 elements
: only
## the first used

## Warning in site_id_l:rel_family_id: numerical expression has 9972 elements
: only
## the first used

## Warning in site_id_l:rel_family_id: numerical expression has 9972 elements
: only
## the first used

## Warning in site_id_l:rel_family_id: numerical expression has 9972 elements
: only
## the first used

## Warning in site_id_l:rel_family_id: numerical expression has 9972 elements
: only
## the first used

## Warning in site_id_l:rel_family_id: numerical expression has 9972 elements
: only
## the first used
```

```
## Warning in site_id_l:rel_family_id: numerical expression has 9972 elements
: only
## the first used

## Linear mixed model fit by maximum likelihood  ['lmerMod']
## Formula: CA_Z_adj ~ South_Asian + Amerindian + East_Asian + African +
##     frac_Black_SIRE_woc + frac_EastAsian_SIRE_woc + frac_SouthAsian_SIRE_w
oc +
##     frac_Native_American_SIRE_woc + frac_Other_Race_SIRE_woc +
##     frac_Hispanic_SIRE_woc + NH_Black_White_only + NH_SouthAsian_White_onl
y +
##     NH_EastAsian_White_only + NH_Native_American_White_only +
##     H_White_only + H_Black_only + H_Other_only + upper_kink +
##     Child_US_Born + general_ses_PCA_z + (1 | site_id_l) + (1 |
##     site_id_l:rel_family_id)
##    Data: merged_df_all_Model3_spec
##
##      AIC      BIC   logLik deviance df.resid
##  25523.4  25696.4 -12737.7  25475.4     9948
##
## Scaled residuals:
##     Min      1Q  Median      3Q     Max
## -3.9088 -0.5421 -0.0481  0.4970  5.0183
##
## Random effects:
##  Groups                  Name        Variance Std.Dev.
##  site_id_l:rel_family_id (Intercept) 0.318111 0.56401
##  site_id_l               (Intercept) 0.008041 0.08967
##  Residual                            0.454931 0.67449
## Number of obs: 9972, groups:  site_id_l:rel_family_id, 8419; site_id_l, 22
##
## Fixed effects:
##                                Estimate Std. Error t value
## (Intercept)                     0.08464    0.06173   1.371
## South_Asian                     0.47092    0.31872   1.478
## Amerindian                     -0.78284    0.12175  -6.430
## East_Asian                      0.63061    0.19711   3.199
## African                        -0.65025    0.12672  -5.132
## frac_Black_SIRE_woc            -0.09775    0.10111  -0.967
## frac_EastAsian_SIRE_woc        -0.31764    0.17758  -1.789
## frac_SouthAsian_SIRE_woc       -0.28412    0.26170  -1.086
## frac_Native_American_SIRE_woc -0.05166    0.13147  -0.393
## frac_Other_Race_SIRE_woc       -0.04434    0.09947  -0.446
## frac_Hispanic_SIRE_woc          0.37824    0.19062   1.984
## NH_Black_White_only             0.11856    0.07357   1.611
## NH_SouthAsian_White_only        0.28393    0.18080   1.570
## NH_EastAsian_White_only        -0.03175    0.09917  -0.320
## NH_Native_American_White_only -0.05843    0.08171  -0.715
## H_White_only                   -0.04267    0.04400  -0.970
```

```
## H_Black_only                     0.06452    0.12024   0.537
## H_Other_only                    -0.07644    0.06717  -1.138
## upper_kink                       0.04466    0.08638   0.517
## Child_US_Born                    0.08967    0.05700   1.573
## general_ses_PCA_z                0.27956    0.01179  23.710
```

```
##
## Correlation matrix not shown by default, as p = 21 > 12.
## Use print(x, correlation=TRUE)  or
##     vcov(x)        if you need it
```

```
performance::icc(model_8a)
```

```
## # Intraclass Correlation Coefficient
##
##      Adjusted ICC: 0.418
##   Conditional ICC: 0.326
```

```
performance::r2(model_8a)
```

```
## # R2 for Mixed Models
##
##   Conditional R2: 0.546
##      Marginal R2: 0.220
```

```
#For Restriction Test
# the test applies to models 2, 3, 6, 8a and 8b

coeffvec <- coef(model_8a)
varcov <- vcov(model_8a, full=FALSE)
```

```
## Warning in site_id_l:rel_family_id: numerical expression has 9972 elements
: only
## the first used

## Warning in site_id_l:rel_family_id: numerical expression has 9972 elements
: only
## the first used

## Warning in site_id_l:rel_family_id: numerical expression has 9972 elements
: only
## the first used

## Warning in site_id_l:rel_family_id: numerical expression has 9972 elements
: only
## the first used

## Warning in site_id_l:rel_family_id: numerical expression has 9972 elements
: only
## the first used
```

```
## Warning in site_id_l:rel_family_id: numerical expression has 9972 elements
: only
## the first used

## Warning in site_id_l:rel_family_id: numerical expression has 9972 elements
: only
## the first used

## Warning in site_id_l:rel_family_id: numerical expression has 9972 elements
: only
## the first used
```

```
varcov
```

```
## 21 x 21 Matrix of class "dgeMatrix"
##                [,1]          [,2]          [,3]          [,4]          [,5
]
##  [1,]  3.810613e-03 -3.028438e-03 -5.050505e-04 -8.132799e-04 -3.003166e-0
4
##  [2,] -3.028438e-03  1.015837e-01  2.660925e-03  1.170635e-02  4.801454e-0
3
##  [3,] -5.050505e-04  2.660925e-03  1.482285e-02  9.287695e-04  2.542400e-0
3
##  [4,] -8.132799e-04  1.170635e-02  9.287695e-04  3.885379e-02  3.339094e-0
3
##  [5,] -3.003166e-04  4.801454e-03  2.542400e-03  3.339094e-03  1.605685e-0
2
##  [6,] -8.274242e-05 -2.574710e-03 -1.550110e-03 -2.492550e-03 -1.207505e-0
2
##  [7,]  3.923823e-06 -9.850360e-03 -4.078969e-04 -3.091528e-02 -3.074560e-0
3
##  [8,]  1.497433e-03 -7.169775e-02 -1.871128e-03 -9.083546e-03 -3.633797e-0
3
##  [9,] -1.367436e-04 -8.537170e-04 -2.169991e-03 -1.707128e-03 -3.454007e-0
3
## [10,] -2.927433e-04 -4.594624e-03 -1.321775e-03 -3.366236e-03 -4.381957e-0
3
## [11,] -2.615675e-04  3.445999e-04 -5.931145e-03  1.070336e-03 -1.585242e-0
3
## [12,] -1.045448e-04 -1.094222e-03 -7.611873e-04 -1.286907e-03 -6.144770e-0
3
## [13,]  8.271006e-04 -3.322530e-02 -8.939964e-04 -4.412793e-03 -1.623529e-0
3
## [14,]  1.595409e-04 -4.909056e-03 -2.671318e-04 -1.554601e-02 -1.394785e-0
3
## [15,] -1.239069e-04  2.951593e-04 -7.020244e-04 -2.223535e-04 -1.782817e-0
4
## [16,] -2.534889e-04 -9.466475e-05 -3.438503e-03 -4.355404e-04 -1.237594e-0
3
## [17,] -2.107689e-04 -1.420793e-03 -2.482619e-03 -1.680641e-03 -8.212494e-0
```

```
3
## [18,] -2.854221e-04 -3.443819e-04 -5.221790e-03 -8.321198e-04 -2.080635e-0
3
## [19,] -1.044473e-04 -4.038132e-04 -4.690650e-04 -3.838164e-04 -2.783608e-0
3
## [20,] -3.224048e-03  1.519110e-03  9.968029e-05  4.506410e-04 -5.254879e-0
5
## [21,] -7.437288e-05 -6.306044e-05  2.688630e-04 -2.318242e-05  2.229803e-0
4
##                [,6]          [,7]          [,8]          [,9]         [,10
]
##  [1,] -8.274242e-05  3.923823e-06  1.497433e-03 -1.367436e-04 -2.927433e-0
4
##  [2,] -2.574710e-03 -9.850360e-03 -7.169775e-02 -8.537170e-04 -4.594624e-0
3
##  [3,] -1.550110e-03 -4.078969e-04 -1.871128e-03 -2.169991e-03 -1.321775e-0
3
##  [4,] -2.492550e-03 -3.091528e-02 -9.083546e-03 -1.707128e-03 -3.366236e-0
3
##  [5,] -1.207505e-02 -3.074560e-03 -3.633797e-03 -3.454007e-03 -4.381957e-0
3
##  [6,]  1.022361e-02  2.431929e-03  2.088053e-03  2.839258e-03  3.621325e-0
3
##  [7,]  2.431929e-03  3.153547e-02  7.927243e-03  1.845409e-03  3.019124e-0
3
##  [8,]  2.088053e-03  7.927243e-03  6.848801e-02  9.323143e-04  3.569012e-0
3
##  [9,]  2.839258e-03  1.845409e-03  9.323143e-04  1.728500e-02  1.382902e-0
3
## [10,]  3.621325e-03  3.019124e-03  3.569012e-03  1.382902e-03  9.893901e-0
3
## [11,]  1.314366e-03 -3.077850e-03 -5.748590e-04 -5.060809e-03  9.301960e-0
4
## [12,]  4.986384e-03  1.342261e-03  9.901618e-04  1.635766e-03  1.912655e-0
3
## [13,]  9.892599e-04  3.948519e-03  2.373882e-02  4.396407e-04  1.677863e-0
3
## [14,]  1.166233e-03  1.263783e-02  4.034418e-03  8.206871e-04  1.512399e-0
3
## [15,]  3.379193e-04  3.098053e-04 -6.867245e-05  8.471200e-04  2.610141e-0
4
## [16,]  1.150196e-03  5.267812e-04  2.571593e-04  9.127086e-04  7.366314e-0
4
## [17,]  6.551513e-03  1.727556e-03  1.292867e-03  2.223744e-03  2.578940e-0
3
## [18,]  1.812416e-03  9.014318e-04  4.607795e-04  1.347702e-03  1.129136e-0
3
## [19,]  1.397494e-03  4.289011e-04  3.637195e-04  5.670579e-04  5.839357e-0
4
## [20,]  7.781056e-05  1.018102e-04 -5.978332e-04  2.016807e-05  2.465960e-0
```

```
4
## [21,]  1.070702e-05  1.114521e-05  2.948499e-07  6.391754e-05  5.886088e-0
5
##               [,11]         [,12]         [,13]         [,14]         [,15
]
##  [1,] -2.615675e-04 -1.045448e-04  8.271006e-04  1.595409e-04 -1.239069e-0
4
##  [2,]  3.445999e-04 -1.094222e-03 -3.322530e-02 -4.909056e-03  2.951593e-0
4
##  [3,] -5.931145e-03 -7.611873e-04 -8.939964e-04 -2.671318e-04 -7.020244e-0
4
##  [4,]  1.070336e-03 -1.286907e-03 -4.412793e-03 -1.554601e-02 -2.223535e-0
4
##  [5,] -1.585242e-03 -6.144770e-03 -1.623529e-03 -1.394785e-03 -1.782817e-0
4
##  [6,]  1.314366e-03  4.986384e-03  9.892599e-04  1.166233e-03  3.379193e-0
4
##  [7,] -3.077850e-03  1.342261e-03  3.948519e-03  1.263783e-02  3.098053e-0
4
##  [8,] -5.748590e-04  9.901618e-04  2.373882e-02  4.034418e-03 -6.867245e-0
5
##  [9,] -5.060809e-03  1.635766e-03  4.396407e-04  8.206871e-04  8.471200e-0
4
## [10,]  9.301960e-04  1.912655e-03  1.677863e-03  1.512399e-03  2.610141e-0
4
## [11,]  3.633766e-02  8.733144e-04  9.809366e-05 -1.483698e-04  5.225947e-0
4
## [12,]  8.733144e-04  5.412899e-03  4.992183e-04  6.743144e-04  2.732446e-0
4
## [13,]  9.809366e-05  4.992183e-04  3.268872e-02  2.060372e-03  4.147527e-0
5
## [14,] -1.483698e-04  6.743144e-04  2.060372e-03  9.833969e-03  2.266268e-0
4
## [15,]  5.225947e-04  2.732446e-04  4.147527e-05  2.266268e-04  6.676801e-0
3
## [16,]  1.939642e-03  6.592617e-04  1.759742e-04  3.317126e-04  3.699171e-0
4
## [17,]  1.801906e-03  3.416495e-03  6.232322e-04  8.516297e-04  3.668215e-0
4
## [18,]  2.809708e-03  1.003249e-03  2.665368e-04  5.068941e-04  4.840023e-0
4
## [19,]  4.483225e-04  1.013793e-03  1.515043e-04  1.716059e-04  5.103002e-0
7
## [20,]  7.917292e-05  1.308757e-05 -4.907737e-04 -1.814400e-04 -3.508767e-0
5
## [21,]  1.091741e-05  2.033967e-05 -2.145044e-05 -1.822603e-05  2.364441e-0
5
##               [,16]         [,17]         [,18]         [,19]         [,20
]
##  [1,] -2.534889e-04 -2.107689e-04 -2.854221e-04 -1.044473e-04 -3.224048e-0
```

```
3
##  [2,] -9.466475e-05 -1.420793e-03 -3.443819e-04 -4.038132e-04  1.519110e-0
3
##  [3,] -3.438503e-03 -2.482619e-03 -5.221790e-03 -4.690650e-04  9.968029e-0
5
##  [4,] -4.355404e-04 -1.680641e-03 -8.321198e-04 -3.838164e-04  4.506410e-0
4
##  [5,] -1.237594e-03 -8.212494e-03 -2.080635e-03 -2.783608e-03 -5.254879e-0
5
##  [6,]  1.150196e-03  6.551513e-03  1.812416e-03  1.397494e-03  7.781056e-0
5
##  [7,]  5.267812e-04  1.727556e-03  9.014318e-04  4.289011e-04  1.018102e-0
4
##  [8,]  2.571593e-04  1.292867e-03  4.607795e-04  3.637195e-04 -5.978332e-0
4
##  [9,]  9.127086e-04  2.223744e-03  1.347702e-03  5.670579e-04  2.016807e-0
5
## [10,]  7.366314e-04  2.578940e-03  1.129136e-03  5.839357e-04  2.465960e-0
4
## [11,]  1.939642e-03  1.801906e-03  2.809708e-03  4.483225e-04  7.917292e-0
5
## [12,]  6.592617e-04  3.416495e-03  1.003249e-03  1.013793e-03  1.308757e-0
5
## [13,]  1.759742e-04  6.232322e-04  2.665368e-04  1.515043e-04 -4.907737e-0
4
## [14,]  3.317126e-04  8.516297e-04  5.068941e-04  1.716059e-04 -1.814400e-0
4
## [15,]  3.699171e-04  3.668215e-04  4.840023e-04  5.103002e-07 -3.508767e-0
5
## [16,]  1.935920e-03  1.262883e-03  1.709036e-03  2.583599e-04  1.225166e-0
4
## [17,]  1.262883e-03  1.445878e-02  1.909890e-03  1.402884e-03  1.627351e-0
4
## [18,]  1.709036e-03  1.909890e-03  4.511544e-03  3.600099e-04  1.809086e-0
4
## [19,]  2.583599e-04  1.402884e-03  3.600099e-04  7.460997e-03  1.659287e-0
4
## [20,]  1.225166e-04  1.627351e-04  1.809086e-04  1.659287e-04  3.249072e-0
3
## [21,]  1.483741e-05 -6.428915e-06  3.704333e-05 -7.162455e-05  1.669474e-0
5
##               [,21]
##  [1,] -7.437288e-05
##  [2,] -6.306044e-05
##  [3,]  2.688630e-04
##  [4,] -2.318242e-05
##  [5,]  2.229803e-04
##  [6,]  1.070702e-05
##  [7,]  1.114521e-05
##  [8,]  2.948499e-07
```

```
##  [9,]  6.391754e-05
## [10,]  5.886088e-05
## [11,]  1.091741e-05
## [12,]  2.033967e-05
## [13,] -2.145044e-05
## [14,] -1.822603e-05
## [15,]  2.364441e-05
## [16,]  1.483741e-05
## [17,] -6.428915e-06
## [18,]  3.704333e-05
## [19,] -7.162455e-05
## [20,]  1.669474e-05
## [21,]  1.390228e-04

#Model 8b: upper kink, orthogonalized SES, US child, ancestry proportions, in
dividual SIRE categories and 7 multi-SIRE categories with smaller data set -
restriction test

orthostep=lm(general_ses_PCA_z ~ South_Asian + Amerindian + East_Asian + Afri
can
             + frac_Black_SIRE_woc
             + frac_EastAsian_SIRE_woc
             + frac_SouthAsian_SIRE_woc
             + frac_Native_American_SIRE_woc
             + frac_Other_Race_SIRE_woc
             + frac_Hispanic_SIRE_woc
             + NH_Black_White_only
             + NH_SouthAsian_White_only
             + NH_EastAsian_White_only
             + NH_Native_American_White_only
             + H_White_only
             + H_Black_only
             + H_Other_only
             + upper_kink, data=merged_df_all_Model3_spec)
ortho_ses <- residuals(orthostep)

model_8b=lmer(CA_Z_adj ~ South_Asian + Amerindian + East_Asian + African
              + frac_Black_SIRE_woc
              + frac_EastAsian_SIRE_woc
              + frac_SouthAsian_SIRE_woc
              + frac_Native_American_SIRE_woc
              + frac_Other_Race_SIRE_woc
              + frac_Hispanic_SIRE_woc
              + NH_Black_White_only
              + NH_SouthAsian_White_only
              + NH_EastAsian_White_only
              + NH_Native_American_White_only
              + H_White_only
              + H_Black_only
              + H_Other_only
```

```
              + upper_kink
              + Child_US_Born
              + ortho_ses
              + (1|site_id_l) + (1|site_id_l:rel_family_id), data=merged_df_a
ll_Model3_spec, REML = FALSE)
summary(model_8b)
```

```
## Warning in site_id_l:rel_family_id: numerical expression has 9972 elements
: only
## the first used

## Warning in site_id_l:rel_family_id: numerical expression has 9972 elements
: only
## the first used

## Warning in site_id_l:rel_family_id: numerical expression has 9972 elements
: only
## the first used

## Warning in site_id_l:rel_family_id: numerical expression has 9972 elements
: only
## the first used

## Warning in site_id_l:rel_family_id: numerical expression has 9972 elements
: only
## the first used

## Warning in site_id_l:rel_family_id: numerical expression has 9972 elements
: only
## the first used

## Warning in site_id_l:rel_family_id: numerical expression has 9972 elements
: only
## the first used

## Warning in site_id_l:rel_family_id: numerical expression has 9972 elements
: only
## the first used

## Warning in site_id_l:rel_family_id: numerical expression has 9972 elements
: only
## the first used

## Warning in site_id_l:rel_family_id: numerical expression has 9972 elements
: only
## the first used

## Warning in site_id_l:rel_family_id: numerical expression has 9972 elements
: only
```

```
## the first used

## Warning in site_id_l:rel_family_id: numerical expression has 9972 elements
: only
## the first used

## Warning in site_id_l:rel_family_id: numerical expression has 9972 elements
: only
## the first used

## Warning in site_id_l:rel_family_id: numerical expression has 9972 elements
: only
## the first used

## Warning in site_id_l:rel_family_id: numerical expression has 9972 elements
: only
## the first used

## Warning in site_id_l:rel_family_id: numerical expression has 9972 elements
: only
## the first used

## Linear mixed model fit by maximum likelihood  ['lmerMod']
## Formula: CA_Z_adj ~ South_Asian + Amerindian + East_Asian + African +
##     frac_Black_SIRE_woc + frac_EastAsian_SIRE_woc + frac_SouthAsian_SIRE_w
oc +
##     frac_Native_American_SIRE_woc + frac_Other_Race_SIRE_woc +
##     frac_Hispanic_SIRE_woc + NH_Black_White_only + NH_SouthAsian_White_onl
y +
##     NH_EastAsian_White_only + NH_Native_American_White_only +
##     H_White_only + H_Black_only + H_Other_only + upper_kink +
##     Child_US_Born + ortho_ses + (1 | site_id_l) + (1 | site_id_l:rel_famil
y_id)
##    Data: merged_df_all_Model3_spec
##
##      AIC      BIC   logLik deviance df.resid
##  25523.4  25696.4 -12737.7  25475.4     9948
##
## Scaled residuals:
##     Min      1Q  Median      3Q     Max
## -3.9088 -0.5421 -0.0481  0.4970  5.0183
##
## Random effects:
##  Groups                  Name        Variance Std.Dev.
##  site_id_l:rel_family_id (Intercept) 0.318111 0.56401
##  site_id_l               (Intercept) 0.008041 0.08967
##  Residual                            0.454931 0.67449
## Number of obs: 9972, groups:  site_id_l:rel_family_id, 8419; site_id_l, 22
##
## Fixed effects:
```

```
##                                  Estimate Std. Error t value
## (Intercept)                       0.20374    0.06142   3.317
## South_Asian                       0.67434    0.31869   2.116
## Amerindian                       -1.31322    0.11960 -10.981
## East_Asian                        0.73163    0.19712   3.712
## African                          -1.13272    0.12530  -9.040
## frac_Black_SIRE_woc              -0.12356    0.10111  -1.222
## frac_EastAsian_SIRE_woc          -0.35045    0.17758  -1.973
## frac_SouthAsian_SIRE_woc         -0.31474    0.26171  -1.203
## frac_Native_American_SIRE_woc -0.26980    0.13142  -2.053
## frac_Other_Race_SIRE_woc         -0.15932    0.09934  -1.604
## frac_Hispanic_SIRE_woc            0.35837    0.19062   1.880
## NH_Black_White_only               0.07349    0.07355   0.999
## NH_SouthAsian_White_only          0.31835    0.18079   1.761
## NH_EastAsian_White_only          -0.00356    0.09915  -0.036
## NH_Native_American_White_only -0.16176    0.08172  -1.979
## H_White_only                     -0.07995    0.04398  -1.818
## H_Black_only                      0.08074    0.12024   0.671
## H_Other_only                     -0.14922    0.06710  -2.224
## upper_kink                        0.22672    0.08618   2.631
## Child_US_Born                     0.08967    0.05700   1.573
## ortho_ses                         0.27956    0.01179  23.710

##
## Correlation matrix not shown by default, as p = 21 > 12.
## Use print(x, correlation=TRUE)  or
##     vcov(x)        if you need it
```

```
performance::icc(model_8b)
```

```
## # Intraclass Correlation Coefficient
##
##      Adjusted ICC: 0.418
##   Conditional ICC: 0.326
```

```
performance::r2(model_8b)
```

```
## # R2 for Mixed Models
##
##   Conditional R2: 0.546
##      Marginal R2: 0.220
```

```
#For Restriction Test
# the test applies to models 2, 3, 6, 8a and 8b

coeffvec <- coef(model_8b)
varcov <- vcov(model_8b, full=FALSE)
```

```
## Warning in site_id_l:rel_family_id: numerical expression has 9972 elements
: only
## the first used
```

```
## Warning in site_id_l:rel_family_id: numerical expression has 9972 elements
: only
## the first used

## Warning in site_id_l:rel_family_id: numerical expression has 9972 elements
: only
## the first used

## Warning in site_id_l:rel_family_id: numerical expression has 9972 elements
: only
## the first used

## Warning in site_id_l:rel_family_id: numerical expression has 9972 elements
: only
## the first used

## Warning in site_id_l:rel_family_id: numerical expression has 9972 elements
: only
## the first used

## Warning in site_id_l:rel_family_id: numerical expression has 9972 elements
: only
## the first used

## Warning in site_id_l:rel_family_id: numerical expression has 9972 elements
: only
## the first used
```

```
varcov
```

```
## 21 x 21 Matrix of class "dgeMatrix"
##                [,1]          [,2]          [,3]          [,4]          [,5
]
##  [1,]  3.772475e-03 -3.066324e-03 -3.617732e-04 -0.0008286291 -1.791825e-0
4
##  [2,] -3.066324e-03  1.015655e-01  2.784280e-03  0.0117032554  4.897951e-0
3
##  [3,] -3.617732e-04  2.784280e-03  1.430307e-02  0.0009745968  2.110545e-0
3
##  [4,] -8.286291e-04  1.170326e-02  9.745968e-04  0.0388551916  3.372978e-0
3
##  [5,] -1.791825e-04  4.897951e-03  2.110545e-03  0.0033729775  1.570128e-0
2
##  [6,] -7.678252e-05 -2.570437e-03 -1.570895e-03 -0.0024911787 -1.209196e-0
2
##  [7,]  1.044963e-05 -9.846722e-03 -4.296411e-04 -0.0309144280 -3.091805e-0
3
##  [8,]  1.499217e-03 -7.170171e-02 -1.872247e-03 -0.0090864033 -3.632449e-0
3
```

```
##  [9,] -9.769570e-05 -8.369377e-04 -2.295241e-03 -0.0017051427 -3.551091e-0
3
## [10,] -2.614378e-04 -4.567464e-03 -1.435547e-03 -0.0033560937 -4.476570e-0
3
## [11,] -2.558403e-04  3.498370e-04 -5.952221e-03  0.0010723585 -1.602880e-0
3
## [12,] -9.343767e-05 -1.085565e-03 -8.005994e-04 -0.0012839194 -6.177140e-0
3
## [13,]  8.160974e-04 -3.323622e-02 -8.526716e-04 -0.0044172140 -1.588596e-0
3
## [14,]  1.502489e-04 -4.918477e-03 -2.320382e-04 -0.0155498663 -1.365039e-0
3
## [15,] -1.082359e-04  2.982806e-04 -7.487704e-04 -0.0002238102 -2.128224e-0
4
## [16,] -2.451481e-04 -8.894926e-05 -3.467334e-03 -0.0004337868 -1.260940e-0
3
## [17,] -2.143863e-04 -1.423260e-03 -2.470126e-03 -0.0016813946 -8.202382e-0
3
## [18,] -2.656976e-04 -3.273462e-04 -5.293399e-03 -0.0008257773 -2.140153e-0
3
## [19,] -1.448244e-04 -4.311182e-04 -3.298522e-04 -0.0003920790 -2.671035e-0
3
## [20,] -3.216936e-03  1.531259e-03  6.800664e-05  0.0004566741 -8.136141e-0
5
## [21,] -1.514321e-05  3.810320e-05  5.105780e-06  0.0000270577 -1.695209e-0
5
##                [,6]          [,7]          [,8]          [,9]         [,10
]
##  [1,] -7.678252e-05  1.044963e-05  1.499217e-03 -9.769570e-05 -2.614378e-0
4
##  [2,] -2.570437e-03 -9.846722e-03 -7.170171e-02 -8.369377e-04 -4.567464e-0
3
##  [3,] -1.570895e-03 -4.296411e-04 -1.872247e-03 -2.295241e-03 -1.435547e-0
3
##  [4,] -2.491179e-03 -3.091443e-02 -9.086403e-03 -1.705143e-03 -3.356094e-0
3
##  [5,] -1.209196e-02 -3.091805e-03 -3.632449e-03 -3.551091e-03 -4.476570e-0
3
##  [6,]  1.022282e-02  2.431150e-03  2.088259e-03  2.835018e-03  3.616765e-0
3
##  [7,]  2.431150e-03  3.153477e-02  7.927775e-03  1.841943e-03  3.014343e-0
3
##  [8,]  2.088259e-03  7.927775e-03  6.848961e-02  9.369651e-04  3.568707e-0
3
##  [9,]  2.835018e-03  1.841943e-03  9.369651e-04  1.726990e-02  1.355300e-0
3
## [10,]  3.616765e-03  3.014343e-03  3.568707e-03  1.355300e-03  9.869001e-0
3
## [11,]  1.313509e-03 -3.078764e-03 -5.749938e-04 -5.066162e-03  9.255863e-0
4
```

```
## [12,]  4.984849e-03  1.340708e-03  9.903415e-04  1.627080e-03  1.904018e-0
3
## [13,]  9.909783e-04  3.950400e-03  2.373933e-02  4.508924e-04  1.686892e-0
3
## [14,]  1.167702e-03  1.263945e-02  4.034908e-03  8.304157e-04  1.520064e-0
3
## [15,]  3.365233e-04  3.089420e-04 -6.574278e-05  8.451418e-04  2.506675e-0
4
## [16,]  1.149110e-03  5.257295e-04  2.575255e-04  9.070736e-04  7.303046e-0
4
## [17,]  6.551983e-03  1.728010e-03  1.292705e-03  2.226176e-03  2.581682e-0
3
## [18,]  1.809550e-03  8.984305e-04  4.606097e-04  1.330399e-03  1.113463e-0
3
## [19,]  1.402721e-03  4.339394e-04  3.618399e-04  5.939258e-04  6.144897e-0
4
## [20,]  7.626905e-05  9.985063e-05 -5.996618e-04  7.141136e-06  2.397296e-0
4
## [21,] -2.129676e-06 -5.172888e-06 -1.493225e-05 -4.456215e-05  1.682562e-0
6
##               [,11]         [,12]         [,13]         [,14]         [,15
]
##  [1,] -2.558403e-04 -9.343767e-05  8.160974e-04  1.502489e-04 -1.082359e-0
4
##  [2,]  3.498370e-04 -1.085565e-03 -3.323622e-02 -4.918477e-03  2.982806e-0
4
##  [3,] -5.952221e-03 -8.005994e-04 -8.526716e-04 -2.320382e-04 -7.487704e-0
4
##  [4,]  1.072359e-03 -1.283919e-03 -4.417214e-03 -1.554987e-02 -2.238102e-0
4
##  [5,] -1.602880e-03 -6.177140e-03 -1.588596e-03 -1.365039e-03 -2.128224e-0
4
##  [6,]  1.313509e-03  4.984849e-03  9.909783e-04  1.167702e-03  3.365233e-0
4
##  [7,] -3.078764e-03  1.340708e-03  3.950400e-03  1.263945e-02  3.089420e-0
4
##  [8,] -5.749938e-04  9.903415e-04  2.373933e-02  4.034908e-03 -6.574278e-0
5
##  [9,] -5.066162e-03  1.627080e-03  4.508924e-04  8.304157e-04  8.451418e-0
4
## [10,]  9.255863e-04  1.904018e-03  1.686892e-03  1.520064e-03  2.506675e-0
4
## [11,]  3.633681e-02  8.717016e-04  9.974573e-05 -1.469700e-04  5.205304e-0
4
## [12,]  8.717016e-04  5.409955e-03  5.024213e-04  6.770438e-04  2.701994e-0
4
## [13,]  9.974573e-05  5.024213e-04  3.268554e-02  2.057691e-03  4.598853e-0
5
## [14,] -1.469700e-04  6.770438e-04  2.057691e-03  9.831707e-03  2.305663e-0
4
```

```
## [15,]  5.205304e-04  2.701994e-04  4.598853e-05  2.305663e-04  6.678316e-0
3
## [16,]  1.938449e-03  6.571462e-04  1.783790e-04  3.337699e-04  3.681324e-0
4
## [17,]  1.802423e-03  3.417411e-03  6.221893e-04  8.507374e-04  3.675886e-0
4
## [18,]  2.806806e-03  9.978166e-04  2.722259e-04  5.117248e-04  4.775326e-0
4
## [19,]  4.540886e-04  1.023989e-03  1.398633e-04  1.616433e-04  8.917681e-0
6
## [20,]  7.798675e-05  1.039585e-05 -4.887183e-04 -1.797566e-04 -4.125857e-0
5
## [21,]  1.039737e-06 -2.075177e-06 -4.334976e-06 -4.208289e-06 -2.774271e-0
5
##              [,16]         [,17]         [,18]         [,19]         [,20
]
##  [1,] -2.451481e-04 -2.143863e-04 -2.656976e-04 -1.448244e-04 -3.216936e-0
3
##  [2,] -8.894926e-05 -1.423260e-03 -3.273462e-04 -4.311182e-04  1.531259e-0
3
##  [3,] -3.467334e-03 -2.470126e-03 -5.293399e-03 -3.298522e-04  6.800664e-0
5
##  [4,] -4.337868e-04 -1.681395e-03 -8.257773e-04 -3.920790e-04  4.566741e-0
4
##  [5,] -1.260940e-03 -8.202382e-03 -2.140153e-03 -2.671035e-03 -8.136141e-0
5
##  [6,]  1.149110e-03  6.551983e-03  1.809550e-03  1.402721e-03  7.626905e-0
5
##  [7,]  5.257295e-04  1.728010e-03  8.984305e-04  4.339394e-04  9.985063e-0
5
##  [8,]  2.575255e-04  1.292705e-03  4.606097e-04  3.618399e-04 -5.996618e-0
4
##  [9,]  9.070736e-04  2.226176e-03  1.330399e-03  5.939258e-04  7.141136e-0
6
## [10,]  7.303046e-04  2.581682e-03  1.113463e-03  6.144897e-04  2.397296e-0
4
## [11,]  1.938449e-03  1.802423e-03  2.806806e-03  4.540886e-04  7.798675e-0
5
## [12,]  6.571462e-04  3.417411e-03  9.978166e-04  1.023989e-03  1.039585e-0
5
## [13,]  1.783790e-04  6.221893e-04  2.722259e-04  1.398633e-04 -4.887183e-0
4
## [14,]  3.337699e-04  8.507374e-04  5.117248e-04  1.616433e-04 -1.797566e-0
4
## [15,]  3.681324e-04  3.675886e-04  4.775326e-04  8.917681e-06 -4.125857e-0
5
## [16,]  1.934435e-03  1.263525e-03  1.705060e-03  2.655005e-04  1.202903e-0
4
## [17,]  1.263525e-03  1.445851e-02  1.911613e-03  1.399795e-03  1.637036e-0
4
```

```
## [18,]  1.705060e-03  1.911613e-03  4.501679e-03  3.792102e-04  1.765623e-0
4
## [19,]  2.655005e-04  1.399795e-03  3.792102e-04  7.426670e-03  1.768010e-0
4
## [20,]  1.202903e-04  1.637036e-04  1.765623e-04  1.768010e-04  3.249072e-0
3
## [21,] -3.702312e-06  1.635365e-06  8.496086e-07  1.891318e-05  1.669474e-0
5
##               [,21]
##  [1,] -1.514321e-05
##  [2,]  3.810320e-05
##  [3,]  5.105780e-06
##  [4,]  2.705770e-05
##  [5,] -1.695209e-05
##  [6,] -2.129676e-06
##  [7,] -5.172888e-06
##  [8,] -1.493225e-05
##  [9,] -4.456215e-05
## [10,]  1.682562e-06
## [11,]  1.039737e-06
## [12,] -2.075177e-06
## [13,] -4.334976e-06
## [14,] -4.208289e-06
## [15,] -2.774271e-05
## [16,] -3.702312e-06
## [17,]  1.635365e-06
## [18,]  8.496086e-07
## [19,]  1.891318e-05
## [20,]  1.669474e-05
## [21,]  1.390228e-04

# I now repeat the same procedures as for models 7 and 8 but without upper_ki
nk to create models 9a,9b,10a and 10b
# model 9a same as model 7a without upper_kink variable

model_9a=lmer(CA_Z_adj ~ South_Asian + Amerindian + East_Asian + African
              + frac_Black_SIRE
              + frac_EastAsian_SIRE
              + frac_SouthAsian_SIRE
              + frac_Native_American_SIRE
              + frac_Other_SIRE
              + frac_Hispanic_SIRE
              + Child_US_Born
              + general_ses_PCA_z
              + (1|site_id_l) + (1|site_id_l:rel_family_id), data=merged_df,
REML = FALSE)
summary(model_9a)

## Warning in site_id_l:rel_family_id: numerical expression has 9972 elements
: only
```

```
## the first used

## Warning in site_id_l:rel_family_id: numerical expression has 9972 elements
: only
## the first used

## Warning in site_id_l:rel_family_id: numerical expression has 9972 elements
: only
## the first used

## Warning in site_id_l:rel_family_id: numerical expression has 9972 elements
: only
## the first used

## Warning in site_id_l:rel_family_id: numerical expression has 9972 elements
: only
## the first used

## Warning in site_id_l:rel_family_id: numerical expression has 9972 elements
: only
## the first used

## Warning in site_id_l:rel_family_id: numerical expression has 9972 elements
: only
## the first used

## Warning in site_id_l:rel_family_id: numerical expression has 9972 elements
: only
## the first used

## Warning in site_id_l:rel_family_id: numerical expression has 9972 elements
: only
## the first used

## Warning in site_id_l:rel_family_id: numerical expression has 9972 elements
: only
## the first used

## Warning in site_id_l:rel_family_id: numerical expression has 9972 elements
: only
## the first used

## Warning in site_id_l:rel_family_id: numerical expression has 9972 elements
: only
## the first used

## Warning in site_id_l:rel_family_id: numerical expression has 9972 elements
: only
## the first used
```

```
## Warning in site_id_l:rel_family_id: numerical expression has 9972 elements
: only
## the first used

## Warning in site_id_l:rel_family_id: numerical expression has 9972 elements
: only
## the first used

## Warning in site_id_l:rel_family_id: numerical expression has 9972 elements
: only
## the first used

## Linear mixed model fit by maximum likelihood  ['lmerMod']
## Formula: CA_Z_adj ~ South_Asian + Amerindian + East_Asian + African +
##     frac_Black_SIRE + frac_EastAsian_SIRE + frac_SouthAsian_SIRE +
##     frac_Native_American_SIRE + frac_Other_SIRE + frac_Hispanic_SIRE +
##     Child_US_Born + general_ses_PCA_z + (1 | site_id_l) + (1 |
##     site_id_l:rel_family_id)
##    Data: merged_df
##
##      AIC      BIC   logLik deviance df.resid
##  25534.1  25649.5 -12751.1  25502.1     9956
##
## Scaled residuals:
##     Min      1Q  Median      3Q     Max
## -3.9196 -0.5387 -0.0476  0.4944  4.9804
##
## Random effects:
##  Groups                  Name        Variance Std.Dev.
##  site_id_l:rel_family_id (Intercept) 0.321015 0.56658
##  site_id_l               (Intercept) 0.008186 0.09048
##  Residual                            0.454384 0.67408
## Number of obs: 9972, groups:  site_id_l:rel_family_id, 8419; site_id_l, 22
##
## Fixed effects:
##                           Estimate Std. Error t value
## (Intercept)                0.08017    0.06177   1.298
## South_Asian                0.43092    0.31740   1.358
## Amerindian                -0.81910    0.11927  -6.867
## East_Asian                 0.62312    0.19631   3.174
## African                   -0.62242    0.12029  -5.174
## frac_Black_SIRE           -0.09968    0.09810  -1.016
## frac_EastAsian_SIRE       -0.21879    0.17074  -1.281
## frac_SouthAsian_SIRE      -0.11478    0.25286  -0.454
## frac_Native_American_SIRE -0.03491    0.10367  -0.337
## frac_Other_SIRE           -0.07731    0.07598  -1.017
## frac_Hispanic_SIRE        -0.03286    0.08142  -0.404
## Child_US_Born              0.10007    0.05694   1.757
## general_ses_PCA_z          0.28020    0.01177  23.808
```

```
##
## Correlation matrix not shown by default, as p = 13 > 12.
## Use print(x, correlation=TRUE)  or
##     vcov(x)        if you need it
```

```
performance::icc(model_9a)
```

```
## # Intraclass Correlation Coefficient
##
##      Adjusted ICC: 0.420
##   Conditional ICC: 0.329
```

```
performance::r2(model_9a)
```

```
## # R2 for Mixed Models
##
##   Conditional R2: 0.546
##      Marginal R2: 0.218
```

```
# model 9b same as model 7b without upper_kink variable

orthostep = lm(general_ses_PCA_z ~ South_Asian + Amerindian + East_Asian + Af
rican
               + frac_Black_SIRE
               + frac_EastAsian_SIRE
               + frac_SouthAsian_SIRE
               + frac_Native_American_SIRE
               + frac_Other_SIRE
               + frac_Hispanic_SIRE, data=merged_df)
ortho_ses <- residuals(orthostep)


model_9b=lmer(CA_Z_adj ~ South_Asian + Amerindian + East_Asian + African
              + frac_Black_SIRE
              + frac_EastAsian_SIRE
              + frac_SouthAsian_SIRE
              + frac_Native_American_SIRE
              + frac_Other_SIRE
              + frac_Hispanic_SIRE
              + Child_US_Born
              + ortho_ses
              + (1|site_id_l) + (1|site_id_l:rel_family_id), data=merged_df,
REML = FALSE)
summary(model_9b)
```

```
## Warning in site_id_l:rel_family_id: numerical expression has 9972 elements
: only
## the first used

## Warning in site_id_l:rel_family_id: numerical expression has 9972 elements
: only
```

```
## the first used

## Warning in site_id_l:rel_family_id: numerical expression has 9972 elements
: only
## the first used

## Warning in site_id_l:rel_family_id: numerical expression has 9972 elements
: only
## the first used

## Warning in site_id_l:rel_family_id: numerical expression has 9972 elements
: only
## the first used

## Warning in site_id_l:rel_family_id: numerical expression has 9972 elements
: only
## the first used

## Warning in site_id_l:rel_family_id: numerical expression has 9972 elements
: only
## the first used

## Warning in site_id_l:rel_family_id: numerical expression has 9972 elements
: only
## the first used

## Warning in site_id_l:rel_family_id: numerical expression has 9972 elements
: only
## the first used

## Warning in site_id_l:rel_family_id: numerical expression has 9972 elements
: only
## the first used

## Warning in site_id_l:rel_family_id: numerical expression has 9972 elements
: only
## the first used

## Warning in site_id_l:rel_family_id: numerical expression has 9972 elements
: only
## the first used

## Warning in site_id_l:rel_family_id: numerical expression has 9972 elements
: only
## the first used

## Warning in site_id_l:rel_family_id: numerical expression has 9972 elements
: only
## the first used
```

```
## Warning in site_id_l:rel_family_id: numerical expression has 9972 elements
: only
## the first used

## Warning in site_id_l:rel_family_id: numerical expression has 9972 elements
: only
## the first used

## Linear mixed model fit by maximum likelihood  ['lmerMod']
## Formula: CA_Z_adj ~ South_Asian + Amerindian + East_Asian + African +
##     frac_Black_SIRE + frac_EastAsian_SIRE + frac_SouthAsian_SIRE +
##     frac_Native_American_SIRE + frac_Other_SIRE + frac_Hispanic_SIRE +
##     Child_US_Born + ortho_ses + (1 | site_id_l) + (1 | site_id_l:rel_famil
y_id)
##    Data: merged_df
##
##      AIC      BIC   logLik deviance df.resid
##  25534.1  25649.5 -12751.1  25502.1     9956
##
## Scaled residuals:
##     Min      1Q  Median      3Q     Max
## -3.9196 -0.5387 -0.0476  0.4944  4.9804
##
## Random effects:
##  Groups                  Name        Variance Std.Dev.
##  site_id_l:rel_family_id (Intercept) 0.321015 0.56658
##  site_id_l               (Intercept) 0.008186 0.09048
##  Residual                            0.454384 0.67408
## Number of obs: 9972, groups:  site_id_l:rel_family_id, 8419; site_id_l, 22
##
## Fixed effects:
##                           Estimate Std. Error t value
## (Intercept)                0.19822    0.06147   3.225
## South_Asian                0.63433    0.31738   1.999
## Amerindian                -1.34634    0.11709 -11.498
## East_Asian                 0.72811    0.19632   3.709
## African                   -1.04807    0.11909  -8.801
## frac_Black_SIRE           -0.15667    0.09808  -1.597
## frac_EastAsian_SIRE       -0.22341    0.17074  -1.308
## frac_SouthAsian_SIRE      -0.12655    0.25287  -0.500
## frac_Native_American_SIRE -0.25296    0.10364  -2.441
## frac_Other_SIRE           -0.18569    0.07583  -2.449
## frac_Hispanic_SIRE        -0.09937    0.08139  -1.221
## Child_US_Born              0.10007    0.05694   1.757
## ortho_ses                  0.28020    0.01177  23.808

##
## Correlation matrix not shown by default, as p = 13 > 12.
```

```
## Use print(x, correlation=TRUE)  or
##     vcov(x)        if you need it

performance::icc(model_9b)

## # Intraclass Correlation Coefficient
##
##      Adjusted ICC: 0.420
##   Conditional ICC: 0.329

performance::r2(model_9b)

## # R2 for Mixed Models
##
##   Conditional R2: 0.546
##      Marginal R2: 0.218

# model 10a same as 8a without upper_kink variable

model_10a=lmer(CA_Z_adj ~ South_Asian + Amerindian + East_Asian + African
               + frac_Black_SIRE_woc
               + frac_EastAsian_SIRE_woc
               + frac_SouthAsian_SIRE_woc
               + frac_Native_American_SIRE_woc
               + frac_Other_Race_SIRE_woc
               + frac_Hispanic_SIRE_woc
               + NH_Black_White_only
               + NH_SouthAsian_White_only
               + NH_EastAsian_White_only
               + NH_Native_American_White_only
               + H_White_only
               + H_Black_only
               + H_Other_only
               + Child_US_Born
               + general_ses_PCA_z
               + (1|site_id_l) + (1|site_id_l:rel_family_id), data=merged_df_
all_Model3_spec, REML = FALSE)
summary(model_10a)

## Warning in site_id_l:rel_family_id: numerical expression has 9972 elements
: only
## the first used

## Warning in site_id_l:rel_family_id: numerical expression has 9972 elements
: only
## the first used

## Warning in site_id_l:rel_family_id: numerical expression has 9972 elements
: only
## the first used
```

```
## Warning in site_id_l:rel_family_id: numerical expression has 9972 elements
: only
## the first used

## Warning in site_id_l:rel_family_id: numerical expression has 9972 elements
: only
## the first used

## Warning in site_id_l:rel_family_id: numerical expression has 9972 elements
: only
## the first used

## Warning in site_id_l:rel_family_id: numerical expression has 9972 elements
: only
## the first used

## Warning in site_id_l:rel_family_id: numerical expression has 9972 elements
: only
## the first used

## Warning in site_id_l:rel_family_id: numerical expression has 9972 elements
: only
## the first used

## Warning in site_id_l:rel_family_id: numerical expression has 9972 elements
: only
## the first used

## Warning in site_id_l:rel_family_id: numerical expression has 9972 elements
: only
## the first used

## Warning in site_id_l:rel_family_id: numerical expression has 9972 elements
: only
## the first used

## Warning in site_id_l:rel_family_id: numerical expression has 9972 elements
: only
## the first used

## Warning in site_id_l:rel_family_id: numerical expression has 9972 elements
: only
## the first used

## Warning in site_id_l:rel_family_id: numerical expression has 9972 elements
: only
## the first used

## Warning in site_id_l:rel_family_id: numerical expression has 9972 elements
```

```
: only
## the first used

## Linear mixed model fit by maximum likelihood  ['lmerMod']
## Formula: CA_Z_adj ~ South_Asian + Amerindian + East_Asian + African +
##     frac_Black_SIRE_woc + frac_EastAsian_SIRE_woc + frac_SouthAsian_SIRE_w
oc +
##     frac_Native_American_SIRE_woc + frac_Other_Race_SIRE_woc +
##     frac_Hispanic_SIRE_woc + NH_Black_White_only + NH_SouthAsian_White_onl
y +
##     NH_EastAsian_White_only + NH_Native_American_White_only +
##     H_White_only + H_Black_only + H_Other_only + Child_US_Born +
##     general_ses_PCA_z + (1 | site_id_l) + (1 | site_id_l:rel_family_id)
##    Data: merged_df_all_Model3_spec
##
##      AIC      BIC   logLik deviance df.resid
##  25521.7  25687.5 -12737.8  25475.7     9949
##
## Scaled residuals:
##     Min      1Q  Median      3Q     Max
## -3.9086 -0.5418 -0.0480  0.4970  5.0183
##
## Random effects:
##  Groups                  Name        Variance Std.Dev.
##  site_id_l:rel_family_id (Intercept) 0.318174 0.56407
##  site_id_l               (Intercept) 0.008022 0.08957
##  Residual                            0.454899 0.67446
## Number of obs: 9972, groups:  site_id_l:rel_family_id, 8419; site_id_l, 22
##
## Fixed effects:
##                                Estimate Std. Error t value
## (Intercept)                     0.08525    0.06171   1.381
## South_Asian                     0.47344    0.31869   1.486
## Amerindian                     -0.77998    0.12163  -6.413
## East_Asian                      0.63296    0.19707   3.212
## African                        -0.63358    0.12255  -5.170
## frac_Black_SIRE_woc            -0.10612    0.09981  -1.063
## frac_EastAsian_SIRE_woc        -0.32022    0.17752  -1.804
## frac_SouthAsian_SIRE_woc       -0.28634    0.26167  -1.094
## frac_Native_American_SIRE_woc -0.05507    0.13131  -0.419
## frac_Other_Race_SIRE_woc       -0.04784    0.09924  -0.482
## frac_Hispanic_SIRE_woc          0.37558    0.19056   1.971
## NH_Black_White_only             0.11249    0.07263   1.549
## NH_SouthAsian_White_only        0.28301    0.18080   1.565
## NH_EastAsian_White_only        -0.03278    0.09915  -0.331
## NH_Native_American_White_only -0.05843    0.08171  -0.715
## H_White_only                   -0.04421    0.04390  -1.007
## H_Black_only                    0.05614    0.11915   0.471
## H_Other_only                   -0.07858    0.06704  -1.172
```

```
## Child_US_Born                  0.08867    0.05697   1.556
## general_ses_PCA_z              0.27999    0.01176  23.805

##
## Correlation matrix not shown by default, as p = 20 > 12.
## Use print(x, correlation=TRUE)  or
##     vcov(x)        if you need it

performance::icc(model_10a)

## # Intraclass Correlation Coefficient
##
##      Adjusted ICC: 0.418
##   Conditional ICC: 0.326

performance::r2(model_10a)

## # R2 for Mixed Models
##
##   Conditional R2: 0.546
##      Marginal R2: 0.220

# model 10b same as model 8b but without upper_kink variable

orthostep=lm(general_ses_PCA_z ~ South_Asian + Amerindian + East_Asian + Afri
can
             + frac_Black_SIRE_woc
             + frac_EastAsian_SIRE_woc
             + frac_SouthAsian_SIRE_woc
             + frac_Native_American_SIRE_woc
             + frac_Other_Race_SIRE_woc
             + frac_Hispanic_SIRE_woc
             + NH_Black_White_only
             + NH_SouthAsian_White_only
             + NH_EastAsian_White_only
             + NH_Native_American_White_only
             + H_White_only
             + H_Black_only
             + H_Other_only, data=merged_df_all_Model3_spec)
ortho_ses <- residuals(orthostep)

model_10b=lmer(CA_Z_adj ~ South_Asian + Amerindian + East_Asian + African
               + frac_Black_SIRE_woc
               + frac_EastAsian_SIRE_woc
               + frac_SouthAsian_SIRE_woc
               + frac_Native_American_SIRE_woc
               + frac_Other_Race_SIRE_woc
               + frac_Hispanic_SIRE_woc
               + NH_Black_White_only
               + NH_SouthAsian_White_only
               + NH_EastAsian_White_only
```

```
                + NH_Native_American_White_only
                + H_White_only
                + H_Black_only
                + H_Other_only
                + Child_US_Born
                + ortho_ses
                + (1|site_id_l) + (1|site_id_l:rel_family_id), data=merged_df_
all_Model3_spec, REML = FALSE)
summary(model_10b)
```

```
## Warning in site_id_l:rel_family_id: numerical expression has 9972 elements
: only
## the first used

## Warning in site_id_l:rel_family_id: numerical expression has 9972 elements
: only
## the first used

## Warning in site_id_l:rel_family_id: numerical expression has 9972 elements
: only
## the first used

## Warning in site_id_l:rel_family_id: numerical expression has 9972 elements
: only
## the first used

## Warning in site_id_l:rel_family_id: numerical expression has 9972 elements
: only
## the first used

## Warning in site_id_l:rel_family_id: numerical expression has 9972 elements
: only
## the first used

## Warning in site_id_l:rel_family_id: numerical expression has 9972 elements
: only
## the first used

## Warning in site_id_l:rel_family_id: numerical expression has 9972 elements
: only
## the first used

## Warning in site_id_l:rel_family_id: numerical expression has 9972 elements
: only
## the first used

## Warning in site_id_l:rel_family_id: numerical expression has 9972 elements
: only
## the first used
```

```
## Warning in site_id_l:rel_family_id: numerical expression has 9972 elements
: only
## the first used

## Warning in site_id_l:rel_family_id: numerical expression has 9972 elements
: only
## the first used

## Warning in site_id_l:rel_family_id: numerical expression has 9972 elements
: only
## the first used

## Warning in site_id_l:rel_family_id: numerical expression has 9972 elements
: only
## the first used

## Warning in site_id_l:rel_family_id: numerical expression has 9972 elements
: only
## the first used

## Warning in site_id_l:rel_family_id: numerical expression has 9972 elements
: only
## the first used

## Linear mixed model fit by maximum likelihood  ['lmerMod']
## Formula: CA_Z_adj ~ South_Asian + Amerindian + East_Asian + African +
##     frac_Black_SIRE_woc + frac_EastAsian_SIRE_woc + frac_SouthAsian_SIRE_w
oc +
##     frac_Native_American_SIRE_woc + frac_Other_Race_SIRE_woc +
##     frac_Hispanic_SIRE_woc + NH_Black_White_only + NH_SouthAsian_White_onl
y +
##     NH_EastAsian_White_only + NH_Native_American_White_only +
##     H_White_only + H_Black_only + H_Other_only + Child_US_Born +
##     ortho_ses + (1 | site_id_l) + (1 | site_id_l:rel_family_id)
##    Data: merged_df_all_Model3_spec
##
##      AIC      BIC   logLik deviance df.resid
##  25521.7  25687.5 -12737.8  25475.7     9949
##
## Scaled residuals:
##     Min      1Q  Median      3Q     Max
## -3.9086 -0.5418 -0.0480  0.4970  5.0183
##
## Random effects:
##  Groups                  Name        Variance Std.Dev.
##  site_id_l:rel_family_id (Intercept) 0.318174 0.56407
##  site_id_l               (Intercept) 0.008022 0.08957
##  Residual                            0.454899 0.67446
```

```
## Number of obs: 9972, groups:  site_id_l:rel_family_id, 8419; site_id_l, 22
##
## Fixed effects:
##                                Estimate Std. Error t value
## (Intercept)                    0.203865   0.061395   3.321
## South_Asian                    0.689227   0.318660   2.163
## Amerindian                    -1.302745   0.119535 -10.898
## East_Asian                     0.744480   0.197068   3.778
## African                       -1.051815   0.121413  -8.663
## frac_Black_SIRE_woc           -0.166786   0.099791  -1.671
## frac_EastAsian_SIRE_woc       -0.363629   0.177512  -2.048
## frac_SouthAsian_SIRE_woc      -0.326581   0.261677  -1.248
## frac_Native_American_SIRE_woc -0.290241   0.131238  -2.212
## frac_Other_Race_SIRE_woc      -0.178356   0.099088  -1.800
## frac_Hispanic_SIRE_woc         0.345424   0.190553   1.813
## NH_Black_White_only            0.041481   0.072588   0.571
## NH_SouthAsian_White_only       0.313697   0.180788   1.735
## NH_EastAsian_White_only       -0.008765   0.099139  -0.088
## NH_Native_American_White_only -0.162652   0.081722  -1.990
## H_White_only                  -0.087275   0.043874  -1.989
## H_Black_only                   0.037741   0.119144   0.317
## H_Other_only                  -0.160358   0.066951  -2.395
## Child_US_Born                  0.088671   0.056969   1.556
## ortho_ses                      0.279987   0.011762  23.805

##
## Correlation matrix not shown by default, as p = 20 > 12.
## Use print(x, correlation=TRUE)  or
##     vcov(x)        if you need it
```

```
performance::icc(model_10b)
```

```
## # Intraclass Correlation Coefficient
##
##      Adjusted ICC: 0.418
##   Conditional ICC: 0.326
```

```
performance::r2(model_10b)
```

```
## # R2 for Mixed Models
##
##   Conditional R2: 0.546
##      Marginal R2: 0.220
```
